# Pleistocene glaciation advances the younger temporal dimension of species diversification in a major biodiversity hotspot

**DOI:** 10.1101/2023.08.10.552668

**Authors:** Santosh Kumar Rana, Hum Kala Rana, Jacob B. Landis, Tianhui Kuang, Juntong Chen, Hengchang Wang, Tao Deng, Charles C. Davis, Hang Sun

## Abstract

- The expansive Himalayan-Hengduan Mountains (HM) and Qinghai-Tibet Plateau (QTP) form a biodiversity hotspot imperiled by global change. The species diversity, resulting from myriad factors led to plant diversification from 10 million years ago till now. However, despite understanding speciation timing better, a lack of in-depth population-level studies and a dearth of organismal sampling among closely related species and populations leaves an incomplete view of diversification trends and biotic influences.
- We delve into the complex factors influencing diversification through genomic and eco-morphological analysis within the *Stellera chamaejasme* L. complex, known for its significant floral diversity.
- Our results uncover four cryptic species, indicating a more recent and younger diversification period (∼2.67–0.9 Mya), driven by Pleistocene glaciation and a complex set of interacting biotic factors. These factors prompted allopatric speciation and advocated cyclical warming–cooling episodes along latitudinal and altitudinal gradients throughout the Pleistocene.
- The study emphasizes the existence of cryptic species within these mountains, broadening our understanding of species diversification to more recent than previously recognized. This novel perspective may reshape evolutionary paradigms in plant science. Additionally, it also brings to light concerns over future warming’s possible impacts on alpine species distribution and speciation, underlining the urgency for increased conservation efforts.

## Introduction

The Himalayan-Hengduan Mountains (HM) and the Qinghai-Tibet Plateau (QTP) constitute the most iconic and highest mountain system globally (Zhang *et al*., 2016), harboring remarkable biodiversity (Wen *et al*., 2014; Rahbek *et al*., 2019; Spicer *et al*., 2020). Acknowledged as birthplaces of evolutionary branching and crucial centers of plant biodiversity, these regions compete with some tropical rainforests in diversity (Sun et al., 2017). Modern macroevolutionary studies using extensive phylogenies in the HM-QTP region have proposed a history of species diversification starting in the Miocene era (23.03–5.33 million years ago; Mya), heavily influenced by mountainous uplifts and monsoonal occurrences (Fig. **1a**) (Xing & Ree, 2017; Chen *et al*., 2018; Ding *et al*., 2020). However, the intricate patterns of this historical diversification, particularly from roughly 5.3 Mya to the present, are largely uncharted.

**Fig. 1.**
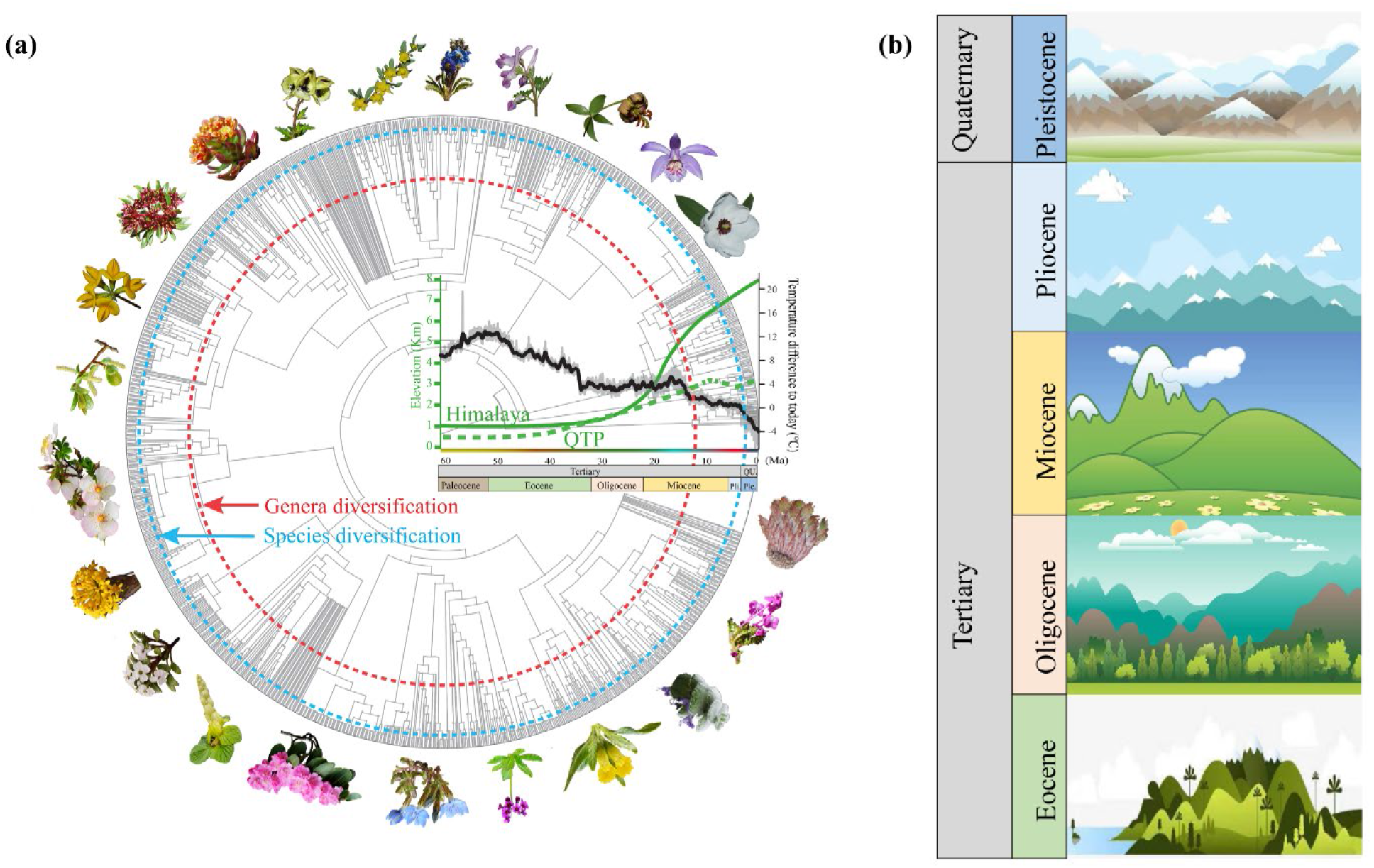
Theoretical framework for plant species diversification in the mountain system. (***a***) The phylogenetic tree was constructed using v.PhyloMaker (Jin & Qian, 2019) with 1520 vascular plant species (representative plant species provided at the edges of the tree) belonging to 235 genera and 56 families from the subnival zone of the Qinghai-Tibet Plateau (QTP) representing genera (red dotted line) and species (cyan dotted line) diversification. The global change in palaeo-temperature (black line; modified from (Westerhold *et al*., 2020) and palaeo-elevation [green solid line for the Himalaya; modified from (Spicer *et al*., 2020) and green dotted line for the QTP; modified from (Zhao *et al*., 2022)] are embedded within the phylogenetic tree (phylogenetic tree not rooted with the geological time period). (***b***) The chronostratigraphic illustrative geological events for the species diversification in the mountain system. The geological events are Pleistocene–the Ice Age, rivers capture and diversified mountain environments; recent diversification of the alpine and temperate species and lineages; Pliocene–the temperature further dropped, and the modern alpine flora formed; Miocene–Asian Moonson formation and intensification, the early diversification of the temperate and alpine genera and species; Oligocene–the plateau further uplifted, the climate gradually cold and dry, and the tropical and subtropical flora replaced by the temperate species; Eocene–the Tethys retreats, the plateau rising, the climate warm and humid, and the tropical and subtropical flora.

The limited understanding of the dynamic evolution of the region partly stems from the phylogenetic sampling of studies (Xing & Ree, 2017; Chen *et al*., 2019; Zhang *et al*., 2021), which predominantly concentrate on broader taxonomic groups rather than densely sampled populations encompassing the entirety of genetic diversity of the taxonomic units involved in these radiations. Such studies may lead to spurious or at least impartial conclusions regarding the true history of the diversification in the region—namely imposing artificially older boundaries that restrict inferences about the historical timing of diversification (Ding *et al*., 2020). We assert that in addition to the previously identified Miocene phenomena, the exclusive exploration of recent glaciation and biotic factors (e.g., pollinator interactions) has driven species diversification in the region. However, the fingerprint of the Pleistocene glaciation and its effect on the biota has remained unexplored. The lack of studies exploring the role of Pleistocene glaciation, in turn, may render the full diversification history incomplete at best, or incorrect at worse, including numerous iconic radiations where primary speciation occurred much later than Pleistocene glaciation [e.g., *Soroseris*, *Syncalathium* (Zhang *et al*., 2011), *Anisodus* (Tu *et al*., 2010), *Saussurea* (Zhang *et al*., 2021), and *Arenaria* (Xu *et al*., 2019)] (Table **1**).

**Table 1.** Representative studies about the cryptic diversity and speciation across the Hengduan Mountains (HM).

| Study species | Species/population | Divergence time (Mya) | Lineage/species diversification (Mya) | Reference |
| --- | --- | --- | --- | --- |
| *Pedicularis siphonantha* complex | 9 species + 12 *Pedicularis* species | 6.04–10.38 | 2.48–5.3 | Liu *et al.*, 2022 |
| *Buddleja alternifolia* | 48 populations | 2.91 | 1.15–1.48 | Ma *et al.*, 2021 |
| *Paeonia* subsect. Delavayanae: Peonies | 2 species (31 *P. delavayi* and 3 *P. ludlowii*) | 10.93 | NA | Zhao *et al.*, 2021 |
| *Roscoea tibetica* | 1 species | 15.4 | NA | Li *et al.*, 2021 |
| Yews (*Taxus wallichiana*) | 43 populations of 1 species | 4.2 | 2.1–2.4 | Liu *et al.*, 2013 |
| White pine (*Pinus armandi*) | 1 species + 6 outgroups species | 9.58 | 4.39–7.36 | Liu *et al.*, 2019 |

The Pleistocene glaciation, around 2.6 Mya (Xing & Ree, 2017), significantly influenced the evolutionary history and diversification in the HM-QTP (Myers *et al*., 2000; Mittermeier *et al*., 2011). Geological processes such as orogeny and monsoons, alongside the uplift of the QTP and the gradual rise of the East Asian monsoon towards the Holocene epoch (Fig. **1b**) (Farnsworth *et al*., 2019), have shaped the macro-diversity of the region (Xing & Ree, 2017; Ding *et al*., 2020). The HM-QTP’s topography, characterized by north-south corridors and east-west isolation (Chen *et al*., 2018), in conjunction with recurrent geographic isolation and river capture events (Sun *et al*., 2022), played a crucial role in driving speciation during the Pleistocene glaciation. These factors, together with glacial history and the “sky islands”, have impacted the evolutionary diversification in the HM-QTP (He *et al*., 2014; Chen *et al*., 2019). Moreover, it is hypothesized that the dynamic glaciation and hydrological changes during the Pleistocene, as well as biotic interactions such as pollinator interactions (Lagomarsiono *et al*., 2016; Weissinger, 2021), herbivory (Niu *et al*., 2014), and human activities (Niu *et al*., 2021), have contributed to the recent diversification of lineages in the region (Eaton *et al*., 2012; Paudel *et al*., 2016). However, the extent to which traits and specific mechanisms like pollen mobility (Wessinger, 2021), the coexistence of differentiated populations (Dynesius & Jansson, 2014; Harvey *et al*., 2017), ecological divergence (Rana *et al*., 2023), and reproductive isolation (Dobzhansky, 1951), have shaped diversity remains poorly understood and necessitates further investigation to unravel their effects on large-scale population-level plant diversification.

*Stellera chamaejasme* L. (Thymelaceae) represents an ideal system for expanding the nuances of diversification due to its broad geographical range, diverse flower color morphs & ecological disparity, high species prevalence, and peculiar evolutionary trajectory (Fig. **2**) (Zhang *et al*., 2010). In the past, it has been classified as two distinct species (Linnaeus, 1753; Léveillé, 1912) or two forms (Huang, 1985), each representing red-white and yellow flower types, respectively. The red-white flower variants, also known as the RW morphs (Fig. **2b**), consist of individuals with red calyx tubes and white lobes and are found widely from the HM-QTP to northeast China and Siberia. Conversely, the purely yellow flower type (hereafter referred to as YY morph; Fig. **2e**), exhibiting a yellow calyx tube and yellow lobes, is located from the southern region of the HM to the Yunnan-Guizhou Plateau (Huang, 1985) (Fig. **2a**). Moreover, we have noticed a pure pink flower type (hereafter referred to as PP morph; Fig. **2c**), with pink calyx tubes and pink lobes, located in Yajiang county of the HM. Additionally, a yellow-white flower variant (hereafter known as YW morph; Fig. **2d**), having a yellow calyx tube and white lobes, is found in the central part of the HM. Given the distinct distribution areas and limited overlap of these four morphs, we are using advanced population genomic techniques to study whether the evolving lineages in the metapopulation introduce ambiguity into the diversification boundaries.

**Fig. 2.**
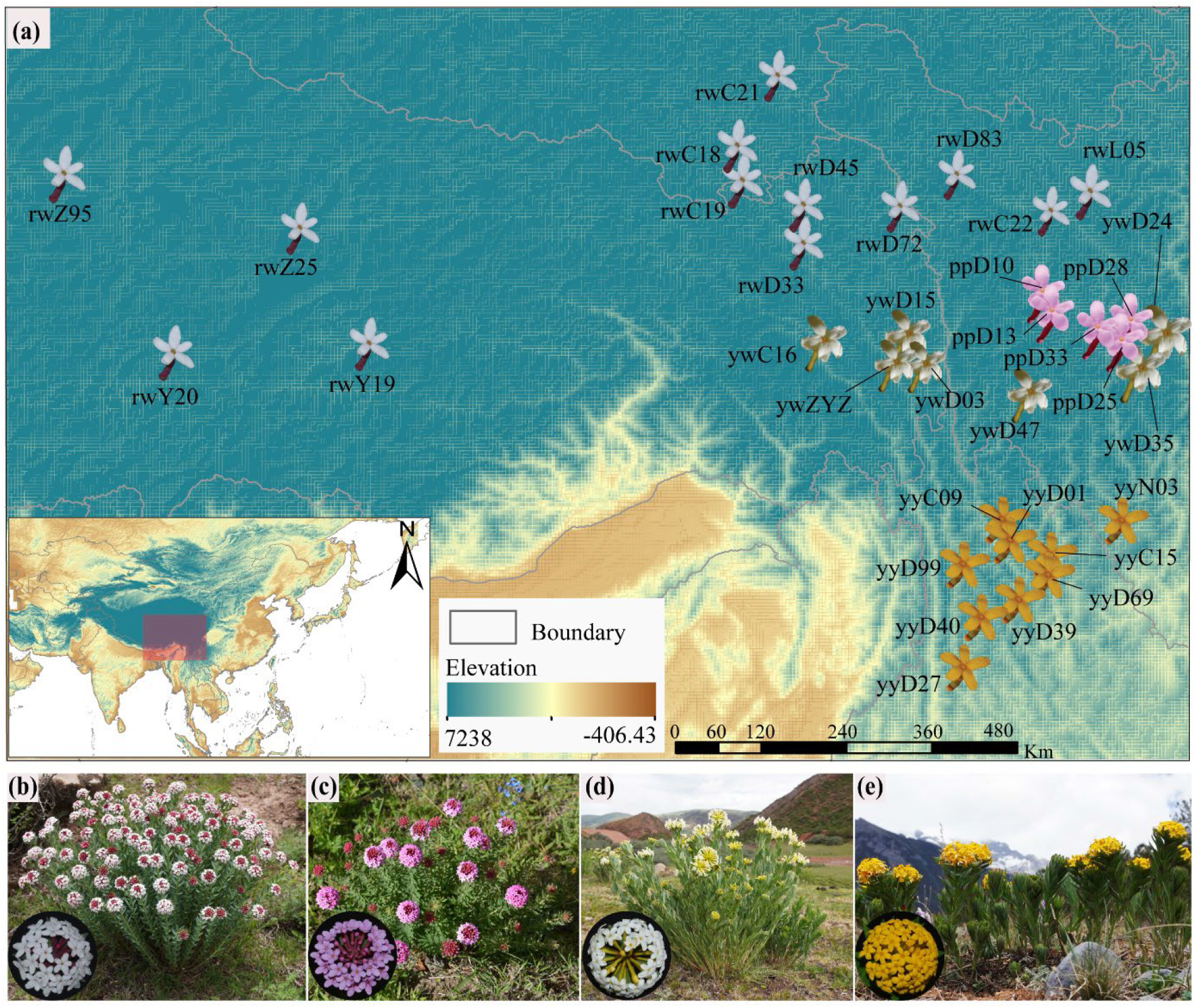
Geographic distribution of populations sampled to infer cryptic speciation scenario in *S. chamaejasme*. (***a***) Sampling localities for 34 populations across the HM-QTP complex (details in Table S2). The four species morphs are (***b***) Red-white morph (RW), (***c***) Pure pink morph (PP), (***d***) Yellow-white morph (YW), and (***e***) Pure yellow morph (YY) in their natural habitats; and were coded same throughout the Figures unless specified for other colors.

This study aims to provide a novel perspective of the diversification paradigm by examining the evolutionary histories of *S. chamaejasme* across different ecological regimes in the HM-QTP. Leveraging population genomics in the study of macroevolutionary dynamics enables us to explore the precise interactions between recent Pleistocene glaciation and ecological phenomena, which is certainly novel. By analyzing genotyped restriction-site association DNA (RAD) loci, we have validated the existence of four independently evolving lineages as cryptic species. This analysis enhances our understanding of the ice age and multiple factors across time and space that drive diversification in this biodiversity hotspot.

## Materials and Methods

### Sampling, RAD sequencing, and data processing

Total genomic DNA was extracted from silica gel-dried leaves of 34 *S. chamaejasme* populations and three outgroups using a Plant Genomic DNA Kit (Tiangen, China). The extracted DNA was used to construct RAD-seq libraries, which were then sequenced on two lanes of the Illumina HiSeq 2000 platform (Illumina, San Diego, CA, USA). The raw sequencing reads will be deposited in NCBI’s Sequence Read Achieve under the accession number. After filtering and processing the clean reads, *de novo* assemblies were generated using Stacks v.2.55 (Catchen *et al*., 2013).

### Genomic screening

Genetic diversity indices (π, *H*_e_, *H*_o_, *F*_IS_) and population genetic differentiation (mean Φ_ST_, *D*_XY_, *F*_ST_’) were calculated using the *populations* pipeline in *Stacks*. The most likely genetic clusters were inferred using Bayesian fastSTRUCTURE (Raj *et al*., 2014) and more with components analysis through DAPC and PCA using the ‘*adegenet’* v2.1.3 (Jombart, 2008). Additionally, shared ancestry was estimated using fineRADstructure v.0.3.2 (Malinsky *et al*., 2018).

By utilizing likelihood and Bayesian approaches with IQ-TREE v.2.1.4_beta (Nguyen *et al*., 2015) and MrBayes v.3.2.7(Ronquist *et al*., 2012), respectively, a concatenated supermatrix was employed to determine the phylogenomic relationships among 251 individuals of *S. chamaejasme*. TreeMix v.1.13 (Pickrell & Pritchard, 2012) was employed to infer historical relationships and gene flow among putative cryptic species. Additionally, the multi-species coalescent model in Beast2 v.2.6.3 (Bouckaert *et al*., 2014) was used to estimate a species tree and divergence times for the cryptic species. Temporal calibration was performed using two fossil calibrations following Zhang *et al*. (2010) for the root age of *S. chamaejasme*. The significance of genetic divergence between the cryptic species of *S. chamaejasme* was assessed using linear models with geographic distance for isolation-by-distance (IBD), implemented with the mantel function in the *‘vegan’* v2.5- 7 (Oksanen *et al*., 2021).

We employed the reversible-jump Bayesian MCMC (rjMCMC) method (Yang & Rannala, 2010; Yang, 2015) using the multi-species coalescent (MSC) model in BPP v.4.4.0 (Rannala & Yang, 2013) to determine cryptic species delimitation and validate taxonomic classifications. Furthermore, Bayes Factor Delimitation (BFD*) method (Grummer *et al*., 2017) was utilized to objectively compare and validate the alternative species delimitation models against the current taxonomy of *S. chamaejasme* in BEAST v2.6.3 (Bouckaert, 2014). Additionally, genomic clustering was performed using unsupervised machine learning (UML) approaches, specifically Random Forest (RF) and t-Distributed Stochastic Neighbor Embedding (t-SNE) (Derkarabetian, 2019).

### Morphological characterization

We analyzed 29 morphological traits (16 continuous and 13 categorical) from 950 herbarium specimens using ridgeplot, performed a multivariate statistical test to assess variation among the putative cryptic species, and visualized significant traits with traitgram in the *‘phytools’* v.0.7-70 (Revell, 2012) R-package using the dated MCMC trees. Morphological traits were further analyzed using principal component analysis (PCA) and Linear discriminant analysis (LDA) to assess the relative contribution of traits to the hypothesized four cryptic species.

### Ecological niche dynamics

The collated species occurrence data from herbaria, field records, and online databases were used for ensemble species distribution modeling (eSDM) using *‘biomod2’* v3.4.12 (Braconnot *et al*., 2007). Models were built using spatially rarefied 307 out of 363 occurrence points and 10000 random pseudo absence points. The potential distribution of four cryptic specie was modeled for the current climatic scenario (c. 1990–2000) and projected to paleoclimatic (last glacial maximum, LGM c. 22 kya) scenario (Thuiller *et al*., 2020) using least correlated predictive variables. The predictive variables were further analyzed using PCA and LDA to evaluate the relative contributions and discriminate the four cryptic species. An ensemble of the ten algorithms was generated using the model with model accuracy≥ 0.80, and the consensus model was converted to a binary model with a 50% probability threshold of suitable habitat that suits the present distribution of the focal species. Niche equivalency testing was performed using the ecospat function in ‘*ENMTools’* v.1.0.3 (Warren & Dinnage, 2021; Warren *et al*., 2021) among the projected niche of *S. chamaejasme*.

See supplementary Methods for more details.

## Results

### Genomic consequences of diversification

A total of 251 individuals representing four distinct cryptic species morphs of *Stellera chamaejasme* (Fig. **2b–e**) were subjected to high-throughput sequencing, resulting in over 3.76 million raw reads on average. After quality control, over 3.47 million clean reads with an effective rate of 93% were used for the assembly of restriction-site associated DNA (RAD) tags (Table **S1**). Subsequent *de novo* assembly and filtration (-p 4 -r 0.8) resulted in a subset of 23,215 RAD loci containing 5,800 single nucleotide polymorphisms (SNPs) for downstream analyses. Additional settings led to the identification of 1,716 loci for Treemix analysis and 25,049 loci for Bayesian phylogeny and phylogeography (BPP). The study detected minimal genetic diversity at the population level (Tables **S2**, **S3**) but observed the highest genetic diversity in the YY morph as determined by *F*_IS_ (0.1327), followed by the PP morph based on *H*_o_ (0.07895), the RW morph based on *He* (0.07756) and π (0.07795) (Table **S2**). Notably, there was a significant level of genetic differentiation within the cryptic species morph, particularly within the Deng8999 population, as indicated by high Φ_ST_ (0.832), *D*_XY_ (0.0058), and *F*_ST_’ (0.824) values (Fig. **S1a** and Tables **S4**, **S5**). However, there was a lack of genetic differentiation observed among populations of the same cryptic species morph (Fig. **S1**). Additionally, the YY morph showed the highest differentiation when compared to the PP morph, RW morph, and YW morph (Table **S6**).

The genetic clustering analysis using fastSTRUCTURE successfully categorized the 34 sampled populations into four distinct ancestral clusters (K=4) (Fig. **3a**, **S2**), consistent with the inferred phylogenetic position, except for Deng9783 (RW morph) which showed a close affinity with the PP morph (Fig. **3c**). Consequently, the populations were assigned to 13 RW, 5 PP, 7 YW, and 9 YY morphs. An analysis with *K*=2 revealed the presence of an admixed population from the YW morph (Fig. **3a**, **S3**). Discriminant analysis of principal components (DAPC) (Fig. **3b**) and principal component analysis (PCA) (Fig. **S4**) confirmed the genetic clusters identified by fastSTRUCTURE with *K*=4. PCA identified two significant principal components (PCs) (Fig. **S4a**, **b**), validated by K-means clustering using the Bayesian information criterion (BIC) curve (Fig. **S4c**, **d**). However, Deng9783 exhibited distinguishable characteristics from the PP morph and was likely misclassified in fastSTRUCTURE due to a high amount of missing data and fewer loci. Bayesian fineRADstructure analysis, including the shared co-ancestry matrix, confirmed the existence of four distinct genetic clusters (*K*=4) (Fig. **S5**) with significant genetic distances (Fig. **S6b**–**f**).

**Fig. 3.**
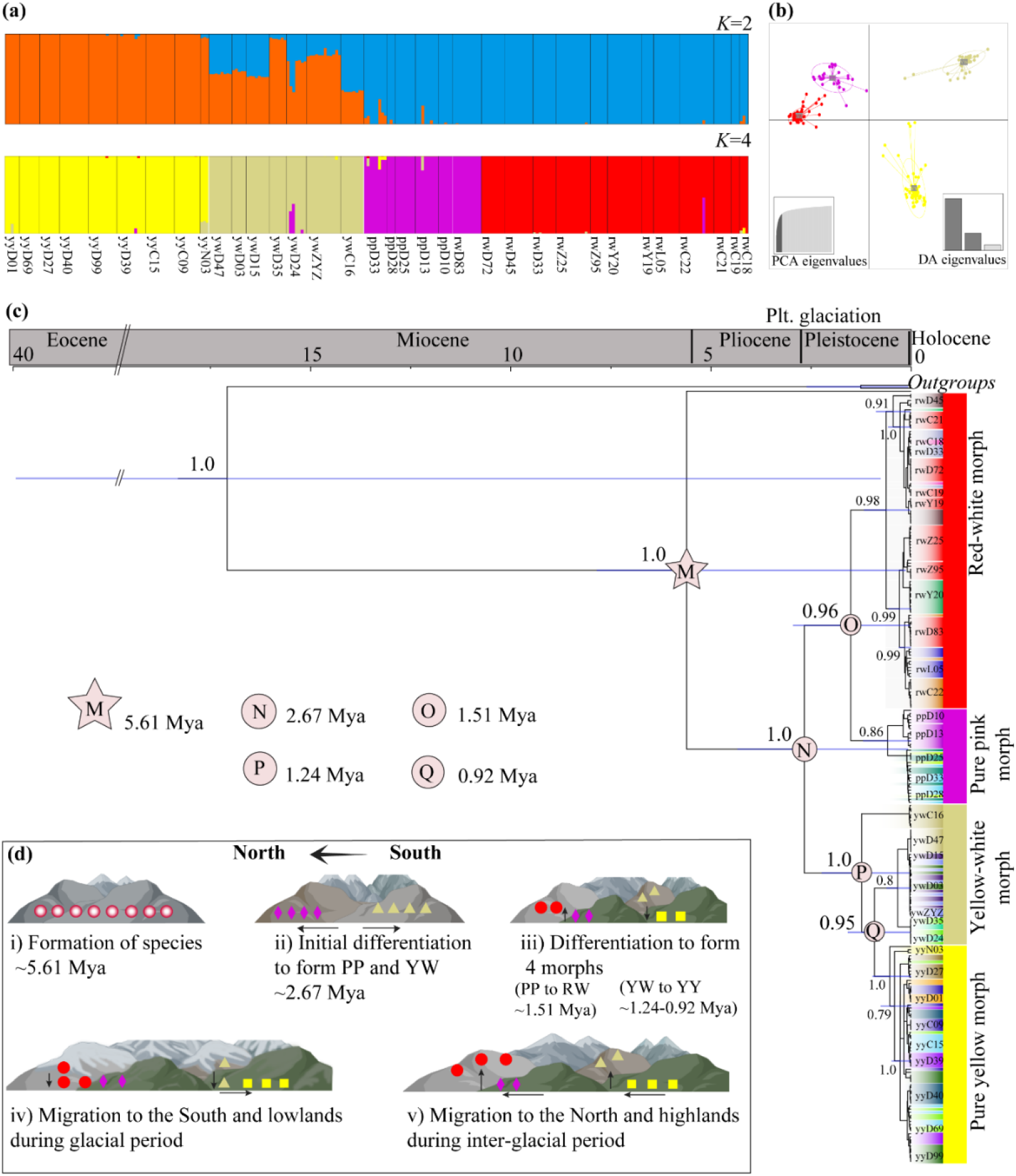
Population divergence of *S. chamaejasme*. (***a***) Hierarchically structured clusters through fastSTRUCTURE analyses under K=2 and K=4. Colors indicate the posterior probability of an assigned individual to a particular cluster. (***b***) Discriminant analysis of principal components displaying the genetic clusters corresponding to four cryptic species morph. (***c***) An estimated divergence time inferred using a Bayesian multi-species coalescent method starBEAST2. The relevant estimated stem age (color star, M) and crown ages (color circles, N–Q) of the cryptic species morph of *S. chamaejasme* are represented in corresponding nodes. Blue bars represent the 95% highest posterior density intervals for node ages, and values nearby nodes indicate posterior probabilities of the major nodes. (***d***) Diagrammatic representation for the temporal dynamics of North-South and up-downslope migration model for diversification of *S. chamaejasme* in the mountain systems. Each species morph is represented by different shape color: PP, pink colored diamond; RW, red colored circle; YW, light yellow colored triangle; YY, yellow colored square.

### Coalescent tree inference and species delimitation

The phylogenomic analysis, conducted using IQ-TREE and MrBayes with the multi-species coalescent model and the best substitution model, consistently identified four major clades representing the four cryptic species morphs (Table **S7**). However, the support values for these clades varied (Figs. **S7–S9**). IQ-TREE showed bootstrap support >86% and SH-aLRT >71 (Figs. **S7**, **S8**) for the RW, PP, YW, and YY morphs, with a notable division observed between the red type (RW and PP morphs) and the yellow type (YW and YY morphs). The PP morph had a more limited geographic distribution compared to the widely distributed RW morph. Interestingly, the HM endemic yellow type appeared as a distinct lineage nested within populations of the YW morph. The phylogenomic tree confirmed the monophyletic nature of the cryptic species morphs, except for the polyphyletic YW morph. It is worth noting that the population Deng9783 formed a major clade with the RW morph, contrary to its clustering with the PP morph in fastSTRUCTURE. The posterior probabilities and rankings of delimited species using BPP varied with different priors for *θ*. Although smaller posterior probabilities were observed (Table **S8**), we focused on the 4- cryptic species morph delimitation model. The 2- and 3-species models had lower posterior probabilities (<0.5) and slightly differed from the 4-species model, with the highest probability of 0.42 under the prior θ ∼ G(2, 0.4) τ ∼ G(2, 0.002) for the 2-species model representing RWPP-YWYY. For the 4-species model, the posterior probability ranged from 0.26 to 0.3 across the range of considered priors (Table **S8**).

To further assess the validity of the previously determined 4-species hypothesis, we utilized BFD* (Bayes Factor Delimitation, *with genomic data) on a dataset (Table **2**) comprising 251 individuals and three closely related taxa (*Daphne longilobata* (Lecomte) Turrill, *Daphne tangutica* Maxim., *Wikstroemia ligustrina* Rehder; not shown). Five different scenarios (listed in Table **2**) were examined, representing distinct partitioning models from the current scenario (single species). BFD results showed increasing likelihood and 2*ln*BF >10 as the number of species increased (Table **2**), supporting a model where individuals are divided into four cryptic species morphs (consistent with clustering and tree estimation under *K*=4). However, the second-ranked 2-species scenarios could also be considered under an admixed model. Convergence of the analysis was confirmed by examining likelihood traces, and the mean effective sample size (ESS) per model was ≥165 (Table **S9**).

**Table 2.** Species delimitation results from BFD* (Bayes Factor Delimitation, *with genomic data) in 251 individuals and three closely related taxa (*Daphne longilobata*, *Daphne tangutica*, *Wikstroemia ligustrina*; not shown) with 5,800 unlinked, bi-allelic SNPs.

| Scenarios | Species delimitation model | marginal L estimate (MLE) | Ln (BF) | 2ln (BF) | Rank |
| --- | --- | --- | --- | --- | --- |
| Null model | Current taxonomy: RWPPYWYY | -110383.59 | - | - | - |
| Alternative 1 | Splitting (Red and yellow morph): 2-species: RWPP-YWYY | -68459.59 | 41924.00 | 83848.00 | 4 |
| Alternative 2 | Splitting (yellow morph): 3-species: RWPP-YW-YY | -32336.66 | 78046.93 | 156093.86 | 2 |
| Alternative 3 | Splitting (red morph): 3-species: RW-PP-YWYY | -36962.92 | 73420.67 | 146841.34 | 3 |
| Alternative 4 | <sup>#</sup>Splitting (red-white, pure red, yellow-white, and pure yellow morph): 4-species: RW-PP-YW-YY | -19020.89 | 91362.70 | 182725.40 | 1 |

Bayes factors [ $2\ln(\text{BF})$ ] were calculated as $2 \times (\text{MLE}_{\text{alternatives}} - \text{MLE}_{\text{null}})$ ; <sup>#</sup> the best-supported models; four cryptic species morphs are RW, Red-white morph; PP, Pure pink morph; YW, Yellow-white morph; YY, Pure yellow morph.

### The tempo of species diversification

The divergence of *S. chamaejasme* from its closely related taxa *W. ligustrina* was estimated to have occurred during the Late Miocene (stem age, c. 5.61 Mya; 95% highest posterior density (HPD) intervals, 0.13–7.85 Mya). Subsequently, within *S. chamaejasme*, the formation of cryptic species morphs began in the Late Pliocene (crown age: approximately 2.67 Mya; 95% HPD: 0.008–4.34 Mya) (Fig. **3c**). Using BEAST2 and SNP datasets, two main clades (red type and yellow type) were identified and further subdivided into four sub-lineages representing the cryptic species morphs (posterior probability ≥ 0.95). The subsequent divergence of the four cryptic species morphs primarily occurred during the Middle Pleistocene (c. 1.51–0.92 Mya). Specifically, the RW and PP morphs diverged around 1.51 Mya (95% HPD: 0.04–2.95 Mya). The YW morph, which is polyphyletic, showed a divergence of one sub-lineage (individuals of Chen0016) at around 1.24 Mya (95% HPD: 0.04–2.24), while the other sub-lineage exhibited nestedness with the YY morph and recent divergence around 0.92 Mya (95% HPD: 0.02–1.59) (Fig. **3c**).

The chord diagram reflects significant genetic differentiation among the cryptic species morphs, with the highest *F*_ST_ observed between the red type-RW/PP and yellow type-YW/YY (Fig. **S1a**). Migration events between the PP and YY morphs were detected, albeit with lesser weight (Fig. **S6a**). However, no apparent gene flow was observed between the other cryptic species morphs. Significant patterns of Isolation-by-distance (IBD) were found, indicating the influence of geographic distance on the genetic structure of *S. chamaejasme*, as supported by a Mantel test (r=0.4785, *P*=0.0001***) (Fig. **S6b**). Within each cryptic species morph, IBD patterns were generally not significant (Fig. **S6c, e**, **f**), except for the PP morph (*r*=0.8875, *P*=0.0167*) (Fig. **S6d**).

### Mode of species diversification: genealogical unsupervised machine learning clustering

The RF and t-SNE run provided more consistent and accurate clustering results (Fig. **4**, **S10**), dividing *S. chamaejasme* into four cryptic species morphs, which align with clustering and tree inference. These machine learning approaches support the discrepancy of the RW morph’s population Deng9783 being grouped with the PP morph by fastSTRUCTURE. However, cMDS (Fig. **4c**) and t-SNE plot (Fig. **4d**) reveal that some individuals of Deng9783 are more closely related to the PP morph. MDS, isoMDS, and t-SNE techniques help resolve the uncertainty regarding the placement of the Deng9783 population within the PP morph. It’s worth noting that MDS and isoMDS exhibited high variability and clustering among the morphs, with fewer exceptions for cMDS. Each clustering technique (Fig. **S10**) and ordination approach (Fig. **4**) demonstrated unique characteristics, but ultimately, the optimal clustering with four clusters (*K*=4) aligns with the four species of *S. chamaejasme*, as confirmed by distance-based *K*-means and PAM clustering (Fig. **S10 a**, **b**) using WSS indices.

**Fig. 4.**
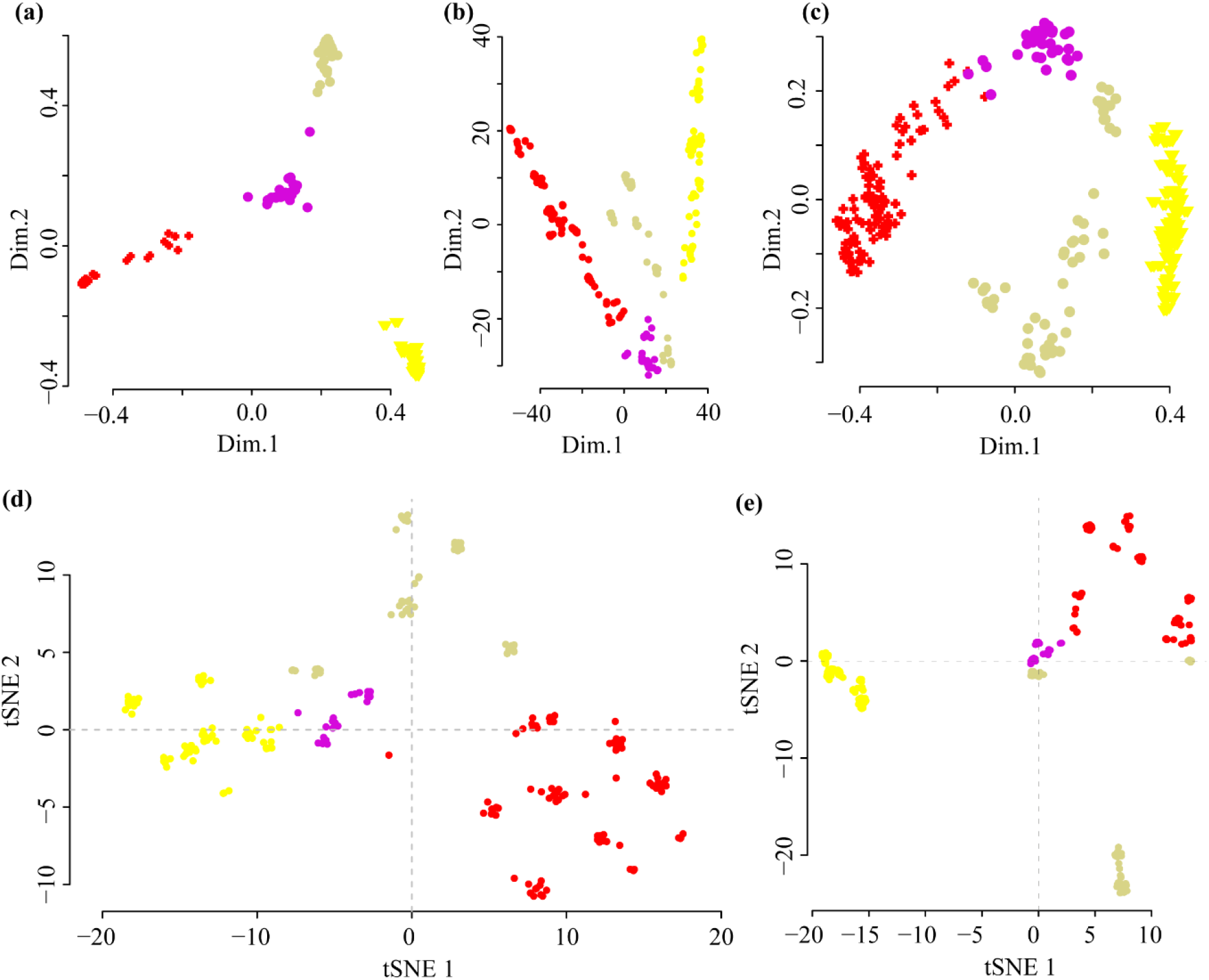
Unsupervised machine learning clustering for the differentiation of cryptic species morph of *S. chamaejasme*. The clustering was identified through (***a***) randomForest multidimensional scaling (MDS) MDSplot function, together with (***b***) isotonic MDS (isoMDS), (***c***) classic MDS (cMDS); and (***d***) Barnes-Hut t-distributed stochastic neighbor embedding (R-tSNE), followed by (***e***) R-tSNE PCA plot.

### Mode of species diversification: morphological basis for the speciation

We analyzed 29 non-normal (*P*<0.05, except PP morph) (Table **S10**) and least correlated (Fig. **S11**) morphological traits (16 continuous, 13 categorical). The continuous traits, such as rhizome length, stem length, leaf length-width ratio, and leaf width base-apex ratio, showed varying densities across the four cryptic species morphs. In contrast, the categorical traits exhibited differing densities, except for leaf apex, midrib appearance, flower number, and bract number (Fig. **S12**). The multivariate analysis of variance (MANOVA) test, assuming homogeneity of variance (Table **S11**), demonstrated the effectiveness of delimiting the cryptic species morphs (Table **S12**) (*F*-stat=17.6, *P*<2.2e-16), although not all traits contributed significantly. Some traits, such as rhizome length, petiole length, bract width, bract length-width ratio, tube diameter, lobe length-width ratio, bract number, and flower number, did not significantly differentiate the four species (at sig. level 0.05) (Table **S12**). Additionally, post-hoc Tukey HSD MANOVA (Table **S13**) identified categorical traits to be more effective in distinguishing the species than continuous traits. For example, leaf shape exhibited greater significance in delimiting all paired species, followed by leaf base, lobe color, and mid-rib color for all paired species except RW and YW (Table **S13**). Among the continuous traits, leaf length-width ratio and leaf width base-apex ratio significantly differentiated the highest number of paired species (*P*<0.05) (Table **S13**).

Based on morphological measurements, traitgrams show overlaps but also reveal differentiation, particularly in the leaf width basal-apex ratio and significantly for leaf shape (Fig. **5a**, **S13**). Ordination analysis using principal component analysis (PCA) on significant traits confirms the distinctiveness of the four species of *S. chamaejasme*, although some traits overlap within certain morphs (Fig. **5b**, **S14a**, **b**). The first two coordinates explain 39.6% of the total variation (Fig. **S14b**). Specifically, the RW morph stands out in terms of leaf width base-apex ratio, flowering season, leaf angle, and stem number, distinguishing it from the other three morphs. The PP morph can only be distinguished based on leaf density. Among the significant traits, except for stem length, bract length, and leaf area, the remaining traits play a more prominent role in differentiating the YW and YY morphs (Fig. **5b**). Linear discriminant analysis (LDA) performed on the significant traits, considering the a priori groups of the four cryptic species morphs, captures 99.87% of the total variation (Fig. **S14c**). The differentiation among morphs primarily relies on variations in categorical traits (Fig. **S15**) rather than the mean values of continuous traits (Fig. **S16**).

**Fig. 5.**
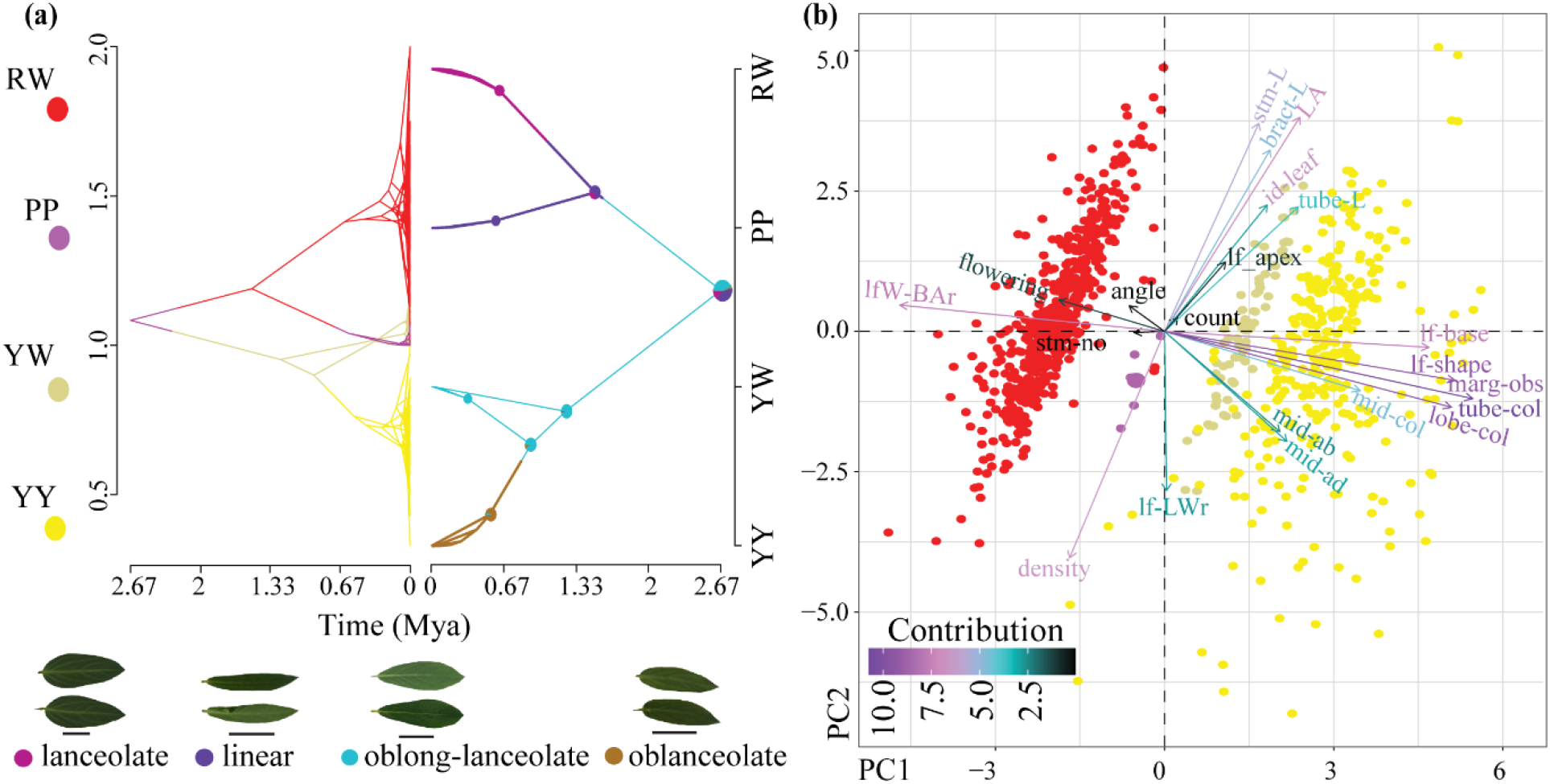
Phenotypic variation of the four cryptic species morphs of *S. chamaejasme*. (***a***) Traitgram depicting the temporal differentiation mapped over dated MCMC tree (time from the root: x-axis) using leaf character (presented as photo plates) categorized as a continuous trait-ratio of leaf width at basal/apex region (lfW_B/Ar) (trait value: y-axes; left side) and categorical trait-leaf shape (species: y-axes; right side). (***b***) The relative contributions of 29 morphological traits to the PCs (PC1, 27.6%; PC2, 12%) according to the clustered cryptic species morph inferred through principal component analysis (PCA) biplots.

### Mode of species diversification: ecological dynamics of speciation

The range of the four cryptic species morphs was determined using the least correlated predictive variables (Table **S14**) from various categories, including bioclimatic, geo-climatic, habitat heterogeneity, ultra-violet radiation, and land cover (Table **S15**, **S16**). However, these ranges are influenced by specific environmental variables associated with each morph. Principal component analysis (PCA) explained 39.6% of the total variation, with PC1 and PC2 accounting for the majority (21.5% and 18.1%, respectively) (Fig. **S17a**). Biplots of the PCA showed that the RW and YY morphs were differentiated by multiple variables, while the PP and YW morphs were primarily influenced by a few specific variables (Fig. **S18a**). The niche assemblage of the RW morph is driven by habitat heterogeneity, followed by geo-climatic, bioclimatic, and land cover factors, whereas the YY morph is characterized by ultra-violet radiation, followed by geo-climatic, land cover, growing days, and bioclimatic factors (Fig. **S18a**). Linear discriminant analysis (LDA) confirmed the grouping of the four morphs based on the predictive environmental variables, with LD1, explaining a significant proportion (61.41%) of the variation (Fig. **S18b**). Common variables that contributed to the niche assemblage of all morphs included bioclimatic factors like isothermality and precipitation seasonality, geo-climatic variables such as elevation and soil pH, habitat heterogeneity measures like the coefficient of variation and correlation, and growing days and land cover characteristics (Fig. **S18a** and Table **S14**).

The eSDM, using various predictive environmental factors and an algorithm (Table **S14**–**S17**), effectively determined the potential distribution and climatic conditions for four cryptic species morphs in the ‘Third Pole’ region, with a focus on the HHM and northeast China, which are major biodiversity hotspots (Fig. **6**). In the eastern range of the HM, the PP and YW morphs of *S. chamaejasme* coexist, while their range overlaps with the RW morph in the northwestern range of the HM (Fig. **2a**). However, the RW morph has a relatively extensive potential distribution across the Himalayas, the QTP, the western part of the HM, and northeast China (Fig. **6**). Analysis of bioclimatic conditions indicated that all morphs have a broader suitable habitat range from the last glacial maximum (LGM, ∼ 22,000 years BP) to the current climatic scenario (c. 1990–2000) (Fig. **S19**). Tests measuring niche similarity showed that the four cryptic species morphs have relatively low similarity (D, I<0.7), indicating distinct suitable environments (Fig. **S20**). Specifically, the RW morph is potentially suited for the Himalayas, the QTP, and northeast China, while the remaining morphs are only suitable for the HM.

**Fig. 6.**
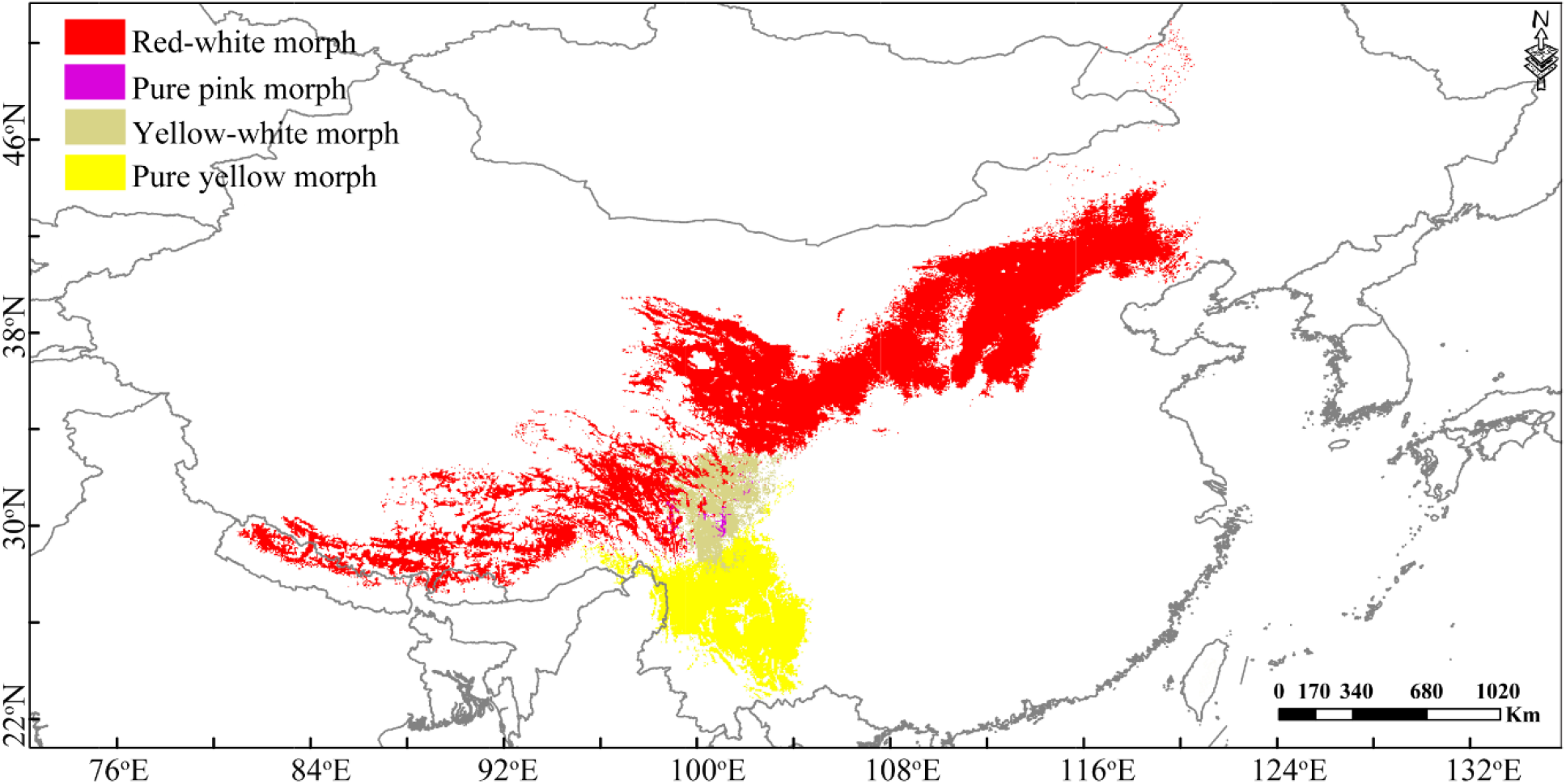
Ecological niche dynamics predicted by ensemble species distribution modeling (eSDM) under current scenarios of environmental variables, as a measure of cryptic species morphs differentiation in *S. chamaejasme*.

## Discussion

### Recent Pleistocene glaciation for species diversification in the polar cap

The Himalayan-Hengduan Mountains and the Qinghai-Tibet Plateau (HM-QTP) complex are globally recognized for their unparalleled species diversity, making it a critical area for examining the organization and distribution of biological communities (Sun *et al*., 2017). Our research aimed to clarify the temporal dynamics of diversification within this complex. Although previous studies hypothesized that the genesis of species within this region was tied to geological upheavals occurring 8–10 million years ago (Mya) (Spicer *et al*., 2003; Deng & Ding, 2015; Ding *et al*., 2020), or ancient monsoonal activities around 5.3 Mya (Chen *et al*., 2018; Ding *et al*., 2020), our investigation into *Stellera chamaejasme* revealed that species diversification likely happened during the more recent Pleistocene glaciation. Extending the timeline for species emergence in the HM-QTP complex allowed us to highlight the profound impact of Pleistocene phenomena on biodiversity. Specifically, we found that *S. chamaejasme* originated during the late Miocene (around 5.61 Mya) and underwent significant diversification during the Pleistocene (around 2.67 Mya). This diversification was primarily driven by the geological alterations associated with the uplift of the HM (Shi *et al*., 1998; Fang *et al*., 2020) (Fig. **3d**). The combination of dynamic geological processes, cyclical environmental changes during the Pleistocene glaciation, facilitated genetic differentiation, leading to the emergence of new species adapted to specific environmental niches (Xing & Ree, 2017).This aligns with the younger time bounds of diversification observed in the HM region (Renner, 2016; Xing & Ree, 2017), which has a distinct history of biodiversity assemblage due to its relatively young age (Luo *et al*., 2016; Sun *et al*., 2017; Xing & Ree, 2017). Our study sheds light on uncharacterized cryptic species, refining the broad time scale of species evolution to a more recent time scale since the Pleistocene epoch. One of the key findings was that one of the four cryptic species morphs, the YY morph, underwent speciation around 0.92 Mya, influenced by the climatic fluctuations of the Pleistocene. Interestingly, this morph showed reduced niche suitability during the Last Glacial Maximum (LGM). During this glaciation period, the YY morph diversified, while the YW population migrated to warmer southern regions and lower altitudes. The glacial history of the HM resulted in the survival of the RW morph, while the PP morph sought refuge in valleys and later migrated northward to higher altitudes. This movement eventually led to the emergence of the RW morph during the interglacial period when the climate warmed up. Consequently, the constrained movement of these species within their geographical range has contributed to their adaptation and survival, resulting in the observed pattern of the four cryptic species morphs (Fig. **3d(v)**). Notably, ice-free pockets of land served as refugia for many alpine plants during glaciation (Pointing, 2014) in the biodiversity hotspot HM (Mittermeier, 2011), expanding our understanding of the timeline of speciation and emphasizing the significant role of Pleistocene phenomena in shaping the region’s evolutionary history. The identification of cryptic species and their ecological adaptations within this temporal framework provides new insights into the dynamics of species diversification in the polar cap.

### North-South and up-downslope migration for the Pleistocene diversification

Explorations of species diversification over time have primarily focused on orogeny and monsoonal events, often overlooking the influence of Pleistocene glaciations on speciation rates (Stanley, 1990; Levsen *et al*., 2012; Xing & Ree, 2017; Ding *et al*., 2020). However, understanding the implications of the Pleistocene epoch, marked by dramatic climatic shifts, is crucial for interpreting recent diversification patterns in mountain ecosystems. The fluctuating Pleistocene climate resulted in substantial reductions in cryptic plant diversity within mountainous regions. Glacial retreat during this time, coupled with decreased precipitation (Farnsworth, 2019) and ice-free temperature conditions (Paillard, 1998; Xing & Ree, 2017), likely spurred plant diversification in the HM region. The HM regions were also extensively covered by ice sheets (Xu *et al*., 2009; Bolch *et al*., 2012) between 0.46 Mya and 0.71 Mya (Jin *et al*., 2005). Yet, we lack a comprehensive understanding of the role the Pleistocene epoch played in current mountain plant diversification. Hypotheses suggest significant diversification occurred due to the impact of Pleistocene glaciation. Diversification during this time appears primarily driven by allopatric speciation, a process involving the geographic separation of populations (Liu *et al*., 2013; Liu *et al*., 2019; Liu *et al*., 2022; Smyčka *et al*., 2022).

Understanding the diversification of four cryptic species morphs necessitates the consideration of a North-South and up-downslope migration model, shaped by fluctuations in climate (Figs. **1**, **3d**). Around 2.67 Mya, climatic fluctuations, and the onset of Pleistocene glaciation likely instigated species diversification. For instance, the transformation of PP into RW can be attributed to a northward and high-altitude migration resulting from warming phenomena at the end of the Xixiabangma Glaciation around 1.51 Mya (Zheng *et al*., 2002). Conversely, the YW to YY transition occurred due to southward and lower-altitude migration, driven by cooling phenomena during the most significant glaciation on the QTP, the Naynayxungla Glaciation (Shi & Ren, 1990; Zhou & Li, 1998). These migration patterns were primarily determined by habitat-specific geographical distribution and have contributed to the differentiation of high-altitude meadow species (RW morph) on mountaintop plateaus from lower-altitude mountain foot valleys (PP morph). A similar distinction exists between high-altitude sub-alpine meadows (or around tree line) as the YW morph and relatively low-altitude southward meadows as the YY morph. As a result, barriers such as the interceding tree line and ecologically distinct niches prevented gene flow among populations with different adaptations, maintaining genetic and morphological distinctiveness.

The north-south-oriented valleys of the HM, unlike the east-west valleys of the Himalayas-QTP, have suggested lower extinction rates and continuous southward migration during Pleistocene glaciation (Xing & Ree, 2017). The development of a major drainage system since the Pleistocene has facilitated the movement of plants from northern or higher regions to southern or lower regions, creating refugia for species (Wang *et al*., 2015; Rana *et al*., 2020; Rana *et al*., 2021; Smyčka *et al*., 2022; Sun *et al*., 2022). This implies that suitable habitats for mountain plants were not drastically reduced but rather displaced southward and downward, resulting in different genetic compositions after the last glacial period and the emergence of the YY morph with similar selection pressures as the YW morph. Moreover, the retreat of glaciers led to reproductive self-incompatibility, contributing to the emergence of four cryptic species morphs under the ice age scenario (Fig. **3d(v)**) (Zhang *et al*., 2015). Some morphs (YW and RW) migrated to lower elevations during glaciation and later returned to higher altitudes, leaving remnant populations at lower elevations during interglacial periods. These remnant populations likely adapted to their new environments, resulting in the formation of new species morphs (YY and PP morphs).

Apart from Pleistocene glaciation, the allopatric speciation within *S. chamaejasme* could have been influenced by orogeny, ancient monsoonal events, and biogeographic barriers such as the Mekong-Salween-Divide (MSD) (Ward, 1921; Luo *et al*., 2017), the 29°N Latitudinal line (Zhang et al., 2009), and the Tsangpo-Brahmaputra Grand-Canyon (TBGC) (Rana *et al*., 2023). Insights from the North-South and up-downslope migration model during glacial-interglacial periods (Li *et al*., 1996; Zheng & Rutter, 1998) illuminate novel Pleistocene diversification and the impact of glaciation together with biotic factors (e.g., pollinator) on speciation in the QTP-HM (Table **1**).

### Allopatric speciation for crypticity and conservation

Our research underlines the pivotal role of allopatric speciation in cryptic species formation and conservation, by focusing on the four distinct metapopulation lineages in *S. chamaejasme*. Despite their morphological similarities, these lineages, which can be considered separate cryptic species, exhibit genetic variation and unique traits. A comprehensive exploration of genomic signatures and significant traits was necessary to delineate these species’ boundaries. We found that genomic evolution and intrinsic genomic incompatibilities played a key role in establishing reproductive isolation (Zhang *et al*., 2011; Dufresnes *et al*., 2021) and fostering diverse evolutionary processes (Struck *et al*., 2018) among these lineages. Their geographical isolation and recent divergence highlight the importance of understanding cryptic species in conservation efforts.

The diversification of the HM-QTP region, typically attributed to geophysical and climatic factors, has been further explored in our study through the lens of ecological interactions, emphasizing their role in shaping biodiversity. We identified biotic factors, especially the role of less-mobile pollinators (Zhang *et al*., 2011; Van der Niet *et al*., 2014; Zhang *et al*., 2021), as instrumental to the diversification and cryptic speciation of *S. chamaejasme*. Reduced gene flow and increased population structure (Zhang *et al*., 2010; Zhang *et al*., 2011; Wessinger, 2021), induced by geographic barriers and the nature of pollinators, are critical to understanding floral macroevolutionary and microevolutionary patterns (Wessinger, 2021).

However, the increased crypticity can lead to an evolutionary slowdown and a loss of phenotypic variation among cryptic species. Overgeneralizing a group of cryptic species as a single entity can distort our understanding of variation (Alizon, 2008) and endanger species conservation. In such cases, “bad taxonomy can kill” (May, 1990), or cryptic species, like the PP morph, can face extinction risks owing to their specific habitat needs and limited distribution. Therefore, acknowledging each cryptic species’ uniqueness and implementing targeted conservation efforts, both in their natural habitats and ex-situ, is fundamental. This approach not only facilitates scientific recognition but also helps assess conservation risks more accurately (Bickford *et al*., 2007; Fišer *et al*., 2018).

In conclusion, our study brings a fresh perspective on species diversification in the biodiversity hotspot, attributing it to recent Pleistocene glaciation and biotic interactions, such as pollinator engagements. Challenging older hypotheses of ancient biodiversity origins, we suggest a more recent emergence of flora diversity in the HM-QTP region, driven largely by allopatric speciation during Pleistocene environmental changes. The uplift of the HM has triggered intricate geological changes, which, in conjunction with environmental fluctuations during the Pleistocene, have played a pivotal role in facilitating genetic differentiation and allopatric speciation. In doing so, our findings challenge previous hypotheses proposing an ancient assemblage of HHM flora (Ding *et al*., 2020) and instead suggest a more recent age for the diversity of HM-QTP mountain plants (Renner, 2016; Sun *et al*., 2017) because of recent glaciation and biotic interactions. The migration model, characterized by movement along a North-South and up-downslope axis, was predominantly driven by allopatric speciation during the Pleistocene’s warming-cooling phenomena.

## Supporting information

Supporting Informations_Tables, Figures, Methods

## Acknowledgments

Thanks to Dr. Zhe Chen for the reflectance measurements and numerous collaborators. This study was funded by the Second Tibetan Plateau Scientific Expedition and Research (STEP) program (2019QZKK0502), the Strategic Priority Research Program of the Chinese Academy of Sciences (XDA20050203), the Key Projects of the Joint Fund of the National Natural Science Foundation of China (U1802232), the Research Fund for International Young Scientists of the National Natural Science Foundation of China (32150410356), the Major Program of the National Natural Science Foundation of China (31590823), the Youth Innovation Promotion Association of Chinese Academy of Sciences (2019382), the Young Academic and Technical Leader Raising Foundation of Yunnan Province (2019HB039), the International Association for Plant Taxonomy (IAPT) Research grant (421064806) and the Chinese Academy of Sciences “Light of West China” Program. The first author is supported by the ‘CAS President’s International Fellowship Initiative’ (PIFI) postdoctoral fellowship (2021PB0034).

## Author contributions

HS, CD, and DT designed research; SKR, HKR, and JC performed research sampling; SKR, HKR, and TK analyzed data; and SKR wrote a paper with the support of HS, CD, TD, JBL, and HW.

## Data Availability

Raw sequence reads obtained by RAD-seq were deposited in the National Center for Biotechnology Information (NCBI) Sequence Read Archive in the Bioproject PRJNA1002599 (https://dataview.ncbi.nlm.nih.gov/object/PRJNA1002599?reviewer=b50aerpftpnsuvhp2bg7v84oiv). All other study materials are included in Supplementary Material online.

The following Supporting Information are available for this article:

## Supplementary Figures

**Fig. S1**. Circos plot showing the measure of genetic variance as indicated by AMOVA-based *F*_ST_ within the paired population level of each cryptic species. (*a*) *Stellera chamaejasme*, (*b*) Red-white morph, (*c*) Pure pink morph, (*d*) Yellow-white morph, and (*e*) Pure yellow morph. Each population is represented by a different gradient of colors.

**Fig. S2**. Model selection for the optimal number of clusters (*K*=4) inferred using fastSTRUCTURE (for *K*=1–10) based on the lowest cross-validation (CV) error (grey color) and the highest marginal likelihood (blue color) estimates.

**Fig. S3**. Hierarchically structured clusters *K*=1–10 identified through fastSTRUCTURE analyses for 34 populations. Colors indicate a posterior probability of assignment of each individual to a particular cluster. The number of clusters *K* for each analysis is displayed on the left side of the bar graph. The cluster for *K*=4 is equivalent to the putative four cryptic species as colored by red – Red-white morph (RW), pink – Pure pink morph (PP), beige – Yellow-white morph (YW), yellow – Pure yellow morph (YY), and coded the same throughout the figures.

**Fig. S4**. Distribution of (*a*) 34 population individuals and (*b*) four morphs of *S. chamaejasme* along principal component (PC) of genetic variation based on single nucleotide polymorphism (SNPs) dataset, along with their eigenvalues plot. Individuals are color-coded according to their population identities (see Fig. 2 for population information). (*c*) The optimal number of clusters *K* based on the Bayesian inference criterion (BIC). The marked (X) indicates the actual number of clusters *K* from where the BIC values decrease slowly on increasing the clusters. (*d*) The optimal number of PCs is retained based on α-score optimization of spline interpolation.

**Fig. S5**. Pairwise co-ancestry heat map depicting variation among individuals of four cryptic species of *S. chamaejasme* inferred through fineRADstructure. The dendrogram shows a clustering of individual samples. Each tip in a tree corresponds to an individual, branches are coded according to clusters. Shared co-ancestry levels are coded from low (yellow) to high (black).

**Fig. S6**. (*a*) Detection of gene flow between the cryptic species morph using TREEMIX. A colored arrow line represents the direction of gene flow. The horizontal scale bar at the bottom (drift parameter) shows the entries’ tenfold average standard error, and the color scale shows the migration weight *w*. Linear regression lines showing the correlations between genetic distance (*F*_ST_, along y-axis) and geographic distances (km, along x-axis) inferred as the function of isolation-by-distance (IBD) for (*b*) *S. chamaejasme*, (*c*) Red-white morph, (*d*) Pure pink morph, (*e*) Yellow-white morph, and (*f*) Pure-yellow morph. Mantel correlation coefficient (r) at a significance level *p* < 0.05 is presented in the figure.

**Fig. S7**. The detailed population-level cryptic species-wise maximum likelihood (ML) phylogenetic tree inferred with IQ-TREE based on the 5,800 SNPs for the Red-white morph (red-colored lineages) and Pure pink morph (pink-colored lineages). Each major node is labeled with SH-aLRT/ultrafast bootstrap support reported as a percentage. The scale bar represents the expected number of substitutions per nucleotide site.

**Fig. S8**. The detailed population-level cryptic species-wise maximum likelihood (ML) phylogenetic tree inferred using IQ-TREE based on the 5,800 SNPs for Yellow-white morph (beige-colored lineages) and Pure yellow morph (yellow-colored lineages). Each major node is labeled with SH-aLRT/ultrafast bootstrap support reported as a percentage. The scale bar represents the expected number of substitutions per nucleotide site.

**Fig. S9**. Bayesian phylogenetic tree inferred with MrBayes analysis based on the 5,800 SNPs of 251 individuals representing four cryptic species of *S. chamaejasme* and three closely related taxa (*Wikstroemia ligustrina*, *Daphne longlobalata*, *Daphne tangutica*). Each major node is labeled with a posterior probability value in the tree. The scale bar represents the expected number of substitutions per nucleotide site.

**Fig. S10**. Genetic distance based unsupervised machine learning (UML) clustering analysis inferred through (*a*) *K*-means clustering and (*b*) Partitioning Around Medoids (PAM) clustering based on within-cluster-sum of squared errors (WSS) index for optimal numbers of clusters *K*. The clusters are represented in two dimensions (Dim1, 36.6%, and Dim2, 10%).

**Fig. S11**. Correlation heatmap of the cryptic species, confirming the least pair correlations across 29 least correlated morphological traits. The figure with blue color indicates positive correlations, and the dark red color indicates negative correlations. Abbreviations for all morphological traits are shown in Supplementary Table S10.

**Fig. S12**. The ridgeplots (*a*–*a3*) visualization for the comparison of all the morphological traits between the four cryptic species of *S. chamaejasme*. The gradient color indicates the trait values provided in the scale. Abbreviations for all morphological traits are shown in **Supplementary** Table S10.

**Fig. S13**. Phylogenetic traitgrams for the significant morphological traits, i.e., (*a*–*j*) continuous traits and (*k*–*t*) categorical traits with strong phylogenetic signals constructed using the phytools. The vertical placement of each tip corresponds to the mean trait values of those individuals for the continuous trait and that of cryptic species for categorical traits. Color in the lineages represents the cryptic species (coded as the same color throughout the figures) for continuous traits, whereas the trait class is represented by color for categorical traits. Abbreviations for all morphological traits are shown in Supplementary Table S10.

**Fig. S14**. (*a*) Top ten contributing morphological traits under the two principal components (blue, PC1; green, PC2) analysis scores. The red dashed line indicates the expected average contribution if variable contributions were uniform. (*b*) The scree plot represents the variance explained by the first ten principal components for morphological traits. Abbreviations for all morphological traits are shown in *SI Appendix*, Table S10. (*c*) The discriminated clusters of the cryptic species into two major components (LD1, 98.89%; LD2, 0.98%) inferred through linear discriminant analysis (LDA) plots of morphological traits. Ellipses are the normal contour lines of each cryptic species with a probability of 90%.

**Fig. S15**. Violin plot (*a*–*j*) to visualize means with standard deviations and the distribution of each significant categorical trait of different cryptic species. Cryptic species are coded in the same color throughout the figures. Abbreviations for all morphological traits are shown in Supplementary Table S10.

**Fig. S16**. Violin plot (*a*–*j*) to visualize mean with standard deviations and the distribution of each significant continuous trait of different cryptic species. Cryptic species are coded in the same color throughout the figures. Abbreviations for all morphological traits are shown in Supplementary Table S10.

**Fig. S17**. Ecological characterization for *Stellera chamaejasme*. (*a*) PCA clustering of all the predictive environmental variables for four cryptic species into two PCs (PC1, 21.5%; PC2, 18.1%). The relative contributions of predictive environmental variables to the PCs for the cryptic species – (*b*) Red-white morph, (*c*) Pure pink morph, (*d*) Yellow-white morph, and (*e*) Pure yellow morph inferred through principal component analysis (PCA) biplots. Abbreviations for all environmental variables are shown in Supplementary Table S14.

**Fig. S18**. Ecological niche analysis as a measure of cryptic species differentiation in *S. chamaejasme*. (*a*) The relative contributions (Cont.) of predictive environmental variables to the PCs (PC1, 21.5%; PC2, 18.1%) inferred through PCA biplots and (*b*) discriminated clusters of the cryptic species into two components (LD1, 61.41%; LD2, 21.15%) inferred through LDA plots. Abbreviations for all environmental variables are shown in Supplementary Table S14.

**Fig. S19**. Habitat suitability predicted by ensemble species distribution modeling (eSDM) for the cryptic species– (*a*, *e*) Red-white morph, (*b*, *f*) Pure pink morph, (*c*, *g*) Yellow-white morph, and (*d*, *h*) Pure yellow morph under the (*a*–*d*) current bioclimatic (c. 1970–2000) and (*e*–*h*) paleoclimatic (last glacial maximum, LGM c. 22 kya) scenario. The LGM scenario is based on the multi-model median ensemble of three global climate models (GCMs), viz. CCSM4, MIROC-ESM, MPI-ESM-P (Braconnot *et al*., 2007).

**Fig. S20**. Niche equivalency test. Pairwise niche (*a*) identity test and (*b*, *c*) background test (*b*, symmetric; *c*, asymmetric) between the four cryptic species morphs through ENMTools. The identity and background test as cryptic species-wise pair comparisons were measured by Schoener’s *D* (in red color) and Hellinger’s-based *I* (in blue color) indices (provided in the respective figure).

## Supplementary Tables

**Table S1**. RAD-seq reads information and quality control statistics for every 251 individuals of *S. chamaejasme* and 3 closely related outgroups.

**Table S2**. Population summary statistics calculated for variant positions of 23,215 restrictions site-associated DNA sequencing (RAD-seq) loci.

**Table S3**. Population summary statistics calculated for all positions of 23,215 restrictions site-associated DNA sequencing (RAD-seq) loci.

**Table S4**. The paired population-level estimation of genetic diversity indices, Φ_ST_ means (lower left diagonal) and *F*_ST_’ means (upper right diagonal) for 34 populations estimated using the *populations* pipeline in Stacks. (*Excel sheet provided for Supplementary Table S4 separately)

**Table S5**. The paired mean absolute differentiation (*D*_XY_) among 34 populations estimated using the *populations* **pipeline** in Stacks. (*Excel sheet provided for Supplementary Table S**5** separately)

**Table S6**. The population-level cryptic species-wise estimation of the genetic diversity indices; Φ_ST_ means, *D*_XY_ means, and *F*_ST_’ means.

**Table S7**. Comparison among the nucleotides substitution model for the selection of best fitting model based on Akaike information criterion (AIC), Bayesian information criterion (BIC), and corrected AIC (AICc) measurements using Modeltest-NG.

**Table S8**. Impact of the prior specification on the number of delimited cryptic species using BPP analysis (A11 = joint species delimitation and species tree estimation). Posterior probability values (*P*) are averaged over two runs.

**Table S9**. Marginal likelihood estimates under different scenarios (N, A1, A2, A3, A4) from Path Sampler analysis for BFD* (Bayes Factor Delimitation, *with genomic data) in 251 individuals and three closely related taxa (*Daphne tangutica*, *Daphne holosericea*, *Wikstroemia ligustrina*).

**Table S10**. Descriptive statistics (mean ± SE) and their Shapiro-Wilk normality test for the normal distribution of the continuous and categorical morphological traits for four cryptic species of *S. chamaejasme* observed from the herbarium specimens.

**Table S11**. Levene’s test of homogeneity of variance assumptions for the morphological traits of *S. chamaejasme*.

**Table S12**. MANOVA test for the significant difference between four cryptic species of *S. chamaejasme* for continuous morphological traits and the trait-wise test of significant difference between the cryptic species.

**Table S13**. Post-hoc TukeyHSD MANOVA test for the trait-species wise significance difference between the four cryptic species of *S. chamaejasme*.

**Table S14**. Predictive variable selection for the ensemble species distribution modeling (eSDM) of each cryptic species of *S. chamaejasme*.

**Table S15**. Variance Inflation Factor (VIF) in different test runs to select explanatory variables for four cryptic species of *S. chamaejasme*. VIF<10 (bold text) are the selected predictive variables.

**Table S16**. Consensus land-cover selection for each cryptic species of *S. chamaejasme* based on the presence of the number of occurrence records.

**Table S17**. Model evaluation indices for the ensemble species distribution modeling (eSDM) of each cryptic species of *S. chamaejasme* using Biomod2 in R-programming language.

## Supplementary Methods

## References

Alizon S, Kucera M, Jansen VAA. 2008. Competition between cryptic species explains variation in rates of lineage evolution. Proceedings of the National Academy of Sciences of the United States of America 105: 12382–12386.

Bickford D, Lohman DJ, Sodhi NS, Ng PKL, Meier R, Winker K, Ingram KK, Das D. 2007. Cryptic species as a window on diversity and conservation. Trends in Ecology and Evolution 22: 148–155.

Bolch T, Kulkarni A, Kääb A, Huggel C, Paul F, Cogley JG, Frey H, Kargel JS, Fujita K, Scheel M et al. 2012. The state and fate of Himalaya. Science 336: 310–314.

Bouckaert R, Heled J, Kühnert D, Vaughan T, Wu CH, Xie D, Suchard MA, Rambaut A, Drummond AJ. 2014. BEAST2: a software platform for Bayesian evolutionary analysis. PLoS Computational Biology 10: e1003537.

Braconnot P, Otto-Bliesner B, Harrison S, Joussaume S, Peterchmitt JY, Abe-Ouchi A, Crucifix M, Driesschaert E, Fichefet Th, Hewitt CD et al. 2007. Results of PMIP2 coupled simulations of the MidHolocene and Last Glacial Maximum–Part1: experiments and large-scale features. Climate of the Past 3: 261–277.

Catchen J, Hohenlohe PA, Bassham S, Amores A, Cresko WA. 2013. Stacks: an analysis tool set for population genomics. Molecular Ecology 22: 3124–3140.

Chen JH, Huang Y, Brachi B, Yun QZ, Zhang W, Lu W, Li HN, Li WQ, Sun XD, Wang GY et al. 2019. Genome-wide analysis of Cushion willow provides insights into alpine plant divergence in a biodiversity hotspot. Nature Communication 10: 5230.

Chen YS, Deng T, Zhou Z, Sun H. 2018. Is the East Asian flora ancient or not? National Science Review 5: 920–932.

Luo D, Yue JP, Sun WG, Xu B, Li ZM, Comes HP, Sun H. 2016. Evolutionary history of the subnival flora of the Himalaya-Hengduan Mountains: first insights from comparative phylogeography of four perennial herbs. Journal of Biogeography 43: 31–43.

de Queiroz K. 2007. Species concepts and species delimitation. Systematic Biology 56: 879–886.

Deng T, Ding L. 2015. Paleoaltimetry reconstructions of the Tibetan Plateau: Progress and contradictions. National Science Review 2: 417–437.

Derkarabetian S, Castillo S, Koo PK, Ovchinnikov S, Hedin M. 2019. A demonstration of unsupervised machine learning in species delimitation. Molecular Phylogenetics and Evolution 139: 106562.

Ding WN, Ree RH, Spicer RA, Xing YW. 2020. Ancient orogenic and monsoon-driven assembly of the world’s richest temperate alpine flora. Science 369: 578–581.

Dobzhansky T. 1951. Genetics and the Origin of Species, third edition. New York: Columbia University Press.

Dufresnes C, Brelsford A, Jeffries DL, Mazepa G, Suchan T, Canestrelli D, Nicieza A, Fumagalli L, Dubey S, Martínez-Solano I et al. 2021. Mass of genes rather than master genes underlie the genomic architecture of amphibian speciation. Proceedings of the National Academy of Sciences of the United States of America 118: e2103963118.

Dynesius M, Jansson R. 2014. Persistence of within-species lineages: A neglected control of speciation rates. Evolution 68: 923–934.

Eaton DAR, Fenster CB, Hereford J, Huang SQ, Ree RH. 2012. Floral diversity and community structure in *Pedicularis* (Orobanchaceae). Ecology 93: S182–S194.

Fang XM, Dupont-Nivet G, Wang CS, Song CH, Meng QQ, Zhang WL, Nie JS, Zhang T, Mao ZQ, Chen Y, 2020. Revised chronology of central Tibet uplift (Lunpola Basin). Science Advances 6: eaba7298.

Farnsworth A, Lunt DJ, Robinson SA, Valdes PJ, Roberts WHG, Clift PD, Markwick P, Su T, Wrobel N, Bragg F et al. 2019. Past East Asian monsoon evolution controlled by paleogeography, not CO_2_. Science Advances 5: eaax1697.

Fišer C, Robinson CT, Malard F. 2018. Cryptic species as a window into the paradigm shift of the species concept. Molecular Ecology 27: 613–635.

Grummer JA, Bryson RW, Reeder TW. 2014. Species delimitation using Bayes factors: Simulations and application to the *Sceloporus sc*alaris species group (Squamata: Phrynosomatidae). Systematic Biology 63: 119–133.

Hu H, Yang Y, Li A, Zheng Z, Zhang J, Liu J. 2022. Genomic divergence of *Stellera chamaejasme* through local selection across the Qinghai-Tibet Plateau and northern China. Molecular Ecology 31: 4782–4796.

Harvey MG, Seeholzer GF, Smith BT, Brumfield RT. 2017. Positive association between population genetic differentiation and speciation rates in New World birds. Proceedings of the National Academy of Sciences of the United States of America 114: 6328–6333.

He K, Jiang XL. 2014. Sky islands of southwest China. I: An overview of phylogeographic patterns. Chinese Science Bulletin 59: 585–597.

Huang SC. 1985. Taxa nova Thymelaeacearum Sinicarum. Acta Botanica Yunnan 7: 277–291.

Jin L, Ganopolski A, Chen F, Claussen M, Wang H. 2005. Impacts of snow and glaciers over Tibetan Plateau on Holocene climate change: sensitivity experiments with a coupled model of intermediate complexity. Geophysical Research Letters 32: L17709.

Jin Y, Qian H. 2019. V.PhyloMaker: an R package that can generate very large phylogenies for vascular plants. Ecography 42: 1353–1359.

Jombart T. 2008. adegenet: an R package for multivariate analysis of genetic markers. Bioinformatics 27: 3070–3071.

Li L, Zhang J, Lu ZQ, Zhao JL, Li QJ. 2021. Genomic data reveal two distinct species from the widespread alpine ginger *Roscoea tibetica* Batalin (Zingiberaceae). Journal of Systematics and Evolution 59, 1232–1243 (2021).

Lagomarsiono LP, Condamine FL, Antonelli A, Mulch A, Davis CC. 2016. The abiotic and biotic drivers of rapid diversification in Andean bellflowers (Campanulaceae). New Phytologist 210: 1430–1442.

Léveillé H. 1912. Decades plantarum novarum. Repertorium Specierum Novarum Regni Vegetabilis 10: 581.

Levsen ND, Tiffin P, Olson MS. 2012. Pleistocene speciation in the genus *Populus* (Salicaceae). Systematic Biology 61: 401–412.

Li J, Feng Z, Zhou S. 1996. The remains of glaciations in Hengduan Mountains. In: Li J, eds. Glaciers in the Hengduan Mountains. Beijing: Science Press, 157–172.

Linnaeus C. 1753.Species Plantarum. Stockholm.

Liu J, Möller M, Provan J, Gao LM, Poudel RC, Li DZ. 2013. Geological and ecological factors drive cryptic speciation of yews in a biodiversity hotspot. New Phytologist 199: 1093–1108.

Liu R, Wang H, Yang JB, Corlett RT, Randle CP, Li DZ, Yu WB. 2022. Cryptic species diversification of the *Pedicularis siphonantha* Complex (Orobanchaceae) in the Mountains of Southwest China since the Pliocene. Frontiers in Plant Science 13: 811206.

Liu Y, Jin W, Wei X, Wang X. 2019. Cryptic speciation in the Chinese white pine (*Pinus armandii*): Implications for the high species diversity of conifers in the Hengduan Mountains, a global biodiversity hotspot. Molecular Phylogenetics and Evolution 138: 114–125.

Luo D, Xu B, Li ZM, Sun H. 2017. The ‘Ward Line–Mekong–Salween divide’ is an important floristic boundary between the eastern Himalaya and Hengduan mountains: evidence from the phylogeographical structure of subnival herbs *Marmoritis complanatum* (Lamiaceae). Botanical Journal of the Linnean Society 185: 482–496.

Ma YP, Wariss HM, Liao RL, Zhang RG, Yun QZ, Olmstead RG, Chau JH, Milne RI, Van de Peer Y, Sun WB. 2021. Genome-wide analysis of butterfly bush (*Buddleja alternifolia*) in three uplands provides insights into biogeography, demography and speciation. New Phytologist 232: 1463–1476.

Malinsky M, Trucchi E, Lawson DJ, Falush D. 2018. RADpainter and fineRADstructure: Population inference from RADseq data. Molecular Biology and Evolution 35: 1284–1290.

May RM. 1990. Taxonomy as destiny. Nature 347: 129–130.

Mittermeier RA, Turner WA, Larsen FW, Brooks TM, Gascon C. 2011. Global biodiversity conservation: the critical role of hotspots. In: Zachos F, Habel J, eds. Biodiversity hotspots. Heidelberg, Berlin: Springer, 3–22.

Myers N, Mittermeier RA, Mittermeier CG, da Fonseca GA, J Kent. 2000. Biodiversity hotspots for conservation priorities. Nature 403: 853–858.

Nguyen LT, Schmidt HA, von Haeseler A, Minh BQ. 2015. IQ-TREE: a fast and effective stochastic algorithm for estimating maximum likelihood phylogenies. Molecular Biology and Evolution 33: 268–274.

Niu Y, Chen G, Peng DL, Song B, Yang Y, Li ZM, Sun H. 2014. Grey leaves in an alpine plant: a cryptic colouration to avoid attack? New Phytologist 203: 953–963.

Niu Y, Stevens M, Sun H. 2021. Commercial harvesting has driven the evolution of camouflage in an Alpine plant. Current Biology 31: 446–449.

Oksanen J, Blanchet FG, Friendly M, Kindt R, Legendre P, McGlinn D, Minchin PR, O’Hara RB, Simpson GL, Solymos P et al. 2021. Vegan: Community Ecology Package. R package Version 2.5–7. URL https://CRAN.R-project.org/package=vegan. [accessed 2 March 2021]

Paillard D. 1998. The timing of Pleistocene glaciations from a simple multiple-state climate model. Nature 391: 378–381.

Paudel BR, Shrestha M, Burd M, Adhikari S, Sun YS, Li QJ. 2016. Coevolutionary elaboration of pollination-related traits in an alpine ginger (*Roscoea purpurea*) and a tabanid fly in the Nepalese Himalayas. New Phytologist 211: 1402–1411.

Pickrell JK, Pritchard JK. 2012. Inference of population splits and mixtures from genome-wide allele frequency data. PLoS Genetics 8: e1002967.

Pointing SB, Bollard-Breen B, Gillman LN. 2014. Diverse cryptic refuges for life during glaciation. Proceedings of the National Academy of Sciences of the United States of America 111: 5452–5453.

Rahbek C, Borregaard MK, Antonelli A, Colwell RK, Holt BG, Nogues-Bravo D, Rasmussen CMØ, Richardson K, Rosing MT, Whittaker RJ et al. 2019. Building mountain biodiversity: Geological and evolutionary processes. Science 365: 1114–1119.

Raj A, Stephens M, Pritchard JK. 2014. fastSTRUCTURE: Variational inference of population structure in large SNP data sets. Genetics 197: 573–589.

Rana HK, Luo D, Rana SK, Sun H. 2020. Geological and climatic factors affect the population genetic connectivity in *Mirabilis himalaica* (Nyctaginaceae): insight from phylogeography and dispersal corridors in the Himalaya-Hengduan biodiversity hotspot. Frontiers in Plant Science 10: 1721.

Rana HK, Rana SK, Luo D, Sun H. 2023. Existence of biogeographic barriers for the long-term Neogene-Quaternary divergence and differentiation of *Koenigia forrestii* in the Himalaya-Hengduan mountains. Botanical Journal of the Linnean Society 201: 230–253.

Rana SK, Luo D, Rana HK, O’Neill AR, Sun H. 2021. Geoclimatic factors influence the population genetic connectivity of *Incarvillea arguta* (Bignoniaceae) in the Himalaya– Hengduan Mountains biodiversity hotspot. Journal of Systematics and Evolution 59: 151–168.

Rannala B, Yang Z. 2013. Improved reversible jump algorithms for Bayesian species delimitation. Genetics 194: 245–253.

Renner SS. 2016. Available data point to a 4-km-high Tibetan Plateau by 40 Ma but 100 molecular-clock papers have linked supposed recent uplift to young node ages. Journal of Biogeography 43: 1479–1487.

Revell LJ. 2012. phytools: An R package for phylogenetic comparative biology (and other things). Methods in Ecology and Evolution 3: 217–223.

Ronquist F, Teslenko M, van der Mark P, Ayres DL, Darling A, Höhna S, Larget B, Liu L, Suchard MA, Huelsenbeck JP. 2012. MrBayes 3.2: Efficient Bayesian phylogenetic inference and model choice across a large model space. Systematic Biology 61: 539–542.

Schluter D, Conte GL. 2009. Genetics and ecological speciation. Proceedings of the National Academy of Sciences of the United States of America 106: 9955–9962.

Shi YF, Li JJ, Li BY. 1998. Uplift and environmental changes of Qinghai-Tibetan Plateau in the Late Cenozoic. Guangzhou, China: Guangdong Science and Technology Press.

Shi YF, Ren JW. 1990. Glacier recession and lake shrinkage indicating a climatic warming and drying trend in central Asia. Annals of Glaciology 14: 261–265.

Smyčka J, Roquet C, Boleda M, Alberti A, Boyer F, Douzet R, Perrier C, Rome M, Valay JG, Denoeud F et al. 2022. Tempo and drivers of plant diversification in the European mountain system. Nature Communication 13: 2750.

Spicer RA, Harris NBW, Widdowson M, Herman AB, Guo SX, Valdes PJ, Wolfe JA, Kelley SP. 2003. Constant elevation of southern Tibet over the past 15 million years. Nature 421: 622– 624.

Spicer RA, Su T, Valdes PJ, Farnsworth A, Wu FX, Shi G, Spicer TEV, Zhou Z. 2020. Why ‘the uplift of the Tibetan Plateau’ is a myth? National Science Review 8(1): nwaa091.

Stanley SM. 1990. In causes of evolution: a paleontological perspective. In: Ross RM, Allmon WD, eds. Chicago: University of Chicago Press, 103–127.

Struck TH, Feder JL, Bendiksby M, Birkeland S, Cerca J, Gusarov VI, Kistenich S, Larsson KH, Liow LH, Nowak MD et al. 2018. Finding evolutionary process hidden in cryptic species. Trends in Ecology and Evolution 33: 153–163.

Sun H, Li Z, Landis JB, Qian L, Zhang T, Deng T. 2022. Effects of drainage reorganization on phytogeographic pattern in Sino-Himalaya. Alpine Botany 132: 141–151.

Sun H, Zhang J, Deng T, Boufford DE. 2017. Origin and evolution of plant diversity in the Hengduan Mountains, China. Plant Diversity 39: 161–166.

Thuiller W, Georges D, Engler R, Breiner F. 2020. Biomod2: Ensemble platform for species distribution modeling. R package version 3.4.12. URL https://cran.r-project.org/web/packages/biomod2/index.html. [accessed 3 November 2020]

Tu T, Volis S, Dillon MO, Sun H, Wen J. 2010. Dispersals of Hyoscyameae and Mandragoreae (Solanaceae) from the New World to Eurasia in the early Miocene and their biogeographic diversification within Eurasia. Molecular Phylogenetics and Evolution 57: 1226–1237.

Van der Niet T, Peakall R, Johnson SD. 2014. Pollinator-driven ecological speciation in plants: new evidence and future perspectives. Annals of Botany 113: 199–211.

Wang ZW, Chen ST, Nie ZL, Zhang JW, Zhou Z, Deng T, Sun H. 2015. Climatic factors drive population divergence and demography: insights based on the phylogeography of a riparian plant species endemic to the Hengduan Mountains and adjacent regions. PloS ONE 10: e0145014.

Ward FK. 1921. The Mekong-Salween divide as a geographical barrier. The Geographical Journal 58: 49–56.

Warren DL, Dinnage R. 2021. ENMTools: Analysis of niche evolution using niche and distribution models. R package version 1.0.3. URL https://cran.r-project.org/web/packages/ENMTools/index.html. [accessed 2 June 2021]

Warren DL, Matzke NJ, Cardillo M, Baumgartner JB, Beaumont LJ, Turelli M, Glor RE, Huron NA, Simões M, Iglesias TL et al. 2021. ENMTools 1.0: an R package for comparative ecological biogeography. Ecography 44: 504–511.

Wen J, Zhang JQ, Nie ZL, Zhong Y, Sun H. 2014. Evolutionary diversifications of plants on the Qinghai Tibetan Plateau. Frontiers in Genetics 5: 4.

Wessinger CA. 2021. From pollen dispersal to plant diversification: genetic consequences of pollination mode. New Phytologist 229: 3125–3132.

Westerhold T, Marwan N, Drury AJ, Liebrand D, Agnini C, Anagnostou E, Barnet JSK, Bohaty SM, de vleeschouwer D, Florindo F et al. 2020. An astronomically dated record of Earth’s climate and its predictability over the last 66 million years. Science 369(6509): 1383– 1387.

Xing Y, Ree RH. 2017. Uplift-driven diversification in the Hengduan Mountains, a temperate biodiversity hotspot. Proceedings of the National Academy of Sciences of the United States of America 114: E3444–E3451.

Xu B, Luo D, Li ZM, Sun H. 2019. Evolutionary radiations of cushion plants on the Qinghai-Tibet Plateau: Insights from molecular phylogenetic analysis of two subgenera of *Arenaria* and *Thylacospermum* (Caryophyllaceae). Taxon 68: 1003–1020.

Xu J, Grumbine RE, Shrestha A, Eriksson M, Yang XF, Wang Y, Wilkes A. 2009. The melting Himalayas: Cascading effects of climate change on water, biodiversity, and livelihoods. Conservation Biology 23: 520–530.

Yang Z, Rannala B. 2010. Bayesian species delimitation using multilocus Proceedings of the National Academy of Sciences of the United States of America 107: 9264–9269.

Yang Z. 2015. The BPP program for species tree estimation and species delimitation. Current Zoology 61: 854–865.

Zhang B, Sun SF, Luo WL, Li JX, Fang QE, Zhang DG, Hu GX. 2021. A new brood-pollination mutualism between *Stellera chamaejasme* and flower thrips *Frankliniella intonsa*. BMC Plant Biology 21: 562.

Zhang D, Ye J, Sun H. 2016. Quantitative approaches to identify floristic units and centres of species endemism in the Qinghai-Tibetan Plateau, south-western China. Journal of Biogeography 43: 2465–2476.

Zhang DC, Boufford DE, Ree RH, Sun H. 2009. The 29°N latitudinal line: an important division in the Hengduan Mountains, a biodiversity hotspot in southwest China. Nordic Journal of Botany 27: 405–412.

Zhang JW, Nie ZL, Wen J, Sun H. 2011. Molecular phylogeny and biogeography of three closely related genera, *Soroseris*, *Stebbinsia*, and *Syncalathium* (Asteraceae, Cichorieae), endemic to the Tibetan Plateau, SW China. Taxon 60: 15–26.

Zhang X, Landis JB, Sun Y, Zhang H, Lin N, Kuang T, Huang X, Deng T, Wang H, Sun H. 2021. Macroevolutionary pattern of *Saussurea* (Asteraceae) provides insights into the drivers of radiating diversification. Proceedings of the Royal Society B: Biological Sciences 288: 20211575.

Zhang Y, Volis S, Sun H. 2010. Chloroplast phylogeny and phylogeography of *Stellera chamaejasme* on the Qinghai-Tibet Plateau and in adjacent regions. Molecular Phylogenetics and Evolution 57: 1162–1172.

Zhang Z, Zhang Y, Sun H. 2011. The reproductive biology of *Stellera chamaejasme* (Thymelaeaceae): A self-incompatible weed with specialized flowers. Flora 206: 567–574.

Zhao JG, Li S, Farnsworth A, Valdes PJ, Reichgelt T, Chen L, Zhou Z, Su T. 2022. The Paleogene to Neogene climate evolution and driving factors on the Qinghai-Tibetan Plateau. Science China Earth Sciences 65(7): 1339–1352.

Zhao Y, Yin G, Pan Y, Tian B, Gong X. 2021. Climatic refugia and geographical isolation contribute to the speciation and genetic divergence in Himalayan-Hengduan tree Peonies (*Paeonia delavayi* and *Paeonia ludlowii*). Frontiers in Genetics 11: 595334.

Zheng B, Rutter N. 1998. On the problem of Quaternary glaciations, and the extent and patterns of Pleistocene ice cover in the Qinghai-Xizang (Tibet) Plateau. Quaternary International 45/46: 109–122.

Zheng B, Xu Q, Shen Y. 2002. The relationship between climate change and Quaternary glacial cycles on the Qinghai–Tibetan Plateau: review and speculation. Quaternary International 97: 93–101.

Zhou SZ, Li JJ. 1998. The sequence of Quaternary glaciation in the Bayan Har Mountains. Quaternary International 45–46: 135–142.

