## Supporting Informations_Tables, Figures, Methods for "Pleistocene glaciation advances the younger temporal dimension of species diversification in a major biodiversity hotspot"

Article title:

Article acceptance date: Click here to enter a date.

The following Supporting Information is available for this article:

**Supplementary Figures**

**Fig. S1**. Circos plot showing the measure of genetic variance as indicated by AMOVA-based $F_{ST}$ within the paired population level of each cryptic species. (*a*) *Stellera chamaejasme*, (*b*) Red-white morph, (*c*) Pure pink morph, (*d*) Yellow-white morph, and (*e*) Pure yellow morph. Each population is represented by a different gradient of colors.

**Fig. S2**. Model selection for the optimal number of clusters (*K*=4) inferred using fastSTRUCTURE (for *K*=1–10) based on the lowest cross-validation (CV) error (grey color) and the highest marginal likelihood (blue color) estimates.

**Fig. S3**. Hierarchically structured clusters *K*=1–10 identified through fastSTRUCTURE analyses for 34 populations. Colors indicate a posterior probability of assignment of each individual to a particular cluster. The number of clusters *K* for each analysis is displayed on the left side of the bar graph. The cluster for *K*=4 is equivalent to the putative four cryptic species as colored by red – Red-white morph (RW), pink – Pure pink morph (PP), beige – Yellow-white morph (YW), yellow – Pure yellow morph (YY), and coded the same throughout the figures.

**Fig. S4**. Distribution of (*a*) 34 population individuals and (*b*) four morphs of *S. chamaejasme* along principal component (PC) of genetic variation based on single nucleotide polymorphism (SNPs) dataset, along with their eigenvalues plot. Individuals are color-coded according to their population identities (see Fig. 2 for population information). (*c*) The optimal number of clusters *K* based on the Bayesian inference criterion (BIC). The marked (X) indicates the actual number of clusters *K* from where the BIC values decrease slowly on increasing the clusters. (*d*) The optimal number of PCs is retained based on α-score optimization of spline interpolation.

**Fig. S5**. Pairwise co-ancestry heat map depicting variation among individuals of four cryptic species of *S. chamaejasme* inferred through fineRADstructure. The dendrogram shows a clustering of individual samples. Each tip in a tree corresponds to an individual, branches are coded according to clusters. Shared co-ancestry levels are coded from low (yellow) to high (black).

**Fig. S6**. (*a*) Detection of gene flow between the cryptic species morph using TREEMIX. A colored arrow line represents the direction of gene flow. The horizontal scale bar at the bottom (drift parameter) shows the entries' tenfold average standard error, and the color scale shows the migration weight *w*. Linear regression lines showing the correlations between genetic distance ($F_{ST}$, along y-axis) and geographic distances (km, along x-axis) inferred as the function of isolation-by-distance (IBD) for (*b*) *S. chamaejasme*, (*c*) Red-white morph, (*d*) Pure pink morph, (*e*) Yellow-white morph, and (*f*) Pure-yellow morph. Mantel correlation coefficient (r) at a significance level $p < 0.05$ is presented in the figure.

**Fig. S7**. The detailed population-level cryptic species-wise maximum likelihood (ML) phylogenetic tree inferred with IQ-TREE based on the 5,800 SNPs for the Red-white morph (red-colored lineages) and Pure pink morph (pink-colored lineages). Each major node is labeled with SH-aLRT/ultrafast bootstrap support reported as a percentage. The scale bar represents the expected number of substitutions per nucleotide site.

**Fig. S8**. The detailed population-level cryptic species-wise maximum likelihood (ML) phylogenetic tree inferred using IQ-TREE based on the 5,800 SNPs for Yellow-white morph (beige-colored lineages) and Pure yellow morph (yellow-colored lineages). Each major node is labeled with SH-aLRT/ultrafast bootstrap support reported as a percentage. The scale bar represents the expected number of substitutions per nucleotide site.

**Fig. S9**. Bayesian phylogenetic tree inferred with MrBayes analysis based on the 5,800 SNPs of 251 individuals representing four cryptic species of *S. chamaejasme* and three closely related taxa (*Wikstroemia ligustrina*, *Daphne longlobalata*, *Daphne tangutica*). Each major node is labeled with a posterior probability value in the tree. The scale bar represents the expected number of substitutions per nucleotide site.

**Fig. S10**. Genetic distance based unsupervised machine learning (UML) clustering analysis inferred through (*a*) *K*-means clustering and (*b*) Partitioning Around Medoids (PAM) clustering based on within-cluster-sum of squared errors (WSS) index for optimal numbers of clusters *K*. The clusters are represented in two dimensions (Dim1, 36.6%, and Dim2, 10%).

**Fig. S11**. Correlation heatmap of the cryptic species, confirming the least pair correlations across 29 least correlated morphological traits. The figure with blue color indicates positive correlations, and the dark red color indicates negative correlations. Abbreviations for all morphological traits are shown in Supplementary Table S10.

**Fig. S12**. The ridgeplots (*a–a3*) visualization for the comparison of all the morphological traits between the four cryptic species of *S. chamaejasme*. The gradient color indicates the trait values provided in the scale. Abbreviations for all morphological traits are shown in Supplementary Table S10.

**Fig. S13**. Phylogenetic traitgrams for the significant morphological traits, i.e., (*a–j*) continuous traits and (*k–t*) categorical traits with strong phylogenetic signals constructed using the phytools. The vertical placement of each tip corresponds to the mean trait values of those individuals for the continuous trait and that of cryptic species for categorical traits. Color in the lineages represents the cryptic species (coded as the same color throughout the figures) for continuous traits, whereas the trait class is represented by color for categorical traits. Abbreviations for all morphological traits are shown in Supplementary Table S10.

**Fig. S14**. (*a*) Top ten contributing morphological traits under the two principal components (blue, PC1; green, PC2) analysis scores. The red dashed line indicates the expected average contribution if variable contributions were uniform. (*b*) The scree plot represents the variance explained by the first ten principal components for morphological traits. Abbreviations for all morphological traits are shown in *SI Appendix*, Table S10. (*c*) The discriminated clusters of the cryptic species into two major components (LD1, 98.89%; LD2, 0.98%) inferred through linear discriminant analysis (LDA) plots of morphological traits. Ellipses are the normal contour lines of each cryptic species with a probability of 90%.

**Fig. S15**. Violin plot (*a–j*) to visualize means with standard deviations and the distribution of each significant categorical trait of different cryptic species. Cryptic species are coded in the same color throughout the figures. Abbreviations for all morphological traits are shown in Supplementary Table S10.

**Fig. S16**. Violin plot (*a–j*) to visualize mean with standard deviations and the distribution of each significant continuous trait of different cryptic species. Cryptic species are coded in the same color throughout the figures. Abbreviations for all morphological traits are shown in Supplementary Table S10.

**Fig. S17**. Ecological characterization for *Stellera chamaejasme*. (*a*) PCA clustering of all the predictive environmental variables for four cryptic species into two PCs (PC1, 21.5%; PC2, 18.1%). The relative contributions of predictive environmental variables to the PCs for the cryptic species – (*b*) Red-white morph, (*c*) Pure pink morph, (*d*) Yellow-white morph, and (*e*) Pure yellow morph inferred through principal component analysis (PCA) biplots. Abbreviations for all environmental variables are shown in Supplementary Table S14.

**Fig. S18**. Ecological niche analysis as a measure of cryptic species differentiation in *S. chamaejasme*. (*a*) The relative contributions (Cont.) of predictive environmental variables to the PCs (PC1, 21.5%; PC2, 18.1%) inferred through PCA biplots and (*b*) discriminated clusters of the cryptic species into two components (LD1, 61.41%; LD2, 21.15%) inferred through LDA plots. Abbreviations for all environmental variables are shown in Supplementary Table S14.

**Fig. S19**. Habitat suitability predicted by ensemble species distribution modeling (eSDM) for the cryptic species– (*a*, *e*) Red-white morph, (*b*, *f*) Pure pink morph, (*c*, *g*) Yellow-white morph, and (*d*, *h*) Pure yellow morph under the (*a*–*d*) current bioclimatic (c. 1970–2000) and (*e*–*h*) paleoclimatic (last glacial maximum, LGM c. 22 kya) scenario. The LGM scenario is based on the multi-model median ensemble of three global climate models (GCMs), viz. CCSM4, MIROC-ESM, MPI-ESM-P (Braconnot *et al*., 2007).

**Fig. S20**. Niche equivalency test. Pairwise niche (*a*) identity test and (*b*, *c*) background test (*b*, symmetric; *c*, asymmetric) between the four cryptic species morphs through ENMTools. The identity and background test as cryptic species-wise pair comparisons were measured by Schoener's *D* (in red color) and Hellinger's-based *I* (in blue color) indices (provided in the respective figure).

### Supplementary Tables

**Table S1**. RAD-seq reads information and quality control statistics for every 251 individuals of *S. chamaejasme* and 3 closely related outgroups.

**Table S2**. Population summary statistics calculated for variant positions of 23,215 restrictions site-associated DNA sequencing (RAD-seq) loci.

**Table S3**. Population summary statistics calculated for all positions of 23,215 restrictions site-associated DNA sequencing (RAD-seq) loci.

**Table S4**. The paired population-level estimation of genetic diversity indices, $\Phi_{ST}$ means (lower left diagonal) and $F_{ST}$' means (upper right diagonal) for 34 populations estimated using the *populations* pipeline in Stacks. (*Excel sheet provided for Supplementary Table S4 separately)

**Table S5**. The paired mean absolute differentiation ($D_{XY}$) among 34 populations estimated using the *populations* pipeline in Stacks. (*Excel sheet provided for Supplementary Table S5 separately)

**Table S6**. The population-level cryptic species-wise estimation of the genetic diversity indices; $\Phi_{ST}$ means, $D_{XY}$ means, and $F_{ST}$' means.

**Table S7**. Comparison among the nucleotides substitution model for the selection of best fitting model based on Akaike information criterion (AIC), Bayesian information criterion (BIC), and corrected AIC (AICc) measurements using Modeltest-NG.

**Table S8**. Impact of the prior specification on the number of delimited cryptic species using BPP analysis (A11 = joint species delimitation and species tree estimation). Posterior probability values ($P$) are averaged over two runs.

**Table S9**. Marginal likelihood estimates under different scenarios (N, A1, A2, A3, A4) from Path Sampler analysis for BFD* (Bayes Factor Delimitation, *with genomic data) in 251 individuals and three closely related taxa (*Daphne tangutica*, *Daphne holosericea*, *Wikstroemia ligustrina*).

**Table S10**. Descriptive statistics (mean ± SE) and their Shapiro-Wilk normality test for the normal distribution of the continuous and categorical morphological traits for four cryptic species of *S. chamaejasme* observed from the herbarium specimens.

**Table S11**. Levene's test of homogeneity of variance assumptions for the morphological traits of *S. chamaejasme*.

**Table S12**. MANOVA test for the significant difference between four cryptic species of *S. chamaejasme* for continuous morphological traits and the trait-wise test of significant difference between the cryptic species.

**Table S13**. Post-hoc TukeyHSD MANOVA test for the trait-species wise significance difference between the four cryptic species of *S. chamaejasme*.

**Table S14**. Predictive variable selection for the ensemble species distribution modeling (eSDM) of each cryptic species of *S. chamaejasme*.

**Table S15**. Variance Inflation Factor (VIF) in different test runs to select explanatory variables for four cryptic species of *S. chamaejasme*. VIF<10 (bold text) are the selected predictive variables.

**Table S16**. Consensus land-cover selection for each cryptic species of *S. chamaejasme* based on the presence of the number of occurrence records.

**Table S17**. Model evaluation indices for the ensemble species distribution modeling (eSDM) of each cryptic species of *S. chamaejasme* using Biomod2 in R-programming language.

**Supplementary Methods**

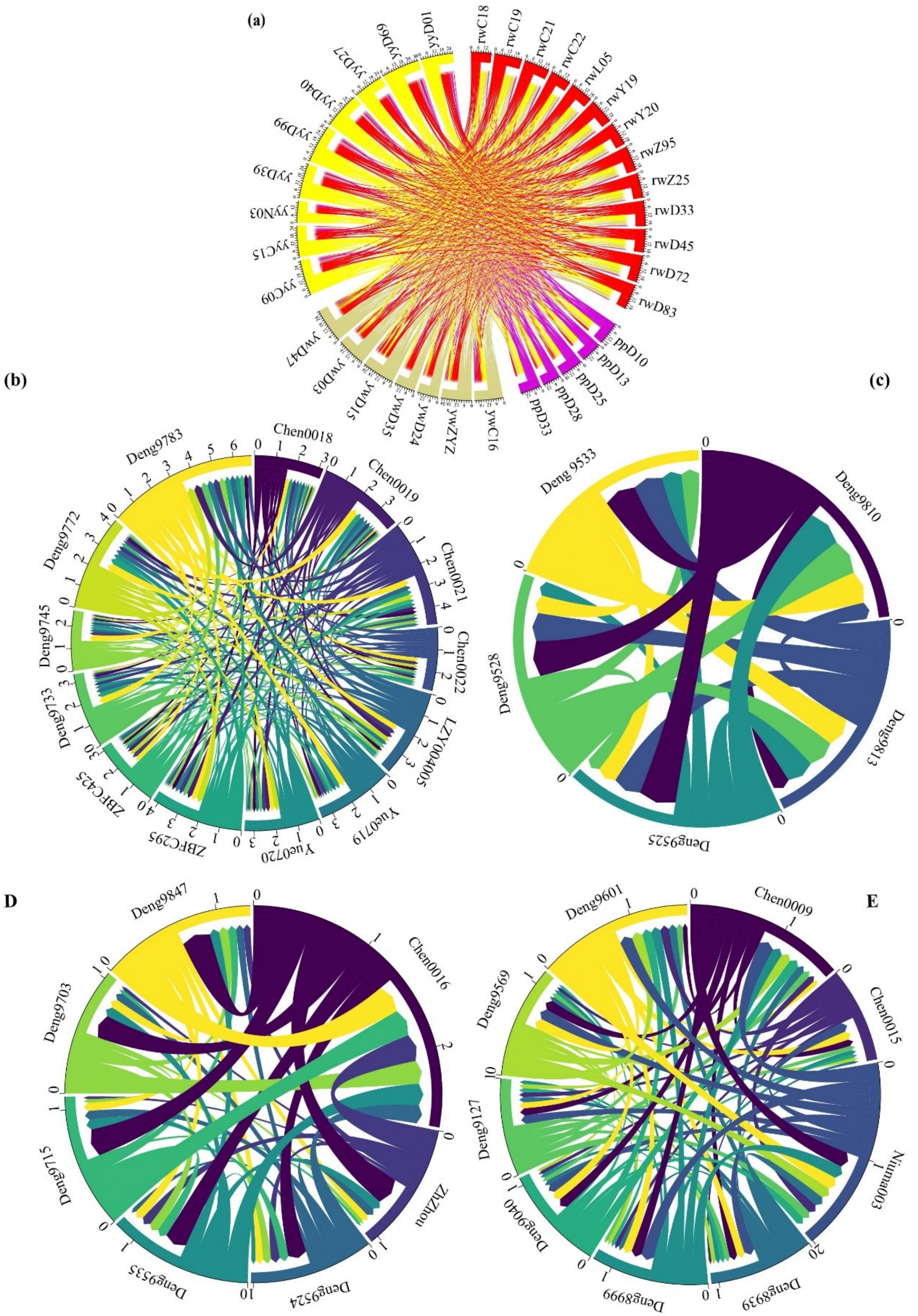


**Fig. S1.** Circos plot showing the measure of genetic variance as indicated by AMOVA-based $F_{ST}$ within the paired population level of each cryptic species. (***a***) *Stellera chamaejasme*, (***b***) Red-white morph, (***c***) Pure pink morph, (***d***) Yellow-white morph, and (***e***) Pure yellow morph. Each population is represented by a different gradient of colors.

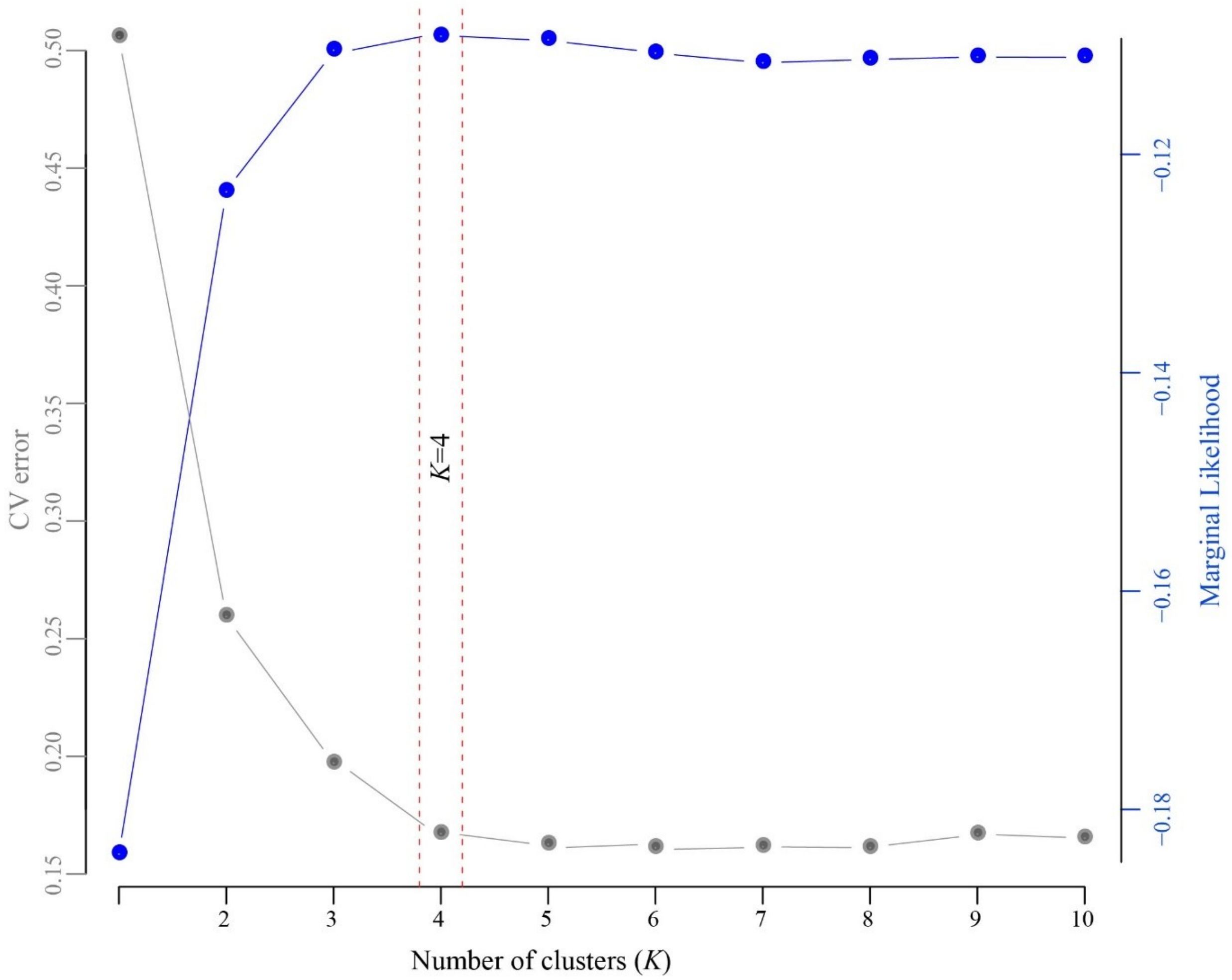


**Fig. S2.** Model selection for the optimal number of clusters (*K*=4) inferred using fastSTRUCTURE (for *K*=1–10) based on the lowest cross-validation (CV) error (grey color) and the highest marginal likelihood (blue color) estimates.

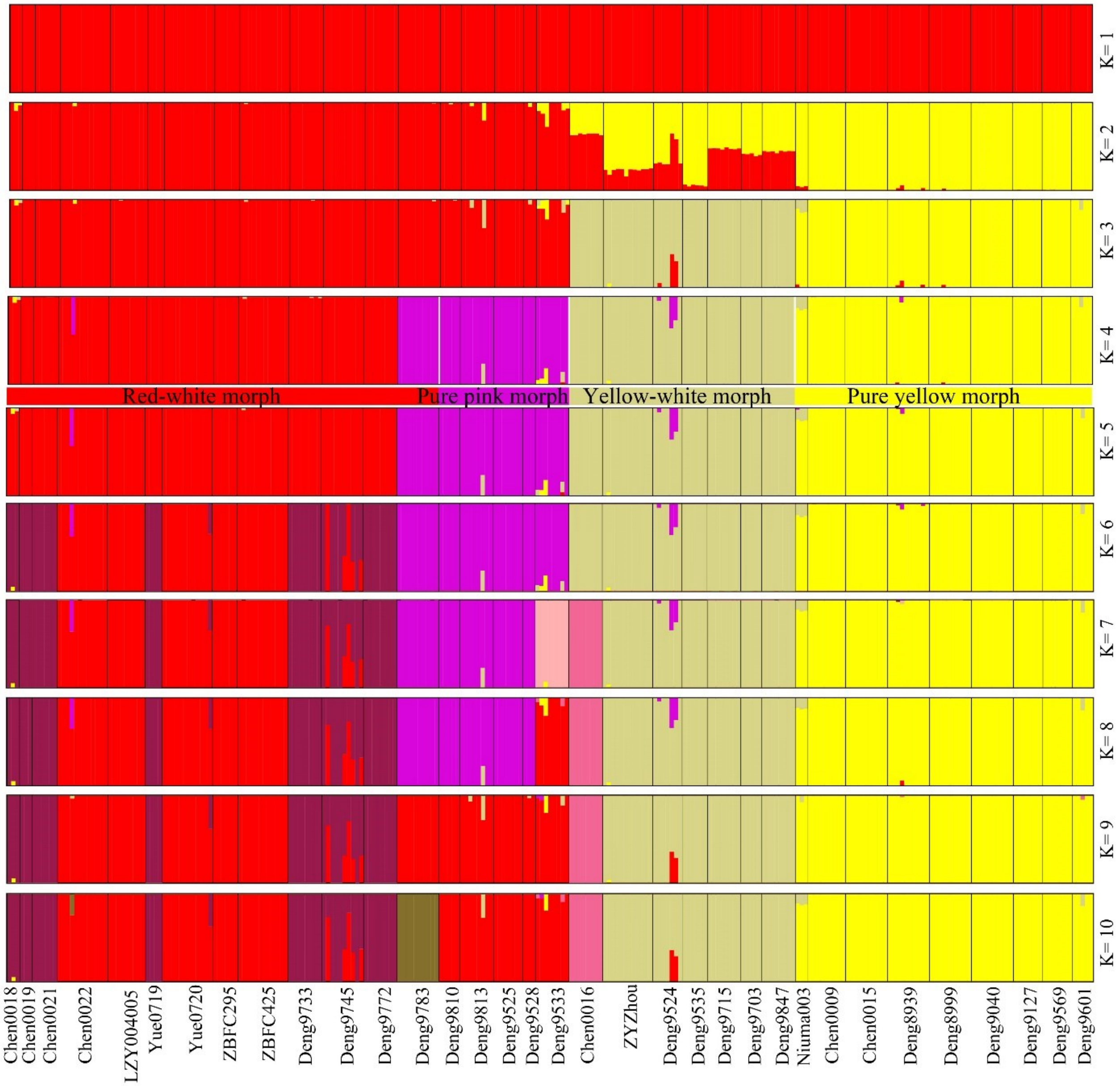


**Fig. S3.** Hierarchically structured clusters K=1–10 identified through fastSTRUCTURE analyses for 34 populations. Colors indicate a posterior probability of assignment of each individual to a particular cluster. The number of clusters K for each analysis is displayed on the left side of the bar graph. The cluster for K=4 is equivalent to the putative four cryptic species as colored by red – Red-white morph (RW), pink – Pure pink morph (PP), beige – Yellow-white morph (YW), yellow – Pure yellow morph (YY), and coded the same throughout the figures.

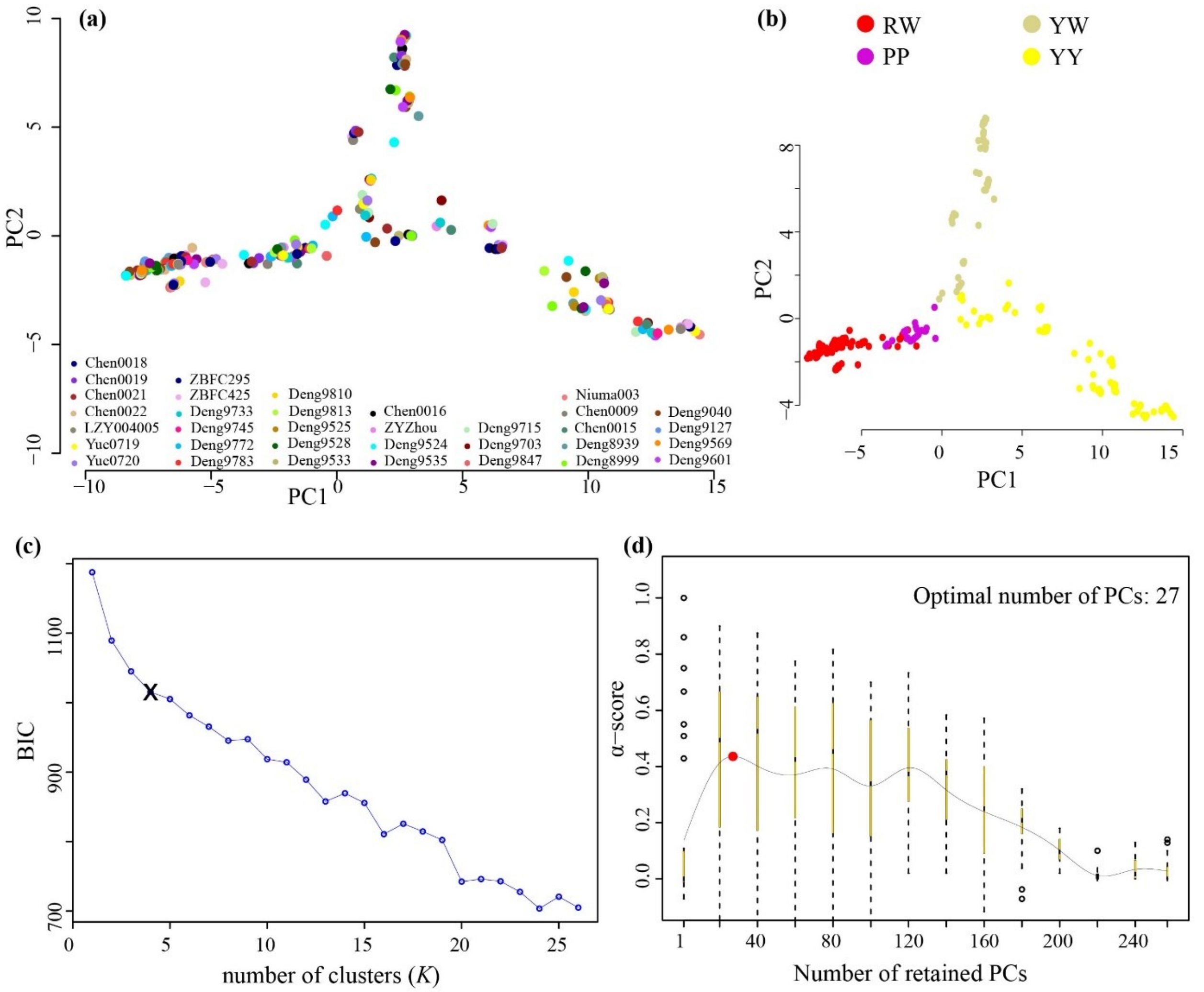


**Fig. S4.** Distribution of (***a***) 34 population individuals and (***b***) four morphs of *S. chamaejasme* along principal component (PC) of genetic variation based on single nucleotide polymorphism (SNPs) dataset, along with their eigenvalues plot. Individuals are color-coded according to their population identities (see **Fig. 2** for population information). (***c***) The optimal number of clusters *K* based on the Bayesian inference criterion (BIC). The marked (X) indicates the actual number of clusters *K* from where the BIC values decrease slowly on increasing the clusters. (***d***) The optimal number of PCs is retained based on α-score optimization of spline interpolation.

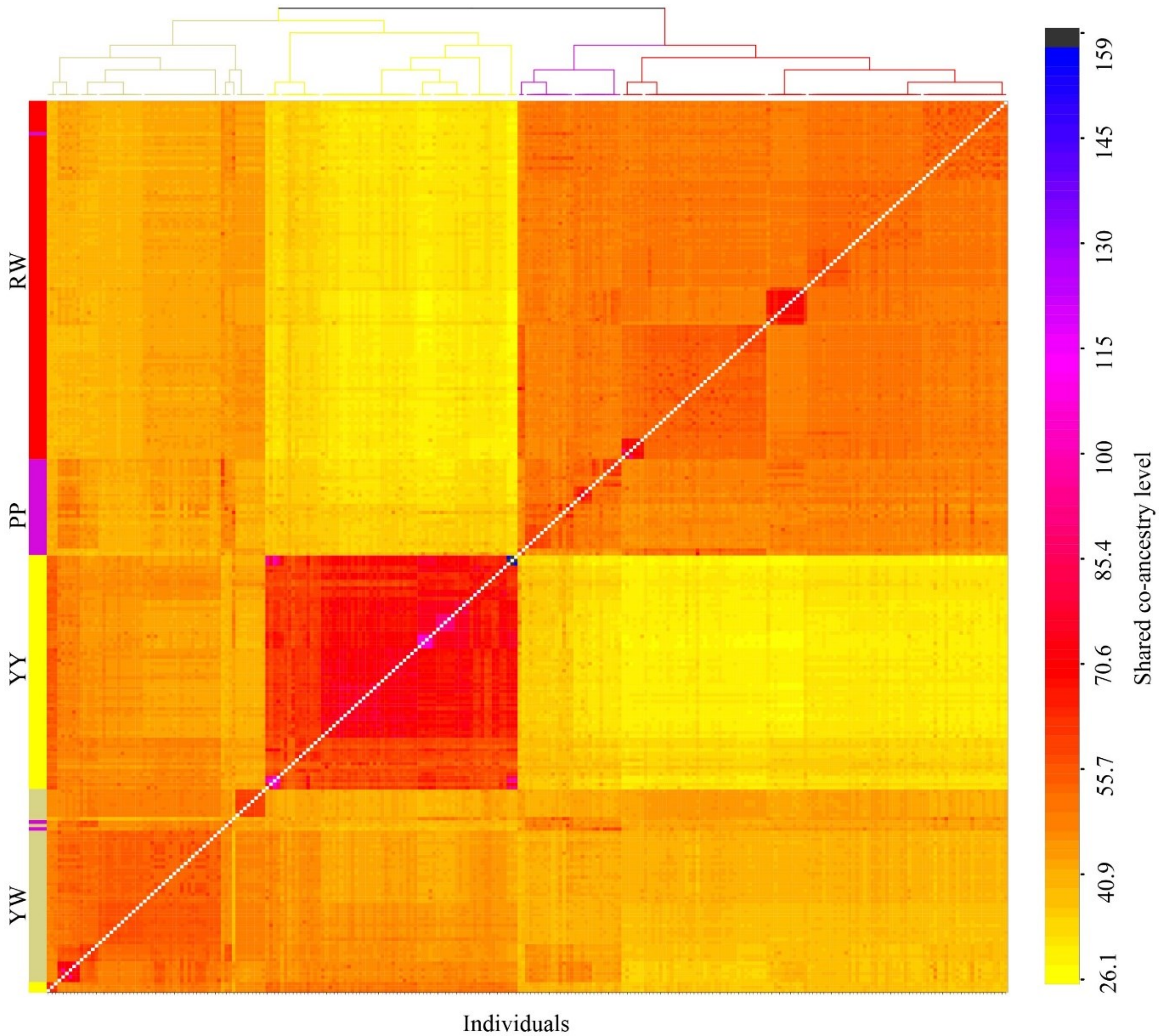


**Fig. S5.** Pairwise co-ancestry heat map depicting variation among individuals of four cryptic species of *S. chamaejasme* inferred through fineRADstructure. The dendrogram shows a clustering of individual samples. Each tip in a tree corresponds to an individual, branches are coded according to clusters. Shared co-ancestry levels are coded from low (yellow) to high (black).

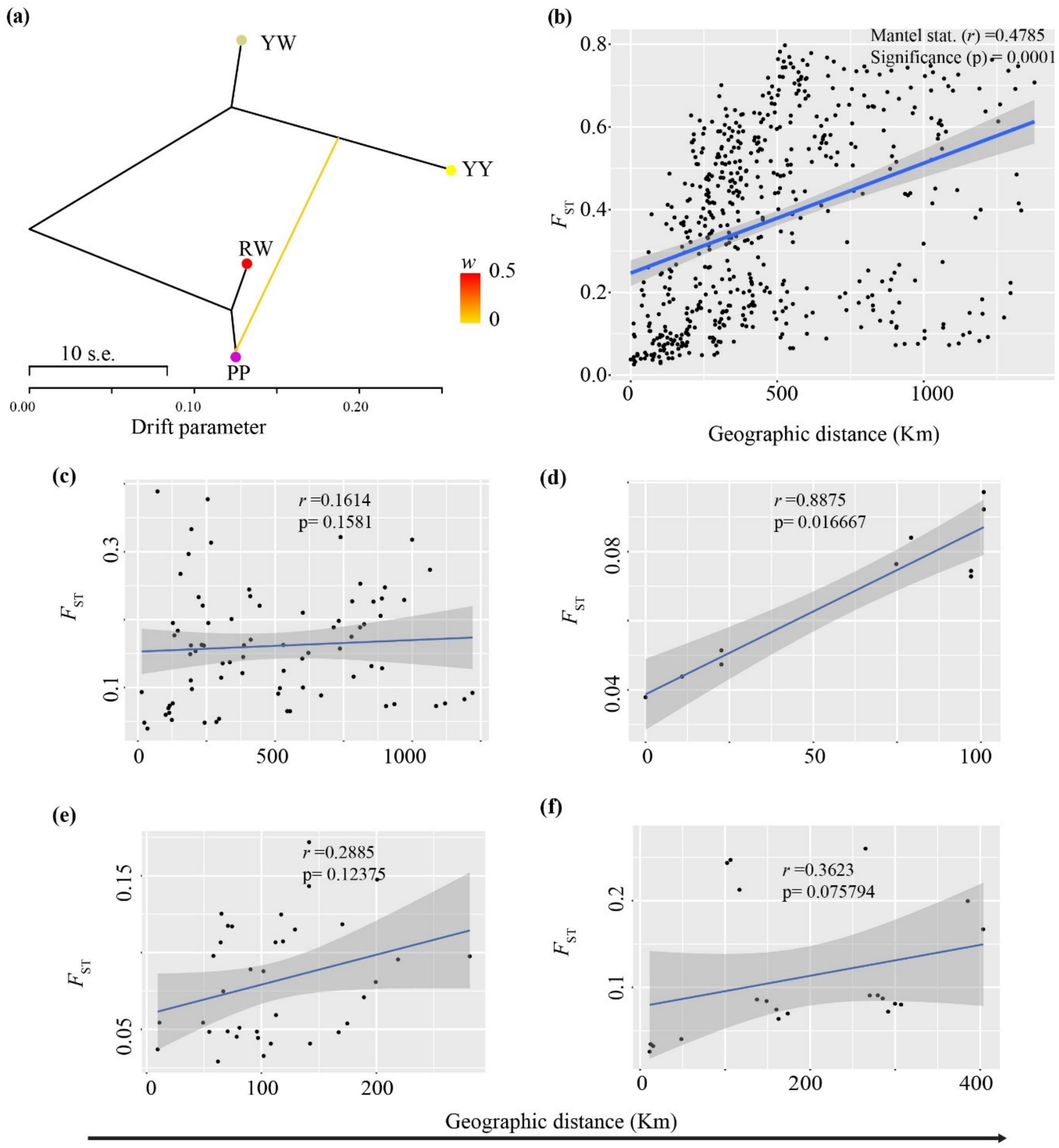


**Fig. S6.** (***a***) Detection of gene flow between the cryptic species morph using TREEMIX. A colored arrow line represents the direction of gene flow. The horizontal scale bar at the bottom (drift parameter) shows the entries' tenfold average standard error, and the color scale shows the migration weight *w*. Linear regression lines showing the correlations between genetic distance ($F_{ST}$, along y-axis) and geographic distances (km, along x-axis) inferred as the function of isolation-by-distance (IBD) for (***b***) *S. chamaejasme*, (***c***) Red-white morph, (***d***) Pure pink morph, (***e***) Yellow-white morph, and (***f***) Pure-yellow morph. Mantel correlation coefficient (r) at a significance level $p < 0.05$ is presented in the figure.

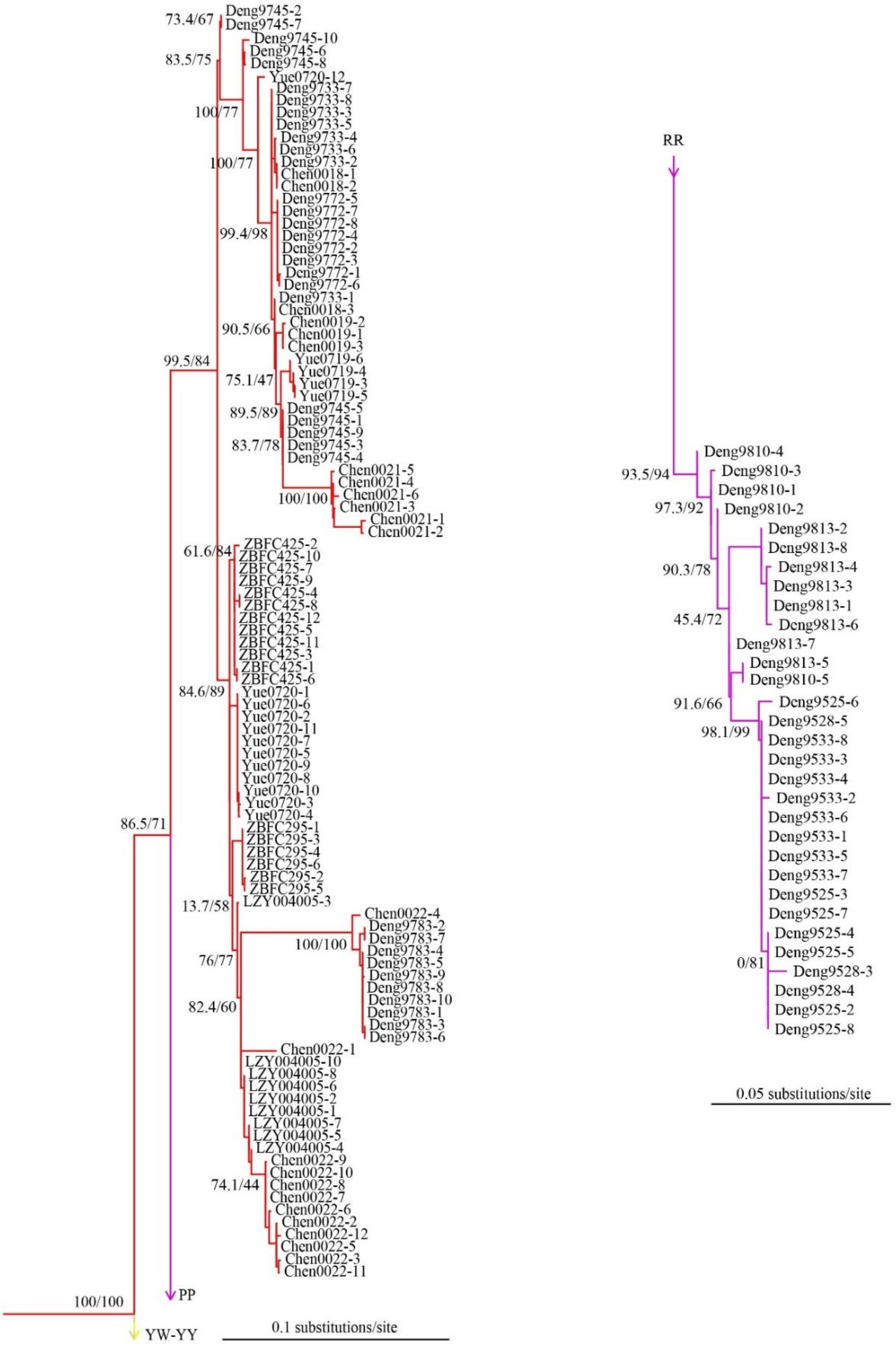


**Fig. S7.** The detailed population-level cryptic species-wise maximum likelihood (ML) phylogenetic tree inferred with IQ-TREE based on the 5,800 SNPs for the Red-white morph (red-colored lineages) and Pure pink morph (pink-colored lineages). Each major node is labeled with SH-aLRT/ultrafast bootstrap support reported as a percentage. The scale bar represents the expected number of substitutions per nucleotide site.

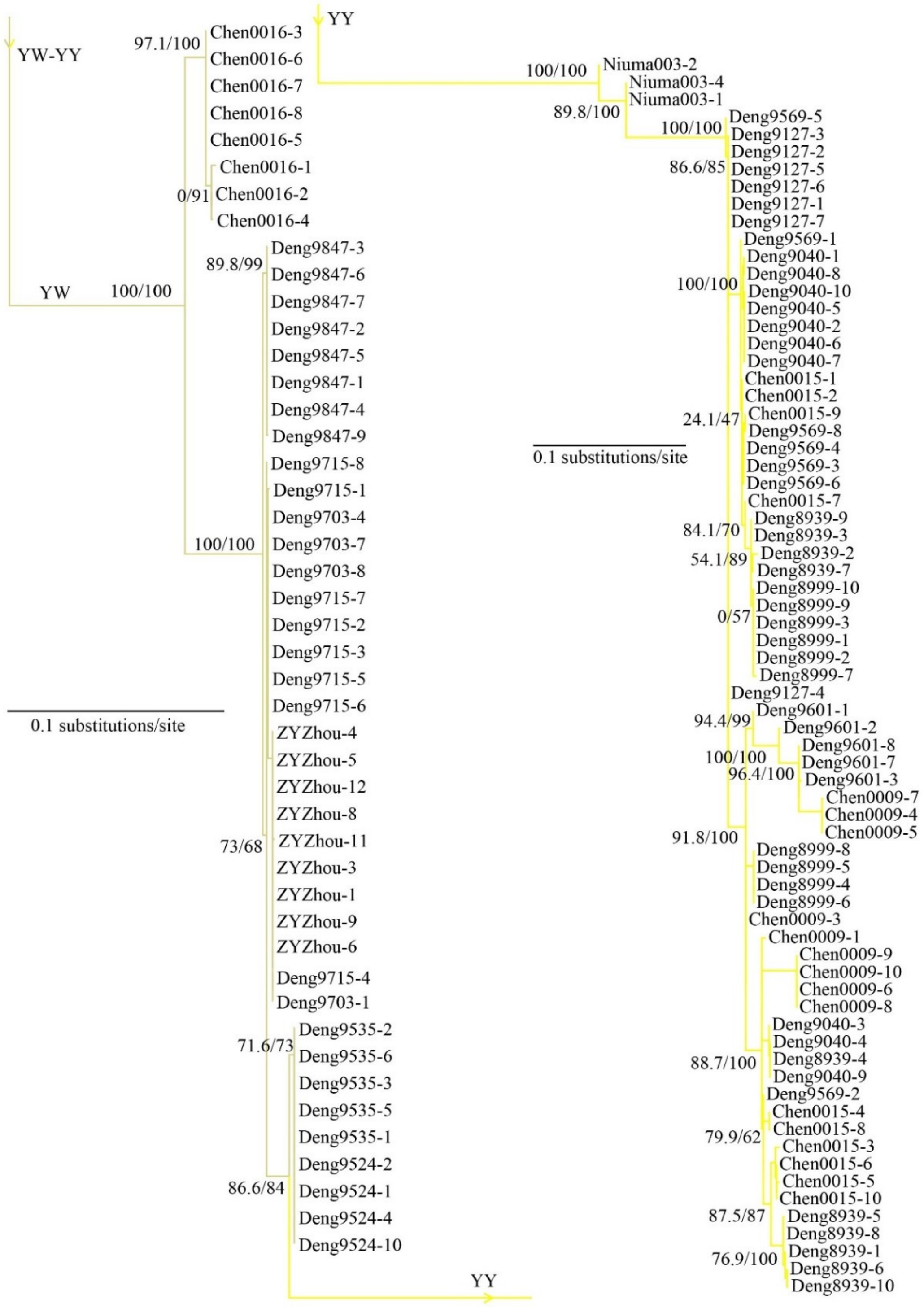


**Fig. S8.** The detailed population-level cryptic species-wise maximum likelihood (ML) phylogenetic tree inferred using IQ-TREE based on the 5,800 SNPs for Yellow-white morph (beige-colored lineages) and Pure yellow morph (yellow-colored lineages). Each major node is labeled with SH-aLRT/ultrafast bootstrap support reported as a percentage. The scale bar represents the expected number of substitutions per nucleotide site.

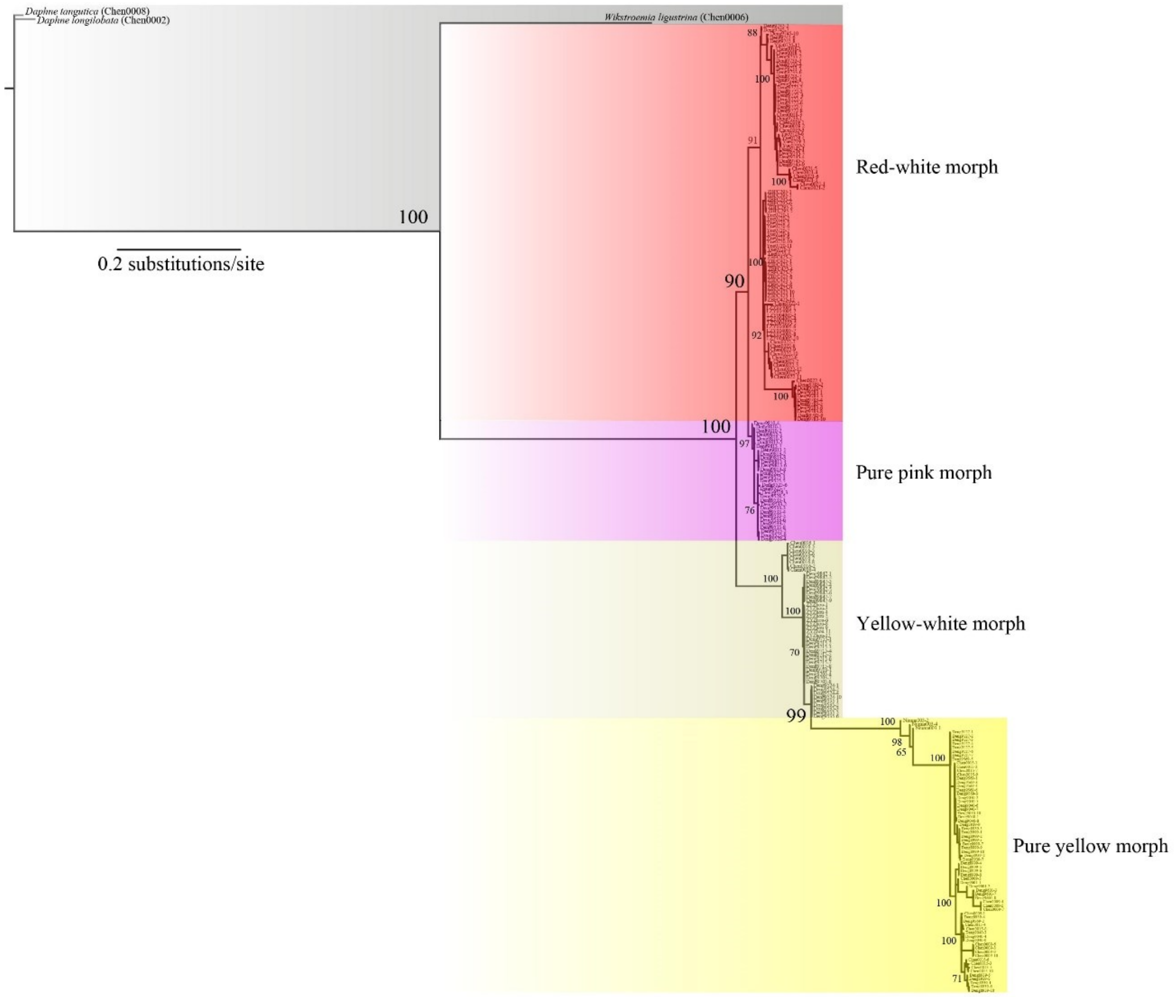


**Fig. S9.** Bayesian phylogenetic tree inferred with MrBayes analysis based on the 5,800 SNPs of 251 individuals representing four cryptic species of *S. chamaejasme* and three closely related taxa (*Wikstroemia ligustrina*, *Daphne longlobalata*, *Daphne tangutica*). Each major node is labeled with a posterior probability value in the tree. The scale bar represents the expected number of substitutions per nucleotide site.

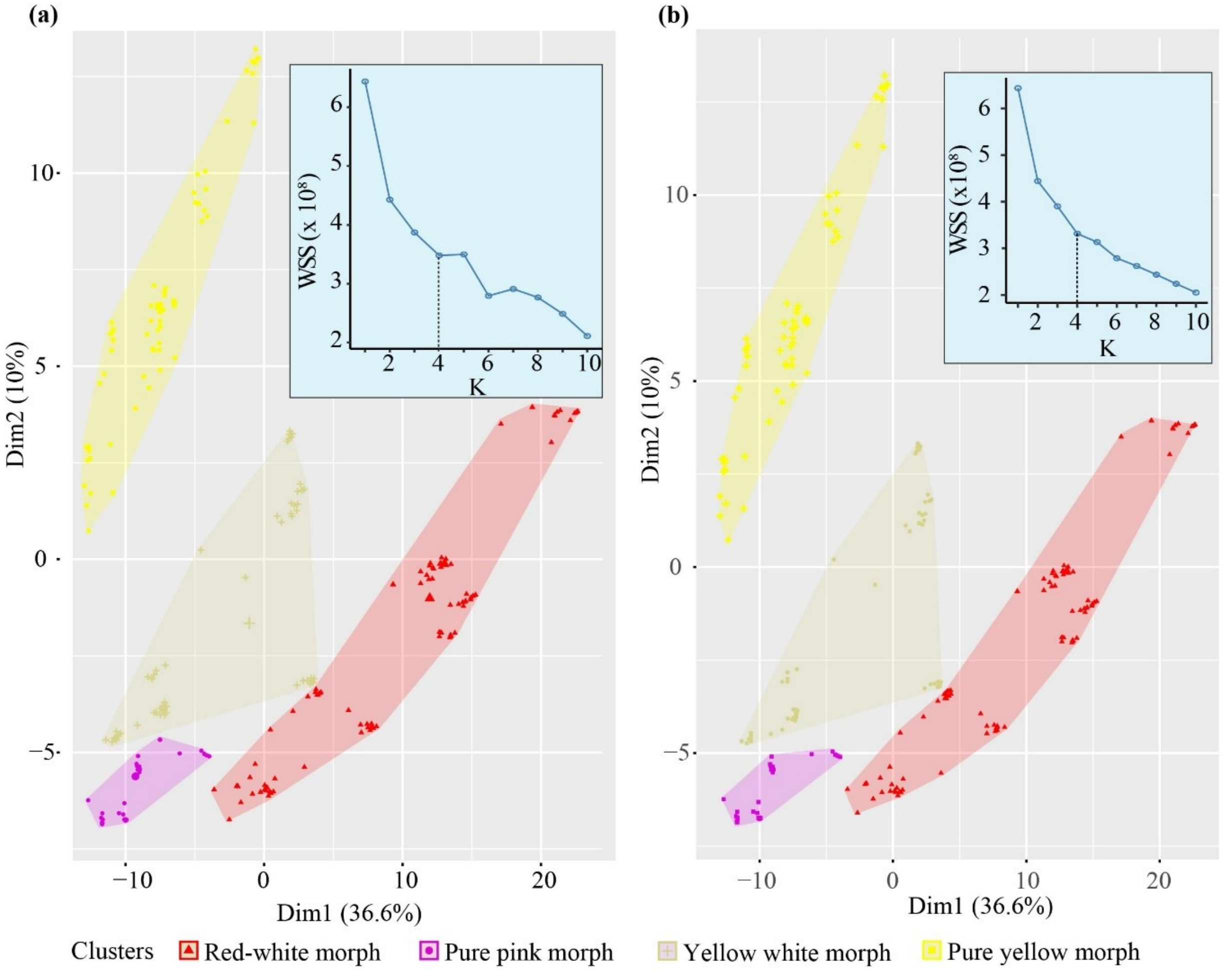


**Fig. S10.** Genetic distance based unsupervised machine learning (UML) clustering analysis inferred through (***a***) *K*-means clustering and (***b***) Partitioning Around Medoids (PAM) clustering based on within-cluster-sum of squared errors (WSS) index for optimal numbers of clusters *K*. The clusters are represented in two dimensions (Dim1, 36.6%, and Dim2, 10%).

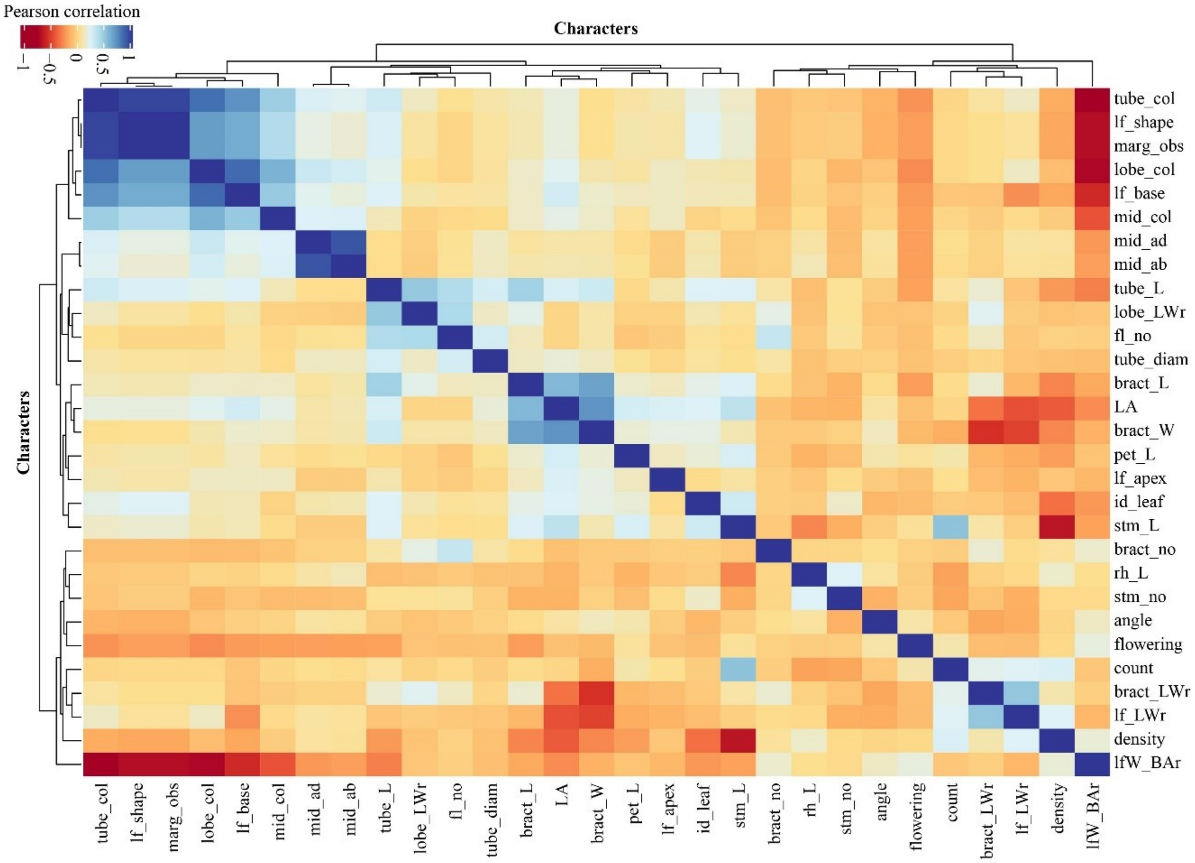


**Fig. S11.** Correlation heatmap of the cryptic species, confirming the least pair correlations across 29 least correlated morphological traits. The figure with blue color indicates positive correlations, and the dark red color indicates negative correlations. Abbreviations for all morphological traits are shown in Supplementary Table S10.

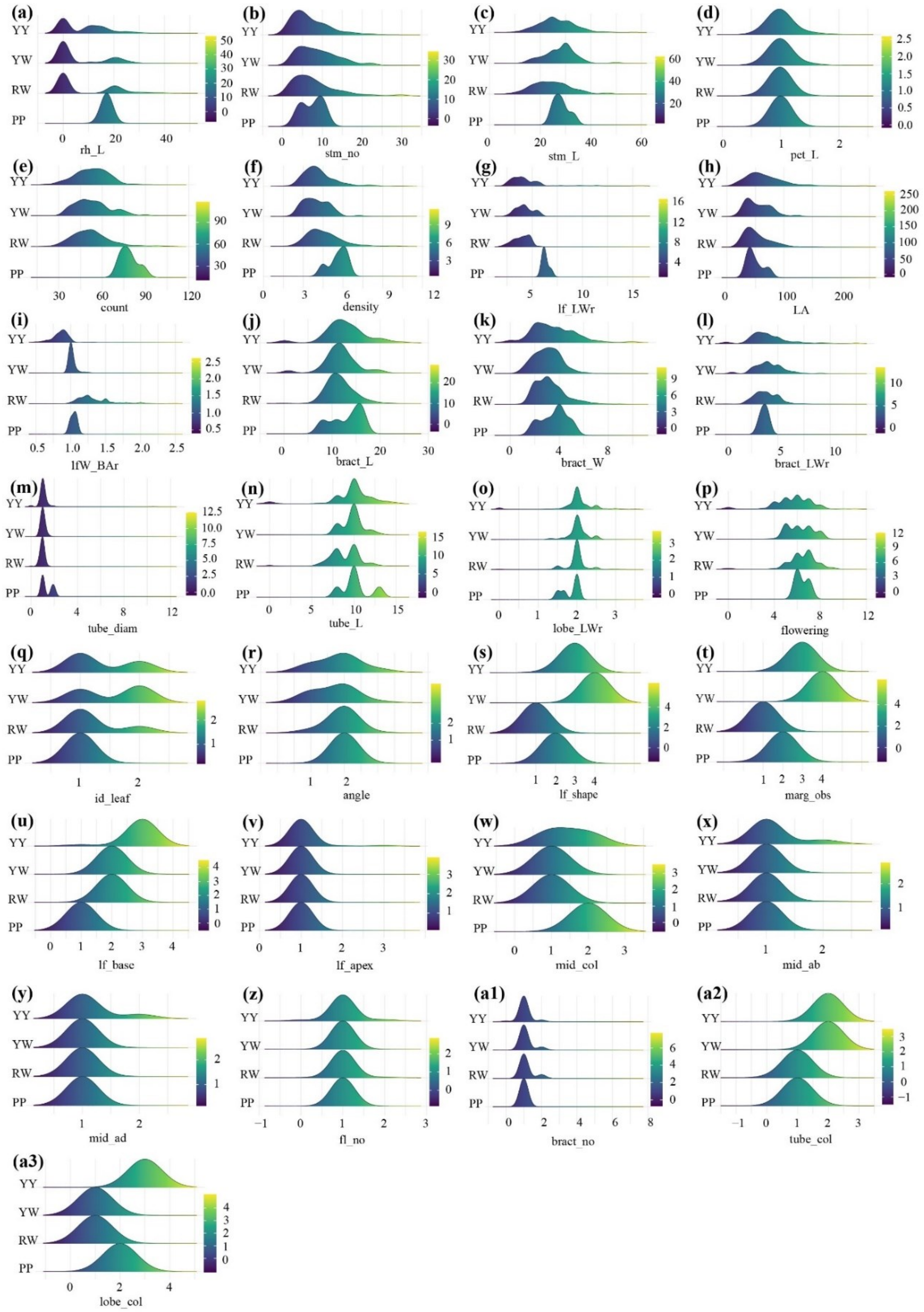


**Fig. S12.** The ridgeplots (***a*–*a3***) visualization for the comparison of all the morphological traits between the four cryptic species of *S. chamaejasme*. The gradient color indicates the trait values provided in the scale. Abbreviations for all morphological traits are shown in Supplementary Table S10.

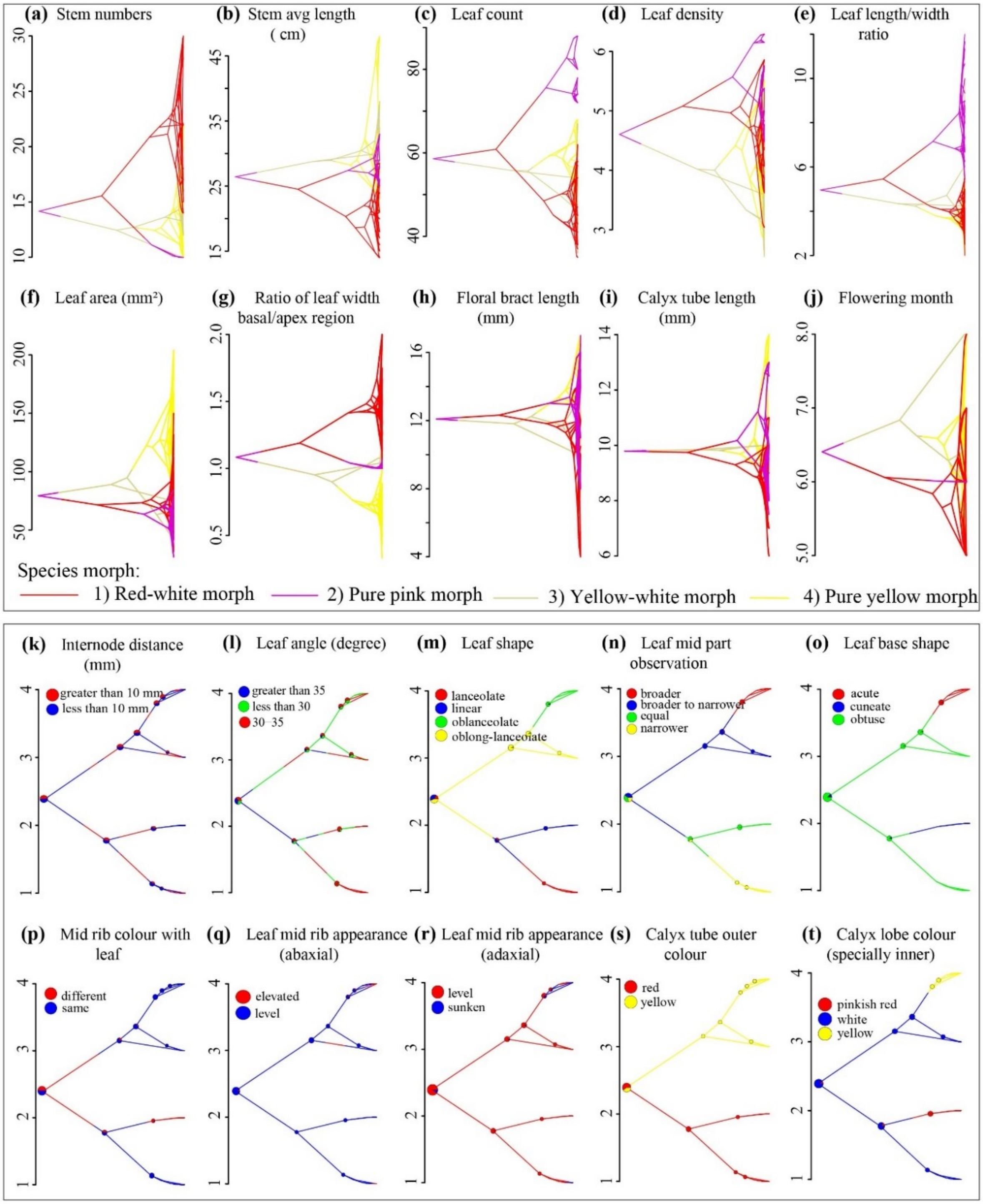


**Fig. S13.** Phylogenetic traitgrams for the significant morphological traits, i.e., (***a–j***) continuous traits and (***k–t***) categorical traits with strong phylogenetic signals constructed using the phytools. The vertical placement of each tip corresponds to the mean trait values of those individuals for the continuous trait and that of cryptic species for categorical traits. Color in the lineages represents the cryptic species (coded as the same color throughout the figures) for continuous traits, whereas the trait class is represented by color for categorical traits. Abbreviations for all morphological traits are shown in Supplementary Table S10.

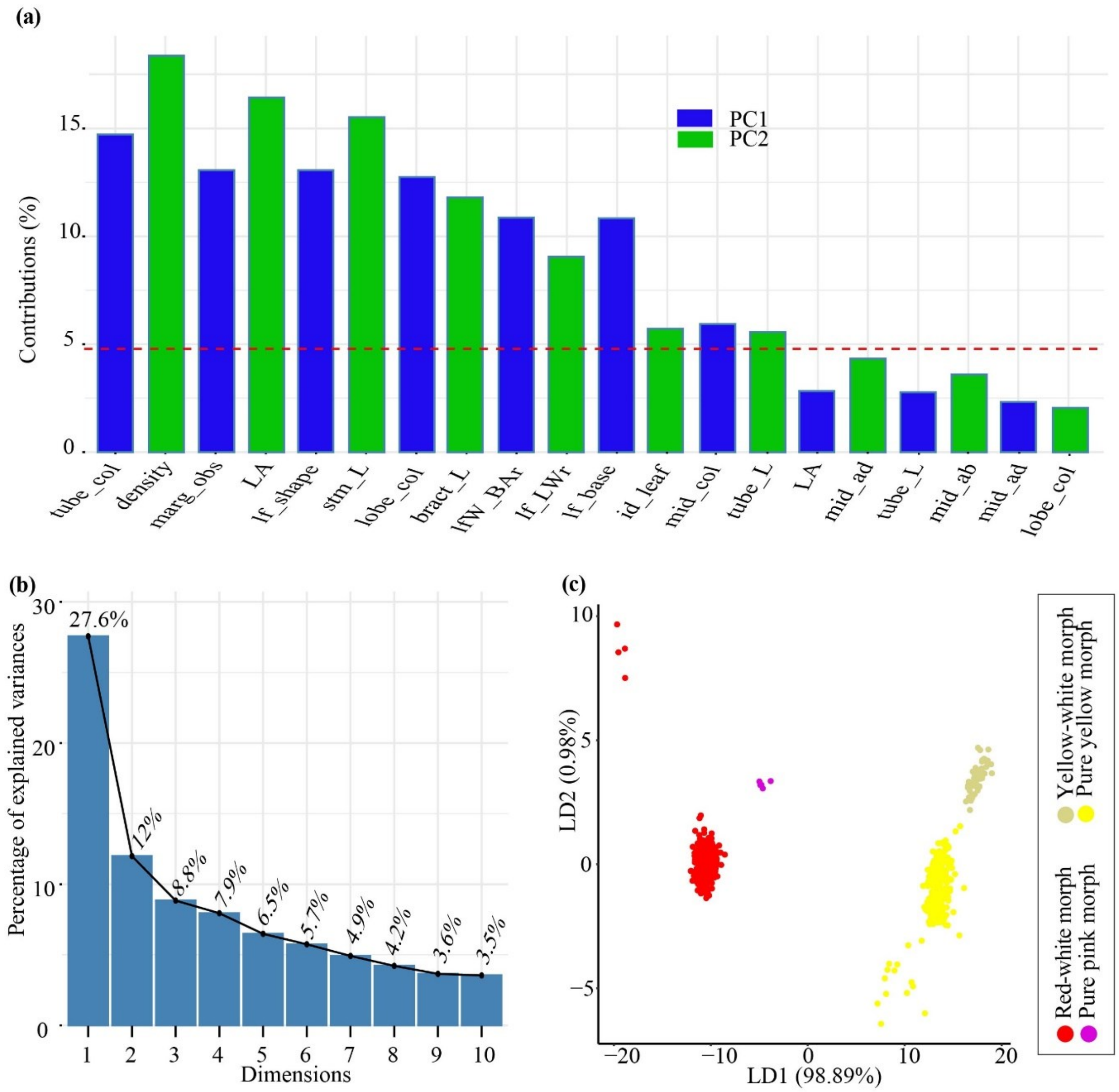


**Fig. S14.** (***a***) Top ten contributing morphological traits under the two principal components (blue, PC1; green, PC2) analysis scores. The red dashed line indicates the expected average contribution if variable contributions were uniform. (***b***) The scree plot represents the variance explained by the first ten principal components for morphological traits. Abbreviations for all morphological traits are shown in *SI Appendix*, Table S10. (***c***) The discriminated clusters of the cryptic species into two major components (LD1, 98.89%; LD2, 0.98%) inferred through linear discriminant analysis (LDA) plots of morphological traits. Ellipses are the normal contour lines of each cryptic species with a probability of 90%.

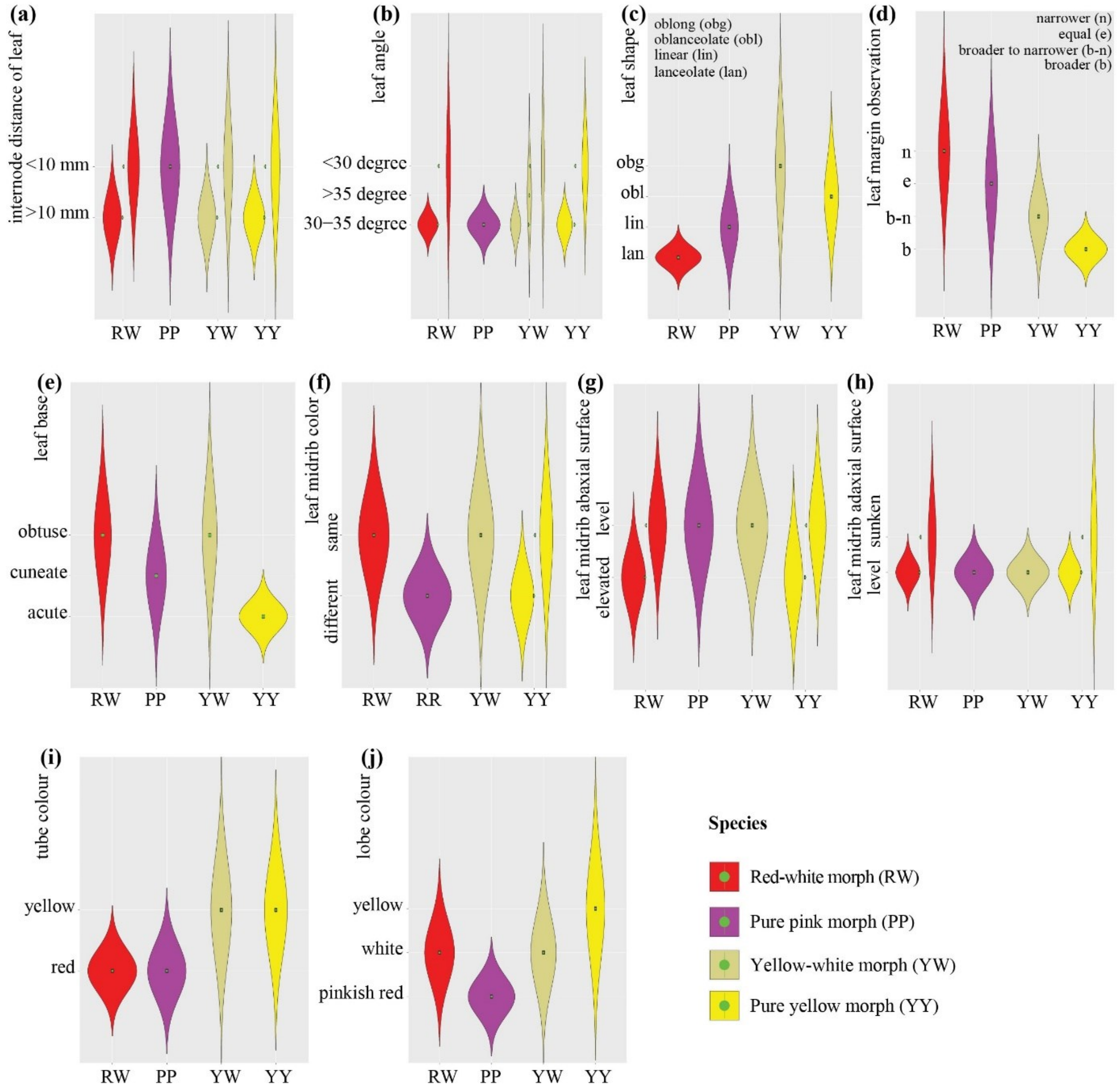


**Fig. S15.** Violin plot (***a*–*j***) to visualize means with standard deviations and the distribution of each significant categorical trait of different cryptic species. Cryptic species are coded in the same color throughout the figures. Abbreviations for all morphological traits are shown in Supplementary Table S10.

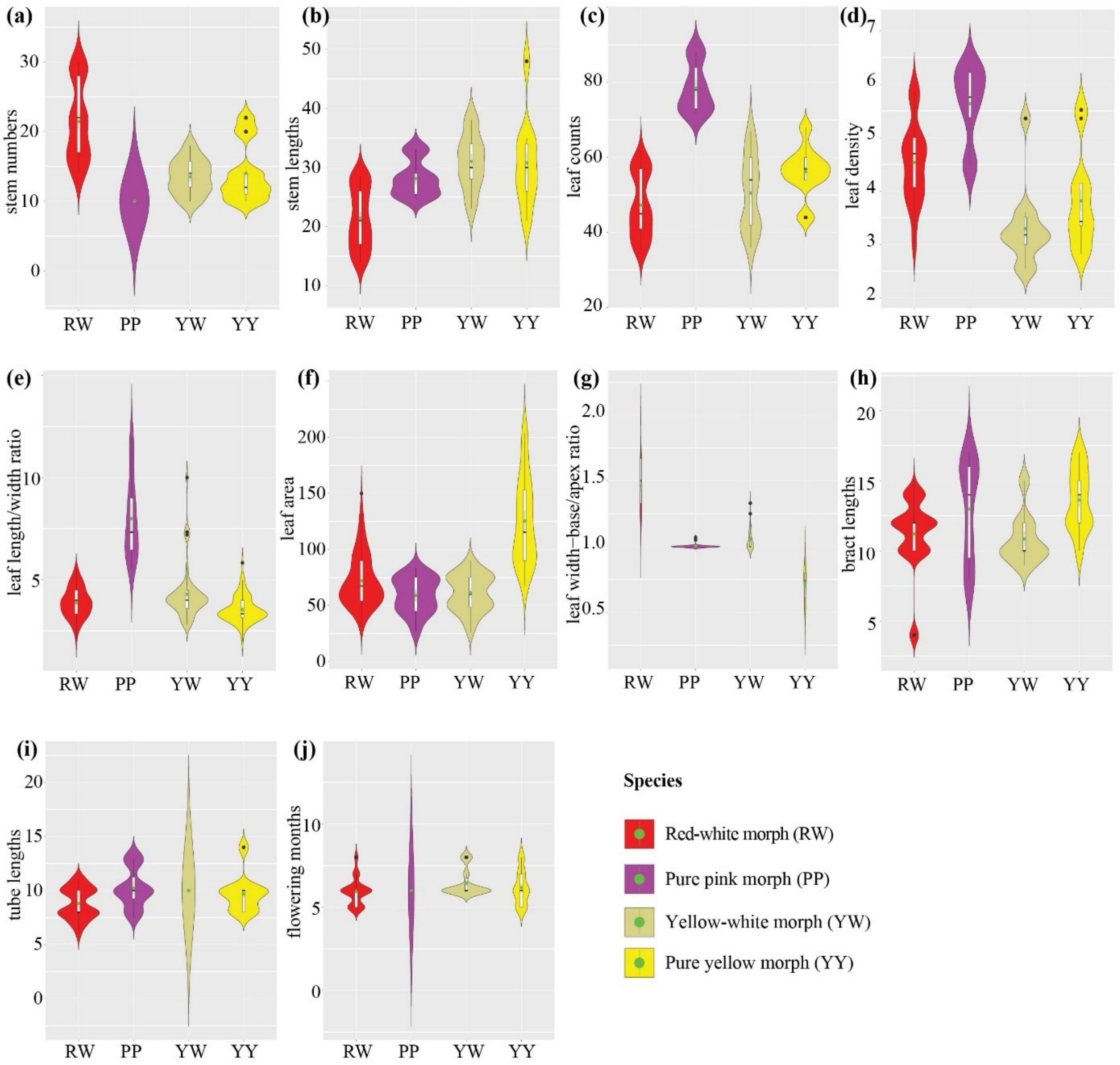


**Fig. S16.** Violin plot (***a–j***) to visualize mean with standard deviations and the distribution of each significant continuous trait of different cryptic species. Cryptic species are coded in the same color throughout the figures. Abbreviations for all morphological traits are shown in Supplementary Table S10.

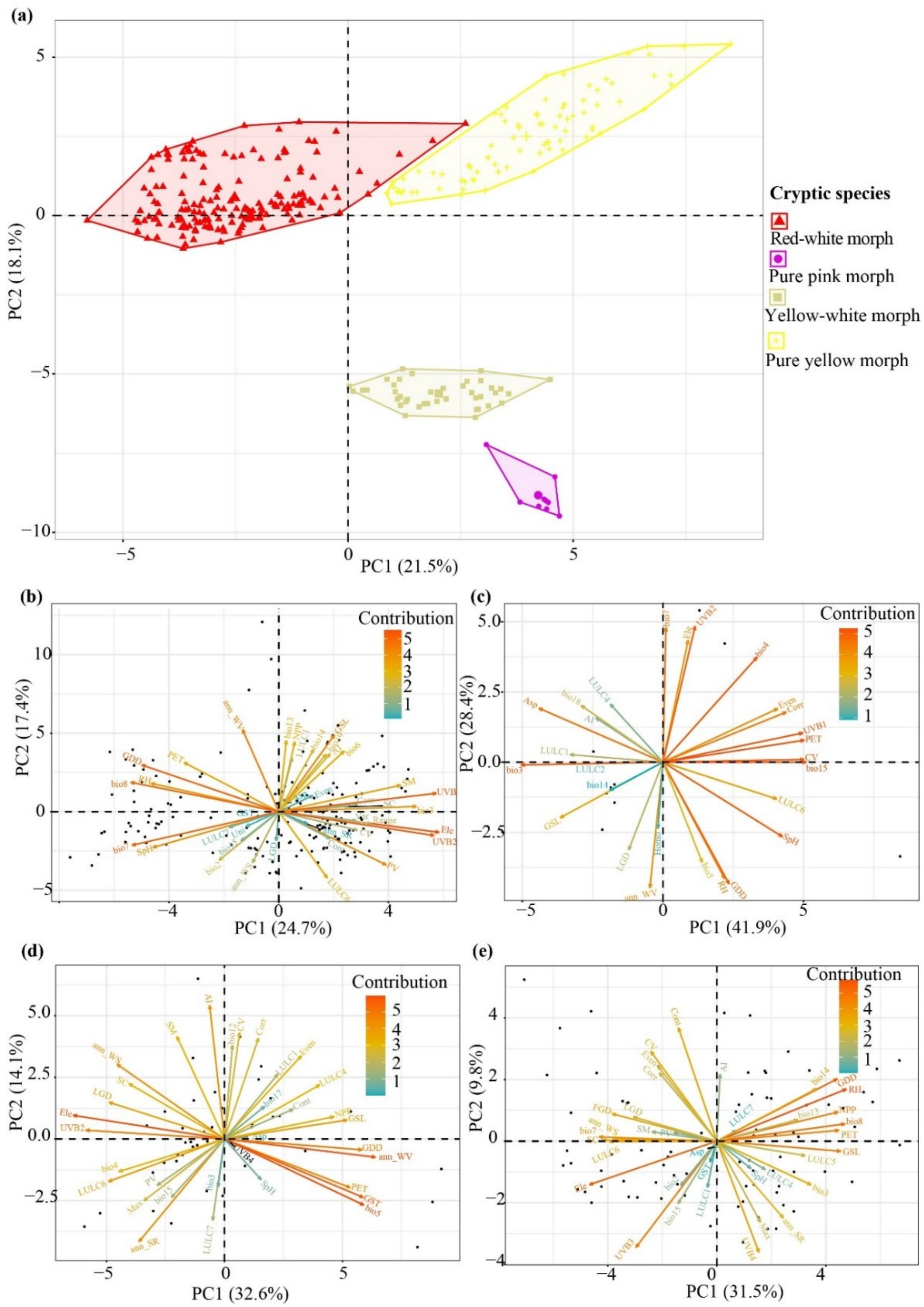


**Fig. S17.** Ecological characterization for *Stellera chamaejasme*. (***a***) PCA clustering of all the predictive environmental variables for four cryptic species into two PCs (PC1, 21.5%; PC2, 18.1%). The relative contributions of predictive environmental variables to the PCs for the cryptic species – (***b***) Red-white morph, (***c***) Pure pink morph, (***d***) Yellow-white morph, and (***e***) Pure yellow morph inferred through principal component analysis (PCA) biplots. Abbreviations for all environmental variables are shown in Supplementary Table S14.

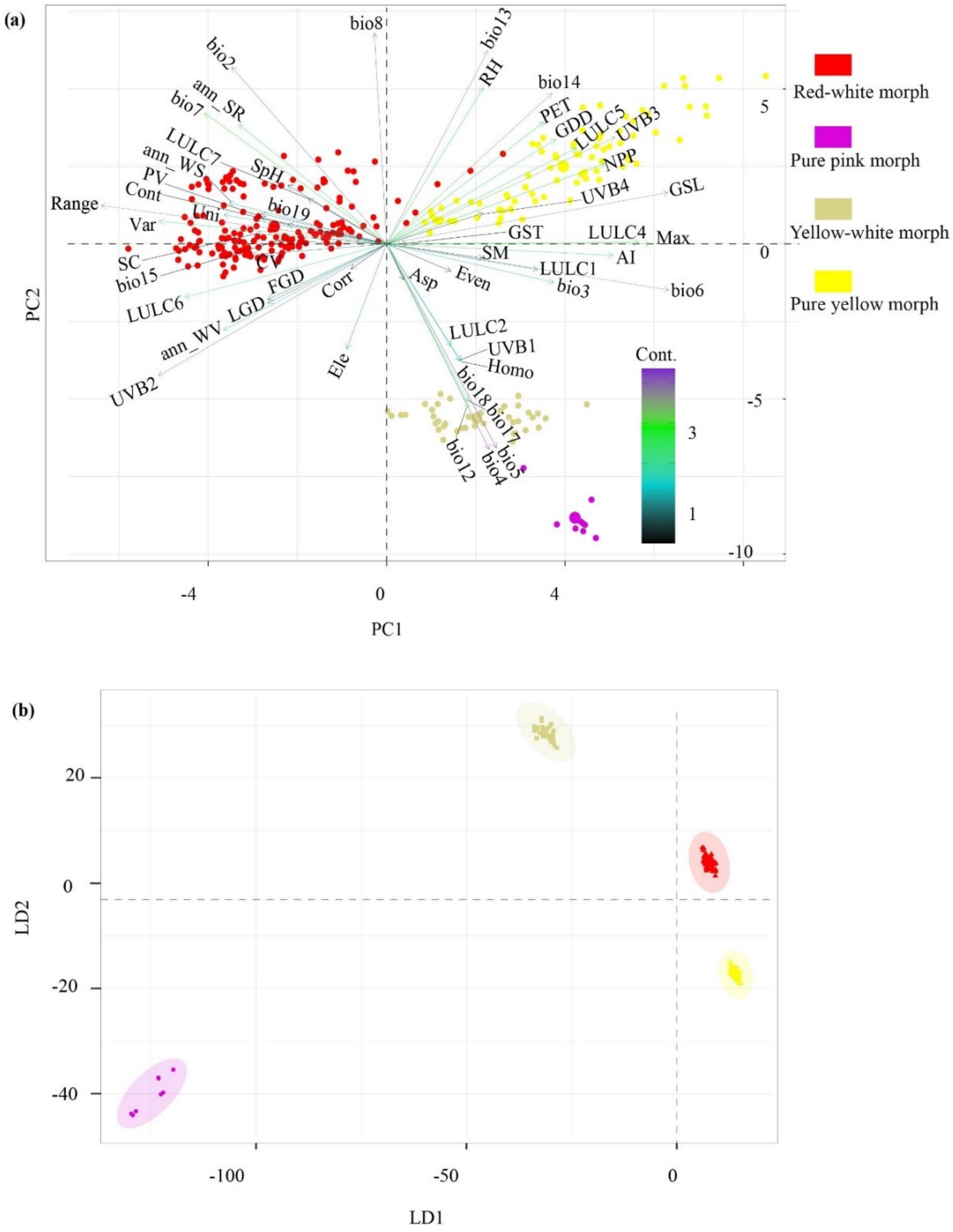


**Fig. S18.** Ecological niche analysis as a measure of cryptic species differentiation in *S. chamaejasme*. (***a***) The relative contributions (Cont.) of predictive environmental variables to the PCs (PC1, 21.5%; PC2, 18.1%) inferred through PCA biplots and (***b***) discriminated clusters of the cryptic species into two components (LD1, 61.41%; LD2, 21.15%) inferred through LDA plots. Abbreviations for all environmental variables are shown in Supplementary Table S14.

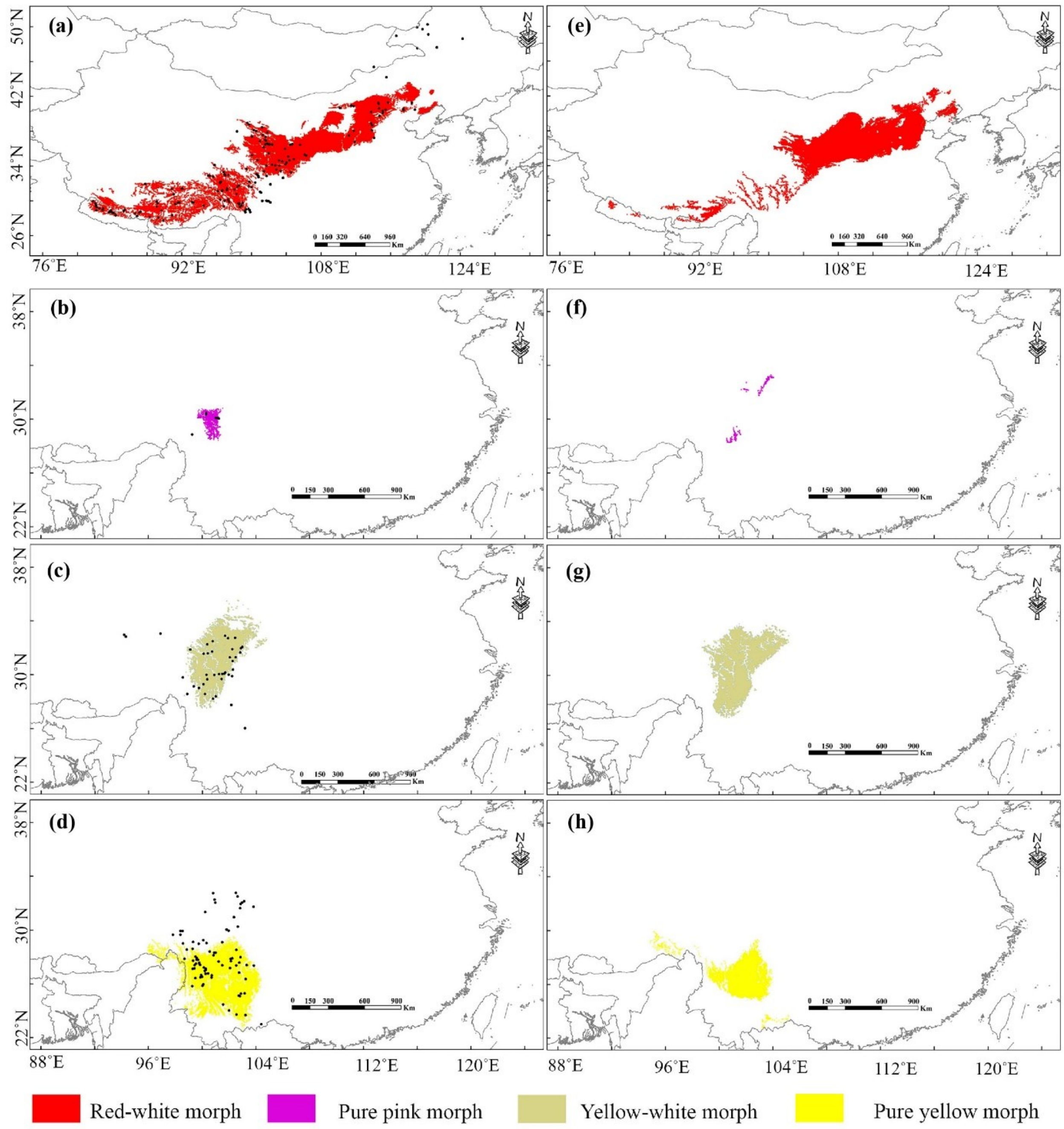


**Fig. S19.** Habitat suitability predicted by ensemble species distribution modeling (eSDM) for the cryptic species– (***a***, ***e***) Red-white morph, (***b***, ***f***) Pure pink morph, (***c***, ***g***) Yellow-white morph, and (***d***, ***h***) Pure yellow morph under the (***a***–***d***) current bioclimatic (c. 1970–2000) and (***e***–***h***) paleoclimatic (last glacial maximum, LGM c. 22 kya) scenario. The LGM scenario is based on the multi-model median ensemble of three global climate models (GCMs), viz. CCSM4, MIROC-ESM, MPI-ESM-P (Braconnot *et al*., 2007).

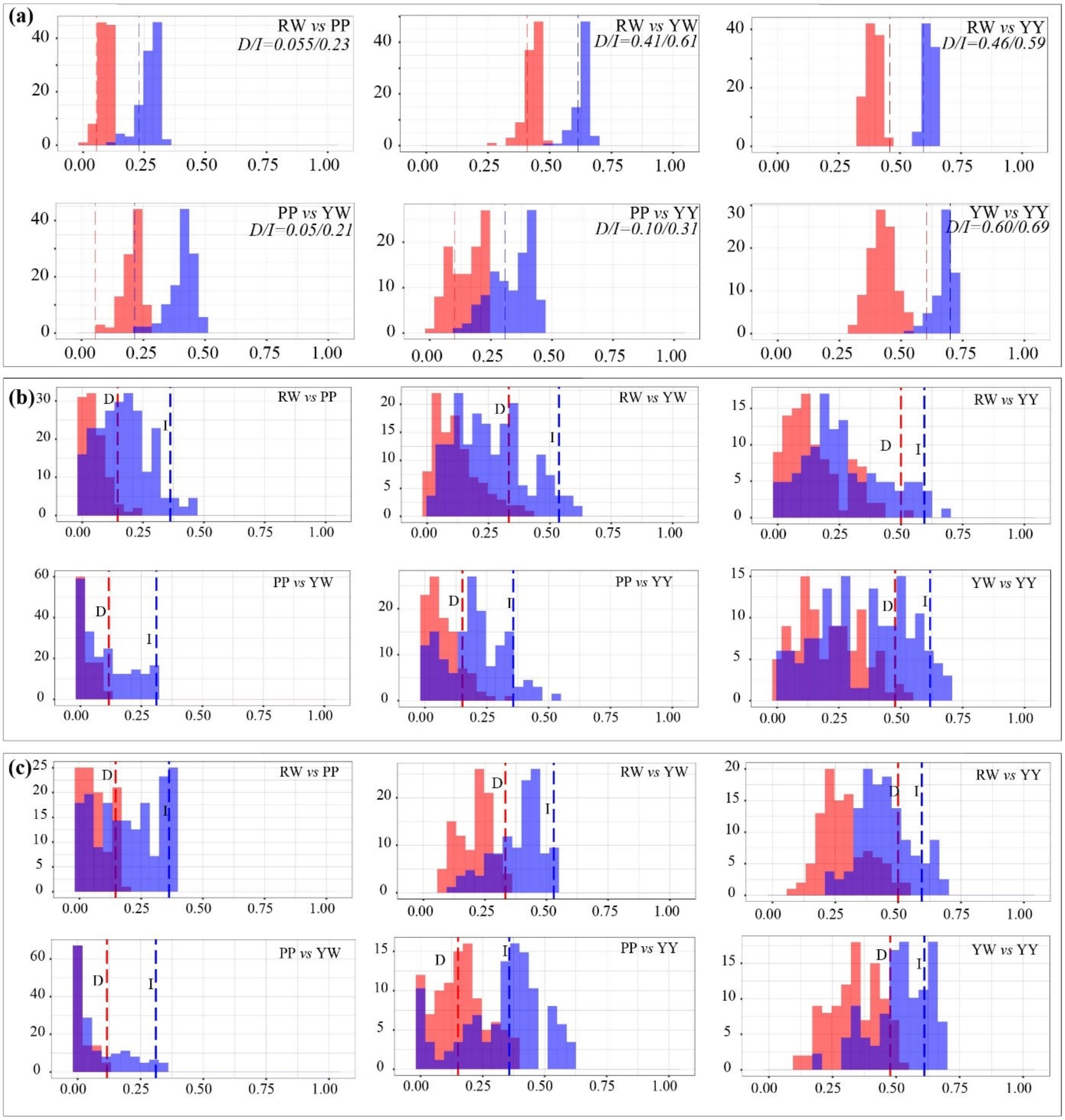


**Fig. S20.** Niche equivalency test. Pairwise niche (***a***) identity test and (***b***, ***c***) background test (***b***, symmetric; ***c***, asymmetric) between the four cryptic species morphs through ENMTools. The identity and background test as cryptic species-wise pair comparisons were measured by Schoener's *D* (in red color) and Hellinger's-based *I* (in blue color) indices (provided in the respective figure).

**Supplementary Tables**

**Table S1.** RAD-seq reads information and quality control statistics for every 251 individuals of *S. chamaejasme* and 3 closely related outgroups.

| Species | Cryptic species | Sample | Individual | Raw Base (bp) | Clean Base (bp) | Effective Rate (%) | Error Rate (%) | Q20 (%) | Q30 (%) | GC Content (%) |
|---|---|---|---|---|---|---|---|---|---|---|
| *Daphne longilobata* | Outgroup | Chen0002 | Chen0002 | 3,032,109,000 | 2,891,013,300 | 95.35 | 0.03 | 96.84 | 91.96 | 39.02 |
| *Wikstroemia ligustrina* | Outgroup | Chen0006 | Chen0006 | 3,947,501,400 | 3,947,475,000 | 100 | 0.04 | 95.48 | 90.05 | 39.97 |
| *Daphne tangutica* | Outgroup | Chen0008 | Chen0008 | 4,169,701,500 | 3,958,197,300 | 94.93 | 0.03 | 95.61 | 89.78 | 38.86 |
| *Stellera chamaejasme* | Red-white | CNQ18-1 | Chen0018-1 | 396,239,400 | 340,122,300 | 85.84 | 0.03 | 95.36 | 89.67 | 42.08 |
| *Stellera chamaejasme* | Red-white | CNQ18-2 | Chen0018-2 | 3,817,346,400 | 3,651,635,100 | 95.66 | 0.04 | 94.44 | 87.62 | 41.16 |
| *Stellera chamaejasme* | Red-white | CNQ18-3 | Chen0018-3 | 3,616,473,600 | 3,531,793,200 | 97.66 | 0.03 | 95.32 | 88.97 | 40.96 |
| *Stellera chamaejasme* | Red-white | CNQ19-1 | Chen0019-1 | 4,535,227,200 | 4,409,727,300 | 97.23 | 0.03 | 95.72 | 89.8 | 39.77 |
| *Stellera chamaejasme* | Red-white | CNQ19-2 | Chen0019-2 | 3,286,938,900 | 3,138,589,800 | 95.49 | 0.03 | 95.66 | 89.87 | 40.23 |
| *Stellera chamaejasme* | Red-white | CNQ19-3 | Chen0019-3 | 3,791,265,300 | 3,690,200,100 | 97.33 | 0.03 | 95.68 | 89.8 | 40.21 |
| *Stellera chamaejasme* | Red-white | CCQ21-1 | Chen0021-1 | 4,091,985,900 | 3,940,084,500 | 96.29 | 0.03 | 96 | 90.55 | 40.02 |
| *Stellera chamaejasme* | Red-white | CCQ21-2 | Chen0021-2 | 3,861,444,000 | 3,685,964,100 | 95.46 | 0.03 | 95.85 | 90.44 | 40.09 |
| *Stellera chamaejasme* | Red-white | CCQ21-3 | Chen0021-3 | 3,377,184,900 | 3,234,676,500 | 95.78 | 0.03 | 95.31 | 89.1 | 39.36 |
| *Stellera chamaejasme* | Red-white | CCQ21-4 | Chen0021-4 | 10,824,196,800 | 2,622,797,100 | 24.23 | 0.07 | 86.81 | 77.84 | 54 |
| *Stellera chamaejasme* | Red-white | CCQ21-5 | Chen0021-5 | 3,142,940,400 | 3,028,536,000 | 96.36 | 0.03 | 95.19 | 88.72 | 40.63 |
| *Stellera chamaejasme* | Red-white | CCQ21-6 | Chen0021-6 | 3,414,583,800 | 3,196,769,700 | 93.62 | 0.03 | 95.78 | 90.24 | 41.49 |
| *Stellera chamaejasme* | Red-white | CLS22-1 | Chen0022-1 | 3,175,448,700 | 2,900,697,900 | 91.35 | 0.03 | 95.7 | 90.3 | 41.55 |
| *Stellera chamaejasme* | Red-white | CLS22-10 | Chen0022-10 | 6,687,529,200 | 5,233,554,900 | 78.26 | 0.04 | 94.47 | 88.7 | 42.26 |

| | | | | | | | | | | |
|---|---|---|---|---|---|---|---|---|---|---|
| *Stellera chamaejasme* | Red-white | CLS22-11 | Chen0022-11 | 5,876,002,200 | 5,408,818,200 | 92.05 | 0.04 | 95.21 | 89.44 | 40.67 |
| *Stellera chamaejasme* | Red-white | CLS22-12 | Chen0022-12 | 3,976,604,400 | 3,570,100,200 | 89.78 | 0.04 | 95.11 | 89.38 | 40.11 |
| *Stellera chamaejasme* | Red-white | CLS22-2 | Chen0022-2 | 3,464,481,300 | 3,335,681,700 | 96.28 | 0.03 | 95.67 | 89.91 | 40.28 |
| *Stellera chamaejasme* | Red-white | CLS22-3 | Chen0022-3 | 4,011,235,800 | 3,781,384,500 | 94.27 | 0.03 | 95.89 | 90.48 | 41.01 |
| *Stellera chamaejasme* | Red-white | CLS22-4 | Chen0022-4 | 4,305,793,800 | 4,084,218,900 | 94.85 | 0.03 | 95.68 | 89.97 | 40.86 |
| *Stellera chamaejasme* | Red-white | CLS22-5 | Chen0022-5 | 5,575,712,700 | 5,316,590,400 | 95.35 | 0.04 | 95.22 | 89.63 | 40.52 |
| *Stellera chamaejasme* | Red-white | CLS22-6 | Chen0022-6 | 3,876,255,000 | 3,772,673,100 | 97.33 | 0.03 | 95.57 | 89.71 | 41.33 |
| *Stellera chamaejasme* | Red-white | CLS22-7 | Chen0022-7 | 3,239,841,300 | 2,889,797,100 | 89.2 | 0.03 | 96.3 | 91.58 | 39.64 |
| *Stellera chamaejasme* | Red-white | CLS22-8 | Chen0022-8 | 4,264,291,200 | 4,096,290,000 | 96.06 | 0.03 | 95.8 | 90.13 | 40.18 |
| *Stellera chamaejasme* | Red-white | CLS22-9 | Chen0022-9 | 3,350,350,800 | 2,963,455,500 | 88.45 | 0.04 | 95.03 | 89.26 | 41.12 |
| *Stellera chamaejasme* | Red-white | DCX33-1 | Deng9733-1 | 4,269,672,900 | 3,843,672,900 | 90.02 | 0.03 | 95.31 | 89.31 | 41.17 |
| *Stellera chamaejasme* | Red-white | DCX33-2 | Deng9733-2 | 3,000,784,800 | 2,923,380,900 | 97.42 | 0.03 | 96.03 | 90.55 | 40.77 |
| *Stellera chamaejasme* | Red-white | DCX33-3 | Deng9733-3 | 3,011,643,000 | 2,922,108,600 | 97.03 | 0.03 | 95.58 | 89.38 | 40.72 |
| *Stellera chamaejasme* | Red-white | DCX33-4 | Deng9733-4 | 3,417,524,400 | 3,317,606,700 | 97.08 | 0.03 | 95.72 | 89.68 | 40.98 |
| *Stellera chamaejasme* | Red-white | DCX33-5 | Deng9733-5 | 3,221,097,000 | 3,082,252,500 | 95.69 | 0.03 | 95.66 | 89.69 | 41.82 |
| *Stellera chamaejasme* | Red-white | DCX33-6 | Deng9733-6 | 3,302,868,300 | 3,208,306,800 | 97.14 | 0.03 | 95.09 | 88.49 | 40.56 |
| *Stellera chamaejasme* | Red-white | DCX33-7 | Deng9733-7 | 3,277,438,500 | 3,148,684,800 | 96.07 | 0.03 | 94.85 | 88.06 | 41.52 |
| *Stellera chamaejasme* | Red-white | DCX33-8 | Deng9733-8 | 4,108,212,900 | 2,678,121,300 | 65.19 | 0.03 | 96.45 | 91.28 | 40.68 |
| *Stellera chamaejasme* | Red-white | DCX45-1 | Deng9745-1 | 3,271,354,500 | 3,179,983,800 | 97.21 | 0.03 | 96.57 | 91.57 | 40.41 |
| *Stellera* | Red-white | DCX45- | Deng9745-10 | 3,528,077,10 | 3,414,369,900 | 96.78 | 0.03 | 95.94 | 89.94 | 40.47 |

| | | | | | | | | | | |
|---|---|---|---|---|---|---|---|---|---|---|
| *chamaejasme* | | 10 | | 0 | | | | | | |
| *Stellera chamaejasme* | Red-white | DCX45-2 | Deng9745-2 | 3,170,193,000 | 3,079,873,200 | 97.15 | 0.03 | 96.5 | 91.47 | 40.66 |
| *Stellera chamaejasme* | Red-white | DCX45-3 | Deng9745-3 | 3,764,043,600 | 3,643,491,900 | 96.8 | 0.03 | 96.06 | 90.45 | 40.44 |
| *Stellera chamaejasme* | Red-white | DCX45-4 | Deng9745-4 | 3,302,476,200 | 3,209,449,500 | 97.18 | 0.03 | 96.45 | 91.31 | 40.06 |
| *Stellera chamaejasme* | Red-white | DCX45-5 | Deng9745-5 | 3,243,729,900 | 3,140,570,400 | 96.82 | 0.03 | 95.12 | 88.53 | 41.3 |
| *Stellera chamaejasme* | Red-white | DCX45-6 | Deng9745-6 | 3,268,750,500 | 3,172,617,900 | 97.06 | 0.03 | 95.94 | 89.89 | 40.58 |
| *Stellera chamaejasme* | Red-white | DCX45-7 | Deng9745-7 | 3,529,694,700 | 3,428,526,600 | 97.13 | 0.03 | 96.03 | 90.06 | 40.47 |
| *Stellera chamaejasme* | Red-white | DCX45-8 | Deng9745-8 | 3,421,904,400 | 3,321,193,500 | 97.06 | 0.03 | 95.8 | 89.58 | 40.12 |
| *Stellera chamaejasme* | Red-white | DCX45-9 | Deng9745-9 | 3,127,361,700 | 3,028,011,900 | 96.82 | 0.03 | 96.27 | 90.94 | 39.83 |
| *Stellera chamaejasme* | Red-white | DJX72-1 | Deng9772-1 | 3,501,546,900 | 3,389,929,500 | 96.81 | 0.03 | 95.1 | 88.44 | 40.81 |
| *Stellera chamaejasme* | Red-white | DJX72-2 | Deng9772-2 | 3,438,599,400 | 3,350,036,400 | 97.42 | 0.03 | 96.56 | 91.53 | 40.53 |
| *Stellera chamaejasme* | Red-white | DJX72-3 | Deng9772-3 | 3,554,420,100 | 3,456,137,100 | 97.23 | 0.03 | 96.42 | 91.3 | 40.23 |
| *Stellera chamaejasme* | Red-white | DJX72-4 | Deng9772-4 | 3,026,032,200 | 2,961,380,700 | 97.86 | 0.03 | 96.38 | 91.18 | 40.44 |
| *Stellera chamaejasme* | Red-white | DJX72-5 | Deng9772-5 | 4,314,350,400 | 4,160,803,800 | 96.44 | 0.03 | 95.35 | 89.25 | 40.8 |
| *Stellera chamaejasme* | Red-white | DJX72-6 | Deng9772-6 | 3,124,585,200 | 3,045,487,200 | 97.47 | 0.03 | 96.59 | 91.63 | 40.67 |
| *Stellera chamaejasme* | Red-white | DJX72-7 | Deng9772-7 | 3,944,062,200 | 3,806,862,600 | 96.52 | 0.03 | 96 | 90.34 | 39.65 |
| *Stellera chamaejasme* | Red-white | DJX72-8 | Deng9772-8 | 4,839,071,100 | 4,629,145,800 | 95.66 | 0.03 | 95.87 | 90.28 | 38.12 |
| *Stellera chamaejasme* | Red-white | DDS83-1 | Deng9783-1 | 3,201,497,100 | 2,906,313,000 | 90.78 | 0.03 | 95 | 88.23 | 41.48 |
| *Stellera chamaejasme* | Red-white | DDS83-10 | Deng9783-10 | 3,026,015,700 | 2,929,932,000 | 96.82 | 0.03 | 96.25 | 90.98 | 39.64 |
| *Stellera chamaejasme* | Red-white | DDS83-2 | Deng9783-2 | 3,856,044,900 | 3,769,456,200 | 97.75 | 0.03 | 95.9 | 89.84 | 40.84 |

| | | | | | | | | | | |
|---|---|---|---|---|---|---|---|---|---|---|
| *Stellera chamaejasme* | Red-white | DDS83-3 | Deng9783-3 | 3,771,792,900 | 3,626,772,000 | 96.16 | 0.03 | 95.84 | 90.15 | 39.55 |
| *Stellera chamaejasme* | Red-white | DDS83-4 | Deng9783-4 | 3,409,242,900 | 3,328,503,300 | 97.63 | 0.03 | 96.24 | 90.63 | 41.47 |
| *Stellera chamaejasme* | Red-white | DDS83-5 | Deng9783-5 | 3,912,135,300 | 3,799,294,500 | 97.12 | 0.03 | 95.82 | 89.83 | 42.27 |
| *Stellera chamaejasme* | Red-white | DDS83-6 | Deng9783-6 | 3,232,731,900 | 3,118,145,700 | 96.46 | 0.03 | 95.69 | 89.81 | 39.88 |
| *Stellera chamaejasme* | Red-white | DDS83-7 | Deng9783-7 | 4,439,429,400 | 4,327,832,700 | 97.49 | 0.03 | 95.86 | 89.8 | 40.02 |
| *Stellera chamaejasme* | Red-white | DDS83-8 | Deng9783-8 | 3,222,197,700 | 3,124,547,700 | 96.97 | 0.03 | 96.8 | 92 | 39.5 |
| *Stellera chamaejasme* | Red-white | DDS83-9 | Deng9783-9 | 3,335,555,100 | 3,220,358,100 | 96.55 | 0.03 | 95.19 | 88.6 | 41.31 |
| *Stellera chamaejasme* | Red-white | LLS5-1 | LZY004-005-1 | 3,803,906,400 | 3,541,698,600 | 93.11 | 0.03 | 96.16 | 91.4 | 39.48 |
| *Stellera chamaejasme* | Red-white | LLS5-10 | LZY004-005-10 | 5,002,508,400 | 4,778,604,600 | 95.52 | 0.03 | 96.84 | 91.98 | 42.18 |
| *Stellera chamaejasme* | Red-white | LLS5-2 | LZY004-005-2 | 3,093,300,600 | 3,093,278,400 | 100 | 0.03 | 95.63 | 90.48 | 41.98 |
| *Stellera chamaejasme* | Red-white | LLS5-3 | LZY004-005-3 | 4,172,979,900 | 3,466,704,900 | 83.08 | 0.03 | 96.08 | 90.92 | 40.19 |
| *Stellera chamaejasme* | Red-white | LLS5-4 | LZY004-005-4 | 3,357,319,200 | 3,264,913,800 | 97.25 | 0.03 | 95.59 | 90.05 | 40.33 |
| *Stellera chamaejasme* | Red-white | LLS5-5 | LZY004-005-5 | 4,873,278,900 | 4,601,533,200 | 94.42 | 0.03 | 96.53 | 91.56 | 41.47 |
| *Stellera chamaejasme* | Red-white | LLS5-6 | LZY004-005-6 | 3,156,871,800 | 3,076,561,800 | 97.46 | 0.03 | 96.47 | 91.3 | 41.16 |
| *Stellera chamaejasme* | Red-white | LLS5-7 | LZY004-005-7 | 4,526,135,400 | 4,356,397,200 | 96.25 | 0.03 | 95.42 | 89.47 | 40.09 |
| *Stellera chamaejasme* | Red-white | LLS5-8 | LZY004-005-8 | 4,775,103,900 | 4,641,288,600 | 97.2 | 0.03 | 96.87 | 92.07 | 41.32 |
| *Stellera chamaejasme* | Red-white | YMX19-3 | Yue0719-3 | 3,422,219,700 | 3,350,753,700 | 97.91 | 0.03 | 96.46 | 91.42 | 40.81 |
| *Stellera chamaejasme* | Red-white | YMX19-4 | Yue0719-4 | 3,441,657,600 | 3,371,088,300 | 97.95 | 0.03 | 96.72 | 91.7 | 40.38 |
| *Stellera chamaejasme* | Red-white | YMX19-5 | Yue0719-5 | 3,706,626,900 | 3,537,345,000 | 95.43 | 0.03 | 95.82 | 90.07 | 40.37 |
| *Stellera* | Red-white | YMX19- | Yue0719-6 | 3,218,475,30 | 3,150,964,500 | 97.9 | 0.03 | 96.7 | 91.68 | 40.31 |

| | | | | | | | | | | |
|---|---|---|---|---|---|---|---|---|---|---|
| *chamaejasme* | | 6 | | 0 | | | | | | |
| *Stellera chamaejasme* | Red-white | YNX20-1 | Yue0720-1 | 3,334,776,000 | 3,229,470,600 | 96.84 | 0.03 | 96.8 | 92.07 | 40.92 |
| *Stellera chamaejasme* | Red-white | YNX20-10 | Yue0720-10 | 3,255,889,200 | 3,153,144,300 | 96.84 | 0.03 | 96.06 | 90.55 | 40.8 |
| *Stellera chamaejasme* | Red-white | YNX20-11 | Yue0720-11 | 3,132,825,900 | 3,037,090,200 | 96.94 | 0.03 | 96.77 | 92.05 | 40.36 |
| *Stellera chamaejasme* | Red-white | YNX20-12 | Yue0720-12 | 3,046,784,700 | 2,959,580,100 | 97.14 | 0.03 | 96.43 | 91.36 | 41.18 |
| *Stellera chamaejasme* | Red-white | YNX20-2 | Yue0720-2 | 3,272,853,600 | 3,153,258,300 | 96.35 | 0.03 | 96.81 | 92.09 | 40.77 |
| *Stellera chamaejasme* | Red-white | YNX20-3 | Yue0720-3 | 3,736,743,600 | 3,597,870,000 | 96.28 | 0.03 | 96.7 | 91.88 | 40.8 |
| *Stellera chamaejasme* | Red-white | YNX20-4 | Yue0720-4 | 3,043,392,000 | 2,882,772,300 | 94.72 | 0.03 | 95.78 | 90.02 | 41.17 |
| *Stellera chamaejasme* | Red-white | YNX20-5 | Yue0720-5 | 3,143,082,600 | 3,030,585,000 | 96.42 | 0.03 | 96.62 | 91.71 | 40.36 |
| *Stellera chamaejasme* | Red-white | YNX20-6 | Yue0720-6 | 3,021,306,600 | 2,915,419,500 | 96.5 | 0.03 | 96.8 | 92.06 | 40.53 |
| *Stellera chamaejasme* | Red-white | YNX20-7 | Yue0720-7 | 3,494,291,100 | 3,309,749,100 | 94.72 | 0.03 | 95.8 | 90.03 | 39.91 |
| *Stellera chamaejasme* | Red-white | YNX20-8 | Yue0720-8 | 3,112,606,500 | 2,999,652,900 | 96.37 | 0.03 | 96.7 | 91.76 | 41.44 |
| *Stellera chamaejasme* | Red-white | YNX20-9 | Yue0720-9 | 3,358,470,300 | 3,264,092,100 | 97.19 | 0.03 | 96.5 | 91.42 | 40.47 |
| *Stellera chamaejasme* | Red-white | ZCX95-1 | ZBFC295-1 | 3,594,273,600 | 3,486,448,500 | 97 | 0.03 | 96.53 | 91.49 | 40.14 |
| *Stellera chamaejasme* | Red-white | ZCX95-2 | ZBFC295-2 | 3,694,402,200 | 3,611,972,100 | 97.77 | 0.03 | 95.98 | 89.99 | 39.8 |
| *Stellera chamaejasme* | Red-white | ZCX95-3 | ZBFC295-3 | 3,162,100,800 | 3,083,473,800 | 97.51 | 0.03 | 95.21 | 88.66 | 40.06 |
| *Stellera chamaejasme* | Red-white | ZCX95-4 | ZBFC295-4 | 3,706,187,700 | 3,612,223,200 | 97.46 | 0.03 | 95.86 | 89.84 | 40.2 |
| *Stellera chamaejasme* | Red-white | ZCX95-5 | ZBFC295-5 | 3,511,997,700 | 3,377,204,400 | 96.16 | 0.03 | 96.12 | 90.5 | 40.36 |
| *Stellera chamaejasme* | Red-white | ZCX95-6 | ZBFC295-6 | 3,405,615,300 | 3,322,074,900 | 97.55 | 0.03 | 96.03 | 90.12 | 41.06 |
| *Stellera chamaejasme* | Red-white | ZBX25-1 | ZBFC425-1 | 4,953,135,600 | 4,764,369,900 | 96.19 | 0.03 | 95.85 | 89.85 | 40.32 |

| | | | | | | | | | | |
|---|---|---|---|---|---|---|---|---|---|---|
| *Stellera chamaejasme* | Red-white | ZBX25-10 | ZBFC425-10 | 3,342,824,700 | 3,183,842,100 | 95.24 | 0.03 | 95.27 | 88.91 | 40.67 |
| *Stellera chamaejasme* | Red-white | ZBX25-11 | ZBFC425-11 | 7,038,783,000 | 2,851,214,400 | 40.51 | 0.03 | 96.19 | 90.99 | 40.19 |
| *Stellera chamaejasme* | Red-white | ZBX25-12 | ZBFC425-12 | 3,879,115,800 | 3,652,238,700 | 94.15 | 0.03 | 96.38 | 91.19 | 41.3 |
| *Stellera chamaejasme* | Red-white | ZBX25-2 | ZBFC425-2 | 3,122,601,600 | 2,968,957,500 | 95.08 | 0.03 | 96.89 | 92.24 | 41.41 |
| *Stellera chamaejasme* | Red-white | ZBX25-3 | ZBFC425-3 | 3,672,768,000 | 3,535,889,700 | 96.27 | 0.03 | 95.8 | 90.09 | 39.85 |
| *Stellera chamaejasme* | Red-white | ZBX25-4 | ZBFC425-4 | 3,645,685,500 | 3,536,750,700 | 97.01 | 0.03 | 96.01 | 90.11 | 39.77 |
| *Stellera chamaejasme* | Red-white | ZBX25-5 | ZBFC425-5 | 4,046,885,400 | 3,890,696,700 | 96.14 | 0.03 | 95.62 | 89.72 | 41 |
| *Stellera chamaejasme* | Red-white | ZBX25-6 | ZBFC425-6 | 3,579,396,000 | 3,310,406,100 | 92.49 | 0.03 | 95.77 | 90.06 | 41.16 |
| *Stellera chamaejasme* | Red-white | ZBX25-7 | ZBFC425-7 | 3,033,328,200 | 2,475,927,600 | 81.62 | 0.04 | 94.13 | 88.21 | 44.49 |
| *Stellera chamaejasme* | Red-white | ZBX25-8 | ZBFC425-8 | 4,955,545,500 | 4,661,142,900 | 94.06 | 0.03 | 95.79 | 90.06 | 41.38 |
| *Stellera chamaejasme* | Red-white | ZBX25-9 | ZBFC425-9 | 4,139,508,300 | 3,930,833,400 | 94.96 | 0.03 | 95.74 | 90 | 40.56 |
| *Stellera chamaejasme* | Pure pink | DYS25-2 | Deng9525-2 | 1,836,029,700 | 1,768,753,200 | 96.34 | 0.04 | 94.94 | 88.81 | 41.86 |
| *Stellera chamaejasme* | Pure pink | DYS25-3 | Deng9525-3 | 3,643,434,900 | 3,529,212,000 | 96.86 | 0.03 | 95.87 | 90.22 | 40.66 |
| *Stellera chamaejasme* | Pure pink | DYS25-4 | Deng9525-4 | 3,679,074,900 | 3,492,600,900 | 94.93 | 0.03 | 95.98 | 90.48 | 41.02 |
| *Stellera chamaejasme* | Pure pink | DYS25-5 | Deng9525-5 | 4,872,174,600 | 4,682,879,400 | 96.11 | 0.03 | 95.86 | 90.17 | 41.19 |
| *Stellera chamaejasme* | Pure pink | DYS25-6 | Deng9525-6 | 1,595,895,000 | 1,549,349,400 | 97.08 | 0.04 | 94.78 | 88.78 | 42.13 |
| *Stellera chamaejasme* | Pure pink | DYS25-7 | Deng9525-7 | 3,340,914,000 | 3,217,763,100 | 96.31 | 0.03 | 95.35 | 89.67 | 41.25 |
| *Stellera chamaejasme* | Pure pink | DYS25-8 | Deng9525-8 | 4,044,463,200 | 3,837,998,100 | 94.9 | 0.03 | 96.07 | 90.81 | 40.45 |
| *Stellera chamaejasme* | Pure pink | DYS28-3 | Deng9528-3 | 3,211,846,200 | 3,140,072,400 | 97.77 | 0.03 | 95.41 | 89.25 | 40.59 |
| *Stellera* | Pure pink | DYS28-4 | Deng9528-4 | 3,216,117,60 | 3,131,730,900 | 97.38 | 0.03 | 95.48 | 89.23 | 41.14 |

| | | | | | | | | | | |
|---|---|---|---|---|---|---|---|---|---|---|
| *chamaejasme* | | | | 0 | | | | | | |
| *Stellera chamaejasme* | Pure pink | DYS28-5 | Deng9528-5 | 5,394,588,300 | 5,091,186,000 | 94.38 | 0.04 | 95.29 | 89.14 | 39.52 |
| *Stellera chamaejasme* | Pure pink | DYS33-1 | Deng9533-1 | 3,263,612,100 | 2,883,810,300 | 88.36 | 0.03 | 96.17 | 91.47 | 40.86 |
| *Stellera chamaejasme* | Pure pink | DYS33-2 | Deng9533-2 | 3,521,598,900 | 3,381,686,100 | 96.03 | 0.04 | 94.36 | 87.6 | 40.95 |
| *Stellera chamaejasme* | Pure pink | DYS33-3 | Deng9533-3 | 3,127,402,200 | 2,987,979,000 | 95.54 | 0.04 | 94.25 | 87.45 | 40.95 |
| *Stellera chamaejasme* | Pure pink | DYS33-4 | Deng9533-4 | 3,383,000,400 | 3,192,415,200 | 94.37 | 0.04 | 93.88 | 86.46 | 41.67 |
| *Stellera chamaejasme* | Pure pink | DYS33-5 | Deng9533-5 | 3,001,218,600 | 2,653,362,600 | 88.41 | 0.04 | 94.7 | 88.23 | 41.5 |
| *Stellera chamaejasme* | Pure pink | DYS33-6 | Deng9533-6 | 3,769,283,700 | 3,660,723,000 | 97.12 | 0.04 | 94.31 | 87.59 | 40.66 |
| *Stellera chamaejasme* | Pure pink | DYS33-7 | Deng9533-7 | 435,121,500 | 313,333,500 | 72.01 | 0.04 | 93.08 | 85.6 | 41.23 |
| *Stellera chamaejasme* | Pure pink | DYS33-8 | Deng9533-8 | 4,072,079,700 | 3,908,985,900 | 95.99 | 0.04 | 94.28 | 87.41 | 40.92 |
| *Stellera chamaejasme* | Pure pink | DLS10-1 | Deng9810-1 | 4,597,954,800 | 3,920,618,400 | 85.27 | 0.03 | 96.31 | 91.58 | 41.49 |
| *Stellera chamaejasme* | Pure pink | DLS10-2 | Deng9810-2 | 3,966,934,200 | 3,755,638,200 | 94.67 | 0.03 | 95.41 | 89.57 | 40.62 |
| *Stellera chamaejasme* | Pure pink | DLS10-3 | Deng9810-3 | 3,228,392,700 | 3,110,802,000 | 96.36 | 0.03 | 94.84 | 87.92 | 41.36 |
| *Stellera chamaejasme* | Pure pink | DLS10-4 | Deng9810-4 | 2,950,151,400 | 2,849,582,100 | 96.59 | 0.03 | 96.77 | 91.97 | 41.89 |
| *Stellera chamaejasme* | Pure pink | DLS10-5 | Deng9810-5 | 3,547,569,600 | 3,413,667,000 | 96.23 | 0.03 | 96.72 | 91.78 | 40.33 |
| *Stellera chamaejasme* | Pure pink | DLS13-1 | Deng9813-1 | 3,525,652,800 | 3,434,803,800 | 97.42 | 0.03 | 96.4 | 91.25 | 40.38 |
| *Stellera chamaejasme* | Pure pink | DLS13-2 | Deng9813-2 | 4,746,599,400 | 4,570,898,700 | 96.3 | 0.04 | 93.75 | 86.28 | 40.48 |
| *Stellera chamaejasme* | Pure pink | DLS13-3 | Deng9813-3 | 3,109,583,100 | 2,935,590,600 | 94.4 | 0.03 | 96.66 | 91.7 | 41.56 |
| *Stellera chamaejasme* | Pure pink | DLS13-4 | Deng9813-4 | 3,263,757,000 | 3,188,502,000 | 97.69 | 0.03 | 96.73 | 91.81 | 41.25 |
| *Stellera chamaejasme* | Pure pink | DLS13-5 | Deng9813-5 | 3,428,648,400 | 3,289,488,900 | 95.94 | 0.03 | 95.47 | 89.64 | 40.06 |

| | | | | | | | | | | |
|---|---|---|---|---|---|---|---|---|---|---|
| *Stellera chamaejasme* | Pure pink | DLS13-6 | Deng9813-6 | 3,486,147,300 | 3,347,839,500 | 96.03 | 0.03 | 96.91 | 92.25 | 42.03 |
| *Stellera chamaejasme* | Pure pink | DLS13-7 | Deng9813-7 | 3,140,209,200 | 3,068,400,000 | 97.71 | 0.03 | 96.72 | 91.75 | 40.73 |
| *Stellera chamaejasme* | Pure pink | DLS13-8 | Deng9813-8 | 3,248,383,800 | 3,165,246,300 | 97.44 | 0.03 | 96.61 | 91.67 | 40.48 |
| *Stellera chamaejasme* | Yellow-white | CZX16-1 | Chen0016-1 | 4,453,794,600 | 4,149,648,300 | 93.17 | 0.03 | 96.5 | 91.46 | 40.37 |
| *Stellera chamaejasme* | Yellow-white | CZX16-2 | Chen0016-2 | 5,642,887,800 | 5,348,057,100 | 94.78 | 0.04 | 95.09 | 89.23 | 40.09 |
| *Stellera chamaejasme* | Yellow-white | CZX16-3 | Chen0016-3 | 3,172,202,400 | 3,024,594,300 | 95.35 | 0.03 | 95.75 | 90.31 | 41.5 |
| *Stellera chamaejasme* | Yellow-white | CZX16-4 | Chen0016-4 | 3,574,028,700 | 3,470,618,400 | 97.11 | 0.03 | 95.74 | 89.86 | 40.38 |
| *Stellera chamaejasme* | Yellow-white | CZX16-5 | Chen0016-5 | 4,800,411,600 | 4,375,990,200 | 91.16 | 0.03 | 95.71 | 89.82 | 41.91 |
| *Stellera chamaejasme* | Yellow-white | CZX16-6 | Chen0016-6 | 3,609,924,300 | 3,533,472,000 | 97.88 | 0.03 | 95.66 | 89.92 | 41.34 |
| *Stellera chamaejasme* | Yellow-white | CZX16-7 | Chen0016-7 | 3,251,660,100 | 3,126,702,900 | 96.16 | 0.03 | 95.81 | 89.97 | 41.32 |
| *Stellera chamaejasme* | Yellow-white | CZX16-8 | Chen0016-8 | 2,156,046,000 | 2,033,022,900 | 94.29 | 0.03 | 95.52 | 89.61 | 41.53 |
| *Stellera chamaejasme* | Yellow-white | DKS24-1 | Deng9524-1 | 3,693,695,100 | 3,492,073,200 | 94.54 | 0.04 | 94.28 | 87.45 | 40.09 |
| *Stellera chamaejasme* | Yellow-white | DKS24-10 | Deng9524-10 | 3,004,965,000 | 2,800,370,100 | 93.19 | 0.03 | 96.52 | 92.01 | 38.92 |
| *Stellera chamaejasme* | Yellow-white | DKS24-2 | Deng9524-2 | 3,845,508,300 | 3,602,776,200 | 93.69 | 0.04 | 94.22 | 87.36 | 41.36 |
| *Stellera chamaejasme* | Yellow-white | DKS24-4 | Deng9524-4 | 3,316,375,500 | 3,152,227,200 | 95.05 | 0.03 | 96.45 | 91.75 | 39.97 |
| *Stellera chamaejasme* | Yellow-white | DKS35-1 | Deng9535-1 | 3,448,819,500 | 3,354,774,000 | 97.27 | 0.03 | 95.77 | 89.84 | 40.83 |
| *Stellera chamaejasme* | Yellow-white | DKS35-2 | Deng9535-2 | 3,743,460,000 | 3,576,734,700 | 95.55 | 0.03 | 95.62 | 89.59 | 41.1 |
| *Stellera chamaejasme* | Yellow-white | DKS35-3 | Deng9535-3 | 3,289,871,100 | 3,093,426,600 | 94.03 | 0.03 | 95.78 | 89.91 | 40.85 |
| *Stellera chamaejasme* | Yellow-white | DKS35-5 | Deng9535-5 | 4,115,716,500 | 3,969,381,300 | 96.44 | 0.03 | 95.67 | 89.59 | 41.02 |
| *Stellera* | Yellow-white | DKS35-6 | Deng9535-6 | 3,743,460,00 | 3,576,893,400 | 95.55 | 0.03 | 95.63 | 89.47 | 40.72 |

| | | | | | | | | | | |
|---|---|---|---|---|---|---|---|---|---|---|
| *chamaejasme* | | | | 0 | | | | | | |
| *Stellera chamaejasme* | Yellow-white | DMX03-1 | Deng9703-1 | 3,486,384,600 | 3,413,860,500 | 97.92 | 0.03 | 95.94 | 89.88 | 41.08 |
| *Stellera chamaejasme* | Yellow-white | DMX03-4 | Deng9703-4 | 4,404,052,200 | 4,287,846,300 | 97.36 | 0.03 | 96.21 | 90.75 | 40.56 |
| *Stellera chamaejasme* | Yellow-white | DMX03-7 | Deng9703-7 | 3,305,228,400 | 3,208,438,200 | 97.07 | 0.03 | 96.13 | 90.65 | 41.05 |
| *Stellera chamaejasme* | Yellow-white | DMX03-8 | Deng9703-8 | 4,802,932,500 | 4,559,589,600 | 94.93 | 0.03 | 96.03 | 90.12 | 41.2 |
| *Stellera chamaejasme* | Yellow-white | DMX15-1 | Deng9715-1 | 3,198,624,000 | 3,038,661,300 | 95 | 0.03 | 96.03 | 90.62 | 41.45 |
| *Stellera chamaejasme* | Yellow-white | DMX15-2 | Deng9715-2 | 3,961,627,800 | 3,744,813,000 | 94.53 | 0.03 | 95.73 | 89.96 | 41.17 |
| *Stellera chamaejasme* | Yellow-white | DMX15-3 | Deng9715-3 | 3,417,001,200 | 3,281,967,600 | 96.05 | 0.03 | 96.05 | 90.75 | 40.78 |
| *Stellera chamaejasme* | Yellow-white | DMX15-4 | Deng9715-4 | 2,983,998,000 | 2,867,157,600 | 96.08 | 0.03 | 96.35 | 91.2 | 40.94 |
| *Stellera chamaejasme* | Yellow-white | DMX15-5 | Deng9715-5 | 4,403,072,400 | 4,185,761,700 | 95.06 | 0.03 | 95.82 | 90.19 | 40.48 |
| *Stellera chamaejasme* | Yellow-white | DMX15-6 | Deng9715-6 | 3,350,170,200 | 3,204,712,800 | 95.66 | 0.03 | 96.13 | 90.91 | 40.47 |
| *Stellera chamaejasme* | Yellow-white | DMX15-7 | Deng9715-7 | 4,231,117,800 | 4,137,555,600 | 97.79 | 0.03 | 95.97 | 89.9 | 40.8 |
| *Stellera chamaejasme* | Yellow-white | DMX15-8 | Deng9715-8 | 3,644,195,700 | 3,559,644,300 | 97.68 | 0.03 | 95.98 | 89.94 | 41.12 |
| *Stellera chamaejasme* | Yellow-white | DGS47-1 | Deng9847-1 | 3,635,957,400 | 3,577,809,600 | 98.4 | 0.03 | 95.86 | 90.9 | 40.71 |
| *Stellera chamaejasme* | Yellow-white | DGS47-2 | Deng9847-2 | 3,720,510,600 | 3,699,852,600 | 99.44 | 0.04 | 95.37 | 90 | 41.3 |
| *Stellera chamaejasme* | Yellow-white | DGS47-3 | Deng9847-3 | 3,285,485,100 | 3,199,882,800 | 97.39 | 0.03 | 96.4 | 91.18 | 40.74 |
| *Stellera chamaejasme* | Yellow-white | DGS47-4 | Deng9847-4 | 3,847,006,200 | 3,733,631,400 | 97.05 | 0.03 | 96.88 | 92.16 | 41.14 |
| *Stellera chamaejasme* | Yellow-white | DGS47-5 | Deng9847-5 | 4,104,339,300 | 4,078,463,700 | 99.37 | 0.04 | 95.44 | 90.09 | 41.49 |
| *Stellera chamaejasme* | Yellow-white | DGS47-6 | Deng9847-6 | 3,028,478,400 | 2,930,835,000 | 96.78 | 0.03 | 96.44 | 91.31 | 40.19 |
| *Stellera chamaejasme* | Yellow-white | DGS47-7 | Deng9847-7 | 3,553,972,800 | 3,522,812,100 | 99.12 | 0.04 | 95.82 | 90.72 | 40.41 |

| | | | | | | | | | | |
|---|---|---|---|---|---|---|---|---|---|---|
| *Stellera chamaejasme* | Yellow-white | DGS47-9 | Deng9847-9 | 3,962,287,200 | 3,641,107,200 | 91.89 | 0.03 | 96.64 | 91.75 | 39.95 |
| *Stellera chamaejasme* | Yellow-white | ZMX-1 | ZYZhou-1 | 4,115,941,200 | 3,908,685,600 | 94.96 | 0.03 | 96.45 | 91.38 | 41.31 |
| *Stellera chamaejasme* | Yellow-white | ZMX-11 | ZYZhou-11 | 3,188,538,300 | 3,028,469,700 | 94.98 | 0.03 | 96.81 | 91.98 | 40.73 |
| *Stellera chamaejasme* | Yellow-white | ZMX-12 | ZYZhou-12 | 3,415,930,200 | 3,249,045,300 | 95.11 | 0.03 | 96.75 | 91.84 | 41.2 |
| *Stellera chamaejasme* | Yellow-white | ZMX-3 | ZYZhou-3 | 3,877,823,700 | 3,698,681,100 | 95.38 | 0.03 | 96.84 | 92.06 | 40 |
| *Stellera chamaejasme* | Yellow-white | ZMX-4 | ZYZhou-4 | 3,459,105,000 | 3,314,609,400 | 95.82 | 0.03 | 96.91 | 92.29 | 40.95 |
| *Stellera chamaejasme* | Yellow-white | ZMX-5 | ZYZhou-5 | 3,943,720,500 | 3,776,125,200 | 95.75 | 0.03 | 96.94 | 92.23 | 40.94 |
| *Stellera chamaejasme* | Yellow-white | ZMX-6 | ZYZhou-6 | 3,100,557,300 | 2,971,787,100 | 95.85 | 0.03 | 97 | 92.44 | 41 |
| *Stellera chamaejasme* | Yellow-white | ZMX-8 | ZYZhou-8 | 4,307,643,000 | 4,176,705,600 | 96.96 | 0.03 | 96.58 | 91.56 | 41.15 |
| *Stellera chamaejasme* | Yellow-white | ZMX-9 | ZYZhou-9 | 3,372,429,300 | 3,266,104,200 | 96.85 | 0.03 | 96.51 | 91.45 | 41.01 |
| *Stellera chamaejasme* | Pure yellow | CSY9-1 | Chen0009-1 | 4,338,585,900 | 4,149,704,400 | 95.65 | 0.03 | 96.04 | 90.62 | 40.84 |
| *Stellera chamaejasme* | Pure yellow | CSY9-10 | Chen0009-10 | 3,624,307,200 | 3,457,748,100 | 95.4 | 0.03 | 95.6 | 89.62 | 40.48 |
| *Stellera chamaejasme* | Pure yellow | CSY9-3 | Chen0009-3 | 4,688,588,700 | 4,310,737,500 | 91.94 | 0.03 | 95.44 | 89.55 | 42.52 |
| *Stellera chamaejasme* | Pure yellow | CSY9-4 | Chen0009-4 | 4,501,227,000 | 4,263,168,000 | 94.71 | 0.03 | 95.6 | 89.78 | 40.73 |
| *Stellera chamaejasme* | Pure yellow | CSY9-5 | Chen0009-5 | 1,740,017,100 | 1,459,449,000 | 83.88 | 0.04 | 93.09 | 85.47 | 41.53 |
| *Stellera chamaejasme* | Pure yellow | CSY9-6 | Chen0009-6 | 5,792,020,800 | 5,443,385,400 | 93.98 | 0.04 | 93.53 | 86.06 | 41.84 |
| *Stellera chamaejasme* | Pure yellow | CSY9-7 | Chen0009-7 | 3,198,561,600 | 3,049,374,600 | 95.34 | 0.03 | 95.22 | 88.78 | 41.59 |
| *Stellera chamaejasme* | Pure yellow | CSY9-8 | Chen0009-8 | 3,667,136,700 | 3,159,592,500 | 86.16 | 0.04 | 94.83 | 89.22 | 40.8 |
| *Stellera chamaejasme* | Pure yellow | CSY9-9 | Chen0009-9 | 3,737,137,200 | 3,531,507,000 | 94.5 | 0.03 | 95.66 | 89.81 | 41.83 |
| *Stellera* | Pure yellow | CLY15-1 | Chen0015-1 | 3,704,745,90 | 3,535,123,200 | 95.42 | 0.03 | 95.59 | 89.81 | 40.76 |

| | | | | | | | | | | |
|---|---|---|---|---|---|---|---|---|---|---|
| *chamaejasme* | | | | 0 | | | | | | |
| *Stellera chamaejasme* | Pure yellow | CLY15-10 | Chen0015-10 | 4,375,714,200 | 4,144,391,400 | 94.71 | 0.04 | 94.33 | 87.42 | 39.7 |
| *Stellera chamaejasme* | Pure yellow | CLY15-2 | Chen0015-2 | 3,872,289,900 | 3,753,764,400 | 96.94 | 0.03 | 95.77 | 90.14 | 40.78 |
| *Stellera chamaejasme* | Pure yellow | CLY15-3 | Chen0015-3 | 3,661,755,900 | 3,513,237,900 | 95.94 | 0.03 | 95.61 | 89.84 | 40 |
| *Stellera chamaejasme* | Pure yellow | CLY15-4 | Chen0015-4 | 3,694,119,300 | 3,461,019,900 | 93.69 | 0.04 | 94.44 | 87.59 | 41.06 |
| *Stellera chamaejasme* | Pure yellow | CLY15-5 | Chen0015-5 | 3,290,056,800 | 3,020,361,300 | 91.8 | 0.03 | 96.3 | 91.52 | 40.02 |
| *Stellera chamaejasme* | Pure yellow | CLY15-6 | Chen0015-6 | 3,630,227,100 | 3,247,975,800 | 89.47 | 0.04 | 94.31 | 87.53 | 40.94 |
| *Stellera chamaejasme* | Pure yellow | CLY15-7 | Chen0015-7 | 4,822,762,500 | 4,561,500,300 | 94.58 | 0.03 | 96.92 | 92.38 | 40.18 |
| *Stellera chamaejasme* | Pure yellow | CLY15-8 | Chen0015-8 | 3,901,402,500 | 2,919,110,100 | 74.82 | 0.03 | 95.87 | 90.98 | 39.84 |
| *Stellera chamaejasme* | Pure yellow | CLY15-9 | Chen0015-9 | 1,674,454,500 | 1,543,710,600 | 92.19 | 0.04 | 93.72 | 86.41 | 41.06 |
| *Stellera chamaejasme* | Pure yellow | DLY39-1 | Deng8939-1 | 6,338,509,200 | 5,903,569,200 | 93.14 | 0.04 | 95.06 | 89.26 | 40.88 |
| *Stellera chamaejasme* | Pure yellow | DLY39-10 | Deng8939-10 | 3,658,836,000 | 2,415,095,700 | 66.01 | 0.04 | 93.37 | 87.05 | 42.73 |
| *Stellera chamaejasme* | Pure yellow | DLY39-2 | Deng8939-2 | 4,608,112,200 | 3,195,641,400 | 69.35 | 0.04 | 93.1 | 86.52 | 44.38 |
| *Stellera chamaejasme* | Pure yellow | DLY39-3 | Deng8939-3 | 3,514,440,000 | 3,383,447,100 | 96.27 | 0.03 | 95.75 | 90.07 | 40.69 |
| *Stellera chamaejasme* | Pure yellow | DLY39-4 | Deng8939-4 | 4,215,881,400 | 4,103,279,700 | 97.33 | 0.03 | 95.82 | 90.2 | 41.69 |
| *Stellera chamaejasme* | Pure yellow | DLY39-5 | Deng8939-5 | 3,494,969,400 | 2,304,907,800 | 65.95 | 0.04 | 92.97 | 86.41 | 44.03 |
| *Stellera chamaejasme* | Pure yellow | DLY39-6 | Deng8939-6 | 3,108,194,400 | 2,971,491,900 | 95.6 | 0.03 | 95.53 | 89.64 | 41.37 |
| *Stellera chamaejasme* | Pure yellow | DLY39-7 | Deng8939-7 | 3,556,806,900 | 2,419,312,800 | 68.02 | 0.04 | 93.79 | 87.58 | 41.28 |
| *Stellera chamaejasme* | Pure yellow | DLY39-8 | Deng8939-8 | 3,116,796,900 | 1,759,206,000 | 56.44 | 0.05 | 89.1 | 81.16 | 48.27 |
| *Stellera chamaejasme* | Pure yellow | DLY39-9 | Deng8939-9 | 3,251,076,900 | 3,093,090,300 | 95.14 | 0.03 | 95.75 | 89.98 | 41.35 |

| | | | | | | | | | | |
|---|---|---|---|---|---|---|---|---|---|---|
| *Stellera chamaejasme* | Pure yellow | DWY99-1 | Deng8999-1 | 3,079,706,100 | 2,985,707,700 | 96.95 | 0.03 | 95.43 | 89.35 | 40.45 |
| *Stellera chamaejasme* | Pure yellow | DWY99-10 | Deng8999-10 | 3,110,756,700 | 2,904,991,200 | 93.39 | 0.03 | 96.69 | 91.98 | 41.47 |
| *Stellera chamaejasme* | Pure yellow | DWY99-2 | Deng8999-2 | 4,170,803,100 | 2,877,803,700 | 69 | 0.04 | 93.22 | 86.91 | 42.51 |
| *Stellera chamaejasme* | Pure yellow | DWY99-3 | Deng8999-3 | 4,048,783,800 | 3,758,392,200 | 92.83 | 0.03 | 95.96 | 90.61 | 41.26 |
| *Stellera chamaejasme* | Pure yellow | DWY99-4 | Deng8999-4 | 2,564,626,200 | 2,123,205,000 | 82.79 | 0.04 | 93.33 | 86.93 | 44.5 |
| *Stellera chamaejasme* | Pure yellow | DWY99-5 | Deng8999-5 | 4,545,090,000 | 4,260,575,700 | 93.74 | 0.04 | 94.47 | 88.44 | 41.9 |
| *Stellera chamaejasme* | Pure yellow | DWY99-6 | Deng8999-6 | 3,344,549,100 | 2,928,506,100 | 87.56 | 0.04 | 95.18 | 89.51 | 41.21 |
| *Stellera chamaejasme* | Pure yellow | DWY99-7 | Deng8999-7 | 7,593,824,400 | 7,179,154,200 | 94.54 | 0.04 | 95.31 | 89.55 | 40.31 |
| *Stellera chamaejasme* | Pure yellow | DWY99-8 | Deng8999-8 | 4,041,486,900 | 3,951,043,500 | 97.76 | 0.03 | 95.8 | 90.25 | 41.55 |
| *Stellera chamaejasme* | Pure yellow | DWY99-9 | Deng8999-9 | 3,680,410,800 | 3,513,967,500 | 95.48 | 0.03 | 95.82 | 90.2 | 41.26 |
| *Stellera chamaejasme* | Pure yellow | DLY40-1 | Deng9040-1 | 4,478,875,500 | 4,390,787,400 | 98.03 | 0.03 | 95.23 | 88.94 | 41.14 |
| *Stellera chamaejasme* | Pure yellow | DLY40-10 | Deng9040-10 | 4,599,324,000 | 4,473,245,400 | 97.26 | 0.03 | 95.81 | 90.12 | 40.95 |
| *Stellera chamaejasme* | Pure yellow | DLY40-2 | Deng9040-2 | 4,517,864,400 | 4,357,637,100 | 96.45 | 0.03 | 95.77 | 90.06 | 40.89 |
| *Stellera chamaejasme* | Pure yellow | DLY40-3 | Deng9040-3 | 3,955,066,200 | 3,889,437,300 | 98.34 | 0.03 | 95.09 | 88.75 | 41.09 |
| *Stellera chamaejasme* | Pure yellow | DLY40-4 | Deng9040-4 | 3,253,315,800 | 3,165,383,700 | 97.3 | 0.03 | 95.38 | 89.19 | 40.97 |
| *Stellera chamaejasme* | Pure yellow | DLY40-5 | Deng9040-5 | 3,822,755,400 | 3,647,760,300 | 95.42 | 0.03 | 95.8 | 90.19 | 41.12 |
| *Stellera chamaejasme* | Pure yellow | DLY40-6 | Deng9040-6 | 3,785,194,200 | 3,667,702,500 | 96.9 | 0.03 | 95.72 | 89.87 | 40.76 |
| *Stellera chamaejasme* | Pure yellow | DLY40-7 | Deng9040-7 | 3,081,477,900 | 2,829,578,400 | 91.83 | 0.03 | 95.85 | 90.6 | 39.89 |
| *Stellera chamaejasme* | Pure yellow | DLY40-8 | Deng9040-8 | 3,146,710,200 | 3,048,523,200 | 96.88 | 0.03 | 94.84 | 88.31 | 41.05 |
| *Stellera* | Pure yellow | DLY40-9 | Deng9040-9 | 3,287,904,90 | 3,181,449,000 | 96.76 | 0.03 | 95.49 | 89.53 | 41.2 |

| | | | | | | | | | | |
|---|---|---|---|---|---|---|---|---|---|---|
| *chamaejasme* | | | | 0 | | | | | | |
| *Stellera chamaejasme* | Pure yellow | DYY27-1 | Deng9127-1 | 3,668,531,700 | 3,450,046,200 | 94.04 | 0.03 | 95.88 | 90.38 | 41.39 |
| *Stellera chamaejasme* | Pure yellow | DYY27-2 | Deng9127-2 | 4,745,922,900 | 4,520,175,300 | 95.24 | 0.03 | 95.88 | 90.28 | 41.74 |
| *Stellera chamaejasme* | Pure yellow | DYY27-3 | Deng9127-3 | 3,409,119,000 | 2,885,270,400 | 84.63 | 0.03 | 96.22 | 91.5 | 40.75 |
| *Stellera chamaejasme* | Pure yellow | DYY27-4 | Deng9127-4 | 4,325,064,000 | 4,119,565,200 | 95.25 | 0.03 | 95.41 | 89.34 | 41.81 |
| *Stellera chamaejasme* | Pure yellow | DYY27-5 | Deng9127-5 | 3,627,124,500 | 3,469,862,100 | 95.66 | 0.03 | 95.5 | 89.82 | 40.21 |
| *Stellera chamaejasme* | Pure yellow | DYY27-6 | Deng9127-6 | 3,515,490,300 | 3,384,054,000 | 96.26 | 0.03 | 95.53 | 90.08 | 41.04 |
| *Stellera chamaejasme* | Pure yellow | DYY27-7 | Deng9127-7 | 3,176,373,900 | 3,013,587,900 | 94.88 | 0.03 | 95.72 | 90.14 | 40.45 |
| *Stellera chamaejasme* | Pure yellow | DLY69-1 | Deng9569-1 | 3,305,442,300 | 3,232,428,900 | 97.79 | 0.03 | 95.85 | 89.91 | 40.52 |
| *Stellera chamaejasme* | Pure yellow | DLY69-2 | Deng9569-2 | 3,764,559,000 | 3,598,658,400 | 95.59 | 0.03 | 95.16 | 89.17 | 40.19 |
| *Stellera chamaejasme* | Pure yellow | DLY69-3 | Deng9569-3 | 4,294,956,900 | 4,079,232,300 | 94.98 | 0.03 | 94.96 | 88.83 | 40.56 |
| *Stellera chamaejasme* | Pure yellow | DLY69-4 | Deng9569-4 | 3,069,584,400 | 2,918,906,700 | 95.09 | 0.03 | 95.05 | 88.93 | 40.16 |
| *Stellera chamaejasme* | Pure yellow | DLY69-5 | Deng9569-5 | 3,265,851,900 | 3,021,408,900 | 92.52 | 0.03 | 95.06 | 89.03 | 39.71 |
| *Stellera chamaejasme* | Pure yellow | DLY69-6 | Deng9569-6 | 5,744,587,200 | 5,528,388,000 | 96.24 | 0.03 | 95.47 | 89.22 | 40.77 |
| *Stellera chamaejasme* | Pure yellow | DLY69-8 | Deng9569-8 | 4,444,466,100 | 4,204,415,100 | 94.6 | 0.03 | 95.38 | 89.21 | 40.75 |
| *Stellera chamaejasme* | Pure yellow | DSY1-1 | Deng9601-1 | 11,565,969,000 | 5,252,239,800 | 45.41 | 0.03 | 95.59 | 89.59 | 40.8 |
| *Stellera chamaejasme* | Pure yellow | DSY1-2 | Deng9601-2 | 3,977,664,300 | 3,349,495,800 | 84.21 | 0.03 | 95.61 | 89.65 | 41.35 |
| *Stellera chamaejasme* | Pure yellow | DSY1-3 | Deng9601-3 | 3,410,224,800 | 3,311,178,900 | 97.1 | 0.03 | 95.07 | 88.49 | 40.47 |
| *Stellera chamaejasme* | Pure yellow | DSY1-7 | Deng9601-7 | 5,777,049,900 | 5,258,548,800 | 91.02 | 0.03 | 95.72 | 89.81 | 40.45 |
| *Stellera chamaejasme* | Pure yellow | DSY1-8 | Deng9601-8 | 3,065,986,200 | 2,965,373,100 | 96.72 | 0.04 | 94.89 | 88.25 | 40.61 |

| | | | | | | | | | | |
|---|---|---|---|---|---|---|---|---|---|---|
| *Stellera chamaejasme* | Pure yellow | NMS3-1 | Niuma003-1 | 3,117,773,10<br>0 | 2,762,506,800 | 88.61 | 0.03 | 96.81 | 92.06 | 42.33 |
| *Stellera chamaejasme* | Pure yellow | NMS3-2 | Niuma003-2 | 3,860,993,70<br>0 | 3,698,373,600 | 95.79 | 0.03 | 96.53 | 91.52 | 40.45 |
| *Stellera chamaejasme* | Pure yellow | NMS3-4 | Niuma003-4 | 3,667,185,00<br>0 | 3,541,804,500 | 96.58 | 0.03 | 96.81 | 92.19 | 41.59 |

**Table S2.** Population summary statistics calculated for variant positions of 23,215 restrictions site-associated DNA sequencing (RAD-seq) loci.

| Code | Population | Co-ordinates | Alt. (m) | Locality | Samples (n) | Private alleles | *H*o | *H*e | π | $F_{IS}$ |
|---|---|---|---|---|---|---|---|---|---|---|
| **Red-white morph** | | | | | 103 | 25 | 0.07085 | 0.07756 | 0.07795 | 0.09063 |
| rwC18 | Chen0018 | 32°11′55.09″N 96°23′58.86″E | 3717 | Nangqên County, Qinghai Province | 3 | 38 | 0.19748 | 0.1219 | 0.14627 | -0.08662 |
| rwC19 | Chen0019 | 32°04′37.83″N 96°24′18.39″E | 4220 | Nangqên County, Qinghai Province | 3 | 42 | 0.08076 | 0.04649 | 0.05579 | -0.04194 |
| rwC21 | Chen0021 | 33°19′55.64″N 96°50′23.34″E | 3661 | Chindu County, Qinghai Province | 6 | 266 | 0.09797 | 0.06971 | 0.07701 | -0.0367 |
| rwC22 | Chen0022 | 31°34′46.59″N 100°23′44.08″E | 3512 | Luhuo County, Sichuan Province | 12 | 290 | 0.10181 | 0.07764 | 0.0814 | -0.00521 |
| rwL05 | LZY004005 | 31°43′15.06″N 100°43′15.68″E | 3685 | Luhuo County, Sichuan Province | 9 | 56 | 0.11293 | 0.06807 | 0.07239 | -0.07856 |
| rwY19 | Yue0719 | 29°50′6.396″N 91°43′20.388″E | 3822 | Maizhokunggar County, Xizang Province | 4 | 55 | 0.10324 | 0.06236 | 0.07127 | -0.05728 |
| rwY20 | Yue0720 | 29°43′27.98″N 89°14′3.77″E | 4088 | Namling County, Xizang Province | 12 | 78 | 0.10266 | 0.06442 | 0.06752 | -0.0576 |
| rwZ95 | ZBFC295 | 31°59′02.48″N 87°48′22.66″E | 4603 | Co'nyi County, Xizang Province | 6 | 96 | 0.0969 | 0.05801 | 0.06385 | -0.06169 |
| rwZ25 | ZBFC425 | 31°24′20.14″N 90°51′24.53″E | 4532 | Baingoin County, Xizang Province | 12 | 86 | 0.11014 | 0.06724 | 0.07056 | -0.07695 |
| rwD33 | Deng9733 | 31°15′38.31″N 97°9′45.85″E | 3246 | Changdu City, Xizang Province | 8 | 53 | 0.10167 | 0.06329 | 0.06792 | -0.0647 |
| rwD45 | Deng9745 | 31°28′04.63″N 97°11′54.57″E | 3338 | Changdu City, Xizang Province | 10 | 112 | 0.10116 | 0.07587 | 0.08049 | -0.03613 |
| rwD72 | Deng9772 | 31°39′03.42″N 98°22′59.02″E | 3376.3 | Jiangda County, Xizang Province | 8 | 34 | 0.08456 | 0.05029 | 0.05396 | -0.05839 |
| rwD83 | Deng9783 | 31°52′06.88″N 99°05′27.18″E | 4050 | Dêgê County, Sichuan Province | 10 | 137 | 0.11096 | 0.06815 | 0.07229 | -0.0743 |
| **Pure pink morph** | | | | | 31 | 0 | 0.07895 | 0.06324 | 0.06439 | -0.03252 |
| ppD10 | Deng9810 | 30°27′34.60″N 100°18′07.87″E | 3068 | Litang County, Sichuan Province | 5 | 61 | 0.1624 | 0.10179 | 0.11497 | -0.08543 |
| ppD13 | Deng9813 | 30°21′43.93″N 100°17′41.13″E | 3399.82 | Litang County, Sichuan Province | 8 | 31 | 0.15085 | 0.09589 | 0.10286 | -0.09038 |
| ppD25 | Deng9525 | 30°32′32.86″N 101°14′04.13″E | 3152.74 | Yajiang County, Sichuan Province | 7 | 49 | 0.1603 | 0.09653 | 0.1047 | -0.10399 |

| Code | Population | Co-ordinates | Alt. (m) | Locality | Samples (n) | Private alleles | *Ho* | *He* | π | $F_{IS}$ |
|---|---|---|---|---|---|---|---|---|---|---|
| ppD28 | Deng9528 | 30°02′32.99″N 101°14′04.22″E | 3150 | Yajiang County, Sichuan Province | 3 | 39 | 0.19577 | 0.11769 | 0.14123 | -0.09204 |
| ppD33 | Deng9533 | 30°05′17.47″N 101°0.33′0″E | 2587.81 | Yajiang County, Sichuan Province | 8 | 15 | 0.12273 | 0.08787 | 0.09438 | -0.05865 |
| **Yellow-white morph** | | | | | 46 | 7 | 0.05696 | 0.05934 | 0.05994 | 0.04703 |
| ywC16 | Chen0016 | 29°56′13.23″N 97°26′30.37″E | 4310 | Zogang County, Xizang Province | 8 | 96 | 0.114 | 0.07061 | 0.07574 | -0.07322 |
| ywZYZ | ZYZhou | 29°42′56.94″N 98°28′17.87″E | 3905 | Mangkang County, Xizang Province | 9 | 52 | 0.12272 | 0.07545 | 0.07912 | -0.0771 |
| ywD24 | Deng9524 | 30°04′42.09″N 101°37′43.61″E | 3583.08 | Kangding, Sichuan Province | 4 | 54 | 0.15719 | 0.10301 | 0.11181 | -0.08693 |
| ywD35 | Deng9535 | 29°40′45.26″N 101°25′31.99″E | 3177.26 | Kangding, Sichuan Province | 5 | 90 | 0.17695 | 0.10403 | 0.1145 | -0.11735 |
| ywD15 | Deng9715 | 29°48′07.99″N 98°32′0.33″E | 4071.49 | Mangkang County, Xizang Province | 8 | 66 | 0.08347 | 0.05104 | 0.05476 | -0.05473 |
| ywD03 | Deng9703 | 29°43′37.84″N 98°37′46.35″E | 3980.1 | Mangkang County, Xizang Province | 4 | 89 | 0.1301 | 0.08042 | 0.09084 | -0.07123 |
| ywD47 | Deng9847 | 29°17′06.89″N 100°04′47.81″E | 4032 | Daocheng County, Sichuan Province | 8 | 79 | 0.06744 | 0.04221 | 0.04526 | -0.04317 |
| **Pure yellow morph** | | | | | 71 | 23 | 0.05321 | 0.07726 | 0.07785 | 0.1327 |
| yyN03 | Niuma003 | 27°41′11.34″N 101°13′21.33″E | 3215 | Muli County, Sichuan Province | 3 | 27 | 0.17645 | 0.1032 | 0.12385 | -0.08815 |
| yyC09 | Chen0009 | 27°35′17.34″N 99°47′28.40″E | 3248 | Shangri-la County, Yunnan Province | 9 | 138 | 0.09721 | 0.09248 | 0.09847 | 0.00552 |
| yyC15 | Chen0015 | 27°09′12.81″N 100°15′25.74″E | 3010 | Yulong County, Yunnan Province | 10 | 96 | 0.15414 | 0.1081 | 0.11464 | -0.0731 |
| yyD39 | Deng8939 | 26°40′52.84″N 99°55′22.65″E | 2700 | Yulong County, Yunnan Province | 10 | 87 | 0.10862 | 0.10047 | 0.10671 | -0.00049 |
| yyD99 | Deng8999 | 27°08′09.53″N 99°17′12.76″E | 2644 | Weixi County, Yunnan Province | 10 | 107 | 0.10622 | 0.08047 | 0.08535 | -0.03953 |
| yyD40 | Deng9040 | 26°27′22.99″N 99°29′51.276″E | 3161 | Lanping County, Yunnan Province | 10 | 147 | 0.09816 | 0.07449 | 0.07896 | -0.03246 |
| yyD27 | Deng9127 | 25°51′23.45″N 99°16′03.09″E | 2642 | Yunlong County, Yunnan Province | 7 | 23 | 0.18486 | 0.10627 | 0.11525 | -0.1313 |
| yyD69 | Deng9569 | 27°04′00.89″N 100°15′47.61″E | 2966.71 | Yulong County, Yunnan Province | 7 | 57 | 0.11634 | 0.07642 | 0.08282 | -0.05519 |

| Code | Population | Co-ordinates | Alt. (m) | Locality | Samples (n) | Private alleles | ***H*o** | ***H*e** | **π** | **$F_{IS}$** |
|---|---|---|---|---|---|---|---|---|---|---|
| yyD01 | Deng9601 | 27°29′17.90″N 99°48′18.57″E | 3155.37 | Shangri-la County, Yunnan Province | 5 | 94 | 0.16454 | 0.10999 | 0.12426 | -0.07156 |

Alt., altitude (meter); n, number of individuals; *H*o; Mean observed heterozygosity; *H*e, Mean expected heterozygosity; π, averaged nucleotide diversity; $F_{IS}$, the inbreeding coefficient of an individual (I) relative to the subpopulation (S)

**Table S3.** Population summary statistics calculated for all positions of 23,215 restrictions site-associated DNA sequencing (RAD-seq) loci.

| Code | Population | n | Total Sites | Variant Sites | Polymorphic Sites | *H*o | *H*e | π | $F_{IS}$ |
|---|---|---|---|---|---|---|---|---|---|
| **Red-white morph** | | 103 | 278341 | 173 | 113 | 0.00004 | 0.00005 | 0.00005 | 0.00006 |
| rwC18 | Chen0018 | 3 | 764958 | 1190 | 373 | 0.00031 | 0.00019 | 0.00023 | -0.00013 |
| rwC19 | Chen0019 | 3 | 2520176 | 2782 | 301 | 0.00009 | 0.00005 | 0.00006 | -0.00005 |
| rwC21 | Chen0021 | 6 | 2630132 | 2976 | 663 | 0.00011 | 0.00008 | 0.00009 | -0.00004 |
| rwC22 | Chen0022 | 12 | 2008284 | 2266 | 763 | 0.00011 | 0.00009 | 0.00009 | -0.00001 |
| rwL05 | LZY004005 | 9 | 1239001 | 1427 | 304 | 0.00013 | 0.00008 | 0.00008 | -0.00009 |
| rwY19 | Yue0719 | 4 | 1736232 | 1988 | 323 | 0.00012 | 0.00007 | 0.00008 | -0.00007 |
| rwY20 | Yue0720 | 12 | 2081791 | 2330 | 512 | 0.00011 | 0.00007 | 0.00008 | -0.00006 |
| rwZ95 | ZBFC295 | 6 | 2711971 | 3143 | 506 | 0.00011 | 0.00007 | 0.00007 | -0.00007 |
| rwZ25 | ZBFC425 | 12 | 1773115 | 1872 | 445 | 0.00012 | 0.00007 | 0.00007 | -0.00008 |
| rwD33 | Deng9733 | 8 | 1548541 | 1688 | 347 | 0.00011 | 0.00007 | 0.00007 | -0.00007 |
| rwD45 | Deng9745 | 10 | 2208204 | 2363 | 600 | 0.00011 | 0.00008 | 0.00009 | -0.00004 |
| rwD72 | Deng9772 | 8 | 1561587 | 1680 | 244 | 0.00009 | 0.00005 | 0.00006 | -0.00006 |
| rwD83 | Deng9783 | 10 | 1873278 | 2240 | 505 | 0.00013 | 0.00008 | 0.00009 | -0.00009 |
| **Pure pink morph** | | 31 | 278341 | 173 | 89 | 0.00005 | 0.00004 | 0.00004 | -0.00002 |
| ppD10 | Deng9810 | 5 | 919460 | 1443 | 396 | 0.00025 | 0.00016 | 0.00018 | -0.00013 |
| ppD13 | Deng9813 | 8 | 651600 | 874 | 278 | 0.0002 | 0.00013 | 0.00014 | -0.00012 |
| ppD25 | Deng9525 | 7 | 631914 | 1006 | 279 | 0.00026 | 0.00015 | 0.00017 | -0.00017 |
| ppD28 | Deng9528 | 3 | 357675 | 836 | 237 | 0.00046 | 0.00028 | 0.00033 | -0.00022 |
| ppD33 | Deng9533 | 8 | 560393 | 574 | 209 | 0.00013 | 0.00009 | 0.0001 | -0.00006 |
| **Yellow-white morph** | | 46 | 278341 | 173 | 95 | 0.00004 | 0.00004 | 0.00004 | 0.00003 |
| ywC16 | Chen0016 | 8 | 1693196 | 1684 | 380 | 0.00011 | 0.00007 | 0.00008 | -0.00007 |
| ywZYZ | ZYZhou | 9 | 1124888 | 1220 | 305 | 0.00013 | 0.00008 | 0.00009 | -0.00008 |
| ywD24 | Deng9524 | 4 | 642181 | 892 | 309 | 0.00022 | 0.00014 | 0.00016 | -0.00012 |
| ywD35 | Deng9535 | 5 | 490976 | 1135 | 311 | 0.00041 | 0.00024 | 0.00026 | -0.00027 |
| ywD15 | Deng9715 | 8 | 1824804 | 1779 | 289 | 0.00008 | 0.00005 | 0.00005 | -0.00005 |

| Code | Population | n | Total Sites | Variant Sites | Polymorphic Sites | $H$o | $H$e | π | $F_{IS}$ |
|---|---|---|---|---|---|---|---|---|---|
| ywD03 | Deng9703 | 4 | 1858184 | 2093 | 473 | 0.00015 | 0.00009 | 0.0001 | -0.00008 |
| ywD47 | Deng9847 | 8 | 1816357 | 1794 | 264 | 0.00007 | 0.00004 | 0.00004 | -0.00004 |
| **Pure yellow morph** | | 71 | 278341 | 173 | 107 | 0.00003 | 0.00005 | 0.00005 | 0.00008 |
| yyN03 | Niuma003 | 3 | 497623 | 801 | 199 | 0.00028 | 0.00017 | 0.0002 | -0.00014 |
| yyC09 | Chen0009 | 9 | 1114783 | 1292 | 361 | 0.00011 | 0.00011 | 0.00011 | 0.00001 |
| yyC15 | Chen0015 | 10 | 890860 | 1261 | 447 | 0.00022 | 0.00015 | 0.00016 | -0.0001 |
| yyD39 | Deng8939 | 10 | 1175784 | 1279 | 424 | 0.00012 | 0.00011 | 0.00012 | 0 |
| yyD99 | Deng8999 | 10 | 1401234 | 1646 | 448 | 0.00012 | 0.00009 | 0.0001 | -0.00005 |
| yyD40 | Deng9040 | 10 | 1703647 | 2041 | 493 | 0.00012 | 0.00009 | 0.00009 | -0.00004 |
| yyD27 | Deng9127 | 7 | 580339 | 844 | 247 | 0.00027 | 0.00015 | 0.00017 | -0.00019 |
| yyD69 | Deng9569 | 7 | 1172977 | 1493 | 337 | 0.00015 | 0.0001 | 0.00011 | -0.00007 |
| yyD01 | Deng9601 | 5 | 993184 | 1386 | 431 | 0.00023 | 0.00015 | 0.00017 | -0.0001 |

n, number of individuals; $H$o; Mean observed heterozygosity; $H$e, Mean expected heterozygosity; π, averaged nucleotide diversity; $F_{IS}$, the inbreeding coefficient of an individual (I) relative to the subpopulation (S)

**Table S4.** The paired population-level estimation of genetic diversity indices, $\Phi_{ST}$ means (lower left diagonal) and $F_{ST}$' means (upper right diagonal) for 34 populations estimated using the *populations* pipeline in Stacks. (*Excel sheet provided for Supplementary Table S4 separately)

**Table S5.** The paired mean absolute differentiation ($D_{XY}$) among 34 populations estimated using the *populations* pipeline in Stacks. (*Excel sheet provided for Supplementary Table S5 separately)

**Table S6.** The population-level cryptic species-wise estimation of the genetic diversity indices; $\Phi_{ST}$ means, $D_{XY}$ means, and $F_{ST}$' means.

| Cryptic species | RW | PP | YW | YY |
|---|---|---|---|---|
| Red-white morph (RW) | - | 0.0541 | 0.3708 | 0.4711 |
| Pure pink morph (PP) | 0.0221/0.00077 | - | 0.3794 | 0.4967 |
| Yellow-white morph (YW) | 0.3439/0.00264 | 0.3658/0.00276 | - | 0.2091 |
| Pure yellow morph (YY) | 0.4361/0.00323 | 0.4762/0.00347 | 0.1778/0.00168 | - |

$\Phi_{ST}$ means/$D_{XY}$ means, Lower left diagonal triangle; $F_{ST}$' means, Upper right diagonal triangle
$\Phi_{ST}$ means, AMOVA-based measure of $F_{ST}$ as the total genetic variance fixation index; $D_{XY}$ means, absolute measure (nonrelative, as opposed to $F_{ST}$ like measures) of locus differentiation, measuring the number of differences between populations, excluding polymorphisms segregating within the two populations; $F_{ST}$' means, haplotype measure of $F_{ST}$ that is scaled to the theoretical maximum $F_{ST}$ value at this locus (upper right diagonal).

**Table S7.** Comparison among the nucleotides substitution model for the selection of best fitting model based on Akaike information criterion (AIC), Bayesian information criterion (BIC), and corrected AIC (AICc) measurements using Modeltest-NG.

| Model | K | *ln*L | *IC*score | delta | weight |
|---|---|---|---|---|---|
| | | | **AIC/BIC/AICc** | | |
| GTR | 8 / 8 / 8 | -124010.24 / -124010.24 / -124010.24 | 253309.79 / 249070.48 / 249093.48 | 0 / 0 / 0 | 0.53 / 0.72 / 0.72 |
| TVM | 7 / 9 / 9 | -124015.4 / -124010.26 / -124010.26 | 253310.03 / 249072.53 / 249095.53 | 0.24 / 2.04 / 2.04 | 0.47 / 0.26 / 0.26 |
| GTR+I | 9 / 7 / 7 | -124010.26 / -124015.4 / -124015.4 | 253319.91 / 249078.8 / 249101.8 | 10.12 / 8.32 / 8.32 | 0 / 0.01/ 0.01 |
| TVM+I | 8 / 8 / 8 | -124015.42 / -124015.42 / -124015.42 | 253320.14 / 249080.83 / 249103.83 | 10.35 / 10.35 / 10.35 | 0 / 0 / 0 |
| TVMef | 4 / 9 / 9 | -124077.2 / -124070.64 / -124070.64 | 253403.41 / 249193.28 / 249216.28 | 93.62 / 122.8 / 122.8 | 0 / 0 / 0 |
| SYM | 5 / 10 / 10 | -124076 / -124070.62 / -124070.62 | 253411.09 / 249195.24 / 249218.24 | 101.3 / 124.76 / 124.76 | 0 / 0 / 0 |
| TVMef+I | 5 / 5 / 5 | -124077.22 / -124076 / -124076 | 253413.53 / 249196.01 / 249219.01 | 103.74 / 125.53 / 125.53 | 0 / 0 / 0 |
| SYM+I | 6 / 4 / 4 | -124076.03 / -124077.2 / -124077.2 | 253421.22 / 249196.4 / 249219.4 | 111.42 / 125.92 / 125.92 | 0 / 0 / 0 |
| GTR+G4 | 9 / 6 / 6 | -124070.64 / -124076.03 / -124076.03 | 253440.67 / 249198.06 / 249221.06 | 130.87 / 127.57 / 127.57 | 0 / 0 / 0 |
| TVM+G4 | 8 / 5 / 5 | -124075.78 / -124077.22 / -124077.22 | 253440.88 / 249198.44 / 249221.44 | 131.08 / 127.96 / 127.96 | 0 / 0 / 0 |

K, Number of estimated parameters; *ln*L, log likelihood; *IC*score, Information criterion scores; delta, Information difference; weight, Information weight; Best fittings model is selected based on the highest likelihood, weight, lowest *IC*score, and possibly zero (0) delta score of *AIC*, *AICc*, *BIC*. For a detailed abbreviated model, refer to the software protocol of ModelTest-NG v.0.1.6 (20).

**Table S8.** Impact of the prior specification on the number of delimited cryptic species using BPP analysis (A11 = joint species delimitation and species tree estimation). Posterior probability values (*P*) are averaged over two runs.

| Models | Priors | Cryptic species (*P*) | | | | Delimited cryptic species (*P*) | | | | | |
|---|---|---|---|---|---|---|---|---|---|---|---|
| | | 2-sp | 3-sp | | 4-sp | RWPP | YWYY | RW | PP | YW | YY |
| | | RWPP-YWYY | RWPP-YW-YY | RW-PP-YWYY | RW-PP-YW-YY | | | | | | |
| model1† | $\theta$ ~ G(3, 0.002) $\tau$ ~ G(3, 0.004) | 0.2 | 0.25 | 0.25 | 0.3* | 0.45 | 0.44 | 0.55 | 0.55 | 0.56 | 0.56 |
| model2 | $\theta$ ~ G(3, 0.02) $\tau$ ~ G(3, 0.004) | 0.27* | 0.26 | 0.24 | 0.23 | 0.53 | 0.52 | 0.47 | 0.47 | 0.48 | 0.48 |
| model3 | $\theta$ ~ G(3, 0.2) $\tau$ ~ G(3, 0.004) | 0.35* | 0.24 | 0.25 | 0.17 | 0.59 | 0.6 | 0.41 | 0.41 | 0.4 | 0.4 |
| model4 | $\theta$ ~ G(3, 0.002) $\tau$ ~ G(3, 0.04) | 0.21 | 0.22 | 0.34* | 0.23 | 0.42 | 0.55 | 0.58 | 0.58 | 0.45 | 0.45 |
| model5 | $\theta$ ~ G(3, 0.002) $\tau$ ~ G(3, 0.4) | 0.41* | 0.1 | 0.27 | 0.21 | 0.52 | 0.69 | 0.48 | 0.48 | 0.31 | 0.31 |
| model6 | $\theta$ ~ G(2, 0.004) $\tau$ ~ G(2, 0.002) | 0.23 | 0.26* | 0.26 | 0.25 | 0.49 | 0.49 | 0.51 | 0.51 | 0.51 | 0.51 |
| model7 | $\theta$ ~ G(2, 0.04) $\tau$ ~ G(2, 0.002) | 0.28* | 0.25 | 0.24 | 0.23 | 0.53 | 0.52 | 0.47 | 0.47 | 0.48 | 0.48 |
| model8 | $\theta$ ~ G(2, 0.4) $\tau$ ~ G(2, 0.002) | 0.42* | 0.23 | 0.23 | 0.12 | 0.65 | 0.65 | 0.35 | 0.35 | 0.35 | 0.35 |
| model9† | $\theta$ ~ G(2, 0.004) $\tau$ ~ G(2, 0.02) | 0.18 | 0.27 | 0.27 | 0.28* | 0.45 | 0.45 | 0.55 | 0.55 | 0.55 | 0.55 |
| model10 | $\theta$ ~ G(2, 0.004) $\tau$ ~ G(2, 0.2) | 0.1 | 0.37* | 0.24 | 0.29 | 0.47 | 0.34 | 0.53 | 0.53 | 0.66 | 0.66 |
| model11† | $\theta$ ~ G(3, 0.002) $\tau$ ~ G(2, 0.004) | 0.21 | 0.25 | 0.27 | 0.27* | 0.46 | 0.48 | 0.54 | 0.54 | 0.52 | 0.52 |
| model12 | $\theta$ ~ G(3, 0.02) $\tau$ ~ G(2, 0.004) | 0.24 | 0.26* | 0.25 | 0.25 | 0.5 | 0.49 | 0.5 | 0.5 | 0.51 | 0.51 |
| model13 | $\theta$ ~ G(3, 0.2) $\tau$ ~ G(2, 0.004) | 0.35* | 0.24 | 0.24 | 0.17 | 0.59 | 0.59 | 0.41 | 0.41 | 0.41 | 0.41 |
| model14† | $\theta$ ~ G(3, 0.002) $\tau$ ~ G(2, 0.04) | 0.22 | 0.3 | 0.18 | 0.3* | 0.52 | 0.4 | 0.48 | 0.48 | 0.6 | 0.6 |
| model15 | $\theta$ ~ G(3, 0.002) $\tau$ ~ G(2, 0.4) | 0.38* | 0.19 | 0.08 | 0.35 | 0.57 | 0.47 | 0.43 | 0.43 | 0.53 | 0.53 |
| model16† | $\theta$ ~ G(2, 0.004) $\tau$ ~ G(3, 0.002) | 0.23 | 0.25 | 0.25 | 0.26* | 0.48 | 0.49 | 0.52 | 0.52 | 0.51 | 0.51 |
| model17 | $\theta$ ~ G(2, 0.04) $\tau$ ~ G(3, 0.002) | 0.31* | 0.25 | 0.24 | 0.2 | 0.55 | 0.55 | 0.45 | 0.45 | 0.45 | 0.45 |
| model18 | $\theta$ ~ G(2, 0.4) $\tau$ ~ G(3, 0.002) | 0.4* | 0.23 | 0.23 | 0.14 | 0.63 | 0.63 | 0.37 | 0.37 | 0.37 | 0.37 |
| model19 | $\theta$ ~ G(2, 0.004) $\tau$ ~ G(3, 0.02) | 0.21 | 0.24 | 0.29* | 0.25 | 0.46 | 0.51 | 0.54 | 0.54 | 0.49 | 0.49 |
| model20 | $\theta$ ~ G(2, 0.004) $\tau$ ~ G(3, 0.2) | 0.24 | 0.18 | 0.31* | 0.27 | 0.42 | 0.55 | 0.58 | 0.58 | 0.45 | 0.45 |

* indicates the selected species with the highest posterior probability (*P*) for the given model of the respective priors; †the priors model with highest posterior probability (*P*) for 4-species; RW, Red-white morph; PP, Pure pink morph; YW, Yellow-white morph; YY, Pure yellow morph.

**Table S9.** Marginal likelihood estimates under different scenarios (N, A1, A2, A3, A4) from Path Sampler analysis for BFD* (Bayes Factor Delimitation, *with genomic data) in 251 individuals and three closely related taxa (*Daphne tangutica*, *Daphne holosericea*, *Wikstroemia ligustrina*).

| **(N) Current taxonomy: RWPPYWYY** | | | | | **(A1) Splitting (Red type and Yellow type): 2-sp: RWPP-YWYY** | | | | |
|---|---|---|---|---|---|---|---|---|---|
| **Steps** | **theta** | **likelihood** | **contribution** | **ESS** | **Steps** | **theta** | **likelihood** | **contribution** | **ESS** |
| 0 | 1.00 | -105675.06 | 0.00 | 17.14 | 0 | 1.00 | -68382.18 | 0.00 | 18.80 |
| 1 | 0.73 | -112964.28 | -30737.54 | 41.53 | 1 | 0.73 | -68394.02 | -18614.13 | 41.87 |
| 2 | 0.51 | -109841.55 | -22870.41 | 2.39 | 2 | 0.51 | -68580.01 | -14748.11 | 35.30 |
| 3 | 0.35 | -116018.79 | -19138.34 | 31.89 | 3 | 0.35 | -69690.15 | -11420.17 | 8.13 |
| 4 | 0.22 | -105675.06 | -13104.79 | 17.14 | 4 | 0.22 | -68382.18 | -8498.07 | 18.80 |
| 5 | 0.13 | -112964.28 | -10059.54 | 41.53 | 5 | 0.13 | -68394.02 | -6091.21 | 41.87 |
| 6 | 0.07 | -109841.55 | -6410.53 | 2.39 | 6 | 0.07 | -68580.01 | -4134.95 | 35.30 |
| 7 | 0.03 | -116018.79 | -4362.62 | 31.89 | 7 | 0.03 | -69690.15 | -2604.94 | 8.13 |
| 8 | 0.01 | -105675.06 | -2232.03 | 17.14 | 8 | 0.01 | -68382.18 | -1447.35 | 18.80 |
| 9 | 0.00 | -112964.28 | -1101.39 | 41.53 | 9 | 0.00 | -68394.02 | -666.84 | 41.87 |
| 10 | 0.00 | -109841.55 | -327.22 | 2.39 | 10 | 0.00 | -68580.01 | -210.32 | 35.30 |
| 11 | 0.00 | -116018.79 | -39.18 | 31.89 | 11 | 0.00 | -69690.15 | -23.50 | 8.13 |
| marginal L estimate = -110383.59 | | | | Sum (ESS) = 278.85 | marginal L estimate = -68459.59 | | | | Sum (ESS) = 312.31 |

| **(A2) Splitting (Yellow type): 3-sp: RWPP-YW-YY** | | | | | **(A3) Splitting (Red type): 3-sp: RW-PP-YWYY** | | | | |
|---|---|---|---|---|---|---|---|---|---|
| **Steps** | **theta** | **likelihood** | **contribution** | **ESS** | **Steps** | **theta** | **likelihood** | **contribution** | **ESS** |
| 0 | 1.00 | -32270.93 | 0.00 | 6.91 | 0 | 1.00 | -36693.26 | 0.00 | 2.34 |
| 1 | 0.73 | -32288.31 | -8785.90 | 25.45 | 1 | 0.73 | -36843.27 | -10025.12 | 51.98 |
| 2 | 0.51 | -32436.50 | -6973.90 | 12.54 | 2 | 0.51 | -37062.22 | -7959.23 | 21.86 |
| 3 | 0.35 | -33472.74 | -5394.78 | 13.65 | 3 | 0.35 | -38442.50 | -6224.61 | 14.52 |
| 4 | 0.22 | -32270.93 | -4010.39 | 6.91 | 4 | 0.22 | -36693.26 | -4556.80 | 2.34 |
| 5 | 0.13 | -32288.31 | -2875.35 | 25.45 | 5 | 0.13 | -36843.27 | -3280.96 | 51.98 |
| 6 | 0.07 | -32436.50 | -1955.59 | 12.54 | 6 | 0.07 | -37062.22 | -2232.47 | 21.86 |

| | | | | | | | | | |
|---|---|---|---|---|---|---|---|---|---|
| 7 | 0.03 | -33472.74 | -1232.18 | 13.65 | 7 | 0.03 | -38442.50 | -1421.44 | 14.52 |
| 8 | 0.01 | -32270.93 | -683.04 | 6.91 | 8 | 0.01 | -36693.26 | -776.45 | 2.34 |
| 9 | 0.00 | -32288.31 | -314.81 | 25.45 | 9 | 0.00 | -36843.27 | -359.22 | 51.98 |
| 10 | 0.00 | -32436.50 | -99.47 | 12.54 | 10 | 0.00 | -37062.22 | -113.65 | 21.86 |
| 11 | 0.00 | -33472.74 | -11.24 | 13.65 | 11 | 0.00 | -38442.50 | -12.97 | 14.52 |
| marginal L estimate = -32336.66 | | | | Sum (ESS) = 175.65 | marginal L estimate = -36962.92 | | | | Sum (ESS) = 272.12 |

**(A4) Splitting (Red-white, Pure red, Yellow-white, and Pure yellow morph): 4-sp: RW-PP-YW-YY**

| Steps | theta | likelihood | contribution | ESS |
|---|---|---|---|---|
| 0 | 1.00 | -18882.48 | 0 | 3.65 |
| 1 | 0.73 | -19018.37 | -5172.64 | 14.67 |
| 2 | 0.51 | -19117.95 | -4107.6 | 29.07 |
| 3 | 0.35 | -19965.65 | -3172.77 | 7.88 |
| 4 | 0.22 | -18882.48 | -2345.43 | 3.65 |
| 5 | 0.13 | -19018.37 | -1693.43 | 14.67 |
| 6 | 0.07 | -19117.95 | -1152.68 | 29.07 |
| 7 | 0.03 | -19965.65 | -726.02 | 7.88 |
| 8 | 0.01 | -18882.48 | -399.59 | 3.65 |
| 9 | 0.00 | -19018.37 | -185.43 | 14.67 |
| 10 | 0.00 | -19117.95 | -58.63 | 29.07 |
| 11 | 0.00 | -19965.65 | -6.67 | 7.88 |
| marginal L estimate = -19020.89 | | | | Sum (ESS) = 165.81 |

The scenarios are N, null model; A1, alternative 1; A2, alternative 2; A3, alternative 3; A4, alternative 4
Four different cryptic species are RW, Red-white morph; PP, Pure pink morph; YW, Yellow-white morph; YY, Pure yellow morph.

**Table S10.** Descriptive statistics (mean ± SE) and their Shapiro-Wilk normality test for the normal distribution of the continuous and categorical morphological traits for four cryptic species of *S. chamaejasme* observed from the herbarium specimens.

| **Continuous morphological traits** | **Abb.** | **Trait value (mean ± SE)** | | | |
|---|---|---|---|---|---|
| | | **RW** | **PP** | **YW** | **YY** |
| Rhizome length (cm) | **rh_L** | 8.19±0.49 | 17.2±0.37* | 6.16±1.06 | 7.9±0.48 |
| Stem number | **stm_no** | 7.91±0.24 | 7.4±1.25* | 8.81±0.55 | 6.81±0.25 |
| Stem avg length (cm) | **stm_L** | 24.53±0.32 | 28.2±1.39* | 27.75±0.67 | 26.44±0.39 |
| Leaf petiole length (mm) | **pet_L** | 0.99±0 | 1±0* | 1±0.02 | 1.01±0.01 |
| Leaf count total (average leaf count of basal, medium and tip part of each stem of plant) | **count** | 52.32±0.57 | 78.4±2.79* | 53.31±1.34 | 52.75±0.61* |
| Leaf density per 2 cm distance of stem length (avg count/cm) | **density** | 4.51±0.05 | 5.61±0.32* | 3.95±0.1 | 4.21±0.07 |
| Average of leaf length measured from the basal, mid and tip part of each stem of plants (mm) | **lf_L** | 15.83±0.12 | 20.06±1.23* | 16.91±0.35* | 17.99±0.19 |
| Average leaf width measured from the basal, mid and tip part of each stem of plants (mm) | **lf_W** | 4.22±0.05 | 3.19±0.23* | 4.16±0.12 | 4.63±0.09 |
| Leaf length/width ratio (lf_L/lf_W) | **lf_LWr** | 3.93±0.03 | 6.31±0.14* | 4.23±0.09 | 4.32±0.1 |
| Leaf area ($mm^2$) = lfL_avg*lfW_avg*0.75 (0.75 is the standard factor) | **LA** | 51.71±0.99 | 48.83±6.6* | 54.45±2.51 | 64.78±1.78 |
| Leaf width (mm) measured at basal region (B) | **lfW_base** | 4.27±0.06 | 3.12±0.26* | 4.22±0.16 | 4.12±0.09 |
| Leaf width (mm) measured at apex region (A) | **lfW_apex** | 3.34±0.06 | 3±0.25* | 4.16±0.16 | 4.8±0.09 |
| Ratio of width at basal to apex region [lfW_base (B)/lfW_apex (A)] | **lfW_BAr** | 1.34±0.01 | 1.04±0.02* | 1.02±0.01 | 0.85±0.01 |
| Floral bract length in mm (min - max) | **bract_L** | 11.62±0.11 | 13±1.55* | 11.63±0.39 | 12.68±0.23 |
| Floral bract width in mm (min - max) | **bract_W** | 3.25±0.05 | 3.6±0.51* | 3.01±0.11 | 3.51±0.1 |
| Floral bract ratio (length/width average) | **bract_LWr** | 3.86±0.05 | 3.67±0.15* | 4.11±0.16 | 4.01±0.1 |
| Calyx tube cylindric diameter (mm) | **tube_diam** | 0.98±0.02 | 1.36±0.22 | 1.02±0.02 | 1.06±0.03 |
| Calyx tube length (mm) | **tube_L** | 8.9±0.1 | 10.2±0.8* | 9.8±0.1 | 9.9±0.1 |
| Calyx lobe length (mm) | **lobe_L** | 2.18±0.02 | 2.6±0.24 | 2.16±0.04 | 2.11±0.03 |
| Calyx lobe width (mm) | **lobe_W** | 1.12±0.01 | 1.46±0.2* | 1.07±0.02 | 1.02±0.02 |
| Calyx lobe Length / Width ratio | **lobe_LWr** | 1.97±0.01 | 1.83±0.11 | 2.03±0.02 | 2.01±0.03 |
| Flowering month | **Flowering** | 6.46±0.05 | 6.4±0.24 | 6.25±0.12 | 5.67±0.09 |
| **Categorical morphological traits** | | | | | |
| Internode distance [1 is the minimum distance, keeping 1 mm constant for all; (1) --10 mm = 1; > 10 mm = 2] | **id_leaf** | 1.22±0.02 | 1±0*nc* | 1.56±0.06*nc* | 1.37±0.03 |
| Leaf angle (degree) (< 30 degree = 1; 30 - 35 degree = 2; > 35 degrees = 3) | **Angle** | 1.92±0.02 | 2±0*nc* | 1.68±0.06*nc* | 1.83±0.03 |
| Leaf shape (lanceolate = 1; linear = 2; oblanceolate =3, oblong-lanceolate= 4) | **lf_shape** | 1±0*nc* | 2±0*nc* | 4±0*nc* | 2.95±0.01 |
| Leaf mid part observation from base (narrower = 1; equal = 2; broader = 3; broader to narrower = 4) | **marg_obs** | 1±0*nc* | 2±0*nc* | 4±0*nc* | 2.95±0.01 |

| | | | | | |
|---|---|---|---|---|---|
| Leaf base shape (cuneate=1, obtuse =2, acute =3) | **lf_base** | 2±0*nc* | 1±0*nc* | 2±0*nc* | 2.9±0.02 |
| Leaf apex shape (acute =1, obtuse =2, acute-obtuse =3) | **lf_apex** | 1.01±0.01 | 1±0*nc* | 1±0*nc* | 1.07±0.02 |
| Mid rib colour with leaf (same = 1; different = 2) | **mid_col** | 1±0*nc* | 2±0*nc* | 1±0*nc* | 1.45±0.03 |
| Leaf mid rib appearance (abaxial) (level = 1; elevated = 2) | **mid_ab** | 1.01±0 | 1±0*nc* | 1±0*nc* | 1.14±0.02 |
| Leaf mid rib appearance (adaxial) (level = 1; sunken = 2) | **mid_ad** | 1±0 | 1±0*nc* | 1±0*nc* | 1.15±0.02 |
| Flower number (< 30 = 1, > 30 = 2) | **fl_no** | 1.01±0.01 | 1±0*nc* | 1±0*nc* | 1.02±0.02 |
| Bract number (< = 6 no. = 1; > 6 no. = 2) | **bract_no** | 1.13±0.01 | 1±0*nc* | 1.11±0.04 | 1.05±0.02 |
| Calyx tube outer colour (red = 1; yellow = 2) | **tube_col** | 0.99±0 | 1±0*nc* | 2±0*nc* | 2±0*nc* |
| Calyx lobe colour (especially inner) (white = 1; pink = 2; yellow = 3) | **lobe_col** | 1±0*nc* | 2±0*nc* | 1±0*nc* | 3±0*nc* |

*normal distribution at significance level 0.05; *nc*, not calculated; four cryptic species are RW, Red-white morph; PP, Pure pink morph; YW, Yellow-white morph; YY, Pure yellow morph.

**Table S11.** Levene's test of homogeneity of variance assumptions for the morphological traits of *S. chamaejasme*.

| **Continuous morphological traits** | | | **Categorical morphological traits** | | |
|---|---|---|---|---|---|
| **Variables** | **Statistics** | ***p-value*** | **Variables** | **Statistics** | ***p-value*** |
| bract_L | 11.53 | $2.00e^{-07}$ | angle | 35.86 | $5.90e^{-22}$ |
| bract_W | 16.03 | $3.60e^{-10}$ | bract_no | 0.64 | $5.87e^{-01}$ |
| bract_LWr | 12.55 | $4.74e^{-08}$ | fl_no | 5.67 | $7.51e^{-04}$ |
| count | 1.56 | $1.97e^{-01}$ | id_leaf | 12.23 | $7.45e^{-08}$ |
| density | 1.56 | $1.97e^{-01}$ | lf_apex | 5.93 | $5.25e^{-04}$ |
| flowering | 5.72 | $6.98e^{-04}$ | lf_base | 10.74 | $6.04e^{-07}$ |
| LA | 9.73 | $2.51e^{-06}$ | lf_shape | 10.74 | $6.04e^{-07}$ |
| lf_LWr | 15.77 | $5.14e^{-10}$ | lobe_col | NA | NA |
| lfW_BAr | 38.91 | $1.03e^{-23}$ | marg_obs | 10.74 | $6.04e^{-07}$ |
| lobe_LWr | 6.48 | $2.41e^{-04}$ | mid_ab | 26.29 | $2.44e^{-16}$ |
| pet_L | 14.92 | $1.71e^{-09}$ | mid_ad | 33.68 | $1.10e^{-20}$ |
| rh_L | 2.24 | $8.24e^{-02}$ | mid_col | 171.33 | $1.06e^{-88}$ |
| stm_L | 3.49 | $1.53e^{-02}$ | tube_col | 1.54 | $2.04e^{-01}$ |
| stm_no | 2.19 | $8.75e^{-02}$ | | | |
| tube_diam | 1.24 | $2.94e^{-01}$ | | | |
| tube_L | 4.96 | $2.03e^{-03}$ | | | |

The significant Levene's test ($p< 0.05$) indicates no homogeneity of variances, whereas not significant ($p> 0.05$) indicates homogeneity of variances.
# df1, 3; df2, 946

**Table S12.** MANOVA test for the significant difference between four cryptic species of *S. chamaejasme* for continuous morphological traits and the trait-wise test of significant difference between the cryptic species.

| **Continuous traits** | | **den Df** | **Pillai** | **approx F** | **Pr(>F)** | **Categorical traits** | | | | | |
|---|---|---|---|---|---|---|---|---|---|---|---|
| **All cryptic species** | | 2799 | 0.69555 | 17.6 | $< 2.2e^{-16}$ *** | All cryptic species | **The ranked categorical morphological traits were not performed.** | | | | |
| **Trait-wise** | | **Sum Sq** | **Mean Sq** | **F value** | **Pr(>F)** | **Trait-wise** | | **Sum Sq** | **Mean Sq** | **F value** | **Pr(>F)** |
| rh_L | species | 718 | 239.23 | 2.24 | 0.08 | id_leaf | species | 11.01 | 3.67 | 18.62 | $9.67e^{-12}$ *** |
| | Residuals | 100942 | 106.70 | | | | Residuals | 186.46 | 0.20 | | |
| stm_no | species | 372.20 | 124.07 | 4.51 | 0.003777 ** | angle | species | 4.978 | 1.66 | 7.42 | $6.49e^{-05}$ *** |
| | Residuals | 26028.3 | 27.51 | | | | Residuals | 211.58 | 0.22 | | |
| stm_L | species | 1245 | 415.03 | 7.96 | 0.00003025*** | lf_shape | species | 1151.92 | 383.97 | 23878 | $< 2.2e^{-16}$ *** |
| | Residuals | 49312 | 52.13 | | | | Residuals | 15.21 | 0.02 | | |
| pet_L | species | 0.13 | 0.04 | 2.31 | 0.08 | marg_obs | species | 1151.92 | 383.97 | 23878 | $< 2.2e^{-16}$ *** |
| | Residuals | 17.57 | 0.02 | | | | Residuals | 15.21 | 0.02 | | |
| count | species | 3410 | 1136.59 | 7.45 | $6.22e^{-05}$ *** | lf_base | species | 181.84 | 60.61 | 942.34 | $< 2.2e^{-16}$ *** |
| | Residuals | 144348 | 152.59 | | | | Residuals | 60.85 | 0.06 | | |
| density | species | 41.46 | 13.82 | 9.03 | $6.71e^{-06}$ *** | lf_apex | species | 0.89 | 0.30 | 5.93 | 0.0005247 *** |
| | Residuals | 1447.47 | 1.53 | | | | Residuals | 47.34 | 0.05 | | |
| lf_LWr | species | 57.01 | 19 | 12.65 | $4.13e^{-08}$ *** | mid_col | species | 47.17 | 15.72 | 184.75 | $< 2.2e^{-16}$ *** |
| | Residuals | 1421.19 | 1.50 | | | | Residuals | 80.51 | 0.09 | | |
| LA | species | 35257 | 11752.30 | 16.84 | $1.15e^{-10}$ *** | mid_ab | species | 3.65 | 1.22 | 26.29 | $2.44e^{-16}$ *** |
| | Residuals | 660027 | 697.70 | | | | Residuals | 43.72 | 0.05 | | |
| lfW_BAr | species | 48.663 | 16.22 | 407.37 | $< 2.2e^{-16}$ *** | mid_ad | species | 4.657 | 1.55 | 33.68 | $< 2.2e^{-16}$ *** |
| | Residuals | 37.67 | 0.04 | | | | Residuals | 43.61 | 0.05 | | |
| bract_L | species | 245 | 81.67 | 7.32 | $7.43e^{-05}$ *** | fl_no | species | 0.02 | 0.01 | 0.12 | 0.95 |
| | Residuals | 10552 | 11.15 | | | | Residuals | 43.83 | 0.05 | | |
| bract_W | species | 21.93 | 7.31 | 3.69 | 0.01167 | bract_no | species | 1.4 | 0.47 | 3.23 | 0.02187 |
| | Residuals | 1874.39 | 1.98 | | | | Residuals | 136.49 | 0.14 | | |

| | | | | | | | | | | | |
|---|---|---|---|---|---|---|---|---|---|---|---|
| bract_LWr | species | 8.11 | 2.70 | 1.35 | 0.26 | tube_col | species | 237.65 | 79.22 | 12630 | < 2.2e$^{-16}$ *** |
| | Residuals | 1888.29 | 2.00 | | | | Residuals | 5.93 | 0.01 | | |
| tube_diam | species | 1.91 | 0.64 | 2.40 | 0.07 | lobe_col | species | 853.39 | 284.46 | 5.07e$^{+29}$ | < 2.2e$^{-16}$ *** |
| | Residuals | 251.23 | 0.27 | | | | Residuals | 0 | 0.00 | | |
| tube_L | species | 249.3 | 83.12 | 20.50 | 7.13e$^{-13}$ *** | | | | | | |
| | Residuals | 3836 | 4.06 | | | | | | | | |
| lobe_LWr | species | 0.64 | 0.21 | 1.61 | 0.19 | | | | | | |
| | Residuals | 126.18 | 0.13 | | | | | | | | |
| flowering | species | 127.26 | 42.42 | 21.30 | 2.34e$^{-13}$ *** | | | | | | |
| | Residuals | 1883.73 | 1.99 | | | | | | | | |

Df (Species: 3, Residuals: 946)
Significance codes: '****' 0 '***' 0.001 '**' 0.01 '*' 0.05

**Table S13.** Post-hoc TukeyHSD MANOVA test for the trait-species wise significance difference between the four cryptic species of *S. chamaejasme*.

| Traits | Term | Group1 | Group2 | null value | p.adj | p.adj. sig. | Traits | Term | Group1 | Group2 | null. value | p.adj | p.adj. sig. |
|---|---|---|---|---|---|---|---|---|---|---|---|---|---|
| **Continuous morphological traits** | | | | | | | **Categorical morphological traits** | | | | | | |
| bract_L | species | PP | RW | 0 | 0.79 | *ns* | angle | species | PP | RW | 0 | 0.98 | *ns* |
| bract_L | species | PP | YW | 0 | 0.81 | *ns* | angle | species | PP | YW | 0 | 0.45 | *ns* |
| bract_L | species | PP | YY | 0 | 1 | *ns* | angle | species | PP | YY | 0 | 0.85 | *ns* |
| bract_L | species | RW | YW | 0 | 1 | *ns* | angle | species | RW | YW | 0 | 0.000121 | *** |
| bract_L | species | RW | YY | 0 | 3.98e$^{-05}$ | **** | angle | species | RW | YY | 0 | 0.03 | * |
| bract_L | species | YW | YY | 0 | 0.05 | *ns* | angle | species | YW | YY | 0 | 0.06 | *ns* |
| bract_LWr | species | PP | RW | 0 | 0.99 | *ns* | bract_no | species | PP | RW | 0 | 0.86 | *ns* |
| bract_LWr | species | PP | YW | 0 | 0.91 | *ns* | bract_no | species | PP | YW | 0 | 0.92 | *ns* |
| bract_LWr | species | PP | YY | 0 | 0.95 | *ns* | bract_no | species | PP | YY | 0 | 0.99 | *ns* |
| bract_LWr | species | RW | YW | 0 | 0.43 | *ns* | bract_no | species | RW | YW | 0 | 0.96 | *ns* |
| bract_LWr | species | RW | YY | 0 | 0.4 | *ns* | bract_no | species | RW | YY | 0 | 0.01 | * |
| bract_LWr | species | YW | YY | 0 | 0.94 | *ns* | bract_no | species | YW | YY | 0 | 0.6 | *ns* |
| bract_W | species | PP | RW | 0 | 0.95 | *ns* | fl_no | species | PP | RW | 0 | 1 | *ns* |
| bract_W | species | PP | YW | 0 | 0.8 | *ns* | fl_no | species | PP | YW | 0 | 1 | *ns* |
| bract_W | species | PP | YY | 0 | 1 | *ns* | fl_no | species | PP | YY | 0 | 1 | *ns* |
| bract_W | species | RW | YW | 0 | 0.47 | *ns* | fl_no | species | RW | YW | 0 | 0.96 | *ns* |
| bract_W | species | RW | YY | 0 | 0.05 | *ns* | fl_no | species | RW | YY | 0 | 1 | *ns* |
| bract_W | species | YW | YY | 0 | 0.02 | * | fl_no | species | YW | YY | 0 | 0.94 | *ns* |
| count | species | PP | RW | 0 | 1.78e$^{-05}$ | **** | id_leaf | species | PP | RW | 0 | 0.70 | *ns* |
| count | species | PP | YW | 0 | 6.86e$^{-05}$ | **** | id_leaf | species | PP | YW | 0 | 0.03 | * |
| count | species | PP | YY | 0 | 2.74e$^{-05}$ | **** | id_leaf | species | PP | YY | 0 | 0.26 | *ns* |
| count | species | RW | YW | 0 | 0.91 | *ns* | id_leaf | species | RW | YW | 0 | 1.19e$^{-09}$ | **** |
| count | species | RW | YY | 0 | 0.96 | *ns* | id_leaf | species | RW | YY | 0 | 8.97e$^{-06}$ | **** |
| count | species | YW | YY | 0 | 0.98 | *ns* | id_leaf | species | YW | YY | 0 | 0.00345 | ** |
| density | species | PP | RW | 0 | 0.2 | *ns* | lf_apex | species | PP | RW | 0 | 1 | *ns* |

| Traits | Term | Group1 | Group2 | null value | p.adj | p.adj. sig. | Traits | Term | Group1 | Group2 | null. value | p.adj | p.adj. sig. |
|---|---|---|---|---|---|---|---|---|---|---|---|---|---|
| **Continuous morphological traits** | | | | | | | **Categorical morphological traits** | | | | | | |
| density | species | PP | YW | 0 | 0.02 | * | lf_apex | species | PP | YW | 0 | 1 | *ns* |
| density | species | PP | YY | 0 | 0.06 | *ns* | lf_apex | species | PP | YY | 0 | 0.9 | *ns* |
| density | species | RW | YW | 0 | 0.000884 | *** | lf_apex | species | RW | YW | 0 | 0.99 | *ns* |
| density | species | RW | YY | 0 | 0.00254 | ** | lf_apex | species | RW | YY | 0 | 0.000347 | *** |
| density | species | YW | YY | 0 | 0.35 | *ns* | lf_apex | species | YW | YY | 0 | 0.0536 | *ns* |
| flowering | species | PP | RW | 0 | 1 | *ns* | lf_base | species | PP | RW | 0 | 0 | **** |
| flowering | species | PP | YW | 0 | 1 | *ns* | lf_base | species | PP | YW | 0 | 0 | **** |
| flowering | species | PP | YY | 0 | 0.66 | *ns* | lf_base | species | PP | YY | 0 | 0 | **** |
| flowering | species | RW | YW | 0 | 0.57 | *ns* | lf_base | species | RW | YW | 0 | 1 | *ns* |
| flowering | species | RW | YY | 0 | 0 | **** | lf_base | species | RW | YY | 0 | 0 | **** |
| flowering | species | YW | YY | 0 | 0.00612 | ** | lf_base | species | YW | YY | 0 | 0 | **** |
| LA | species | PP | RW | 0 | 1 | *ns* | lf_shape | species | PP | RW | 0 | 0 | **** |
| LA | species | PP | YW | 0 | 0.97 | *ns* | lf_shape | species | PP | YW | 0 | 0 | **** |
| LA | species | PP | YY | 0 | 0.54 | *ns* | lf_shape | species | PP | YY | 0 | 0 | **** |
| LA | species | RW | YW | 0 | 0.82 | *ns* | lf_shape | species | RW | YW | 0 | 0 | **** |
| LA | species | RW | YY | 0 | $2.14e^{-11}$ | **** | lf_shape | species | RW | YY | 0 | 0 | **** |
| LA | species | YW | YY | 0 | 0.0092 | ** | lf_shape | species | YW | YY | 0 | 0 | **** |
| lf_LWr | species | PP | RW | 0 | 0.000106 | *** | lobe_col | species | PP | RW | 0 | 0 | **** |
| lf_LWr | species | PP | YW | 0 | 0.00143 | ** | lobe_col | species | PP | YW | 0 | 0 | **** |
| lf_LWr | species | PP | YY | 0 | 0.00198 | ** | lobe_col | species | PP | YY | 0 | 0 | **** |
| lf_LWr | species | RW | YW | 0 | 0.18 | *ns* | lobe_col | species | RW | YW | 0 | 0.80 | *ns* |
| lf_LWr | species | RW | YY | 0 | $3.75e^{-05}$ | **** | lobe_col | species | RW | YY | 0 | 0 | **** |
| lf_LWr | species | YW | YY | 0 | 0.93 | *ns* | lobe_col | species | YW | YY | 0 | 0 | **** |
| lfW_BAr | species | PP | RW | 0 | 0.00616 | ** | marg_obs | species | PP | RW | 0 | 0 | **** |
| lfW_BAr | species | PP | YW | 0 | 0.99 | *ns* | marg_obs | species | PP | YW | 0 | 0 | **** |
| lfW_BAr | species | PP | YY | 0 | 0.15 | *ns* | marg_obs | species | PP | YY | 0 | 0 | **** |
| lfW_BAr | species | RW | YW | 0 | 0 | **** | marg_obs | species | RW | YW | 0 | 0 | **** |

| Traits | Term | Group1 | Group2 | null value | p.adj | p.adj. sig. | Traits | Term | Group1 | Group2 | null. value | p.adj | p.adj. sig. |
|---|---|---|---|---|---|---|---|---|---|---|---|---|---|
| **Continuous morphological traits** | | | | | | | **Categorical morphological traits** | | | | | | |
| lfW_BAr | species | RW | YY | 0 | 0 | **** | marg_obs | species | RW | YY | 0 | 0 | **** |
| lfW_BAr | species | YW | YY | 0 | $3.25e^{-10}$ | **** | marg_obs | species | YW | YY | 0 | 0 | **** |
| lobe_LWr | species | PP | RW | 0 | 0.85 | *ns* | mid_ab | species | PP | RW | 0 | 1 | *ns* |
| lobe_LWr | species | PP | YW | 0 | 0.64 | *ns* | mid_ab | species | PP | YW | 0 | 1 | *ns* |
| lobe_LWr | species | PP | YY | 0 | 0.72 | *ns* | mid_ab | species | PP | YY | 0 | 0.48 | *ns* |
| lobe_LWr | species | RW | YW | 0 | 0.41 | *ns* | mid_ab | species | RW | YW | 0 | 0.98 | *ns* |
| lobe_LWr | species | RW | YY | 0 | 0.40 | *ns* | mid_ab | species | RW | YY | 0 | 0 | **** |
| lobe_LWr | species | YW | YY | 0 | 0.94 | *ns* | mid_ab | species | YW | YY | 0 | $1.57e^{-06}$ | **** |
| pet_L | species | PP | RW | 0 | 1 | *ns* | mid_ad | species | PP | RW | 0 | 1 | *ns* |
| pet_L | species | PP | YW | 0 | 1 | *ns* | mid_ad | species | PP | YW | 0 | 1 | *ns* |
| pet_L | species | PP | YY | 0 | 1 | *ns* | mid_ad | species | PP | YY | 0 | 0.40 | *ns* |
| pet_L | species | RW | YW | 0 | 0.69 | *ns* | mid_ad | species | RW | YW | 0 | 1 | *ns* |
| pet_L | species | RW | YY | 0 | 0.05 | * | mid_ad | species | RW | YY | 0 | 0 | **** |
| pet_L | species | YW | YY | 0 | 0.98 | *ns* | mid_ad | species | YW | YY | 0 | $1.24e^{-07}$ | **** |
| rh_L | species | PP | RW | 0 | 0.21 | *ns* | mid_col | species | PP | RW | 0 | $1.86e^{-13}$ | **** |
| rh_L | species | PP | YW | 0 | 0.09 | *ns* | mid_col | species | PP | YW | 0 | $1.21e^{-12}$ | **** |
| rh_L | species | PP | YY | 0 | 0.19 | *ns* | mid_col | species | PP | YY | 0 | 0.000198 | *** |
| rh_L | species | RW | YW | 0 | 0.35 | *ns* | mid_col | species | RW | YW | 0 | 1 | *ns* |
| rh_L | species | RW | YY | 0 | 0.98 | *ns* | mid_col | species | RW | YY | 0 | 0 | **** |
| rh_L | species | YW | YY | 0 | 0.53 | *ns* | mid_col | species | YW | YY | 0 | 0 | **** |
| stm_L | species | PP | RW | 0 | 0.67 | *ns* | tube_col | species | PP | RW | 0 | 0.99 | *ns* |
| stm_L | species | PP | YW | 0 | 1 | *ns* | tube_col | species | PP | YW | 0 | 0 | **** |
| stm_L | species | PP | YY | 0 | 0.95 | *ns* | tube_col | species | PP | YY | 0 | 0 | **** |
| stm_L | species | RW | YW | 0 | 0.00111 | ** | tube_col | species | RW | YW | 0 | 0 | **** |
| stm_L | species | RW | YY | 0 | 0.00095 | *** | tube_col | species | RW | YY | 0 | 0 | **** |
| stm_L | species | YW | YY | 0 | 0.47 | *ns* | tube_col | species | YW | YY | 0 | 1 | *ns* |
| stm_no | species | PP | RW | 0 | 1 | *ns* | | | | | | | |

| Traits | Term | Group1 | Group2 | null value | p.adj | p.adj. sig. | Traits | Term | Group1 | Group2 | null. value | p.adj | p.adj. sig. |
|---|---|---|---|---|---|---|---|---|---|---|---|---|---|
| **Continuous morphological traits** | | | | | | | **Categorical morphological traits** | | | | | | |
| stm_no | species | PP | YW | 0 | 0.94 | *ns* | | | | | | | |
| stm_no | species | PP | YY | 0 | 1 | *ns* | | | | | | | |
| stm_no | species | RW | YW | 0 | 0.47 | *ns* | | | | | | | |
| stm_no | species | RW | YY | 0 | 0.02 | * | | | | | | | |
| stm_no | species | YW | YY | 0 | 0.01 | * | | | | | | | |
| tube_diam | species | PP | RW | 0 | 0.35 | *ns* | | | | | | | |
| tube_diam | species | PP | YW | 0 | 0.48 | *ns* | | | | | | | |
| tube_diam | species | PP | YY | 0 | 0.56 | *ns* | | | | | | | |
| tube_diam | species | RW | YW | 0 | 0.88 | *ns* | | | | | | | |
| tube_diam | species | RW | YY | 0 | 0.13 | *ns* | | | | | | | |
| tube_diam | species | YW | YY | 0 | 0.95 | *ns* | | | | | | | |
| tube_L | species | PP | RW | 0 | 0.47 | *ns* | | | | | | | |
| tube_L | species | PP | YW | 0 | 0.98 | *ns* | | | | | | | |
| tube_L | species | PP | YY | 0 | 0.99 | *ns* | | | | | | | |
| tube_L | species | RW | YW | 0 | 0.000519 | *** | | | | | | | |
| tube_L | species | RW | YY | 0 | $1.42e^{-12}$ | **** | | | | | | | |
| tube_L | species | YW | YY | 0 | 0.98 | *ns* | | | | | | | |

RW, Red-white morph; PP, Pure pink morph; YW, Yellow-white morph; YY, Pure yellow morph; *ns*, not significant
Significance codes: ‘****’ 0 ‘***’ 0.001 ‘**’ 0.01 ‘*’ 0.05

**Table S14.** Predictive variable selection for the ensemble species distribution modeling (eSDM) of each cryptic species of *S. chamaejasme*.

| Categories | Descriptions | Abb. (Range) | Presence (+) or absence (-) of variables (vif / Pearson) | | | |
|---|---|---|---|---|---|---|
| | | | PP | RW | YW | YY |
| **Bioclimatic variables (*V1*)** *(http://www.worldclim.org/version2)* | Mean diurnal range (°C) | bio2 (7.8–16.6) | - | + / + | - | + / + |
| | Isothermality [(bio2/bio7) × 100] | bio3 (23–49) | + / + | - / + | + / + | + / + |
| | Temperature seasonality (standard deviation) | bio4 (47.49–76.95) | - / + | - | + / + | - |
| | Max temperature of warmest month (°C) | bio5 (12.4–28.4) | + / + | - | + / + | - |
| | Min temperature of coldest month (°C) | bio6 (-33.3–3.0) | - | + / + | - | - |
| | Temperature annual range (bio5 - bio6) (°C) | bio7 (19.9–58.4) | + / + | - / + | - | - / + |
| | Mean temperature of wettest quarter (°C) | bio8 (5.8–24.2) | - | + / + | - | + / + |
| | Annual precipitation (mm) | bio12 (460–994) | - | - | + / + | - |
| | Precipitation of wettest month (mm) | bio13 (57–534) | - | + / + | - | + / + |
| | Precipitation of driest month (mm) | bio14 (1–14) | + / + | + / - | - | + / + |
| | Precipitation seasonality (coefficient of variation) | bio15 (59–150) | - / + | + / + | + / + | + / + |
| | Precipitation of driest quarter (mm) | bio17 (7–30) | - | - | + / + | - |
| | Precipitation of warmest quarter (mm) | bio18 (363–435) | - / + | - | - | - |
| | Precipitation of coldest quarter (mm) | bio19 (1–134) | - | + / + | - | - |
| | Annual mean temperature (°C) | bio1 | - | - | - | - |
| | Mean temperature of driest quarter (°C) | bio9 | - | - | - | - |
| | Mean temperature of warmest quarter (°C) | bio10 | - | - | - | - |
| | Mean temperature of coldest quarter (°C) | bio11 | - | - | - | - |
| | Precipitation of wettest quarter (mm) | bio16 | - | - | - | - |
| **Geo-climatic (*V2*)** *(http://www.worldclim.org/; version2http://nelson.wisc.edu/sage; http://www.cgiar-csi.org)* | Annual aridity (ratio) | AI (2115.03–14133.70) | - / + | + / + | + / + | + / + |
| | Aspect (degree) | Asp (10.40–346.41) | + / - | + / + | + / + | + / + |
| | Elevation (m) | Ele (282.90–5043.75) | + / + | + / + | + / + | + / + |
| | Net primary productivity (kg-carbon/m²/year) | NPP (0.01–0.95) | - | + / + | + / + | + / + |
| | Potential evapotranspiration (mm) | PET (6.23–100.89) | + / + | + / + | + / + | + / + |
| | Potential vegetation (15 biomes categories) | PV (1–13) | - | + / + | + / + | + / + |
| | Annual Relative Humidity (%) | RH (-8–76) | + / - | + / + | - | + / + |
| | Soil carbon (Kg-Carbon/m² to 1 m depth) | SC (3.49–18.81) | - | + / + | - / + | + / + |
| | Soil moisture (mm based on 150 mm water holding capacity) | SM (1.06–150) | - | + / + | + / + | + / + |
| | Soil pH (acidity-alkaline | SpH (4.69–7.89) | *bs** | + / + | + / + | + / + |

| | | | | | | |
|---|---|---|---|---|---|---|
| | level) | | | | | |
| | Annual solar radiation (kJ/m²/day) | ann_SR (11501.40–17629.30) | - | + / + | + / + | + / + |
| | Annual wind speed (m/s) | ann_WS (1.47–4.31) | - | + / + | + / + | + / - |
| | Annual water vapour (kPa) | ann_WV (0.27–1.68) | + / + | + / + | + / + | - |
| **Habitat heterogeneity (*V3*)** *(http://www.earthenv.org/texture)* | Coefficient of variation (normalized dispersion of EVI, x ≥0) | CV (0.06–0.8916) | + / - | + / + | + / + | + / + |
| | Contrast (exponentially weighted difference in EVI between adjacent pixels, x ≥0) | Cont (1.79–63.48) | - | + / + | + / + | + / + |
| | Correlation (linear dependency of EVI on adjacent pixels, 1≥ x ≥-1) | Corr (0.51–0.92) | + / - | + / + | + / + | + / + |
| | Evenness (evenness of EVI, 1≥ x ≥0) | Even (0.63–0.97) | + / + | + / + | + / + | + / + |
| | Homogeneity (similarity of EVI between adjacent pixels, 1≥ x ≥0) | Homo (0.3–0.34) | + / + | - | - | - |
| | Maximum (dominance of EVI combinations between adjacent pixels, 1≥ x ≥0) | Max (0.01–0.05) | - | - | + / + | + / + |
| | Range (Range of EVI, x ≥0) | Range (0.08–0.56) | - | + / + | - | - |
| | Uniformity (orderliness of EVI, 1≥ x ≥0) | Uni (0.003–0.12) | - | + / + | - | - |
| | Variance (Dispersion of EVI combinations between adjacent pixels, x ≥0) | Var (2.38–176.005) | - | + / + | - | - |
| | Simpson index | Simpson | - | - | - | - |
| | Standard deviation | Std | - | - | - | - |
| | Shannon index | Shannon | - | - | - | - |
| | Dissimilarity | Dissimilarity | - | - | - | - |
| | Entrophy | Entrophy | - | - | - | - |
| **Growing days (*V4*)** *(https://chelsa-climate.org/)* | First day of growing season (Julian day) | FGD (38.71–152.33) | - | - | - | + / + |
| | Growing degree days (°C based on 5-degree base temperature) | GDD (3–5325) | + / + | + / + | + / + | + / + |
| | Growing season length (number of days) | GSL (45.82–351.55) | + / + | + / + | + / + | + / + |
| | Growing season temperature (°C/10) | GST (5–19.34) | - | + / + | + / + | + / + |
| | Last day of growing season (Julian day) | LGD (15.66–322.80) | + / + | + / + | + / + | + / + |
| **Ultra-violet radiations (*V5*)** *(https://www.ufz.de/gluv/)* | Annual mean UV-B (J/m²/day) | UVB1 (5674.93–5921.47) | + / + | - | - | - |
| | UV-B seasonality (J/m²/day) | UVB2 (118219–280208) | + / + | + / + | + / + | - |
| | Mean UV-B of highest month (J/m²/day) | UVB3 (4240.67–9333.17) | - | - | - | + / + |
| | Mean UV-B of lowest month (J/m²/day) | UVB4 (160.02–2952.21) | - | + / - | + / + | + / + |
| | Sum of monthly mean UV-B during highest quarter | UVB5 | - | - | - | - |
| | Sum of monthly mean UV-B during lowest quarter | UVB6 | - | - | - | - |
| **Consensus landcover (*V6*)** *(http://www.ea* | Evergreen/deciduous Needleleaf trees (%) | LULC1 (0.02–60.46) | + | + | + | + |
| | Evergreen broadleaf trees (%) | LULC2 (0.13–48.90) | + | - | - | - |

| | | | | | | |
|---|---|---|---|---|---|---|
| *rthenv.org/landcover)* | Mixed/other trees (%) | LULC4 (0.36–60.17) | + | - | + | + |
| | Shrubs (%) | LULC5 (0.03–45.43) | - | - | - | + |
| | Herbaceous vegetation (%) | LULC6 (0.01–92.89) | + | + | + | + |
| | Cultivated and managed vegetation (%) | LULC7 (0.01–93.50) | - | + | + | + |

°C, degree Celsius; mm, millimeter; kg, kilogram; kJ, kilo Joules; s, second; kPa, kilo Pascal; %, percentage; EVI, Enhanced Vegetation Index; *bs**, significant biological variables; four cryptic species are: RW, Red-white morph; PP, Pure pink morph; YW, Yellow-white morph; YY, Pure yellow morph

**Table S15.** Variance Inflation Factor (VIF) in different test runs to select explanatory variables for four cryptic species of *S. chamaejasme*. VIF<10 (bold text) are the selected predictive variables.

a) Red-white morph (RW)

| Variables | Run1 | Run2 | Run3 | Run4 | Run5 | Run6 | Run7 | Run8 | Run9 | Run10 | Run11 | Run12 |
|---|---|---|---|---|---|---|---|---|---|---|---|---|
| **Bioclimatic variables** | | | | | | | | | | | | |
| **bio8** | 207.24 | 201.8 | 178.5 | 150.61 | 91.63 | 57.99 | 56.9 | 56.36 | 55.63 | 7.83 | 6.98 | 1.4 |
| **bio2** | 103.52 | 80.69 | 52.56 | 49.21 | 25.49 | 19.14 | 18.68 | 18.54 | 9.88 | 4.87 | 4.42 | 1.94 |
| **bio6** | 107.21 | 106.83 | 99.87 | 99.86 | 99.56 | 79.55 | 79 | 78.1 | 20.78 | 15.53 | 14.67 | 2.03 |
| **bio13** | 138.73 | 138.68 | 138.67 | 121.39 | 107.53 | 88.39 | 31.2 | 30.72 | 30.63 | 29.55 | 2.17 | 2.07 |
| **bio15** | 10.66 | 8.85 | 8.17 | 8.14 | 7.85 | 7.22 | 7.11 | 7.03 | 7.03 | 6.95 | 2.18 | 2.1 |
| **bio19** | 107.15 | 107.15 | 107.12 | 104.88 | 104.69 | 102.45 | 100.32 | 5.14 | 4.68 | 3.15 | 2.84 | 2.76 |
| **bio14** | 8.44 | 8.37 | 8.14 | 8.14 | 6.63 | 6.44 | 6.39 | 3.55 | 3.51 | 3.4 | 3.4 | 3.4 |
| bio3 | 73.24 | 70.46 | 54.68 | 54.01 | 49.45 | 45.87 | 42.61 | 42.52 | 39.24 | 24.51 | 23.81 | |
| bio12 | 127.45 | 125.84 | 94.92 | 84 | 84 | 78.01 | 33.74 | 31.78 | 31.55 | 30.69 | | |
| bio5 | 486.27 | 433.01 | 286.76 | 261.61 | 80.33 | 74.84 | 74.39 | 73.87 | 68.47 | | | |
| bio9 | 932.58 | 598.13 | 315.43 | 286.33 | 182.45 | 89.36 | 88.72 | 87.37 | | | | |
| bio17 | 144.86 | 144.59 | 142.72 | 140.43 | 134.51 | 134.42 | 122.51 | | | | | |
| bio18 | 1056.66 | 1022.86 | 998.83 | 223.02 | 203.99 | 170.78 | | | | | | |
| bio1 | 1778.75 | 1061.21 | 417.02 | 415.74 | 237.75 | | | | | | | |
| bio10 | 2682.28 | 1612.89 | 865.16 | 857.48 | | | | | | | | |
| bio16 | 1249.69 | 1226.58 | 1226.58 | | | | | | | | | |
| bio4 | 8066.32 | 1902.44 | | | | | | | | | | |
| bio11 | 10386.53 | | | | | | | | | | | |
| bio7 | alias | | | | | | | | | | | |
| **Geo-climatic** | | | | | | | | | | | | |
| **Asp** | 1.06 | | | | | | | | | | | |
| **NPP** | 1.84 | | | | | | | | | | | |
| **ann_WS** | 2.19 | | | | | | | | | | | |
| **SC** | 2.4 | | | | | | | | | | | |
| **ann_SR** | 2.49 | | | | | | | | | | | |
| **PV** | 2.7 | | | | | | | | | | | |
| **AI** | 2.83 | | | | | | | | | | | |

| | | | | | | | | |
|---|---|---|---|---|---|---|---|---|
| **PET** | 2.87 | | | | | | | |
| **RH** | 2.9 | | | | | | | |
| **ann_WV** | 3.93 | | | | | | | |
| **SM** | 4.05 | | | | | | | |
| **SpH** | 4.74 | | | | | | | |
| **Habitat heterogeneity** | | | | | | | | |
| **CV** | 4.53 | 4.53 | 4.46 | 4.28 | 3.88 | 3.59 | 3.29 | 2.94 |
| **Corr** | 10.03 | 10.03 | 10.02 | 9.94 | 6.92 | 4.89 | 4.47 | 3.63 |
| **Uni** | 66.02 | 65.82 | 65.54 | 55.21 | 55.05 | 40 | 5.82 | 5.11 |
| **Even** | 20.14 | 19.7 | 19.5 | 15.27 | 15.27 | 12.1 | 10.35 | 5.82 |
| **Cont** | 104.14 | 100.87 | 10.24 | 10.19 | 8.12 | 7.61 | 7.1 | 7.07 |
| **Range** | 28.75 | 28.66 | 27.72 | 20.44 | 17 | 14.27 | 14.04 | 8.09 |
| **Var** | 56.4 | 55.3 | 54.77 | 54.19 | 12.3 | 9.58 | 9.47 | 9.09 |
| Homo | 168.81 | 94.6 | 28.9 | 23.57 | 19.57 | 15.64 | 15.5 | |
| Max | 63.34 | 63.02 | 62.46 | 60.78 | 57.04 | 56.99 | | |
| Simpson | 181.29 | 162.75 | 155.51 | 63.01 | 61.44 | | | |
| Std | 171.42 | 166.25 | 166.21 | 151.01 | | | | |
| Shannon | 430.24 | 217.86 | 198.96 | | | | | |
| Dissimilarity | 244.28 | 230.06 | | | | | | |
| Entrophy | 671.43 | | | | | | | |
| **Growing days** | | | | | | | | |
| **GDD** | 1.72 | 1.06 | | | | | | |
| **GSL** | 4.06 | 1.14 | | | | | | |
| **GST** | 2.48 | 1.96 | | | | | | |
| **LGD** | 775.59 | 2.02 | | | | | | |
| FGD | 818.15 | | | | | | | |
| **Ultra-violet radiation** | | | | | | | | |
| **UVB2** | 4895.44 | 4885.71 | 576.88 | 17.84 | 2.63 | | | |
| **UVB4** | 1462.14 | 1282.64 | 971.34 | 771.81 | 2.63 | | | |
| UVB6 | 7442.02 | 2982.99 | 972.09 | 955.68 | | | | |
| UVB3 | 2974.09 | 2810.56 | 1575.11 | | | | | |
| UVB5 | 10344.81 | 7581.51 | | | | | | |
| UVB1 | 13196.4 | | | | | | | |

b) Pure pink morph (PP)

| Variables | Run1 | Run2 | Run3 | Run4 |
|---|---|---|---|---|
| **Bioclimatic variables** | | | | |
| **bio3** | 25184.36 | 16897.72 | 287.87 | 1.1 |
| **bio14** | 5.64 | 1.9 | 1.73 | 1.26 |
| **bio7** | 16303.17 | 5471.13 | 275.07 | 1.42 |
| **bio5** | 23822.5 | 29.95 | 2.89 | 1.44 |
| bio2 | 8467.54 | 2897.37 | 582.73 | |
| bio4 | 23630.99 | 20809.39 | | |
| bio6 | 53466.17 | | | |
| bio1 | alias | | | |
| bio8 | alias | | | |
| bio9 | alias | | | |
| bio10 | alias | | | |
| bio11 | alias | | | |
| bio12 | alias | | | |
| bio13 | alias | | | |
| bio15 | alias | | | |
| bio16 | alias | | | |
| bio17 | alias | | | |
| bio18 | alias | | | |
| bio19 | alias | | | |
| **Geo-climatic** | | | | |
| **ann_WV** | 169.6 | 23.55 | 23.54 | 2.12 |
| **RH** | 5256.57 | 110.25 | 3.24 | 2.68 |
| **PET** | 44970.32 | 33.79 | 19.91 | 6.5 |
| **Asp** | 1220.41 | 26.16 | 8.85 | 8.57 |
| PV | 1715.46 | 45.74 | 39.94 | |
| NPP | 4349.64 | 168.59 | | |

| | | | | |
|---|---|---|---|---|
| ann_SR | 53035.76 | | | |
| SC | alias | | | |
| SM | alias | | | |
| SpH | alias | | | |
| ann_WS | alias | | | |
| AI | alias | | | |
| **Habitat heterogeneity** | | | | |
| **Homo** | 127.75 | 125.18 | 32.29 | 2.32 |
| **CV** | 247.57 | 217.32 | 40.58 | 4.15 |
| **Corr** | 5945.91 | 254.04 | 28.41 | 5.21 |
| **Even** | 128.04 | 13.52 | 8.95 | 5.55 |
| Cont | 2033.18 | 1662.67 | 40.59 | |
| Dissimilarity | 21419.27 | 2043.63 | | |
| Entrophy | 22665.19 | | | |
| Range | alias | | | |
| Shannon | alias | | | |
| Simpson | alias | | | |
| Std | alias | | | |
| Uni | alias | | | |
| Var | alias | | | |
| Max | alias | | | |
| **Growing days** | | | | |
| **GSL** | 4.32 | 1.1 | 1.01 | |
| **GDD** | 2.5 | 1.36 | 1.19 | |
| **LGD** | 98.63 | 12.24 | 1.21 | |
| GST | 1537.7 | 13.15 | | |
| FGD | 2251.61 | | | |
| **Ultra-violet radiation** | | | | |
| **UVB2** | 2557.78 | 1.2 | | |

| | | |
|---|---|---|
| **UVB1** | 6407.44 | 1.2 |
| UVB3 | 12279.37 | |
| UVB4 | alias | |
| UVB5 | alias | |
| **UVB6** | alias | |

c) Yellow-white morph (YW)

| variables | Run1 | Run2 | Run3 | Run4 | Run5 | Run6 | Run7 | Run8 | Run9 | Run10 | Run11 | Run12 | Run13 | Run14 |
|---|---|---|---|---|---|---|---|---|---|---|---|---|---|---|
| **Bio-climatic variables** | | | | | | | | | | | | | | |
| **bio3** | 750.33 | 414.07 | 380.37 | 267.83 | 251.18 | 243.12 | 169.5 | 156.12 | 2.08 | 1.99 | 1.98 | 1.41 | 1.4 | 1.32 |
| **bio4** | 1829.83 | 1467.34 | 1460.36 | 1274.96 | 602.67 | 580.6 | 549.92 | 268.85 | 4.14 | 3.95 | 3.6 | 2.39 | 1.85 | 1.58 |
| **bio12** | 84.88 | 76.35 | 76.03 | 74.17 | 71.46 | 71.01 | 60.85 | 45.66 | 41.12 | 24 | 24 | 23.43 | 1.78 | 1.67 |
| **bio5** | 6339.44 | 6041.49 | 5119.58 | 2002.2 | 1721.38 | 419.99 | 356.36 | 44.8 | 44.64 | 39.15 | 34.54 | 1.99 | 1.82 | 1.68 |
| **bio15** | 25.74 | 25.69 | 23.48 | 23.47 | 19.82 | 18.12 | 16.48 | 16.26 | 15.3 | 11.3 | 10.24 | 9.08 | 2.68 | 2.35 |
| **bio17** | 647.16 | 634.67 | 528.66 | 521.48 | 264.59 | 233.36 | 147.74 | 116.22 | 81.88 | 74.62 | 14.2 | 14 | 13.72 | 2.44 |
| bio14 | 31.61 | 29.33 | 27.26 | 27.17 | 23.54 | 22.78 | 22.53 | 22.44 | 19.49 | 18.08 | 14.93 | 14.93 | 14.64 | |
| bio13 | 141 | 137.24 | 133.64 | 132.03 | 128.18 | 128 | 122.69 | 121.56 | 119.9 | 29.94 | 29.88 | 29.52 | | |
| bio8 | 63.06 | 62.32 | 60.48 | 60.39 | 55.22 | 55.11 | 53.49 | 53.36 | 52.46 | 43.04 | 37.65 | | | |
| bio19 | 722.3 | 690.21 | 570.74 | 564.11 | 321.13 | 266.31 | 179.68 | 131.98 | 93.29 | 77.86 | | | | |
| bio16 | 249 | 246.53 | 240.31 | 232.6 | 228.12 | 227.17 | 225.92 | 202.71 | 189.05 | | | | | |
| bio2 | 2241.19 | 1953.52 | 1953.52 | 1318.59 | 1049.6 | 1048.73 | 328.89 | 322.49 | | | | | | |
| bio9 | 1553.21 | 1419.95 | 1281.97 | 1133.34 | 973.58 | 800.85 | 682.44 | | | | | | | |
| bio7 | 4739.92 | 4152.14 | 4062.7 | 1706.57 | 1376.55 | 1318.6 | | | | | | | | |
| bio10 | 4521.38 | 4521.37 | 4517.22 | 4516.12 | 2529.55 | | | | | | | | | |
| bio11 | 16011.67 | 11502.29 | 10788.91 | 9869.68 | | | | | | | | | | |
| bio6 | 20560.55 | 15885.16 | 13360.55 | | | | | | | | | | | |
| bio1 | 28164.74 | 18603.49 | | | | | | | | | | | | |
| bio18 | alias | | | | | | | | | | | | | |
| **Geo-climatic** | | | | | | | | | | | | | | |
| **Asp** | 1.47 | 1.21 | 1.19 | | | | | | | | | | | |

| | | | | | | | | | | |
|---|---|---|---|---|---|---|---|---|---|---|
| **SpH** | 20.38 | 1.86 | 1.55 | | | | | | | |
| **PV** | 3.3 | 3.3 | 1.62 | | | | | | | |
| **NPP** | 4.79 | 4.12 | 3.37 | | | | | | | |
| **SM** | 4.28 | 4.24 | 3.73 | | | | | | | |
| **PET** | 5.61 | 4.97 | 4.97 | | | | | | | |
| **AI** | 6.42 | 6.1 | 5.06 | | | | | | | |
| **ann_SR** | 9.48 | 5.44 | 5.44 | | | | | | | |
| **ann_WV** | 27.52 | 7.42 | 6.6 | | | | | | | |
| **ann_WS** | 8.46 | 8.42 | 7.32 | | | | | | | |
| RH | 10.93 | 10.57 | | | | | | | | |
| SC | 29.15 | | | | | | | | | |
| **Habitat heterogeneity** | | | | | | | | | | |
| **Cont** | 294.43 | 18.42 | 18.09 | 17.83 | 15.46 | 8.26 | 7.86 | 7.2 | 6.53 | 3.11 |
| **Corr** | 23.3 | 19.34 | 19.13 | 17.04 | 16.81 | 7.32 | 7.21 | 4.13 | 4.12 | 4.02 |
| **Max** | 25.13 | 25.1 | 23.43 | 23.28 | 18.57 | 18.21 | 9.18 | 8.83 | 8.68 | 4.32 |
| **Even** | 18.74 | 18.72 | 18.4 | 17.54 | 17.07 | 7.21 | 6.18 | 4.5 | 4.47 | 4.43 |
| **CV** | 18.76 | 18.48 | 16.98 | 16.9 | 16.21 | 16.18 | 16.17 | 16.17 | 4.94 | 4.89 |
| Homo | 139.55 | 58.4 | 55.47 | 55.32 | 19.1 | 17.25 | 13.87 | 12.38 | 12.31 | |
| Var | 184.31 | 165.91 | 165.88 | 38.61 | 38.49 | 27.44 | 24.37 | 17.75 | | |
| Range | 107.9 | 106.86 | 94.67 | 61.45 | 61.36 | 28.92 | 25.12 | | | |
| Uni | 107.82 | 76.24 | 75.06 | 70.62 | 66.81 | 33.74 | | | | |
| Simpson | 398.72 | 397.76 | 329.71 | 327.29 | 209.96 | | | | | |
| Entrophy | 719.98 | 450.33 | 351.1 | 333.33 | | | | | | |
| Std | 523.82 | 504.21 | 487.39 | | | | | | | |
| Shannon | 542 | 515.27 | | | | | | | | |
| Dissimilarity | 1014.16 | | | | | | | | | |
| **Growing days** | | | | | | | | | | |
| **LGD** | 58.93 | 1.79 | | | | | | | | |
| **GSL** | 7.28 | 1.79 | | | | | | | | |

| | | | | | |
|---|---|---|---|---|---|
| **GST** | 8.96 | 2.62 | | | |
| **GDD** | 2.86 | 2.85 | | | |
| FGD | 60.95 | | | | |
| **Ultra-violet radiation** | | | | | |
| **UVB4** | 404.52 | 329.49 | 285.56 | 104.9 | 1 |
| **UVB2** | 2944.48 | 986 | 704.6 | 13.63 | 1 |
| **UVB6** | 440.72 | 440.72 | 124.37 | 114.07 | |
| UVB3 | 792 | 788.97 | 719.81 | | |
| UVB1 | 2322.54 | 1575.59 | | | |
| UVB5 | 4595.42 | | | | |

d) Pure yellow morph

| **Variables** | **Run1** | **Run2** | **Run3** | **Run4** | **Run5** | **Run6** | **Run7** | **Run8** | **Run9** | **Run10** | **Run11** | **Run12** | **Run13** |
|---|---|---|---|---|---|---|---|---|---|---|---|---|---|
| **Bio-climatic** | | | | | | | | | | | | | |
| **bio8** | 189.72 | 189.15 | 180.75 | 180.56 | 151.18 | 58.26 | 57.55 | 57.42 | 52.41 | 2.26 | 1.78 | 1.78 | 1.74 |
| **bio13** | 160.31 | 156.75 | 155.53 | 146.61 | 137.55 | 134.83 | 132.37 | 38.7 | 38.64 | 34.51 | 32.39 | 1.85 | 1.84 |
| **bio2** | 235.3 | 196.6 | 167.58 | 163.24 | 161.44 | 159.36 | 17.83 | 17.69 | 17.15 | 16.99 | 2.56 | 2.2 | 2.16 |
| **bio15** | 10.52 | 10.51 | 9.75 | 9.19 | 8.87 | 8.85 | 8.07 | 7.24 | 7.23 | 6.18 | 6.02 | 3.72 | 3 |
| **bio3** | 192.82 | 176.68 | 173.9 | 173.77 | 173.76 | 173.76 | 52.28 | 52.1 | 49.61 | 49.48 | 4.01 | 3.18 | 3.1 |
| **bio14** | 79.84 | 77.64 | 75.81 | 73.92 | 66.81 | 59.41 | 54.71 | 53.46 | 32.69 | 26.91 | 21.59 | 21.01 | 4.61 |
| bio19 | 129.06 | 128.8 | 124.29 | 122.55 | 116.74 | 96.23 | 87.97 | 85.08 | 36.2 | 28.99 | 23.7 | 23.14 | |
| bio12 | 98.25 | 98.08 | 95.94 | 95.09 | 70.6 | 68.28 | 61.85 | 47.25 | 47.2 | 40.42 | 37.04 | | |
| bio4 | 290.74 | 288.35 | 277.29 | 155.47 | 76.69 | 68.81 | 65.37 | 64.9 | 64.88 | 50.88 | | | |
| bio9 | 252.89 | 248.22 | 241.59 | 235.61 | 231.39 | 121.65 | 120.84 | 119.89 | 106.85 | | | | |
| bio17 | 160.53 | 155.51 | 154.64 | 153.66 | 141.73 | 140.01 | 139.33 | 133.01 | | | | | |
| bio16 | 248.1 | 245.31 | 243.61 | 234.87 | 184.86 | 176.94 | 175.88 | | | | | | |
| bio7 | 3328.08 | 3282.16 | 446.44 | 390.83 | 278.37 | 210.49 | | | | | | | |
| bio5 | 7559.62 | 7511.53 | 1134.1 | 1100.02 | 289.09 | | | | | | | | |
| bio10 | 2620.74 | 2517.69 | 1927.95 | 1241.97 | | | | | | | | | |

| | | | | | | | | | | |
|---|---|---|---|---|---|---|---|---|---|---|
| bio11 | 4570.84 | 4568.05 | 2235.78 | | | | | | | |
| bio6 | 14055.55 | 13985.79 | | | | | | | | |
| bio1 | 21675.28 | | | | | | | | | |
| bio18 | alias | | | | | | | | | |
| **Geo-climatic** | | | | | | | | | | |
| **Asp** | 1.37 | 1.36 | | | | | | | | |
| **PV** | 1.62 | 1.62 | | | | | | | | |
| **SpH** | 2.27 | 1.76 | | | | | | | | |
| **ann_SR** | 2.12 | 2.09 | | | | | | | | |
| **SC** | 2.94 | 2.93 | | | | | | | | |
| **AI** | 3.84 | 3.75 | | | | | | | | |
| **NPP** | 4.87 | 4.4 | | | | | | | | |
| **ann_WS** | 6.65 | 4.47 | | | | | | | | |
| **RH** | 7.44 | 5.9 | | | | | | | | |
| **SM** | 6.44 | 6.41 | | | | | | | | |
| **PET** | 9.44 | 9 | | | | | | | | |
| ann_WV | 12.35 | | | | | | | | | |
| **Habitat heterogeneity** | | | | | | | | | | |
| **Cont** | 180.38 | 148.8 | 148.79 | 16.2 | 15.83 | 12.79 | 10.61 | 6.92 | 3.92 | 3.04 |
| **Max** | 16.43 | 16.13 | 15.5 | 15.46 | 11.43 | 11.4 | 9.32 | 8.05 | 3.11 | 3.1 |
| **Corr** | 21.66 | 19.98 | 12.83 | 12.7 | 9.32 | 6.58 | 6.57 | 3.76 | 3.52 | 3.39 |
| **CV** | 12.62 | 12.33 | 11.41 | 10.6 | 10.5 | 9.93 | 9.84 | 8.89 | 8.88 | 3.47 |
| **Even** | 18.56 | 17.27 | 17.26 | 16.6 | 11.19 | 9.5 | 7.55 | 3.79 | 3.78 | 3.78 |
| Var | 126.23 | 126.22 | 20.15 | 18.78 | 16.13 | 15 | 12.82 | 11.62 | 11.37 | |
| Homo | 121.19 | 115.84 | 113.78 | 50.05 | 43.17 | 24 | 14.14 | 14.05 | | |
| Range | 56.37 | 51.03 | 43.05 | 38.96 | 29.26 | 25.49 | 21.12 | | | |
| Uni | 224.14 | 112.15 | 108.43 | 102.72 | 42.28 | 27.17 | | | | |
| Entrophy | 574.31 | 247.33 | 238.37 | 217.02 | 156.09 | | | | | |

| | | | | | |
|---|---|---|---|---|---|
| Simpson | 551.73 | 270.83 | 268.08 | 267.93 | |
| Dissimilarity | 419.71 | 374.59 | 371.51 | | |
| Std | 466.45 | 400.38 | | | |
| Shannon | 1195.93 | | | | |
| **Growing days** | | | | | |
| **GST** | 1.05 | | | | |
| **LGD** | 1.55 | | | | |
| **GDD** | 1.56 | | | | |
| **FGD** | 2.15 | | | | |
| **GSL** | 2.37 | | | | |
| **Ultra-violet radiation** | | | | | |
| **UVB4** | 285.94 | 266.17 | 95.56 | 94.82 | 1.46 |
| **UVB3** | 799.57 | 794.23 | 273.45 | 45.76 | 1.46 |
| UVB6 | 345.44 | 345.28 | 324.92 | 210.65 | |
| UVB1 | 1301.88 | 1007.74 | 580.23 | | |
| UVB2 | 1846.86 | 1099.66 | | | |
| UVB5 | 1900.95 | | | | |

**Table S16.** Consensus land-cover selection for each cryptic species of *S. chamaejasme* based on the presence of the number of occurrence records.

| **Consensus Land-cover** | **Count (% of occurrence)** | | | |
|---|---|---|---|---|
| | Pure pink morph (PP) | Red-white morph (RW) | Pure yellow morph (YY) | Yellow-white morph (YW) |
| Evergreen/deciduous needleleaf trees (LULC1) | **9 (100)** | **96 (54.55)** | **75 (91.46)** | **29 (72.5)** |
| Evergreen broadleaf trees (LULC 2) | **9 (100)** | 19 (10.80) | 41 (50) | 19 (47.5) |
| Deciduous broadleaf trees (LULC 3) | 8 (88.89) | 27 (15.34 | 27 (32.93) | 6 (15) |
| Mixed/other trees (LULC 4) | **9 (100)** | 84 (47.73) | **76 (92.68)** | **29 (72.5)** |
| Shrubs (LULC 5) | 6 (66.67) | 60 (34.09) | **58 (70.73)** | 13 (32.5) |
| Herbaceous vegetation (LULC 6) | **9 (100)** | **170 (96.59)** | **79 (96.34)** | **40 (100)** |
| Cultivated and managed vegetation (LULC 7) | 6 (66.67) | **173 (98.30)** | **78 (95.12)** | **38 (95)** |
| Regularly flooded vegetation (LULC 8) | 0 | 0 | 1 (1.22) | 0 |
| Urban/built-up (LULC 9) | 0 | 14 (7.95) | 8 (9.76) | 1 (2.5) |
| Snow/ice (LULC 10) | 0 | 14 (7.95) | 4 (4.88) | 2 (5) |
| Barren (LULC 11) | 0 | 37 (21.02) | 1 (1.22) | 2 (5) |
| Open water (LULC 12) | 0 | 8 (4.55) | 10 (12.20) | 1 (2.5) |
| Threshold (%) | 100% | >50% | >50% | >50% |

% of occurrence (count/total occurrence*100)

**Table S17.** Model evaluation indices for the ensemble species distribution modeling (eSDM) of each cryptic species of *S. chamaejasme* using Biomod2 in R-programming language.

a) Bioclimatic variables only

| **Models** | | ***Indices for each* cryptic species** | | | | | | | | | | | |
|---|---|---|---|---|---|---|---|---|---|---|---|---|---|
| | | TSS pp | TSS rw | TSS yw | TSS yy | Kappa pp | Kappa rw | Kappa yw | Kappa yy | AUC pp | AUC rw | AUC yw | AUC yy |
| Generalized Additive Models | GAM | 0.996 | 0.881 | 0.916 | 0.888 | 0.888 | 0.802 | 0.525 | 0.748 | 1 | 0.983 | 0.976 | 0.985 |
| Generalized Boosted Models | GBM | 0.996 | 0.897 | 0.934 | 0.949 | 0.888 | 0.825 | 0.727 | 0.826 | 1 | 0.987 | 0.992 | 0.994 |
| Generalized Linear Models | GLM | 0.882 | 0.687 | 0.757 | 0.738 | 0.154 | 0.483 | 0.269 | 0.476 | 0.949 | 0.893 | 0.922 | 0.929 |
| Classification Tree Analysis | CTA | 0.841 | 0.62 | 0.572 | 0.693 | 0.713 | 0.663 | 0.488 | 0.639 | 0.978 | 0.852 | 0.845 | 0.891 |
| Artificial Neural Network | ANN | 0.93 | 0.81 | 0.767 | 0.787 | 0.154 | 0.781 | 0.543 | 0.728 | 0.949 | 0.943 | 0.894 | 0.953 |
| Flexible Discriminant Analysis | FDA | 0.88 | 0.763 | 0.79 | 0.748 | 0.623 | 0.684 | 0.44 | 0.657 | 0.939 | 0.956 | 0.959 | 0.913 |
| Multiple Adaptive Regression Splines | MARS | 0.874 | 0.827 | 0.859 | 0.888 | 0.665 | 0.699 | 0.485 | 0.709 | 0.983 | 0.965 | 0.967 | 0.984 |
| Random Forest | RF | 0.994 | 0.989 | 0.965 | 0.987 | 0.874 | 0.962 | 0.856 | 0.931 | 1 | 0.999 | 0.997 | 0.999 |
| Surface Range Envelop | SRE | 0.657 | 0.622 | 0.813 | 0.647 | 0.423 | 0.575 | 0.356 | 0.504 | 0.828 | 0.811 | 0.907 | 0.823 |
| Maximum Entropy | MaxEnt | 0.829 | 0.852 | 0.866 | 0.761 | 0.102 | 0.845 | 0.47 | 0.792 | 0.919 | 0.977 | 0.966 | 0.912 |
| Maxnet | MaxEnt2 | 0.829 | 0.85 | 0.871 | 0.889 | 0.12 | 0.757 | 0.46 | 0.707 | 0.92 | 0.978 | 0.965 | 0.979 |

b) Environmental variables

| **Models** | | ***Indices for each* cryptic species** | | | | | | | | | | | |
|---|---|---|---|---|---|---|---|---|---|---|---|---|---|
| | | TSS pp | TSS rw | TSS yw | TSS yy | Kappa pp | Kappa rw | Kappa yw | Kappa yy | AUC pp | AUC rw | AUC yw | AUC yy |
| Generalized Additive Models | GAM | 0.996 | 0.766 | NA | 0.993 | 0.888 | 0.655 | NA | 0.944 | 1 | 0.954 | NA | 1 |
| Generalized Boosted Models | GBM | 0.996 | 0.922 | 0.91 | 0.97 | 0.888 | 0.864 | 0.815 | 0.921 | 1 | 0.991 | 0.988 | 0.998 |
| Generalized Linear Models | GLM | 0.916 | 0.881 | 0.874 | 0.889 | 0.874 | 0.802 | 0.45 | 0.75 | 0.993 | 0.983 | 0.964 | 0.984 |
| Classification Tree Analysis | CTA | 0.911 | 0.789 | 0.737 | 0.698 | 0.776 | 0.789 | 0.619 | 0.717 | 0.989 | 0.962 | 0.901 | 0.934 |
| Artificial Neural Network | ANN | 0.889 | 0.627 | 0.848 | 0.784 | 0.1 | 0.552 | 0.397 | 0.524 | 0.945 | 0.854 | 0.924 | 0.906 |
| Flexible Discriminant Analysis | FDA | 0.886 | 0.839 | 0.81 | 0.876 | 0.76 | 0.784 | 0.582 | 0.801 | 0.943 | 0.978 | 0.909 | 0.956 |
| Multiple Adaptive Regression splines | MARS | 0.996 | 0.897 | 0.931 | 0.912 | 0.888 | 0.844 | 0.72 | 0.748 | 1 | 0.989 | 0.99 | 0.985 |
| Random Forest | RF | 0.991 | 0.995 | 0.965 | 0.993 | 0.888 | 0.98 | 0.874 | 0.944 | 0.999 | 1 | 0.998 | 0.999 |
| Surface Range Envelop | SRE | 0.556 | 0.374 | 0.478 | 0.284 | 0.713 | 0.458 | 0.436 | 0.387 | 0.778 | 0.687 | 0.739 | 0.642 |
| Maximum Entropy | MaxEnt | 0.777 | 0.89 | 0.913 | 0.81 | 0.822 | 0.924 | 0.684 | 0.819 | 0.889 | 0.977 | 0.988 | 0.933 |
| Maxnet | MaxEnt2 | 0.883 | 0.904 | 0.9 | 0.943 | 0.874 | 0.907 | 0.679 | 0.919 | 0.985 | 0.991 | 0.986 | 0.997 |

Kappa = *Cohen's kappa*; TSS = True Skill Statistics; AUC = Area Under the Curve; rw, Red-white morph; pp, Pure red morph; yw, Yellow-white morph; yy, Pure yellow morph

### Supplementary Methods

#### Sampling, library preparation, and RAD sequencing

The study involved 251 individuals of *S. chamaejasme* populations, which encompass four color morphs (Zhang *et al.*, 2016) and were collected from the Qinghai Tibetan Plateau (QTP), the Hengduan Mountains (HM), and northeast China (Qiu, 2008) (Table **S1**). As outgroups, three single-individual of closely related species (*Daphne longilobata* (Lecomte) Turrill, *Daphne tangutica* Maxim., and *Wikstroemia ligustrina* Rehder) were included. Voucher specimens for each population were deposited in the Herbaria of Kunming Institute of Botany, CAS (KUN). The purity and integrity of an extracted genomic DNA were analyzed by 1% agarose gel electrophoresis and quantified using a NanoDrop® spectrophotometer (ND-2000, Thermo Fisher Scientific, USA). DNA concentration was accurately quantified with a QubitTM 2.0 fluorometer (Invitrogen, Carlsbad, CA, USA) and stored at −20 °C for further analysis.

To construct the restriction site-associated DNA (RAD) library and perform sequencing, the genomic DNA underwent several steps. First, it was digested with the EcoRI restriction enzyme in a 30-µl reaction, followed by ligation of the P1 adapter using T4 ligase. The resulting fragments were then pooled, randomly sheared, and size-selected to a range of 300-700 bp. Subsequently, a second adapter (P2) was ligated to the fragments. The ligation products underwent purification and PCR amplification. Gel purification and size selection to the range of 300-700 bp was performed. The library was qualified and quantified using the Agilent 2100 Bioanalyzer and quantitative (q)PCR. Sequencing was carried out on the Illumina HiSeq 2000 platform, generating paired-end reads with a length of 150 bp. Two lanes were used for sequencing, and each sample produced approximately 3.56 Gb of raw data. The raw sequencing reads will be deposited in the NCBI Sequence Read Archive with an accession number.

#### Illumina paired-end reads processing

The raw Illumina reads underwent initial quality checks using FastQC v0.11.4 software (Andrews, 2010) to assess per-base sequence quality and identify overrepresented sequences indicating adapter contamination. Reads that did not meet quality thresholds, such as having nucleotide quality above Q30 (<0.1% sequencing error) and more than 99% above Q20 (<1% sequencing error), raw reads with ≥10% unidentified nucleotides (N), >50% bases having Phred quality <5, with >10 nt aligned to the adapter, or enzyme sequences, were discarded.

For data processing, Stacks v.2.55 (Catchen *et al.*, 2013; http://catchenlab.life.illinois.edu/stacks/) software was utilized. Sequence reads were demultiplexed and filtered using *process_radtags*, and *de novo* assemblies were generated. Reads with an average Phred score of ≥30, an unambiguous barcode, and a restriction cut site were retained. The *denovo_map.pl* wrapper program executed three main pipeline components (*ustacks*, *cstacks*, and *sstacks*), along with *tsv2bam*, *gstacks*, and *populations* modules (Catchen *et al.*, 2013). The important parameters for Stacks are *M* parameter of *ustacks*, which controls the number of mismatches allowed between the two alleles in a sample; and the n parameter of *cstacks*, which controls the number of mismatches allowed between any two alleles in the population. In addition, the *M* parameter of *ustacks* controls the number of identical reads required to create a new putative allele (Rochette & Catchen, 2017). The Stacks parameters were optimized based on Rochette & Catchen (2017) suggested protocol. Various values of *M* and *n* (ranging from 1 to 9 with *M*=*n* and *m*=3) were tested to achieve optimal data assembly. For final assemblies, a minimum of three identical, raw reads were required to identify loci in each individual and call SNPs (*denovo_map -m* parameter). Three mismatches were allowed between loci when processing a single individual (-*M* parameter) and building the catalog (-*n* parameter). These assembly options (-*m* 3 -*M* 3 -*n* 3) resulted in the identification of 23,215 loci. Finally, the *populations* pipeline was run with additional filters to create assemblies for subsets of the entire dataset for downstream analyses.

To create subsets of the dataset for downstream analyses, the *populations* pipeline was run with additional filters. A filtering parameter of -r 0.8 was used to process loci with ≥80% individuals in a population, and loci showing high heterozygosity (>0.5) were excluded. The dataset was further filtered based on minor allele frequency (MAF) >0.01, and only biallelic SNPs were retained. Unlinked loci were selected using the *write_single_snp* flag. For specific analyses, separate *populations* pipeline was called for Treemix and Bayesian phylogeny and phylogeography (BPP). Treemix analysis utilized cryptic species-wise categorized individuals, resulting in 1,716 loci containing 173 SNPs. BPP analysis involved 102 selected individuals, resulting in 25,049 loci containing 7,208 SNPs. These datasets were used for their respective analyses.

**Genetic distinctness and clustering**

We estimated the genetic diversity indices, including nucleotide diversity (π), expected heterozygosity ($H_e$), observed heterozygosity ($H_o$), and inbreeding coefficient ($F_{IS}$) using the *populations* pipeline in Stacks v.2.55 (Catchen *et al.*, 2013). The population and cryptic species-wise estimation of genetic differentiation indices include $\Phi_{ST}$means (AMOVA-based measure of $F_{ST}$) as the total genetic variance fixation index, $D_{XY}$means as the absolute measure of locus differentiation among genetic populations excluding polymorphisms segregating within the two populations, and $F_{ST}$'means as the haplotype measure of $F_{ST}$ that is scaled to the theoretical maximum $F_{ST}$ value at this locus. Furthermore, the $F_{ST}$ was graphically visualized with chord plots using the '*chorddiag'* v.0.1.3 R-package (Flor, 2021).

A combination of fastSTRUCTURE (Raj *et al.*, 2014), DAPC (Jombart *et al.*, 2010), and PCA were used to infer population genetic structure. The Bayesian clustering program fastSTRUCTURE determined the most likely number of genetic clusters at different ancestral populations (K) numbers ranging from 1 to 10 using 10-fold cross-validation. We compared the cross-validation profile identifying the number of K for which the prediction error is minimized with the output of the function 'chooseK.py' based on admixture proportion (Q values) to find the best model that explains the structure. The function computes two values for *K*: one that maximizes the log-marginal likelihood lower bound (LLBO) of the dataset (K*ε) to identify strong structure, and another one reports the model components that have a cumulative ancestry contribution of at least 99% (Køc) to capture additional weak underlying structure (Raj *et al.*, 2014). To evaluate the repeatability across runs and rule out true multimodality (as opposed to cluster labels switching), we ran the program CLUMPP v.1.1.2 using the Greedy algorithm (Jakobsson & Rosenberg, 2007) and graphically visualized the results using DISTRUCT (Rosenberg 2004). The discriminant functions that maximize the differentiation between groups while minimizing the variation within them were evaluated by DAPC using the optimal number of principal components (α-score optimization). We employed 5,800 SNP datasets, each of which contained a single SNP sampled at random from each locus. We chose the number of clusters by running *K-means* clustering at different values of *K* and compared different numbers of clusters using the Bayesian information criterion (*BIC*) according to Jombart *et al.* (2010). The individual and cryptic species-wise PCA on allele frequency data of 5,800 SNPs were also performed using the R-package '*adegenet'* v2.1.3 (Jombart, 2008; Jombart & Ahmed, 2011).

The fineRADstructure v.0.3.2 (Malinsky, 2018), a modified version of fineSTRUCTURE software (Lawson *et al.*, 2012) mainly designed for RADseq data that does not require information about the location of loci on chromosomes or phased haplotypes, was used to estimate population structure through shared ancestry. The reordered RAD loci according to linkage disequilibrium in *sampleLD.r* script was analyzed for the co-ancestry matrix in RAD painter (scripts implemented in fineRADstructure package). Finally, fineSTRUCTURE v1.1 was used to allocate the individuals to populations, which involved 100,000 Markov chain Monte Carlo (MCMC) iterations, 100,000 burn-in sampling per 1,000 iterations, and 10,000 hill-climbing iterations for constructing a tree. We utilized the R scripts *fineradstructureplot.r* and *finestructurelibrary.r* (available at *http://cichlid.gurdon.cam.ac.uk/fineRADstructure.html*) to display the results.

**Species tree inference and divergence**

The relationships among the individuals were reconstructed using the phylogenomic tree through the concatenated super matrix of 23,739 loci by maximum likelihood (ML) and Bayesian inference (BI) using IQ-TREE v.2.1.4_beta (Nguyen *et al.*, 2015) and MrBayes v.3.2.7 (Ronquist *et al.*, 2012), respectively. The super matrix was constructed using 23,215 loci of 251 individuals and 23,726 loci of three closely related outgroup species *Daphne longilobata* (Chen0002), *Daphne tangutica* (Chen0008), and *Wikstroemia ligustrina* (Chen0006). Due to computational restrictions, IQ-TREE's optimum nucleotide substitution model was limited to the general time-reversible (GTR) model (Kalyaanamoorthy *et al.*, 2017). Clade support was calculated for 50 runs with 1,000 µLtra-fast bootstrap replicates (Minh *et al.*, 2013; Hoang *et al.*, 2018) and the SH-aLRT branch test with 1,000 replicates. The Akaike information criterion (*AIC*), corrected *AIC* (*AICc*), and Bayesian information criterion (*BIC*) from ModelTest-NG v.0.1.6 (Darriba *et al.*, 2020) were used to evaluate the best-fitted substitution model for BI analysis. The Bayesian analysis was assessed using four independent MCMC for $2\times10^7$ generations each, sampling every 1,000 generations and removing the first 20% as burn-in, the GTRCAT model as the best-fitted substitution model, the number of substitution types was set to 6, the rate matrix and the base frequencies were set as estimated. Finally, the likelihood and Bayesian tree were visualized in FIGTREE v.1.4.4 (Rambaut, 2021).

TreeMix v.1.13 (Pickrell & Pritchard, 2012) was used to infer the historical relationship and gene flow in a phylogenetic context between the four cryptic species as determined by genetic clustering approaches (DAPC, fastSTRUCTURE). TreeMix estimates an ML species tree and the direction and weight (*w*) of gene flow between taxa using allele frequencies. Using the supermatrix of 5,800 SNPs, we applied the multispecies coalescent model in StarBEAST2 template (Ogilvie *et al.*, 2017) in Beast2 v.2.6.3 (Bouckaert *et al.*, 2014) to concurrently estimate a species tree with divergence times of cryptic species. Although StarBEAST2 is designed for multilocus analyses, it may also be used with single-locus genomic data sets such as RAD-seq, which are usually a good fit for it (Ogilvie *et al.*, 2016). Individuals were classified into cryptic species using a combination of current taxonomy (e.g., Grey-Wilson, 1995; Wang & Gilbert, 2007; Zhang *et al.*, 2016) and additional phylogenomic and clustering results. Multiple individuals were included for each cryptic species to account for intraspecific diversity. A total of four cryptic species and the outgroups were defined. Analyses used a relaxed clock lognormal species tree clock model, analytical population size integration, a GTR substitution model identified by Modeltest-NG, and a calibrated yule tree prior (Heled & Drummond, 2010). The remaining priors were set to their default values. Two independent chains were run for $5\times10^7$ generations each, sampling every 5,000. Tracer v.1.7.2 (Rambaut *et al.*, 2018; 2021) was used to monitor ESS values to make sure that analyses were run for a sufficient length of time (target ESS>200). LogCombiner was used to combine the tree files from both runs, following a burn-in of 10% of trees. TreeAnnotator was then used to construct an MCC tree. For temporal calibration, we followed Zhang *et al.* (2010), who used two fossil information to calibrate the root age of *Stellera* L. The first calibration was constrained between 33.9–23.03 Mya for the split between *Aquilaria* Lam. and other species based on the known *Aquilaria* fossil record (Mai *et al.*, 1978) and estimated origin of *Aquilaria* (32–26 Mya) (Wikström *et al.*, 2001). The second fossil calibration was based on the fossils of *Daphne* L. set between 16.4–5.3 Mya for the split between *Thymelaea* Mill. and *Daphne* (Palmarev, 1989). Therefore, we calibrated the root node of *Stellera* as the split of *S. chamaejasme* and *Wikstroemia* Endl. as 6.5892 Mya (Zhang *et al.*, 2010) using a normal birth rate prior and the normal distribution (mean of 6.5892, sigma of 1.0 and zero offset).

To test whether the genetic divergence between the four cryptic species was significant beyond what would be expected from geographic isolation alone, we compared linear models with geographic distance and each cryptic species as fixed effects as the function of isolation-by-distance (IBD). Mantel tests were used to evaluate the correlation between genetic and geographic distance within each cryptic species. The test of the significance of the relationship between geographic distance and genetic distance among the populations was implemented in the R-package '*vegan*' v2.5-7 (Oksanen *et al.*, 2021) using the mantel function with 9,999 permutations. The uncorrected pairwise genetic distance between all the populations of the cryptic species was obtained from the '*populations*' module of Stacks. In contrast, the geographic distance among the populations of the cryptic species was calculated using the *dist* function in the R-package '*stats*' v.3.6.2

(*https://rdocumentation.org/packages/stats/versions/3.6.2*) as a part of R-programming language (R Core Team, 2021).

**Cryptic species delimitation**

The multispecies coalescent (MSC) model implemented in the program BPP v.4.4.0 (Yang, 2015) uses the reversible-jump Bayesian MCMC (rjMCMC) method of Yang & Rannala (2010) to generate the posterior probabilities of different species delimitation based on support for recent coalescence using a fixed species tree and four putative cryptic species. The likelihood-based implementation of the multispecies coalescent model accounts for the species' phylogeny while accommodating uncertainty due to unknown gene trees (topologies and branch lengths) at individual loci and incomplete lineage sorting. Basic assumptions include no recombination within a locus and free recombination between loci, neutral evolution at a constant rate at each locus under the JC69 model of nucleotide substitution, and no gene flow between species (Yang, 2015). For each run, we randomly pruned the samples to 3 individuals for 34 populations of four different cryptic species and comprised the entire sequence length of 7,208 bp obtained from *Stacks* pipeline with *-r* 0.8, *-p* 4, and *MAF* >0.01. We performed joint species delimitation and species-tree estimation (analysis A11) for 102 individuals (13 RW population, 5 PP population, 7 YW population, and 7 YY population). The analyses used the uniform rooted trees for the species model prior and the species guide tree from the IQ-TREE analysis. Gamma priors were assigned on population size ($\theta$) and divergence time ($\tau$) parameters on 20 different combinations of priors (Table **S8**). Both the $\theta$ and $\tau$ are expressed in units of the expected number of mutations per site. Sites with ambiguities were retained and used in the likelihood calculation (Yang, 2015). The analysis was run for $10^7$ MCMC iterations after a burn-in of 10,000 iterations, sampling every 5 iterations (for 2,000,000 samples). We allowed fine-tune parameters to be automatically adjusted as recommended by the software authors. The convergence of the two runs was analyzed and compared for each combination of priors using rjMCMC algorithm 1 (with parameters $\alpha = 2$ and $m = 1$).

The Bayes Factor Delimitation (BFD*) method (Leaché *et al.*, 2014) is mainly used for comparing alternative species delimitation models in an explicit multispecies coalescent framework using genome-wide SNP data. We calculated marginal likelihood estimates (MLE) of four different alternative scenarios of the species delimitation model obtained from BPP on a supermatrix of unlinked SNPs from 34 populations of *S. chamaejasme*. We then compared the MLE of the null model based on the current taxonomy of the species. The 251 individuals and three closely related taxa (*Daphne longilobata*, *Daphne tangutica*, *Wikstroemia ligustrina*) were used for MLE estimation under different delimitation models. For each model, individuals were assigned to different cryptic species. Of the alternative scenarios ("Lump, Split, Re-assign") stated by Grummer *et al*. (2014), we are more focused on splitting the current taxonomy model into four species models. MLE was obtained using the implementation of BFD* in the SNAPP (Bryant *et al*., 2012) plug-in for BEAST v2.6.3 (Bouckaert *et al*., 2014). We performed a path sampling of 4 runs (MCMC length 10,000, pre burn-in 1,000), following (Bryant *et al*., 2012). The convergence of 4 runs was triplicated for ESS>200 and MLE estimates using a Path sampler analyzer (*alpha*=0.3, *nr of steps*=12, *burn-in percentage*=20, *cross*=0, *repeats*=2000). The strength of support for the compared model was evaluated from the Bayes factor scale, *2ln* (BF), using the framework of Kass & Raftery (1995). The BF scale is stated as: 0 < *2ln* (BF) < 2 is not worth more than a bare mention, 2 < *2ln* (BF) < 6 means positive evidence, 6 < *2ln*(BF) < 10 represents strong support, and *2ln*(BF) > 10 represents decisive support.

**Unsupervised machine learning**

The unsupervised machine learning (UML) approaches Random Forest (RF), and t-Distributed stochastic neighbor embedding (t-SNE) were run and visualized using an R script developed by Derkarabetian *et al*. (2019), more specifically based on R-package '*randomForest*' v.4.6.-14 (Liaw & Wiener, 2002), and '*Rtsne*' v.0.15 (Krijthe, 2021) respectively. The data were represented as scaled principal components (*N*=27 axes) generated in R-package '*adgenet*'. RF was performed with *ntree*=5,000 and *mtry* left as default for classification analysis. The resulting proximity matrix was then used in classic multidimensional scaling (cMDS) using the *MDSplot* function in R-package '*randomForest*' and isotonic MDS (isoMDS) using the *isoMDS* function in R-package '*MASS*' v.7.3-53.1 (Venables & Ripley, 2002). t-SNE was executed with parameters maximum iterations (*max_iter*=10,000), *perplexity*=5, initial dimensions (*initial_dims*=5), and number of dimensions for the resulting embedding (*K*=4). More specifically, for large datasets, we performed t-SNE using the results of the initial PCA as an input (Pedregosa *et al*., 2011).

Furthermore, the two common distance-based clustering analyses were conducted on UML outputs to derive optimal *K* for RF and t-SNE analyses. The analyses were performed to confirm that the UML cluster assignments are equivalent to DAPC and fastSTRUCTURE analyses. We used *K*-means (MacQueen, 1967) and Partitioning Around Medoids (*PAM*) (Kaufman & Rousseeuw, 1990) clustering methods to find the optimal number of clusters (*K*) based on the within-cluster-sum of squared errors (*WSS*) index. The analyses were executed by R-package '*NbClust*' v.3.0 (Charrad *et al*., 2014) for *K*-means clustering and R-package '*cluster*' v.2.10 (Maechler *et al*., 2021) for *PAM* clustering, subsequently followed by R-package '*factoextra*' v.1.0.7 (Kassambara & Mundt, 2020) for visualization.

**Morphological analysis**

For morphological analysis, the 950 specimens were examined from our original collections and the herbaria (KUN, PE, IBSC, and KATH). Twenty-nine (16 quantitative and 13 discrete) morphological traits were measured from the representative cryptic species of *S. chamaejasme*. Preliminarily, ridgeplot and corplot heatmap were visualized for the morphological differentiation and correlation between all the morphological traits, respectively. Furthermore, a multivariate analysis of variance (MANOVA) was used to analyze the variance among the non-normal data of presumed four cryptic species. Post-hoc pair-wise comparison of MANOVA was based on Tukey's multiple comparison test. *P*-values of less than 0.05 were considered statistically significant. These statistical analyses were performed using R-package '*car*' v.3.0-10 (Fox & Weisberg, 2019) and '*MASS*' (Venables & Ripley, 2002).

Additionally, morphological character states were analyzed using principal component analysis (PCA) and Linear discriminant analysis (LDA) using the R-package '*factoextra*' and '*MASS'*, respectively. A PCA was used to detect morphological variation patterns and determine whether the data could be partitioned into taxonomically meaningful groups solely based on morphological character states. Furthermore, significant morphological traits were visualized using traitgram for the mean trait value of the sampled individuals in each population using R-package '*phytools*' v.0.7-70 (Revell, 2012) using the dated MCMC trees.

**Ensemble Species Distribution Modelling (eSDM)**

We constructed the range maps for each putative cryptic species based on the previous clustering analysis. Species occurrence data were collated from herbarium records at the National Herbarium and Plant Laboratories (KATH, Nepal) and Kunming Institute of Botany, CAS (KUN, China). In addition to field-gathered occurrence points, online databases of Chinese National Herbarium (PE; *http://pe.ibcas.ac.cn/en/*), Chinese Virtual Herbarium (CVH; *http://www.cvh.ac.cn/*), Global Biodiversity Information Facility (GBIF; *https://www.gbif.org/*), the Royal Botanical Garden at Edinburgh (RBGE, United Kingdom; *http://data.rbge.org.uk/search/herbarium/*) and the Herbarium at the University of Tokyo (TI, Japan; *http://umdb.um.u-tokyo.ac.jp/DShokubu/*) were utilized. The random pseudo-absence data were selected within a 25–50 km radius from the point of occurrence following Barbet-Massin *et al*. (2012). These points were spatially rarefied at the minimal filter distance of 10 km, which is more appropriate for the mountainous region with high spatial heterogeneity (Boria *et al*., 2014; Rana *et al*., 2020) The use of rarefication is more prevalent when developing the distribution model to prevent spatial clustering and model inflation (Boria *et al*., 2014; Rana *et al*., 2020; 2021). Out of 681 records collected, only 363 occurrence points have well-discriminated cryptic species. Finally, 307/363 occurrence points (of total) [RW-176/200, PP-9/9, YW-40/54 and YY-82/100] were used for building the models.

The potential distribution of four cryptic species was modeled for the current climatic scenario (c. 1990–2000) and projected to the paleoclimatic (Last glacial maximum, LGM c. 22kya) scenario. The climatic projection was based on the multi-model median (MMM) ensemble of 3 General circulation models (GCMs) for the 19 climatic variables provided by Paleoclimate modeling inter-comparisons project phase 5 (Braconnot *et al*., 2007). The climatic space under the current conditions was analyzed with bioclimatic (*http://www.worldclim.org/version2*), geo-climatic (*http://www.worldclim.org/version2*; *http://nelson.wisc.edu/sage*; *http://www.cgiar-csi.org*), habitat heterogeneity (Tuanmu & Jetz, 2015; *http://www.earthenv.org/texture*), growing days (*https://chelsa-climate.org/*), ultra-violet radiation (Beckmann *et al*., 2014; *https://www.ufz.de/gluv/*) and consensus land-cover (Tuanmu & Jetz, 2014; *http://www.earthenv.org/landcover*) variables to further discriminate the suitability range of four cryptic species along with their relative contributions (Table **S14**). All the predictive variables were used with a spatial resolution of c. 2.5 arc-min.

The variance inflation factor (VIF; Fox & Weisberg, 2019), calculated using R-package '*car*' with elevation as a response variable was used to test for and remove highly correlated and/or redundant variables. A VIF >10 indicates strong collinearity (Quinn & Keough, 2002), and variables with high VIF adversely affect the modeling results. Therefore, several test runs were performed on the extent of occurrence points for climatic and other environmental variables until a set of predictors with VIF values lower than 10 was retained (Table **S15**). Besides, the global consensus land-cover variables with occurrence points greater than the 50% threshold (100% for pure pink morph) out of total occurrence points were selected for the cryptic species (Table **S16**). The resulting 6–9 bio-climatic variables (9/7/6/7 for RW/PP/YW/YY morph), 7–13 geo-climatic variables (13/7/12/12 for RW/PP/YW/YY morph), 4–7 habitat heterogeneity (7/4/5/5 for RW/PP/YW/YY morph), 3–5 growing days (4/3/4/5 for RW/PP/YW/YY morph), 2 ultra-violet radiations (2 for each morph) and 3–5 consensus land-cover (3/4/4/5 for RW/PP/YW/YY morph) were used as a subset of predictive variables (Tables **S14**, **S16**). Statistically, all the predictive variables were analyzed to evaluate the relative contributions of the variables and discriminate the four cryptic species using PCA and LDA using R-package '*factoextra*' and '*MASS*', respectively.

Species distribution modeling was carried out using R-package '*Biomod2*' v3.4.12 (Thuiller *et al.*, 2020) as an ensemble of species distribution models (Guisan & Zimmermann, 2000). Moreover, the modeling process was projected to the past (LGM) scenario in response to current climatic conditions using the multi-model median (MMM) of GCMs for the four cryptic species. The default parameters were used for each *biomod2* sub-model. For proper evaluation, models were 4-fold cross-validated using 80% of the data for training the models. In comparison, the remaining 20% were used for evaluating the predictive power of each model (Araujo, 2005) using TSS (True Skill Statistics), Cohen's kappa, and AUC (Area Under Curve-Receiver Operating characteristics) statistics. TSS and Cohen's kappa are the threshold-dependent measures of model accuracy and are independent of prevalence–the ratio of presence to pseudo-absence data in the presence-absence predictions (Allouche, 2006). It deals with sensitivity and specificity, with values ranging from − 1 to + 1, where + 1 indicates perfect agreement, and scores ranging from 0.6 to 0.9 specify fair to good model performance (Allouche, 2006). At the same time, AUC is a threshold-independent model evaluation indicator and independent of prevalence (Phillips & Anderson, 2006). It assesses the discrimination capacity of the models (the probability that a randomly selected presence will have a higher model output than a randomly selected absence). AUC values below 0.6 were considered weak, 0.6–0.9 moderate, and > 0.9 good (Phillips & Anderson, 2006). An ensemble of the ten algorithms was generated using the model with TSS ≥ 0.80, whereas Cohen's kappa and AUC are considered as the indices for the model accuracy (Table **S17**). The sampling procedure was replicated 5 times. The consensus model was then projected onto a paleo-climatic scenario (LGM). The contribution of each model to the final ensemble model was proportional to their goodness-of-fit statistics. So obtained consensus ensemble model was converted to a binary model (presence/absence), applying thresholds that allow a maximum of 50% probability of suitable habitat (Forester *et al.*, 2013; Rana *et al.*, 2020; 2021) that suits the present distribution of the focal species.

**Niche equivalency test**

Additionally, the hypothesis testing as a function of the niche equivalency test among the cryptic species was performed using *ecospat* test in R-package '*ENMTools*' v.1.0.3 (Warren

& Dinnage, 2021). The extent of occurrence points selected for each cryptic species lies within 50 km radius of each genomics sampled population. The R version of ENMTools provides a simplified user interface for ecospat hypothesis tests using *enmtools.species* objects conducting principal component analysis for multiple predictors layers to reduce the predictors to a two-dimensional environment space (Warren *et al*., 2021). The climatic niches were also quantified and validated using Schoener's '*D*' and Hellinger's-based '*I*' implemented in *enmtools.ecospat.id* function of identity test and *enmtools.ecospat.bg* function of background test (symmetric and asymmetric) (Warren *et al*., 2010; 2021). Schoener's *D* measures the suitable range based on the probability of occupied grid cells. Hellinger's-based *I* work similarly to Schoener's *D* but without its assumption (Warren & Seifert, 2011). The pairwise similarity values of *D* and *I* indices ranged between 0 (complete divergence/no overlap) and 1 (high similarity/complete overlap); the higher the score indicates more niche overlap.

**References**

**Allouche O, Tsoar A, Kadmon R. 2006.** Assessing the accuracy of species distribution models: prevalence, kappa and the true skill statistic (TSS). *Journal of Applied Ecology* 43: 1223–1232.

**Andrews S. 2010.** FastQC: A quality control tool for high throughput sequence data. Babraham Bioinformatics. Version 0.11.4. URL http://www.bioinformatics.babraham.ac.uk/projects/fastqc. [accessed 1 November 2020]

**Araujo MB, Pearson RG, Thuiller W, Erhard M. 2005.** Validation of species–climate impact models under climate change. *Global Change Biology* 11: 1504–1513.

**Barbet-Massin M, Jiguet F, Albert CH, Thuiller W. 2012.** Selecting pseudo-absences for species distribution models: how, where and how many? *Methods in Ecology and Evolution* 3: 327–338.

**Beckmann M, Václavík T, Manceur AM, Šprtová L, von Wehrden H, Welk E, Cord AF. 2014.** glUV: a global UV-B radiation data set for macroecological studies. *Methods in Ecology and Evolution* 5: 372–383.

**Boria RA, Olson LE, Goodman SM, Anderson RP. 2014.** Spatial filtering to reduce sampling bias can improve the performance of ecological niche models. *Ecological Modelling* 275: 73–77.

**Braconnot P, Otto-Bliesner B, Harrison S, Joussaume S, Peterchmitt JY, Abe-Ouchi A, Crucifix M, Driesschaert E, Fichefet T, Hewitt CD et al. 2007.** Results of PMIP2 coupled simulations of the Mid-Holocene and Last Glacial Maximum–Part1: experiments and large-scale features. *Climate of the Past* 3: 261–277.

**Bryant D, Bouckaert R, Felsenstein J, Rosenberg NA, RoyChoudhury A. 2012.** Inferring species trees directly from biallelic genetic markers: bypassing gene trees in a full coalescent analysis. *Molecular Biology and Evolution* 29: 1917–1932.

**Catchen J, Hohenlohe PA, Bassham S, Amores A, Cresko WA. 2013.** Stacks: an analysis tool set for population genomics. *Molecular Ecology* 22: 3124–3140.

**Charrad M, Ghazzali N, Boiteau V, Niknafs A. 2014.** NbClust: An R-package for determining the relevant number of clusters in a data set. *Journal of Statistical Software* 61: 1–36.

**Darriba D, Posada D, Kozlov AM, Stamatakis A, Morel B, Flouri T. 2020.** ModelTest-NG: A new and scalable tool for the selection of DNA and protein evolutionary models. *Molecular Biology and Evolution* 37: 291–294.

**Derkarabetian S, Castillo S, Koo PK, Ovchinnikov S, Hedin M. 2019.** A demonstration of unsupervised machine learning in species delimitation. *Molecular Phylogenetics and Evolution* 139: 106562.

**Pedregosa F, Varoquaux G, Gramfort A, Michel V, Thirion B, Grisel O, Blondel M, Prettenhofer P, Weiss R, Dubourg V et al. 2011.** Scikit-learn: Machine learning in Python. *Journal of Machine Learning Research* 12: 2825–2830.

**Flor M. 2021.** chorddiag: Interactive chord diagrams. R-package version 0.1.3. URL https://github.com/mattflor/chorddiag/. [accessed 3 January 2021]

**Forester BR, DeChaine EG, Bunn AG. 2013.** Integrating ensemble species distribution modelling and statistical phylogeography to inform projections of climate change impacts on species distributions. *Diversity and Distributions* 19: 1480–1495.

**Fox J, Weisberg S. 2019.** *An R Companion to Applied Regression, Third Edition*. California: Sage, Thousand Oaks.

**Grey-Wilson C. 1995.** *Stellera chamaejasme*: an overview. *The New Plantsman* 2: 43–49.

**Grummer JA, Bryson RW, Reeder TW. 2014.** Species delimitation using Bayes factors: Simulations and application to the *Sceloporus scalaris* species group (Squamata: Phrynosomatidae). *Systematic Biology* 63: 119–133.

**Guisan A, Zimmermann NE. 2000.** Predictive habitat distribution models in ecology. *Ecological Modelling* 135: 147–186.

**Heled J, Drummond AJ. 2010.** Bayesian inference of species trees from multilocus data. *Molecular Biology and Evolution* 27: 570–580.

**Hoang DT, Chernomor O, von Haeseler A, Minh BQ, Vinh LS. 2018.** UFBoot2: improving the ultrafast bootstrap approximation. *Molecular Biology and Evolution* 35: 518–522.

**Jakobsson M, Rosenberg NA. 2007.** CLUMPP: A cluster matching and permutation program for dealing with label switching and multimodality in analysis of population structure. *Bioinformatics* 23: 1801–1806.

**Jombart T, Ahmed I. 2011.** adegenet 1.3-1: new tools for the analysis of genome-wide SNP data. *Bioinformatics* 27: 3070–3071.

**Jombart T, Devillard S, Balloux F. 2010.** Discriminant analysis of principal components: A new method for the analysis of genetically structured populations. *BMC Genetics* 11: 94.

**Jombart T. 2008.** adegenet: a R-package for the multivariate analysis of genetic markers. *Bioinformatics* 24: 1403–1405.

**Kalyaanamoorthy S, Minh BQ, Wong TKF, von Haeseler A, Jermiin LS. 2017.** ModelFinder: Fast model selection for accurate phylogenetic estimates. *Nature Methods* 14: 587–589.

**Kass RE, Raftery AE. 1995.** Bayes factors. *Journal of the American Statistical Association* 90: 773–795.

**Kassambara A, Mundt F. 2020.** factoextra: Extract and visualize the results of multivariate data analyses. R-package version 1.0.7. URL https://cran.r-project.org/web/packages/factoextra/index.html. [accessed 3 January 2021]

**Kaufman L, Rousseeuw PJ. 1990.** Partitioning around medoids (Program PAM). In: Kaufman L, Rousseeuw PJ, eds. *Finding groups in data: An introduction to cluster analysis*. Hoboken: John Wiley & Sons, Inc, 68–125.

**Krijthe JH. 2015.** Rtsne: T-distributed stochastic neighbor embedding using a Barnes-Hut implementation. R-package version 0.15. URL https://github.com/jkrijthe/Rtsne. [accessed 3 January 2021]

**Lawson DJ, Hellenthal G, Myers S, Falush D. 2012.** Inference of population structure using dense haplotype data. *PLoS Genetics* 8: e1002453.

**Leaché AD, Fujita MK, Minin VN, Bouckaert RR. 2014.** Species delimitation using genome-wide SNP data. *Systematic Biology* 63: 534–542.

**Liaw A, Wiener M. 2002.** Classification and regression by randomForest. *R News* 2: 18–22.

**MacQueen J. 1967.** Some methods for classification and analysis of multivariate observations. In: Cam LML, Neyman J, eds. *Proceedings of the fifth Berkeley symposium on mathematical statistics and probability, vol 1: Statistics*. Berkeley, California: University of California Press, 281–297.

**Maechler M, Rousseeuw P, Struyf A, Hubert M, Hornik K. 2019.** cluster: Cluster analysis basics and extensions. R-package version 2.1.0. URL https://cran.r-project.org/web/packages/cluster/index.html. [accessed 3 January 2021]

**Mai DH, Walther H. 1978.** Die Floren des Haselbacher Serie im Weisselster-Becken (Bezirk Leipzig, DDR). *Abhandlungen des Staatlichen Museums für Mineralogie und Geologie zu Dresden 28*: 1–200.

**Malinsky M, Trucchi E, Lawson DJ, Falush D. 2018.** RADpainter and fineRADstructure: Population inference from RADseq data. *Molecular Biology and Evolution* 35: 1284–1290.

**Minh BQ, Nguyen MAT, von Haeseler A. 2013.** Ultrafast approximation for phylogenetic bootstrap. *Molecular Biology and Evolution* 30: 1188–1195.

**Nguyen LT, Schmidt HA, von Haeseler A, Minh BQ. 2015.** IQ-TREE: a fast and effective stochastic algorithm for estimating maximum likelihood phylogenies. *Molecular Biology and Evolution* 33: 268–274.

**Ogilvie HA, Bouckaert RR, Drummond AJ. 2017.** Starbeast2 brings faster species tree inference and accurate estimates of substitution rates. *Molecular Biology and Evolution* 34: 2101–2114.

**Ogilvie HA, Heled J, Xie D, Drummond AJ. 2016.** Computational performance and statistical accuracy of *BEAST and comparisons with other methods. *Systematic Biology* 65: 381–396.

**Oksanen J, Blanchet FG, Friendly M, Kindt R, Legendre P, McGlinn D, Minchin PR, O'Hara RB, Simpson GL, Solymos P et al. 2020.** vegan: Community ecology package. R-package version 2.5-7. URL https://cran.r-project.org/web/packages/vegan/index.html. [accessed 3 January 2021]

**Palmarev E. 1989.** Paleobotanical evidence of the Tertiary history and origin of the Mediterranean sclerophyll dendroflora. *Plant Systematics and Evolution* 162: 93–107.

**Phillips SJ, Anderson RP, Schapire RE. 2006.** Maximum entropy modeling of species geographic distributions. *Ecological Modelling* 190: 231–259.

**Pickrell JK, Pritchard JK. 2012.** Inference of population splits and mixtures from genome-wide allele frequency data. *PLoS Genetics* 8: e1002967.

**Qiu J. 2008.** The Third Pole. *Nature* 454: 393–396.

**Quinn GP, Keough MJ. 2002.** *Experimental Design and Data Analysis for Biologists*. Cambridge: Cambridge University Press.

**R Core Team. 2021.** R: A language and environment for statistical computing. Vienna, Austria: R Foundation for Statistical Computing. URL https://www.R-project.org/. [accessed 2 April 2020]

**R. Bouckaert, Heled J, Kühnert D, Vaughan T, Wu CH, Xie D, Suchard MA, Rambaut A, Drummond AJ. 2014.** BEAST2: a software platform for Bayesian evolutionary analysis. *PLoS Computational Biology* 10: e1003537.

**Raj A, Stephens M, Pritchard JK. 2014.** fastSTRUCTURE: Variational inference of population structure in large SNP data sets. *Genetics* 197: 573–589.

**Rambaut A, Drummond AJ, Xie D, Baele G, Suchard MA. 2018.** Posterior summarization in Bayesian phylogenetics using Tracer 1.7. *Systematic Biology* 67: 901–904.

**Rambaut A, Suchard MA, Xie D, Drummon AJ. 2014.** Tracer. Version 1.6. URL http://tree.bio.ed.ac.uk/software/tracer/. [accessed 12 March 2021]

**Rambaut A. 2021.** FigTree. Version 1.4.4. URL https://github.com/rambaut/figtree. [accessed 12 March 2021]

**Rana HK, Luo D, Rana SK, Sun H. 2020.** Geological and climatic factors affect the population genetic connectivity in *Mirabilis himalaica* (Nyctaginaceae): Insight from phylogeography and dispersal corridors in the Himalaya-Hengduan Biodiversity Hotspot. *Frontiers in Plant Science* 10: 1721.

**Rana SK, Rana HK, Luo D, Sun H. 2021.** Estimating climate-induced 'Nowhere to go' range shifts of the Himalayan *Incarvillea* Juss. using multi-model median ensemble species distribution models. *Ecological Indicator* 121: 107127.

**Revell LJ. 2012.** phytools: An R package for phylogenetic comparative biology (and other things). *Methods in Ecology and Evolution* 3: 217–223.

**Rochette N, Catchen J. 2017.** Deriving genotypes from RAD-seq short-read data using Stacks. *Nature Protocols* 12: 2640–2659.

**Ronquist F, Teslenko M, van der Mark P, Ayres DL, Darling A, Höhna S, Larget B, Liu L, Suchard MA, Huelsenbeck JP. 2012.** MrBayes 3.2: Efficient Bayesian phylogenetic inference and model choice across a large model space. *Systematic Biology* 61: 539–542.

**Rosenberg NA. 2004.** DISTRUCT: a program for the graphical display of population structure. *Molecular Ecology Notes* 4: 137–138.

**Thuiller W, Georges D, Engler R, Breiner F. 2020.** Biomod2: Ensemble platform for species distribution modeling. R-package version 3.4.12. URL https://cran.r-project.org/web/packages/biomod2/index.html. [accessed 3 November 2020]

**Tuanmu MN, Jetz W. 2014.** A global 1-km consensus land-cover product for biodiversity and ecosystem modeling. *Global Ecology and Biogeography* 23: 1031–1045.

**Tuanmu MN, Jetz W. 2015.** A global, remote sensing-based characterization of terrestrial habitat heterogeneity for biodiversity and ecosystem modelling. *Global Ecology and Biogeography* 24: 1329–1339.

**Venables WN, Ripley BD. 2002.** *Modern Applied Statistics with S (Statistics and Computing), Fourth Edition*. Springer.

**Wang YZ, Gilbert MG. 2007.** *Stellera* Linnaeus. In: Wu CY, Raven PH, Hong DY, eds. *Flora of China, vol. 13.* Beijing: Science Press, 250.

**Warren DL, Dinnage R. 2021.** ENMTools: Analysis of niche evolution using niche and distribution models. R-package version 1.0.3. URL https://cran.r-project.org/web/packages/ENMTools/index.html. [accessed 2 June 2021]

**Warren DL, Glor RE, Turelli M. 2010.** ENMTools: a toolbox for comparative studies of environmental niche models. *Ecography* 33: 607–611.

**Warren DL, Matzke NJ, Cardillo M, Baumgartner JB, Beaumont LJ, Turelli M, Glor RE, Huron NA, Simões M, Iglesias TL et al. 2021.** ENMTools 1.0: an R package for comparative ecological biogeography. *Ecography* 44: 504–511.

**Warren DL, Seifert SN. 2011.** Ecological niche modeling in Maxent: the importance of model complexity and the performance of model selection criteria. *Ecological Applications* 21: 335–342.

**Wikström N, Savolainen V, Chase MW. 2001.** Evolution of the angiosperms: calibrating the family tree. *Proceedings of the Royal Society B: Biological Sciences* 268: 2211–2220.

**Yang Z, Rannala B. 2010.** Bayesian species delimitation using multilocus sequence data. *Proceedings of the National Academy of Sciences of the United States of America* 107: 9264–9269.

**Yang Z. 2015.** The BPP program for species tree estimation and species delimitation. *Current Zoology* 61: 854–865.

**Zhang D, Ye J, Sun H. 2016.** Quantitative approaches to identify floristic units and centres of species endemism in the Qinghai-Tibetan Plateau, south-western China. *Journal of Biogeography* 43: 2465–2476.
