## Supporting Information_Table S4 for "Pleistocene glaciation advances the younger temporal dimension of species diversification in a major biodiversity hotspot"

**Table S4.** The paired population-level estimation of genetic diversity indices; $\Phi_{ST}$means (lower left diagonal) and $F_{ST}$'means (upper right diagonal) f

| Populations | Chen0018 | Chen0019 | Chen0021 | Chen0022 | LZY004005 | Yue0719 | Yue0720 | ZBFC295 | ZBFC425 | Deng9733 | Deng9745 |
|---|---|---|---|---|---|---|---|---|---|---|---|
| Chen0018 | - | -0.057 | 0.121 | 0.1 | 0.099 | -0.027 | 0.096 | 0.09 | 0.054 | -0.014 | -0.015 |
| Chen0019 | -0.03 | - | 0.139 | 0.111 | 0.182 | -0.042 | 0.18 | 0.168 | 0.123 | -0.002 | -0.007 |
| Chen0021 | 0.154 | 0.139 | - | 0.149 | 0.221 | 0.087 | 0.169 | 0.178 | 0.192 | 0.131 | 0.126 |
| Chen0022 | 0.163 | 0.082 | 0.168 | - | 0.005 | 0.104 | 0.04 | 0.044 | 0.044 | 0.119 | 0.095 |
| LZY004005 | 0.171 | 0.188 | 0.251 | 0.017 | - | 0.152 | 0.038 | 0.026 | 0.033 | 0.16 | 0.123 |
| Yue0719 | 0.019 | -0.025 | 0.113 | 0.098 | 0.167 | - | 0.12 | 0.124 | 0.112 | -0.001 | 0.003 |
| Yue0720 | 0.195 | 0.182 | 0.208 | 0.069 | 0.071 | 0.131 | - | 0.002 | 0.015 | 0.152 | 0.104 |
| ZBFC295 | 0.159 | 0.186 | 0.2 | 0.05 | 0.056 | 0.141 | 0.013 | - | 0.005 | 0.177 | 0.104 |
| ZBFC425 | 0.128 | 0.105 | 0.213 | 0.065 | 0.054 | 0.12 | 0.032 | 0.017 | - | 0.117 | 0.086 |
| Deng9733 | 0.051 | -0.007 | 0.158 | 0.131 | 0.178 | 0.008 | 0.174 | 0.195 | 0.125 | - | 0.003 |
| Deng9745 | 0.051 | -0.01 | 0.159 | 0.114 | 0.151 | 0.019 | 0.129 | 0.12 | 0.103 | 0.015 | - |
| Deng9772 | 0.094 | 0.023 | 0.225 | 0.136 | 0.229 | 0.047 | 0.246 | 0.302 | 0.173 | 0.004 | 0.035 |
| Deng9783 | 0.197 | 0.37 | 0.329 | 0.208 | 0.254 | 0.306 | 0.224 | 0.257 | 0.211 | 0.328 | 0.321 |
| Deng9810 | 0.104 | 0.133 | 0.191 | 0.037 | 0.032 | 0.085 | 0.09 | 0.095 | 0.068 | 0.122 | 0.107 |
| Deng9813 | 0.147 | 0.121 | 0.211 | 0.083 | 0.085 | 0.13 | 0.103 | 0.107 | 0.084 | 0.154 | 0.121 |
| Deng9525 | 0.198 | 0.339 | 0.338 | 0.065 | 0.026 | 0.251 | 0.101 | 0.119 | 0.075 | 0.194 | 0.196 |
| Deng9528 | 0.064 | 0.108 | 0.123 | 0.054 | 0.037 | 0.124 | 0.16 | 0.144 | 0.084 | 0.074 | 0.09 |
| Deng9533 | 0.077 | 0.098 | 0.191 | 0.079 | 0.055 | 0.117 | 0.099 | 0.096 | 0.081 | 0.112 | 0.121 |
| Chen0016 | 0.287 | 0.437 | 0.464 | 0.337 | 0.476 | 0.381 | 0.476 | 0.455 | 0.435 | 0.4 | 0.389 |
| ZYZhou | 0.482 | 0.639 | 0.602 | 0.51 | 0.594 | 0.593 | 0.579 | 0.606 | 0.578 | 0.586 | 0.551 |
| Deng9524 | 0.36 | 0.521 | 0.423 | 0.379 | 0.4 | 0.464 | 0.452 | 0.424 | 0.492 | 0.492 | 0.434 |
| Deng9535 | 0.431 | 0.552 | 0.494 | 0.339 | 0.366 | 0.46 | 0.409 | 0.423 | 0.427 | 0.48 | 0.421 |
| Deng9715 | 0.481 | 0.645 | 0.607 | 0.469 | 0.625 | 0.603 | 0.587 | 0.614 | 0.638 | 0.599 | 0.549 |
| Deng9703 | 0.424 | 0.608 | 0.568 | 0.453 | 0.56 | 0.541 | 0.558 | 0.573 | 0.582 | 0.555 | 0.496 |
| Deng9847 | 0.567 | 0.721 | 0.639 | 0.497 | 0.641 | 0.664 | 0.613 | 0.657 | 0.69 | 0.673 | 0.629 |
| Chen0009 | 0.577 | 0.744 | 0.727 | 0.681 | 0.718 | 0.724 | 0.727 | 0.737 | 0.725 | 0.736 | 0.723 |
| Chen0015 | 0.604 | 0.709 | 0.7 | 0.623 | 0.683 | 0.691 | 0.666 | 0.666 | 0.681 | 0.703 | 0.651 |

| | | | | | | | | | | | |
|---|---|---|---|---|---|---|---|---|---|---|---|
| **Niuma003** | 0.403 | 0.616 | 0.5 | 0.491 | 0.502 | 0.538 | 0.509 | 0.522 | 0.584 | 0.545 | 0.481 |
| **Deng8939** | 0.568 | 0.738 | 0.73 | 0.681 | 0.729 | 0.74 | 0.732 | 0.737 | 0.733 | 0.748 | 0.735 |
| **Deng8999** | 0.65 | 0.808 | 0.779 | 0.731 | 0.787 | 0.797 | 0.783 | 0.799 | 0.784 | 0.802 | 0.79 |
| **Deng9040** | 0.603 | 0.777 | 0.771 | 0.697 | 0.751 | 0.758 | 0.754 | 0.762 | 0.762 | 0.753 | 0.759 |
| **Deng9127** | 0.54 | 0.677 | 0.657 | 0.584 | 0.595 | 0.632 | 0.637 | 0.63 | 0.639 | 0.649 | 0.608 |
| **Deng9569** | 0.62 | 0.786 | 0.77 | 0.719 | 0.762 | 0.756 | 0.767 | 0.769 | 0.766 | 0.777 | 0.767 |
| **Deng9601** | 0.536 | 0.701 | 0.691 | 0.629 | 0.648 | 0.67 | 0.652 | 0.704 | 0.689 | 0.701 | 0.653 |

ΦST means, AMOVA-based measure of FST (lower left diagonal); Fst' means, haplotype measure of FST that is scaled to the theoretical maximum [illegible] or 34 populations estimated using populations platform in Stacks.

| Deng9772 | Deng9783 | Deng9810 | Deng9813 | Deng9525 | Deng9528 | Deng9533 | Chen0016 | ZYZhou | Deng9524 | Deng9535 | Deng9715 |
|---|---|---|---|---|---|---|---|---|---|---|---|
| 0.004 | 0.136 | 0.071 | 0.1 | 0.13 | 0.01 | 0.086 | 0.239 | 0.416 | 0.339 | 0.394 | 0.431 |
| 0.018 | 0.373 | 0.115 | 0.135 | 0.321 | 0.079 | 0.105 | 0.435 | 0.638 | 0.53 | 0.523 | 0.646 |
| 0.192 | 0.306 | 0.172 | 0.192 | 0.308 | 0.11 | 0.174 | 0.445 | 0.58 | 0.409 | 0.462 | 0.591 |
| 0.125 | 0.192 | 0.017 | 0.055 | 0.044 | 0.043 | 0.041 | 0.326 | 0.49 | 0.352 | 0.313 | 0.46 |
| 0.21 | 0.233 | 0.005 | 0.057 | 0.014 | 0.01 | 0.027 | 0.463 | 0.573 | 0.379 | 0.349 | 0.611 |
| 0.02 | 0.297 | 0.057 | 0.128 | 0.214 | 0.056 | 0.116 | 0.374 | 0.579 | 0.448 | 0.423 | 0.594 |
| 0.236 | 0.212 | 0.048 | 0.067 | 0.072 | 0.131 | 0.048 | 0.464 | 0.565 | 0.418 | 0.377 | 0.579 |
| 0.278 | 0.247 | 0.061 | 0.075 | 0.066 | 0.093 | 0.073 | 0.445 | 0.59 | 0.404 | 0.384 | 0.606 |
| 0.163 | 0.189 | 0.021 | 0.054 | 0.05 | 0.041 | 0.03 | 0.428 | 0.568 | 0.46 | 0.399 | 0.631 |
| -0.013 | 0.319 | 0.093 | 0.138 | 0.181 | 0.034 | 0.086 | 0.4 | 0.579 | 0.47 | 0.456 | 0.593 |
| 0.007 | 0.308 | 0.077 | 0.096 | 0.175 | 0.066 | 0.076 | 0.383 | 0.535 | 0.404 | 0.395 | 0.539 |
| - | 0.386 | 0.113 | 0.149 | 0.311 | 0.088 | 0.127 | 0.471 | 0.605 | 0.499 | 0.515 | 0.661 |
| 0.397 | - | 0.091 | 0.096 | 0.083 | 0.074 | 0.127 | 0.488 | 0.579 | 0.415 | 0.36 | 0.647 |
| 0.161 | 0.104 | - | -0.013 | 0 | -0.032 | 0.034 | 0.401 | 0.443 | 0.239 | 0.214 | 0.468 |
| 0.171 | 0.12 | -0.012 | - | 0 | -0.005 | 0.044 | 0.384 | 0.39 | 0.226 | 0.249 | 0.403 |
| 0.322 | 0.116 | 0.025 | 0.023 | - | -0.089 | 0.001 | 0.426 | 0.451 | 0.218 | 0.205 | 0.439 |
| 0.122 | 0.116 | -0.012 | 0 | -0.072 | - | -0.025 | 0.296 | 0.325 | 0.16 | 0.118 | 0.303 |
| 0.155 | 0.158 | 0.041 | 0.059 | 0.02 | -0.033 | - | 0.34 | 0.41 | 0.301 | 0.341 | 0.399 |
| 0.479 | 0.497 | 0.425 | 0.402 | 0.438 | 0.316 | 0.354 | - | 0.233 | 0.1 | 0.136 | 0.221 |
| 0.611 | 0.597 | 0.47 | 0.41 | 0.461 | 0.346 | 0.439 | 0.242 | - | 0.011 | 0.032 | -0.026 |
| 0.519 | 0.443 | 0.257 | 0.239 | 0.228 | 0.138 | 0.306 | 0.141 | 0.072 | - | -0.035 | 0.023 |
| 0.535 | 0.392 | 0.24 | 0.273 | 0.219 | 0.156 | 0.349 | 0.169 | 0.067 | -0.018 | - | 0.025 |
| 0.668 | 0.656 | 0.489 | 0.414 | 0.452 | 0.317 | 0.416 | 0.227 | -0.02 | 0.055 | 0.064 | - |
| 0.629 | 0.584 | 0.374 | 0.355 | 0.38 | 0.224 | 0.392 | 0.193 | -0.014 | 0.023 | 0.031 | -0.034 |
| 0.727 | 0.709 | 0.521 | 0.41 | 0.442 | 0.35 | 0.446 | 0.251 | 0.051 | 0.05 | 0.058 | 0.036 |
| 0.774 | 0.756 | 0.622 | 0.576 | 0.514 | 0.443 | 0.55 | 0.637 | 0.465 | 0.326 | 0.219 | 0.58 |
| 0.728 | 0.72 | 0.575 | 0.549 | 0.506 | 0.374 | 0.589 | 0.52 | 0.433 | 0.279 | 0.199 | 0.421 |

| | | | | | | | | | | | |
|---|---|---|---|---|---|---|---|---|---|---|---|
| 0.584 | 0.572 | 0.391 | 0.331 | 0.355 | 0.188 | 0.359 | 0.218 | 0.174 | 0.008 | 0.075 | 0.144 |
| 0.76 | 0.765 | 0.632 | 0.586 | 0.544 | 0.487 | 0.547 | 0.59 | 0.463 | 0.334 | 0.203 | 0.552 |
| 0.815 | 0.832 | 0.677 | 0.648 | 0.594 | 0.493 | 0.607 | 0.628 | 0.493 | 0.339 | 0.241 | 0.633 |
| 0.786 | 0.795 | 0.656 | 0.578 | 0.512 | 0.388 | 0.571 | 0.592 | 0.498 | 0.278 | 0.195 | 0.615 |
| 0.647 | 0.639 | 0.491 | 0.515 | 0.45 | 0.385 | 0.509 | 0.492 | 0.327 | 0.174 | 0.17 | 0.31 |
| 0.793 | 0.786 | 0.666 | 0.591 | 0.564 | 0.446 | 0.63 | 0.65 | 0.531 | 0.334 | 0.213 | 0.61 |
| 0.737 | 0.69 | 0.589 | 0.538 | 0.515 | 0.379 | 0.567 | 0.555 | 0.463 | 0.22 | 0.172 | 0.448 |

FST value at this locus (upper right diagonal)

| Deng9703 | Deng9847 | Chen0009 | Chen0015 | Niuma003 | Deng8939 | Deng8999 | Deng9040 | Deng9127 | Deng9569 | Deng9601 |
|---|---|---|---|---|---|---|---|---|---|---|
| 0.399 | 0.515 | 0.542 | 0.572 | 0.385 | 0.539 | 0.61 | 0.555 | 0.498 | 0.595 | 0.519 |
| 0.605 | 0.723 | 0.744 | 0.706 | 0.594 | 0.745 | 0.809 | 0.777 | 0.668 | 0.786 | 0.701 |
| 0.561 | 0.623 | 0.711 | 0.686 | 0.492 | 0.72 | 0.766 | 0.759 | 0.647 | 0.761 | 0.679 |
| 0.446 | 0.487 | 0.651 | 0.591 | 0.457 | 0.659 | 0.715 | 0.679 | 0.564 | 0.705 | 0.592 |
| 0.547 | 0.626 | 0.696 | 0.661 | 0.462 | 0.714 | 0.774 | 0.736 | 0.573 | 0.75 | 0.609 |
| 0.529 | 0.654 | 0.716 | 0.678 | 0.501 | 0.738 | 0.791 | 0.75 | 0.608 | 0.752 | 0.656 |
| 0.542 | 0.604 | 0.699 | 0.638 | 0.467 | 0.706 | 0.769 | 0.738 | 0.612 | 0.751 | 0.615 |
| 0.56 | 0.646 | 0.726 | 0.652 | 0.472 | 0.729 | 0.792 | 0.754 | 0.619 | 0.761 | 0.682 |
| 0.574 | 0.685 | 0.7 | 0.654 | 0.545 | 0.711 | 0.769 | 0.746 | 0.621 | 0.751 | 0.65 |
| 0.549 | 0.662 | 0.714 | 0.684 | 0.506 | 0.733 | 0.792 | 0.738 | 0.63 | 0.765 | 0.674 |
| 0.492 | 0.619 | 0.704 | 0.628 | 0.449 | 0.716 | 0.778 | 0.746 | 0.588 | 0.755 | 0.627 |
| 0.62 | 0.723 | 0.757 | 0.713 | 0.544 | 0.753 | 0.809 | 0.774 | 0.627 | 0.783 | 0.709 |
| 0.567 | 0.702 | 0.735 | 0.699 | 0.537 | 0.749 | 0.824 | 0.781 | 0.619 | 0.773 | 0.656 |
| 0.36 | 0.492 | 0.601 | 0.558 | 0.365 | 0.62 | 0.659 | 0.642 | 0.48 | 0.659 | 0.571 |
| 0.345 | 0.398 | 0.552 | 0.52 | 0.311 | 0.557 | 0.62 | 0.549 | 0.496 | 0.576 | 0.506 |
| 0.366 | 0.432 | 0.489 | 0.482 | 0.325 | 0.525 | 0.576 | 0.498 | 0.432 | 0.541 | 0.494 |
| 0.205 | 0.322 | 0.424 | 0.343 | 0.165 | 0.483 | 0.461 | 0.359 | 0.361 | 0.424 | 0.35 |
| 0.39 | 0.429 | 0.527 | 0.57 | 0.378 | 0.532 | 0.586 | 0.552 | 0.502 | 0.624 | 0.559 |
| 0.179 | 0.24 | 0.617 | 0.491 | 0.168 | 0.573 | 0.615 | 0.573 | 0.469 | 0.634 | 0.52 |
| -0.025 | 0.029 | 0.421 | 0.393 | 0.109 | 0.429 | 0.468 | 0.468 | 0.302 | 0.501 | 0.42 |
| 0 | 0.013 | 0.285 | 0.241 | -0.002 | 0.306 | 0.305 | 0.233 | 0.144 | 0.305 | 0.187 |
| 0.001 | 0.03 | 0.181 | 0.166 | 0.024 | 0.181 | 0.205 | 0.167 | 0.138 | 0.184 | 0.139 |
| -0.036 | 0.024 | 0.549 | 0.39 | 0.098 | 0.533 | 0.618 | 0.595 | 0.284 | 0.592 | 0.415 |
| - | 0.022 | 0.475 | 0.348 | 0.062 | 0.476 | 0.546 | 0.502 | 0.239 | 0.527 | 0.341 |
| 0.047 | - | 0.565 | 0.388 | 0.116 | 0.524 | 0.612 | 0.595 | 0.294 | 0.587 | 0.426 |
| 0.495 | 0.596 | - | 0.03 | 0.128 | 0.044 | 0.087 | 0.089 | 0.041 | 0.083 | 0.002 |
| 0.373 | 0.416 | 0.075 | - | 0.047 | -0.025 | 0.003 | 0.002 | 0.01 | -0.007 | 0.044 |

| | | | | | | | | | | |
|---|---|---|---|---|---|---|---|---|---|---|
| 0.095 | 0.179 | 0.156 | 0.077 | - | 0.076 | 0.091 | 0.02 | -0.007 | 0.038 | 0.044 |
| 0.483 | 0.54 | 0.092 | -0.015 | 0.097 | - | 0.021 | 0.027 | 0.028 | 0.007 | 0.053 |
| 0.554 | 0.625 | 0.137 | 0.021 | 0.135 | 0.031 | - | -0.003 | -0.008 | -0.009 | 0.06 |
| 0.522 | 0.613 | 0.134 | 0.016 | 0.058 | 0.04 | 0.017 | - | 0.003 | -0.022 | 0.067 |
| 0.255 | 0.314 | 0.062 | 0.028 | 0.021 | 0.041 | 0.001 | 0.016 | - | -0.013 | -0.005 |
| 0.54 | 0.605 | 0.117 | -0.005 | 0.06 | 0.016 | 0.005 | -0.015 | 0 | - | 0.07 |
| 0.359 | 0.458 | 0.017 | 0.087 | 0.072 | 0.077 | 0.1 | 0.106 | 0.014 | 0.102 | - |
