## Supporting Information_Table S5 for "Pleistocene glaciation advances the younger temporal dimension of species diversification in a major biodiversity hotspot"

**Table S5.** The paired mean absolute differentiation ($D_{XY}$) among 34 populations estimated using *populations* platform in Stacks.

| **Populations** | **Chen0018** | **Chen0019** | **Chen0021** | **Chen0022** | **LZY004005** | **Yue0719** | **Yue0720** | **ZBFC295** | **ZBFC425** | **Deng9733** |
|---|---|---|---|---|---|---|---|---|---|---|
| **Chen0018** | - | | | | | | | | | |
| **Chen0019** | 0.0022 | - | | | | | | | | |
| **Chen0021** | 0.0027 | 0.0025 | - | | | | | | | |
| **Chen0022** | 0.0023 | 0.0018 | 0.0021 | - | | | | | | |
| **LZY004005** | 0.0024 | 0.0029 | 0.0028 | 0.0015 | - | | | | | |
| **Yue0719** | 0.0022 | 0.0025 | 0.0026 | 0.0021 | 0.0028 | - | | | | |
| **Yue0720** | 0.0026 | 0.0032 | 0.0029 | 0.0016 | 0.002 | 0.0031 | - | | | |
| **ZBFC295** | 0.0027 | 0.0031 | 0.003 | 0.0016 | 0.0021 | 0.0032 | 0.0021 | - | | |
| **ZBFC425** | 0.0024 | 0.0027 | 0.0027 | 0.0016 | 0.002 | 0.0028 | 0.0017 | 0.002 | - | |
| **Deng9733** | 0.002 | 0.0021 | 0.0024 | 0.0019 | 0.0025 | 0.002 | 0.0027 | 0.003 | 0.0025 | - |
| **Deng9745** | 0.002 | 0.0022 | 0.0025 | 0.0018 | 0.0024 | 0.0022 | 0.0025 | 0.0025 | 0.0024 | 0.0021 |
| **Deng9772** | 0.0023 | 0.0025 | 0.0025 | 0.0019 | 0.0028 | 0.0025 | 0.0033 | 0.0035 | 0.0026 | 0.0019 |
| **Deng9783** | 0.0029 | 0.0038 | 0.0034 | 0.0024 | 0.003 | 0.0036 | 0.003 | 0.0034 | 0.003 | 0.0034 |
| **Deng9810** | 0.0025 | 0.0031 | 0.0028 | 0.0019 | 0.0024 | 0.0028 | 0.0024 | 0.0031 | 0.0024 | 0.0026 |
| **Deng9813** | 0.0022 | 0.0024 | 0.0025 | 0.0016 | 0.0018 | 0.0024 | 0.0019 | 0.0024 | 0.0017 | 0.0023 |
| **Deng9525** | 0.0027 | 0.0039 | 0.0036 | 0.0023 | 0.0024 | 0.0035 | 0.0023 | 0.0028 | 0.0024 | 0.0028 |
| **Deng9528** | 0.0028 | 0.0031 | 0.0028 | 0.0027 | 0.0028 | 0.003 | 0.0032 | 0.0033 | 0.0026 | 0.0025 |
| **Deng9533** | 0.002 | 0.0017 | 0.002 | 0.0013 | 0.0013 | 0.0017 | 0.0013 | 0.0015 | 0.0012 | 0.0014 |
| **Chen0016** | 0.0034 | 0.004 | 0.0039 | 0.0031 | 0.0041 | 0.0039 | 0.0041 | 0.004 | 0.0039 | 0.004 |
| **ZYZhou** | 0.0039 | 0.0048 | 0.0044 | 0.0039 | 0.0044 | 0.0046 | 0.0045 | 0.0047 | 0.0045 | 0.0046 |
| **Deng9524** | 0.0035 | 0.0043 | 0.0037 | 0.0031 | 0.0036 | 0.004 | 0.0036 | 0.0039 | 0.0039 | 0.0041 |
| **Deng9535** | 0.0038 | 0.0046 | 0.0042 | 0.0033 | 0.0037 | 0.004 | 0.0036 | 0.0042 | 0.0039 | 0.0041 |
| **Deng9715** | 0.0038 | 0.0048 | 0.0045 | 0.0036 | 0.0046 | 0.0047 | 0.0045 | 0.0048 | 0.0047 | 0.0045 |
| **Deng9703** | 0.0039 | 0.0047 | 0.0044 | 0.0037 | 0.0045 | 0.0045 | 0.0044 | 0.0047 | 0.0045 | 0.0045 |
| **Deng9847** | 0.004 | 0.005 | 0.0045 | 0.0036 | 0.0046 | 0.0048 | 0.0045 | 0.0049 | 0.0049 | 0.0047 |
| **Chen0009** | 0.0044 | 0.0053 | 0.0052 | 0.0048 | 0.005 | 0.0053 | 0.0052 | 0.0053 | 0.0052 | 0.0052 |
| **Chen0015** | 0.0046 | 0.0052 | 0.0051 | 0.0045 | 0.0049 | 0.0051 | 0.0048 | 0.005 | 0.0048 | 0.005 |
| **Niuma003** | 0.0036 | 0.0048 | 0.0041 | 0.004 | 0.0041 | 0.0045 | 0.0041 | 0.0044 | 0.0043 | 0.0042 |

| | | | | | | | | | | |
|---|---|---|---|---|---|---|---|---|---|---|
| **Deng8939** | 0.0043 | 0.0054 | 0.0052 | 0.0048 | 0.0052 | 0.0054 | 0.0051 | 0.0053 | 0.0051 | 0.0052 |
| **Deng8999** | 0.0046 | 0.0057 | 0.0055 | 0.0051 | 0.0055 | 0.0057 | 0.0054 | 0.0056 | 0.0055 | 0.0055 |
| **Deng9040** | 0.0044 | 0.0056 | 0.0055 | 0.0049 | 0.0053 | 0.0054 | 0.0053 | 0.0056 | 0.0054 | 0.0052 |
| **Deng9127** | 0.004 | 0.005 | 0.005 | 0.0043 | 0.0046 | 0.0047 | 0.0045 | 0.005 | 0.0047 | 0.0048 |
| **Deng9569** | 0.0047 | 0.0055 | 0.0054 | 0.0051 | 0.0054 | 0.0053 | 0.0053 | 0.0054 | 0.0054 | 0.0054 |
| **Deng9601** | 0.0044 | 0.0053 | 0.0051 | 0.0046 | 0.0048 | 0.0051 | 0.0047 | 0.0052 | 0.0049 | 0.0051 |

Dxy means, absolute measure (nonrelative, as opposed to FSTlike measures) of locus differentiation, measuring the number of difference

| Deng9745 | Deng9772 | Deng9783 | Deng9810 | Deng9813 | Deng9525 | Deng9528 | Deng9533 | Chen0016 | ZYZhou | Deng9524 |
|---|---|---|---|---|---|---|---|---|---|---|
| - | | | | | | | | | | |
| 0.002 | - | | | | | | | | | |
| 0.0034 | 0.0038 | - | | | | | | | | |
| 0.0028 | 0.0026 | 0.0028 | - | | | | | | | |
| 0.0022 | 0.0022 | 0.0021 | 0.0021 | - | | | | | | |
| 0.0028 | 0.0036 | 0.0024 | 0.0027 | 0.0021 | - | | | | | |
| 0.0028 | 0.0029 | 0.0029 | 0.0028 | 0.0023 | 0.0026 | - | | | | |
| 0.0015 | 0.0017 | 0.0018 | 0.0016 | 0.0013 | 0.0014 | 0.0016 | - | | | |
| 0.0039 | 0.0043 | 0.0042 | 0.0041 | 0.004 | 0.0043 | 0.0039 | 0.003 | - | | |
| 0.0044 | 0.0047 | 0.0045 | 0.0041 | 0.0038 | 0.0042 | 0.0042 | 0.0034 | 0.0031 | - | |
| 0.0037 | 0.0041 | 0.0036 | 0.0032 | 0.0031 | 0.0032 | 0.0032 | 0.0028 | 0.0022 | 0.0018 | - |
| 0.0039 | 0.0045 | 0.0035 | 0.0034 | 0.0034 | 0.0033 | 0.0032 | 0.0032 | 0.0031 | 0.0025 | 0.002 |
| 0.0044 | 0.0049 | 0.0048 | 0.0042 | 0.0039 | 0.0042 | 0.0039 | 0.0033 | 0.003 | 0.002 | 0.0018 |
| 0.0043 | 0.0049 | 0.0045 | 0.004 | 0.0039 | 0.004 | 0.0037 | 0.0034 | 0.0029 | 0.002 | 0.002 |
| 0.0046 | 0.0051 | 0.005 | 0.0043 | 0.0038 | 0.004 | 0.004 | 0.0033 | 0.0028 | 0.0017 | 0.0018 |
| 0.0052 | 0.0055 | 0.0053 | 0.0049 | 0.0044 | 0.0041 | 0.0043 | 0.0041 | 0.0046 | 0.0036 | 0.003 |
| 0.0048 | 0.0051 | 0.0052 | 0.0046 | 0.0044 | 0.0042 | 0.0038 | 0.0043 | 0.004 | 0.0036 | 0.003 |
| 0.0039 | 0.0043 | 0.0045 | 0.0039 | 0.0035 | 0.0036 | 0.0036 | 0.0031 | 0.0029 | 0.0025 | 0.0021 |

| | | | | | | | | | | |
|---|---|---|---|---|---|---|---|---|---|---|
| 0.0052 | 0.0053 | 0.0054 | 0.0049 | 0.0044 | 0.0043 | 0.0044 | 0.0042 | 0.0043 | 0.0036 | 0.0031 |
| 0.0055 | 0.0057 | 0.0058 | 0.0051 | 0.0048 | 0.0046 | 0.0045 | 0.0043 | 0.0045 | 0.0039 | 0.0031 |
| 0.0054 | 0.0055 | 0.0056 | 0.0051 | 0.0045 | 0.0043 | 0.0041 | 0.0042 | 0.0044 | 0.004 | 0.0027 |
| 0.0046 | 0.0047 | 0.0048 | 0.0041 | 0.0043 | 0.0041 | 0.004 | 0.0039 | 0.0039 | 0.0033 | 0.0026 |
| 0.0054 | 0.0055 | 0.0055 | 0.0051 | 0.0045 | 0.0044 | 0.0043 | 0.0045 | 0.0047 | 0.004 | 0.003 |
| 0.0048 | 0.0052 | 0.0049 | 0.0049 | 0.0044 | 0.0044 | 0.0044 | 0.0043 | 0.0042 | 0.004 | 0.0029 |

s between populations, excluding polymorphisms segregating within the two populations

| **Deng9535** | **Deng9715** | **Deng9703** | **Deng9847** | **Chen0009** | **Chen0015** | **Niuma003** | **Deng8939** | **Deng8999** | **Deng9040** | **Deng9127** |
|---|---|---|---|---|---|---|---|---|---|---|
| - | | | | | | | | | | |
| 0.0024 | - | | | | | | | | | |
| 0.0026 | 0.0021 | - | | | | | | | | |
| 0.0027 | 0.0018 | 0.0021 | - | | | | | | | |
| 0.0028 | 0.0043 | 0.004 | 0.0044 | - | | | | | | |
| 0.0029 | 0.0036 | 0.0035 | 0.0036 | 0.0021 | - | | | | | |
| 0.0025 | 0.0025 | 0.0029 | 0.0026 | 0.0025 | 0.0025 | - | | | | |

| | | | | | | | | | | |
|---|---|---|---|---|---|---|---|---|---|---|
| 0.0026 | 0.0042 | 0.004 | 0.0041 | 0.0019 | 0.002 | 0.0024 | - | | | |
| 0.0032 | 0.0047 | 0.0044 | 0.0046 | 0.0021 | 0.0022 | 0.0027 | 0.002 | - | | |
| 0.003 | 0.0046 | 0.0042 | 0.0046 | 0.0023 | 0.0022 | 0.0024 | 0.0021 | 0.0019 | - | |
| 0.0028 | 0.0033 | 0.0033 | 0.0034 | 0.0022 | 0.0024 | 0.0026 | 0.0022 | 0.0023 | 0.0024 | - |
| 0.0032 | 0.0044 | 0.0042 | 0.0045 | 0.0023 | 0.0022 | 0.0025 | 0.002 | 0.002 | 0.0022 | 0.0023 |
| 0.0032 | 0.004 | 0.0038 | 0.0039 | 0.0021 | 0.0024 | 0.0027 | 0.0021 | 0.0023 | 0.0024 | 0.0023 |

| Deng9569 | Deng9601 |
| --- | --- |

| | |
|---|---|
| - | |
| 0.0024 | - |
